# Parental age at reproduction accelerates offspring adult emergence with no apparent effects on reproductive success

**DOI:** 10.64898/2026.04.14.718189

**Authors:** Mark D. Pitt, Brendan O’Connor, Timothy D. Sheen, Davide M. Dominoni, Tom Tregenza, Jelle J. Boonekamp

## Abstract

Parental age at reproduction has been widely observed to induce detrimental effects on offspring adult lifespan (i.e., The Lansing Effect). In addition to The Lansing Effect, parental age might elicit a host of effects on linked life-history traits, but this has received considerably less attention. Two demographic processes could underpin a Lansing Effect, with different fitness implications: (i) Offspring from old parents may display greater mortality from conception (i.e., frailty), constraining their survival from birth. (ii) Alternatively, they may exhibit an increased age-specific mortality (i.e., accelerated senescence), with detrimental effects on survival that manifest after maturity. Our focal species, *Gryllus bimaculatus,* is often used as a model in evolutionary ecology as their survival and reproduction are highly sensitive to changes in temperature. Using a longitudinal within-parent study, we investigated how parental age influenced offspring longevity (i.e., through increased frailty or accelerated senescence), alongside its joint effects on a suite of other life-history traits. Additionally, we exposed parental pairs to one of three naturally-encountered temperatures (25.5°C, 28.0°C, or 30.5°C) and asked whether parental temperature exposure influenced the severity of the parental age effect. Contrary to our predictions, we found no evidence for The Lansing Effect, nor that parental temperature exposure (despite increasing mortality) mediated the observed parental age effects. In comparison to offspring from young parents, those from old parents exhibited accelerated development and growth rates, a smaller adult mass, and extended adult lifespans, opposite of a Lansing Effect. Moreover, there were no apparent negative effects of parental age on offspring reproduction. The increased adult lifespan in offspring from older parents did not arise from a reduced initial frailty, but from a delayed onset and slowed progression of senescence. This slowed senescence is unlikely to reflect a direct effect of parental age. Rather, it may arise indirectly through causal effects of parental age on adult size. Meanwhile, the accelerated development of offspring from old parents could be adaptive, enabling individuals to overcome seasonal time constraints. Determining whether these parental age effects accumulate across generations, or persist in the wild, will be essential in establishing their role in the evolution of ageing.

## 1. Introduction

Classical theories of ageing have implicitly assumed that senescence, the progressive increase in mortality and reduction in reproduction with increasing chronological age, is an exclusively within-individual phenomenon (Hamilton, 1966; Abrams, 1991; Monaghan *et al*., 2008; Nussey *et al*., 2013). Aside from the possible effects of reduced parental care, offspring fitness was said to be entirely independent from the parents age at conception, with these evolutionary theories excluding the possibility that the germline could age (Van Den Heuvel *et al*., 2016). Only somatic cells were thought to accumulate damage over time (e.g., DNA replication errors, accelerated telomere attrition, abnormal DNA methylation patterns, heightened mitochondrial dysfunction, and the loss of proteostasis; discussed in Ogrodnik *et al.,* 2019). Meanwhile, the germ cells were assumed to be protected through a phenomenon referred to as the “Weissmann barrier” (Weismann, 1893; Kirkwood & Cremer, 1982). Conceptually, this barrier was thought to function to preserve the integrity of the germline across generations. The existence of the Weismann barrier has faced increasing scrutiny (Monaghan & Metcalfe, 2019; Sciamanna *et al*., 2019; Bline *et al*., 2020), with evidence now suggesting that even though germline DNA is offered some special protection (Strome & Updike, 2015), it is not invulnerable to senescence (Keefe & Liu, 2008; Yamada-Fukunaga *et al*., 2013; Maklakov & Immler, 2016; Chieffi Baccari *et al*., 2023). Thus, the consensus is that the effects of senescence *can* be transmitted intergenerationally through the germ cells (Monoghan & Metcalfe, 2019 and references therein), affecting offspring fitness. It is crucial to understand the importance of these parental age effects in the evolution of senescence, as their role has previously been understated (Moorad & Nussey, 2016; Van Den Heuvel *et al*., 2016; Barks & Laird, 2020; Hernández *et al*., 2020).

Evidence for negative effects of parental age on offspring traits, hereafter ‘*parental effect senescence’* (Moorad & Nussey, 2016), continues to accumulate from both captive and wild animal populations (Ivimey-Cook & Moorad, 2020; Ivimey-Cook *et al*., 2023). These effects have been most widely reported across invertebrates, including in Rotifers (e.g., *Philodina citrina*: Lansing, 1947; *Brachionus manjavacas:* Gribble *et al*., 2014; Bock *et al*., 2019; *Brachionus plicatilis:* Liguori *et al*., 2024), Nematodes (*Caenorhabditis elegans*: Klass, 1977; Lenzi *et al*., 2023; *Heligmosomoides polygyrus:* Lippens *et al*., 2017), Fruit flies *(Drosophila melanogaster:* Kern *et al*., 2001; Priest *et al*., 2002; Bloch Qazi *et al*., 2017), Water fleas *(Daphnia pulex:* Plaistow *et al*., 2015; *Daphnia magna:* Anderson *et al*., 2022), Crickets *(Gryllus bimaculatus:* Noguera, 2021), Lepidopterans (*Pieris brassicae*: Ducatez *et al*., 2012), and Copepods (*Cyclops kolensis*: Jamieson & Santer, 2003). Although less frequently reported, parental effect senescence has been observed in some free-living mammal populations, including red squirrels (*Tamiasciurus hudsonicus*: Descamps *et al*., 2008), Asian elephants (*Elephas maximus*: Reichert *et al*., 2020), Soay sheep (*Ovis aries*: Ravindran *et al*., 2025), and even pre-industrial humans (Gillespie *et al*., 2013). Some of the earliest evidence for this phenomenon comes from Lansing (1947), who demonstrated that old-aged selection lines of the rotifer *P. citrina* produced offspring with abbreviated adult lifespans, with this effect propagating through subsequent generations. These effects on offspring adult lifespan are among one of the most well-known manifestations of parental age, widely referred to as *‘The Lansing Effect’* (Comfort, 1953). However, Lansing’s initial findings were subsequently questioned (Comfort, 1953; Lints & Hoste, 1974; King, 1983), owing to issues with the experimental design that inadvertently created selection on reproductive scheduling, independent of an effect of parental age at reproduction *per se* (King, 1983). More recently, a meta-analysis investigating the general tendency for The Lansing Effect in 15 species by Ivemy-Cook *et al*. (2023) demonstrated that a one-unit increase in maternal age was associated with a 17 to 22% reduction in the offspring’s adult lifespan, on average. Yet, this result is mainly driven by laboratory invertebrate studies in the existing literature. Disentangling the effects of taxonomy from the study environment remains difficult (Promislow *et al*., 2022). Most studies on short-lived invertebrates have been conducted in controlled laboratory environments whereas the applicability of The Lansing Effect to natural insect populations, where extrinsic mortality is high, remains uncertain (Zajitschek *et al*., 2020; Angell *et al*., 2022; Tregenza *et al*., 2026). Alongside the offspring’s adult lifespan, detrimental effects of advanced parental age have been observed on embryo viability (e.g., in *D. melanogaster*: Bloch Qazi *et al*., 2017; *G. bimaculatus:* Noguera, 2021), pre-adult survival (e.g., in *D. melanogaster*: Kern *et al.,* 2001; Red-billed choughs (*Pyrrhocorax pyrrhocorax)*: Reid *et al.,* 2010; Blue-footed boobies (*Sula nebouxii)*: Torres *et al*., 2011), development time and growth rate (e.g., in Seed beetles (*Callosobruchus maculatus*): Fox & Dingle, 1994; Lind *et al*. 2015; Zig-zag ladybirds (*Cheilomenes sexmaculata*): Singh & Omkar, 2009), the size at birth and maturity (e.g., in Rice weevils (*Sitophilus oryzae*): Opit & Throne, 2007; Mosquitofish (*Gambusia holbrooki*): Vega – Trejo *et al*., 2018), and reproductive performance (e.g., Three-spined sticklebacks (*Gasterosteus aculeatus*): Kim *et al*., 2026; *Drosophila serrata*: Hercus & Hoffmann, 2000; House sparrows (*Passer domesticus*): Schroeder *et al.,* 2015; Rotifers (*B. manjavacas*): Van Daalen *et al*., 2022). Despite this prevailing evidence, there is a growing uncertainty over the ubiquity of parental effect senescence; studies frequently find conflicting results between laboratory or genetic strains (Priest *et al*., 2002; Plaistow *et al*., 2015; Anderson *et al*., 2022; Liguori *et al*., 2024), including neutral (Ivimey-Cook & Moorad, 2018) or positive parental age effects (Plaistow *et al*., 2007; Perez *et al*., 2017; Ivimey-Cook & Moorad, 2020; Travers *et al*., 2021), even on adult lifespan (an *‘Inverse Lansing Effect’*; Marshall *et al*., 2010; Anderson *et al*., 2022). Furthermore, parental effect senescence often incorporates sex-specific interactions, with increasing maternal and paternal age shown to have differing impacts on the fitness of daughters and sons (Priest *et al*., 2002; Ducatez *et al*., 2012; Bouwhuis *et al*., 2015; Fay *et al*., 2016; Noguera, 2021). This lack of consistency raises challenges in identifying the physiological pathways, biological factors, and environments that are important for parental effect senescence (Priest *et al*., 2002; Monaghan *et al*., 2020).

Despite an extensive history of research, there are numerous issues with previous study designs that could have biased or masked the observed parental age effect. First, most studies appear to follow a typical cross-sectional comparison of young and old parents (see: Priest *et al*., 2002; Plaistow *et al*., 2015; Bloch Qazi *et al*., 2017; Bock *et al*., 2019; Wylde *et al*., 2019; Angell *et al*., 2022), rather than longitudinally tracking changes to offspring quality across an individual’s lifespan. Such cross-sectional studies cannot control the differential survival of parental phenotypes (e.g., *selective disappearance’*: Van De Pol & Verhulst, 2006; Nussey *et al*., 2008; Van De Pol & Wright, 2009; Ivimey-Cook & Moorad, 2018). For example, the selective disappearance of low-quality or high early-life investment parental phenotypes may lead to an overrepresentation of high-quality parents, or those adopting a low reproductive investment strategy, in the old-aged cohort (Monaghan *et al*., 2020). At the population level, this then artificially creates correlations between individual quality, reproductive investment, and age, masking the true within-individual changes to offspring quality associated with senescence (Van Noordwijk & De Jong, 1986; Van De Pol & Verhulst, 2006).

Second, studies into parental age effects are often correlational, describing general associations between parental age and offspring traits. There is a growing necessity to tease apart the causal effects of parental senescence (both genetic and environmental) on offspring performance (Kern *et al*., 2001; Bock *et al*., 2019). Previous work has demonstrated that parental effect senescence can exhibit genotypic heterogeneity, differing between strains artificially selected for early-or late-life performance (Kern *et al*., 2001; Priest *et al*., 2002; Yılmaz *et al*., 2008; Anderson *et al*., 2022; Liguori *et al*., 2024). It is increasingly recognised that environmental variation can also govern the severity of parental effect senescence, with non-additive effects on offspring phenotypes (Benton *et al*., 2008; Ducatez *et al*., 2012; Cooper & Kruuk, 2018; Cope *et al*., 2022). For instance, resource limited or physiologically stressful environments could either ameliorate or reinforce the deleterious effects of senescence on survival and reproduction (Gibbs *et al*., 2010; Van Den Heuvel *et al*., 2016; Lemaître & Gaillard, 2017), so that any adverse effects of parental age on offspring fitness are context dependent (Fox & Dingle, 1994). For example, Gribble *et al*. (2014) demonstrated that in the rotifer *B. manjavacas*, maternal caloric restriction not only reduced the mother’s mortality risk, but also partially rescued the detrimental effects of advanced maternal age on the longevity of female offspring. Conversely, environments that increase the parents’ mortality risk may exacerbate any negative effects of parental age (Gibbs *et al*., 2010). The role of temperature in shaping parental effect senescence has received limited attention, despite the known importance of temperature in modulating the lifespan of ectothermic organisms, who lack physiological means of thermoregulation (Deutsch *et al*., 2008; Kingsolver *et al*., 2013; González – Tokman *et al*., 2020). The routes through which temperature governs longevity in ectothermic insects are complex, with chronic exposure to high temperatures either raising their baseline mortality (i.e., temperature induced frailty; Vaupel *et al*., 1979; Vindenes *et al*., 2025) or accelerating actuarial senescence (Flouris & Piantoni, 2015; Burraco *et al*., 2020). These routes could have different implications for offspring fitness (i.e., the offspring’s total genetic contribution to future generations; Wadgymar *et al*., 2024), especially if only senescent damage can be inherited by the offspring. Sublethal thermal stress has also been increasingly shown to reduce gamete quality in both sexes (Reinhardt *et al*., 2015; Gasparini *et al*., 2018; Chevrier *et al*., 2019; Walsh *et al*., 2019; Zhou *et al*., 2025; Snook *et al*., 2026), alongside their pre– and post-copulatory behaviours (Abram *et al*., 2017; Baur *et al*., 2024; Grandela *et al*., 2024), well before any detrimental effects are observed on survival. For example, *G. bimaculatus* males briefly exposed to sublethal heatwave temperatures displayed lower fertility relative to those maintained under standard rearing conditions, despite no observable difference in their lifetime survival (Ratz *et al*., 2024). The broader effects of temperature on parental effect senescence remain unclear and thus deserve more attention.

Third, few have considered how the variability of offspring phenotypes changes with parental age. Parental effect senescence may not only govern the offspring’s average trait value, but also generate greater variation in offspring phenotypes, as has been similarly observed for other parental environmental stressors (e.g., increasing temperature; De Jong, 1995; O’Dea *et al*., 2016, 2019). Any age-dependent change to the variability of offspring phenotypes may be adaptive (i.e., bet-hedging; Crean & Marshall, 2009) or reflect physiological constraints (Debat & David, 2001; Olsson & Uller, 2002), such as the senescent instability of the parents’ reproductive systems (Monaghan & Metcalfe, 2019).

Fourth, despite the evolutionary importance of parental age effects in understanding senescence, most studies have tested for their isolated effects on offspring longevity – the end point of senescence (Promislow *et al*., 2022). However, this precludes any effects of parental age on offspring age-specific mortality (i.e., *how* the offspring senesce) and reproductive performance. Assessing the joint effects of parental age on offspring mortality and reproduction is a necessary step in establishing the fitness, demographic, and evolutionary consequences of parental effect senescence (Kirkpatrick & Lande, 1989; Nussey *et al*., 2013; Bloch Qazi *et al*., 2017; Monaghan & Metcalfe, 2019).

In their review, Monaghan *et al*. (2020) identified that increasing parental age could reduce offspring longevity through two non-exclusive demographic processes. First, offspring from old-age parents may be conceived with an increased frailty (Vaupel *et al*., 1979). Here, their mortality is higher from conception as they initiate life with either the senescent damage accumulated by their aged parents (Díaz & Esponda, 2004; Kong *et al*., 2012; Monaghan & Metcalfe, 2019; Dutt & Laird, 2024), or experience age-dependent reductions in either pre– or post-natal parental provisioning (Bouwhuis *et al*., 2015; Lemaître & Gaillard, 2017; Muller *et al*., 2017; Ivimey-Cook & Moorad, 2018). This increased frailty could persist, but does not necessarily worsen, as the offspring age. Alternatively, offspring from old-aged parents may exhibit increased age-specific mortality (i.e., accelerated senescence), with the negative effects of the accumulated senescent damage manifesting after maturity (Monaghan *et al*., 2020. Despite the breadth of studies demonstrating parental effect senescence, few have assessed its demographic underpinnings. Indeed, those that have reported conflicting results, with either increased frailty (Dutt & Laird, 2024), accelerated senescence (Plaistow *et al*., 2015) or both (Bock *et al*., 2019), being implicated in reducing offspring longevity. Although both demographic processes reduce survival, they may have different fitness consequences. If offspring exhibit increased frailty, then their fitness is likely constrained from birth. Conversely, if parental age accelerates offspring senescence, but has no effect on frailty, then offspring may undertake a phenotypically plastic switch to a fast pace-of-life strategy. In this latter scenario, offspring may accelerate development to reach reproductive maturity earlier in life, increasing investment in their early life fitness contributions prior to their hastened onset of senescence (King, 1983; Bouwhuis *et al*., 2010; Monaghan *et al*., 2020). Whether these shifts in offspring life history represent an adaptive strategy to overcome the inheritance of senescent damage remains unclear. Older mothers may increase their provisioning (e.g., lipids, carbohydrates, metabolites, organelles) to offspring conceived later in life, especially if this enables these offspring to remain competitive against their siblings that were conceived earlier in the year (Plaistow *et al*., 2015; Travers *et al*., 2021; Liguori *et al*., 2024). Offspring from old parents may then accelerate their growth rates and increase investment towards early-life reproduction at the expense of reducing their longevity (Metcalfe & Monaghan, 2003; Takeshita, 2024), thereby culminating in a Lansing Effect. Regardless of the mechanism, increasing parental age may have no effect on offspring fitness. Rather, the offspring produced at the extreme ends of the parent’s lifespan could differ in their life-histories (Van Noordwijk & De Jong, 1986; Travers *et al*., 2021; Anderson *et al*., 2022). These are important questions that merit further investigation.

Using the two-spotted cricket (*Gryllus bimaculatus*), and an experimental, longitudinal within-individual study design, we investigated how parental age affected offspring mortality trajectories and life-history traits. This species is multivoltine and undergoes uninterrupted development, although the reproduction of populations in Mediterranean Europe is limited to the summer months (Ferreira and Ferguson, 2010). *G. bimaculatus* is an ideal system to study the influence of temperature exposure on parental effect senescence for several reasons. First, crickets (Orthoptera) have long been studied to answer several key questions in evolutionary ecology, including on the forces that influence sexual selection (Gray & Cade, 1999; Bailey & Zuk, 2008; Simmons and Beveridge, 2010), body size variation (Simmons, 1988; Mousseau & Roff, 1989; Carrière *et al*., 1996; Whitman, 2008), and voltinism (Bradford & Roff, 1995; Matsuda *et al*., 2018). More recently, there exists a growing recognition that senescence forms an integral part of the life-histories of both captive (Zajitschek *et al*., 2009, 2012) and wild (Rodríguez-Muñoz *et al*., 2019a, 2019b, 2019c; Zajitschek *et al*., 2020) cricket populations. For example, natural populations of *Gryllus campestris* have been demonstrated to display both actuarial and reproductive senescence, although both exhibit substantial interannual variation (Rodríguez-Muñoz *et al*. 2019b). Second, the copulatory behavior of *G. bimaculatus* is highly tractable and their reproductive biology has been well described and is likely representative of many Orthoptera species (Simmons, 1988; Simmons & Beveridge, 2010; Horch *et al*., 2017). They also display no post-natal care, with any parental investment ceasing after oviposition (Zuk & Simmons, 1997). Thus, any direct effects of parental age (e.g., on gamete quality) are unlikely to be confounded by indirect effects arising through age-dependent changes to post-natal care quality (Ivimey-Cook & Moorad, 2018). Moreover, field crickets are highly sensitive to temperature fluctuations which affect their development, physiology, survival, and reproduction (Hoffmann, 1974; Nespolo *et al*., 2003; Gasparini *et al*., 2018; Odhiambo *et al*., 2022). Crucially, *G. bimaculatus* has been previously shown to display negative effects of parental age on offspring traits, including longevity (Noguera, 2021). However, this was a cross-sectional study in which group housed offspring of old and young parents were compared, rather than through any direct impact of *within* parent senescence on individually housed offspring unaffected by group competition effects.

In this study, we first addressed a key conceptual issue: does experimentally induced environmental heterogeneity in parent mortality mediate the expression of parental effect senescence? To this end, we manipulated the temperature both parent animals experienced over their adult lifespan (either 25.5°C, 28.0°C, or 30.5°C), which created variation in parent longevity. These treatments encompass the thermal range naturally encountered by our study population (Ratz *et al*., 2024). We then asked whether parental effect senescence manifested differently depending on the parents’ temperature exposure, predicting that any adverse parental age effects would be exacerbated under the highest, and most stressful, temperature treatment. We reared all offspring under a common environment (28.0°C), to confine our inferences of temperature variation to the parent generation. Second, we sought to disentangle which demographic process (frailty or age-specific mortality) was implicated in reducing offspring longevity. Finally, we determined if a suite of other life-history traits were affected by increasing parental age, to test for a possible switch to a fast pace-of-life strategy. We predicted an increase in egg size, no effect on pre-adult survival, a decrease in post-natal development time and adult size, an increase in growth rate, and an improvement to the offspring’s own reproductive performance.

## 2. Methods

### 2.1 Study species and laboratory conditions

*G. bimaculatus* De Geer 1773 is found over a large geographic range, being native to the arid and semi-arid regions of Africa, Asia, and southern Europe (Ferreira & Ferguson, 2010). Males fight over access to high-quality territories (i.e., cracks or burrows in the ground) and will repeatedly sing from these territories to attract and court passing females (Adamo & Hoy, 1995; Tachon *et al*., 1999). During mating, the female mounts the male to enable genital coupling, before the male transfers his spermatophore (i.e., the protein capsule filled with egg-laying stimulants and spermatozoa), and females can repeatedly mate throughout life (Simmons, 1986, 1987c). After copulation, females will continuously oviposit thousands of eggs in damp soil (Lorenz, 2007). These eggs require warmth and moisture to develop, and the survival of early-instar nymphs is constrained below 16.6°C (Ferreira & Ferguson, 2010; Magara *et al*., 2024). Once oviposited, the eggs typically take 10-12 days to hatch, with the nymphs then passing through eight to 10 instars before adult eclosion (Behrens *et al*., 1983; Donoughe & Extavour, 2016). Although growth and survival are dictated by climactic conditions (Magara *et al*., 2024), those *G. bimaculatus* populations maintained under group-housed laboratory conditions typically take five to 11 weeks to reach adult maturity (Rantala & Roff, 2005), and spend a further seven to eight weeks as a reproductively active adult (Behrens *et al*., 1983).

We conducted the study outlined below from September 2023 to September 2024, using an outbred laboratory-reared population of *G. bimaculatus*, located at the Scottish Centre for Ecology and the Natural Environment (SCENE), University of Glasgow. These animals were sourced from the University of Munich in spring of 2023 and originated from a population of wild-caught animals in Seville (Spain) captured during the summer of 2021 (see Ratz *et al*. 2024 for further details on the wild population). All animals were individually housed in plastic containers (75H × 125W × 185Lmm for the parent generation, 75H × 108W × 108Lmm for the offspring generation), containing *ad-libitum* rabbit pellets (Supreme Petfoods Science Selective Adult Rabbit food), a water vial with a soaked cotton stopper, and an egg carton for shelter. We individually housed animals to remove any effects that intra-specific competition may have on their life-history traits (Simmons, 1987), and to limit the selective disappearance of low-quality individuals from our study population. Individuals were kept in incubators set to a 14:10 hour day: night cycle with a relative humidity (RH) of ∼50%. We continuously monitored temperature and relative humidity through the experimental period with loggers (Elitech RC 5+, RS-191A Temperature Loggers + DOOMAY mini hygrometer). We rotated all individuals amongst the incubators weekly, ensuring animals were alternated between incubators and shelve locations.

### 2.2 Creating the parent generation

The parent generation used in the experiment descended from 12 pairs of individually housed colony animals. We continuously maintained these colony animals under the standard rearing temperature (28.0°C) that was previously used for the source population at the University of Munich (Ratz *et al*., 2024). Due to constraints on the availability of colony pairs needed to avoid inbreeding, we could not mate individuals immediately on reproductive maturity. Colony pairs had an adult age (mean ± standard deviation (SD)) of 2.42 ± 1.51 weeks at first mating. These animals were mated a single time, after which we collected eggs from the colony mothers every other week. The average age (mean ± SD) of the colony animals across all sampling points from which we harvested the parent generation was 3.95 ± 2.18 weeks. We collected the eggs ad-hoc from the oviposition substrate (damp cotton) before these were plated and incubated in a climate room for nine days at 28.5°C (14:10 hour day: night cycle; RH of 40-60%), after which we initiated daily hatch checks. Once hatched, the experimental parent generation were housed individually in incubators (set temperature 25.0-26.5°C; equally experienced by all offspring). These cooler temperatures were used to delay their adult emergence to the start of our experimental period (January 2024).

From January 5^th^ 2024, we monitored the late-stage instars daily to record their exact date of adult eclosion. As adults can be damaged shortly after eclosing, we waited at least 24 hours after observed emergence to ensure their cuticle had hardened before measuring their mass (to the nearest 0.0001g) using a VWR SMG4i precision balance (readability ± 0.01mg). We photographed all individuals (Panasonic Lumix DC-FZ82 camera + Raynox DCR-250 macro lens (focal length: 60mm) + set to *“autofocus”* mode) to measure pronotum width and length (mm). To prevent individuals from moving during photography, we placed them in a petri dish (Greiner 6-point 60mm x 15mm) filled with cotton pads, with a scale attached (10mm strip of 1 x 1 mm graph paper). We measured pronotum width and length (in mm) from the images with the line tool in IMAGEJ (Schneider *et al*., 2012). Morphological measurements (both photography and mass) were taken between 13:00 – 14:30, with this remaining consistent throughout the experiment. Images were analysed by a single observer (Mark D. Pitt; MDP) to eliminate any between-observer measurement bias. Pronotum width and length are assumed to be more fixed indicators of structural size than adult mass, as daily variation in hydration, satiation, and oviposition can cause mass to have low within-individual repeatability (Gershman *et al*., 2022. However, we observed reasonably tight correlations between all three measures of adult body size (pronotum width ∼ pronotum length, Pearson’s correlation coefficient *r* [95% Credible Intervals (CI)] = 0.856 [0.760, 0.916]; pronotum width ∼ adult body mass, Pearson’s correlation coefficient *r* [95% CI] = 0.758 [0.610, 0.855]; pronotum length ∼ adult body mass, Pearson’s correlation coefficient *r* [95% CI] = 0.808 [0.685, 0.886], using a subset of photographs, N = 51 animals). Therefore, we selected adult body mass as our only proxy of adult size, assuming this measurement captured relevant biological variation (e.g., the time since last feeding and drinking). Survival was monitored weekly, at which point we also replaced food, water, and cleaned individual housing.

### 2.3 Parent mating design and temperature treatments

An overview of our experimental design is depicted in *Figure 1*. We paired parent animals pseudo-randomly so that inbreeding was minimised and that the members of a pair were age matched, with maternal and paternal age being tightly correlated (Pearson’s correlation coefficient *r* [95% CI] = 0.98 [0.97, 0.98]). As such, our study does not control the separate contribution of paternal and maternal age effects, which could have differing effects on the fitness of daughters and sons, as has been previously demonstrated in *G. bimaculatus* (Noguera, 2021). Parents were first mated in their second week of adult life (mean adult age (± SD) = 1.4 ± 0.67 weeks). This allowed time for the oocytes in the females to mature, which usually occurs in the initial 10 days after adult eclosion (Lorenz, 2007). During the first mating round, the parent pair were placed into a plastic container (75H × 125W × 185Lmm) and allowed to interact for 15 minutes. We watched the pair continuously and assumed copulation was successful after visual confirmation of spermatophore transfer. If this first mating was unsuccessful, we provided the pair with a further two opportunities to mate on the same day. If the third attempt was unsuccessful, both individuals were given the opportunity to mate with a different partner, who had also previously been unsuccessful, on the following day. Any pairs that failed to mate with their second partner were excluded from the study. Out of the 132 initial pairs, 120 successfully mated with their first partner, eight pairs mated with their second partner, and four pairs unsuccessfully mated and were removed from the study.

**Figure 1.**
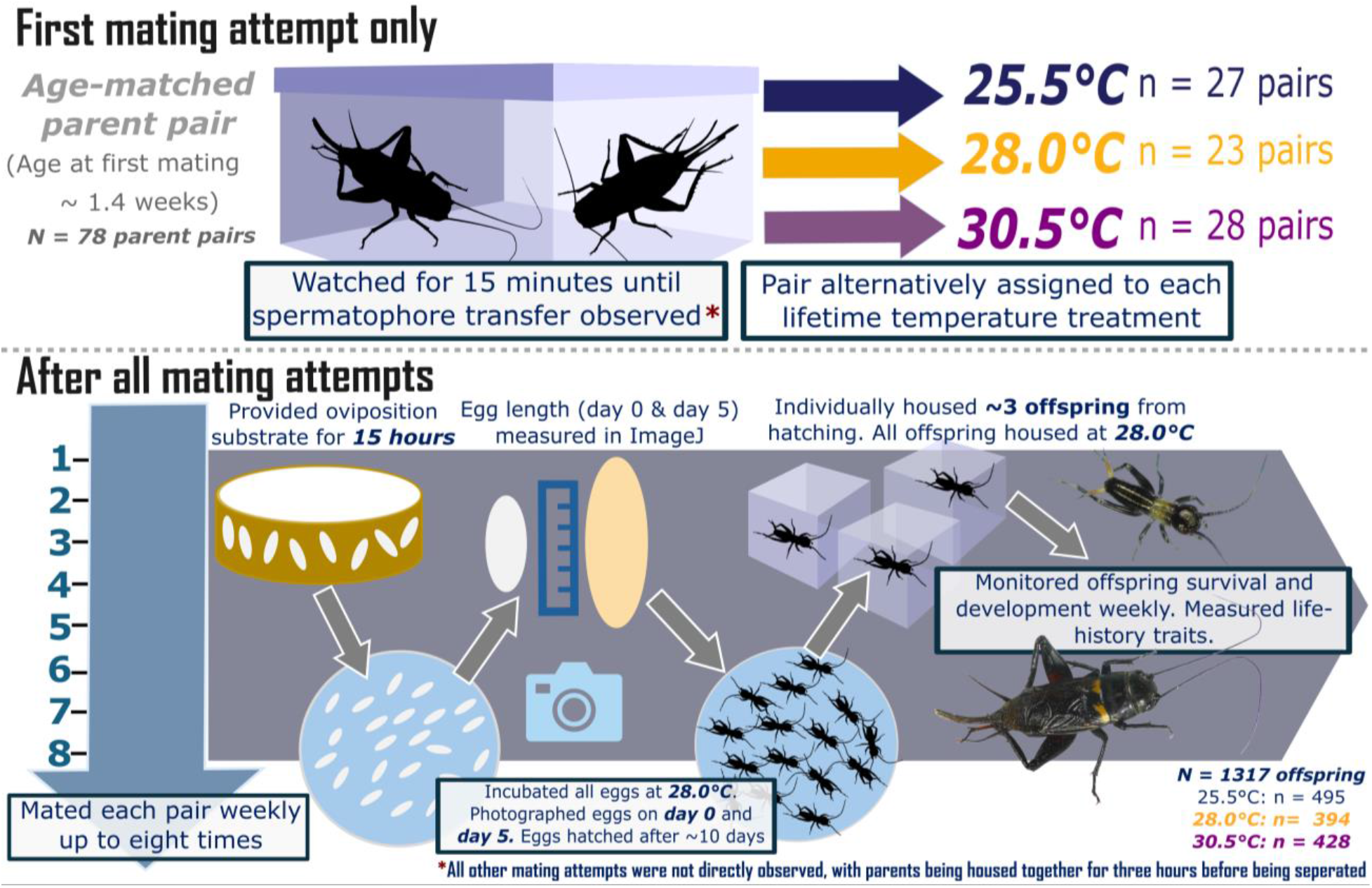
Overview of the experimental design, describing the procedure for the staged matings and the creation of the offspring generation. After the first successful mating, the pair were immediately assigned to be continuously maintained under one of three temperature treatments. Each pair was mated weekly (at their set temperatures), up to eight times in total. After each mating attempt, we collected and photographed the eggs (on the day of laying and five days into incubation). We individually housed up to three offspring per parent pair from each mating attempt. Offspring lifetime development and survival were monitored weekly, and we measured their life-history traits as discussed below. Photographs taken by Mark Pitt, and schematic designed in Inkscape.

After the first successful mating, pairs were immediately moved from their rearing temperatures and alternatively assigned to be continuously maintained under one of the three temperature treatments (25.5°C, 28.0°C, or 30.5°C). Previous experimental work has revealed that the survival of *Gryllus spp.* is sensitive to experimental temperature manipulations, both constant and fluctuating, over their adult lifespan (Centeno Filho *et al*., 2023). Additionally, male gamete quality and courtship behavior are negatively affected by heatwave temperatures (39°C) in this species (Ratz *et al*., 2024). Finally, the average summer temperature experienced by the wild population in Seville ranges from 24.5°C to 28.4°C, with maximum temperatures rarely exceeding 30.6°C – 35.0°C (Ratz *et al*., 2024). Although the critical temperature limit (*CT_max_*) for *G. bimaculatus* is unknown, the optimal temperatures for development, growth, and survival range anywhere from 27°C – 34°C, depending on locality (Behrens *et al*., 1983). We assumed that our selected temperatures (25.5°C, 28.0°C, or 30.5°C) captured the range naturally experienced by the wild population during the summer (i.e., not excessively high or low as to induce heat or cold shock, which could completely impair reproduction; Tattersall *et al*., 2012), with 30.5°C representing the most stressful conditions. Our experimental temperature manipulations created variation in parent longevity, with the Cox proportional hazards model revealing that increasing temperature reduced parent survival (*Likelihood ratio test*: *X^2^ =* 31.27*, p <* 0.005*; Appendix S1: Section S1.1*). Relative to those parents at 25.5°C, those maintained at 28.0°C or 30.5°C both exhibited an increased mortality risk (*25.5°C vs. 28.0°C*: *Hazard Ratio* (*HR [95% Confidence Intervals (CI)]* = 3.65 [1.53, 8.67], *p =* 0.003*; 25.5°C vs. 30.5°C: HR =* 7.37, 95% CI [3.31, 16.43], *p* < 0.005). Parent mortality was also higher at 30.5°C relative to 28.0°C (*28.0°C vs. 30.5°C: HR =* 2.02, 95% CI [0.32, 2.20], *p =* 0.03).

We performed the first round of matings (i.e., the pair formation round) at room temperature (24°C ± 1.5°C). Parents were then repeatedly mated on a weekly basis, up to a maximum of eight times, which ensured that we had representative sample of offspring from across the parents’ entire reproductive lifespan (Behrens *et al*., 1983). After the first mating, all other mating attempts in the sequence occurred under each pair’s specific temperature treatment and were not directly observed. Instead, pairs were housed together for three hours, before being separated and housed individually between each mating attempt. We repeatedly mated the parents to reduce the likelihood that stored sperm senescence confounded our results, as this could artificially create a parental age effect (Ribou & Reinhardt, 2012; Monaghan *et al*., 2020.

Of the 128 pairs that initially mated, only 78 pairs produced viable eggs (i.e., that showed signs of development and hatched) from this first mating attempt. To avoid wasted sampling effort repeatedly mating infertile parents, only the 78 pairs that produced viable eggs from their first mating attempt remained in the study. This selection criteria could have removed low-quality animals, or those that successfully produced viable eggs at a later attempt. However, experimentally induced selective disappearance is likely to be minimal; many parent pairs who entered the study stopped producing viable eggs well before the end of the sampling period, with only 36 of the initial 78 pairs generating offspring from their eighth mating attempt (*Appendix S1: Tables S1 & S2* for further details on sample size). For four pairs, we observed that one member of the pair had died after a staged mating, though we were unable to identify if this was a natural death or cannibalism. Due to capacity constraints, those parents who no longer produced viable eggs (i.e., following two mating attempts with no successful hatchlings; n = 13 pairs), and parents still alive by March 26^th^, 2024 (i.e., after the end of the sampling period; n = 34 pairs) were moved from their temperature treatment to a climate room (28.5°C; 14:10 hour day: night cycle; RH of 40-60%; for further details, see *Appendix S1: Section S1.1*). All offspring that these pairs produced up until their point of removal remained in our study. Although increasing temperature affected parent survival, it did not impact the parents’ number of successful mating attempts (i.e., mating attempts that resulted in successfully hatched offspring; *Appendix S1: Section S1.2*). Our final sample size consists of 78 parent pairs, with the number of pairs in each temperature treatment as follows: 27 pairs at 25.5°C, 23 pairs at 28.0°C, and 28 pairs at 30.5°C (for further details, see *Appendix S1: Tables S1 & S2*).

### 2.4 Egg length measurements and creating the offspring generation

After each staged mating, we allowed females 15 hours to lay, providing them with a petri dish (35mm x 10mm) filled with 20.0 ± 1.11 grams (mean ± SD) of wet sand (Children’s Play Sand; grain size ≤ 250 µm), covered with a damp cotton pad for laying. *G. bimaculatus* can lay eggs continuously when provided with an oviposition substrate, producing many thousands of eggs over their reproductive lifespan (Lorenz, 2007). As we could not continually siphon eggs during the study, we limited the window over which the female could lay. Here, we only provided sand for three consecutive days after each mating attempt, with fresh sand provisioned on each day. Sand was given to females on each day at 17:00, and eggs were collected the following morning at 09:00 (*Appendix S1: Section S2* for further details on egg collection). Due to capacity constraints and time limitations, we could only collect eggs from the first and third batches of sand. Eggs from each batch were photographed as a group (Panasonic LUMIX G7, 12-60mm lens; to estimate fecundity for another study) before being incubated on a petri dish (60mm x 15mm; filled with damp cotton) in the climate room. To measure egg length (in mm; for further details, see *Appendix S1: Section S2*), we randomly selected five eggs from each mating attempt to be measured on the day of laying (day zero) and again five days into incubation (day five, marking the point in embryonic development where egg size peaks; Donoughe & Extavour, 2016). Crucially, we did not obtain repeated measures from the same egg on both days, and only viable eggs showing signs of development were measured on day five (further details in *Appendix S1: Section S2*). In total, 4216 eggs across 461 egg batches were measured from 77 parent pairs, with 2285 eggs measured on day zero and a further 1931 eggs measured on day five. Egg length measurements were conducted in IMAGEJ by a single observer (Timothy Sheen; TS) to reduce observer bias. Egg batches were inspected daily during incubation and, if hatchlings were observed, they were photographed as a group (to estimate hatching success; for another study) before being housed individually for this study (*details below*). We defined a successful hatching as any nymph that had completely removed itself from its egg casing. Any unhatched eggs were returned to the climate room. Hatching checks then continued daily until all the remaining eggs on a dish had hatched or up to 14-days after first incubation, after which we assumed the remaining unhatched eggs were non-viable.

From each mating attempt, we individually housed up to three offspring (mean ± SD = 2.70 ± 0.54) per parent pair to create the offspring generation for this study. When tallied across all mating attempts, we housed a total (mean ± SD) of 16.91 ± 5.91 offspring per parent pair (*Appendix S1: Tables S1 & S2*). Due to capacity constraints, all offspring were maintained at a single intermediate temperature (28.0°C) from hatching, irrespective of their parents’ temperature treatment. This temperature is optimal for survival and growth (Behrens *et al*., 1983; Ratz *et al*., 2024), and reflects the standard rearing conditions used in the source population at the University of Munich (Ratz *et al*., 2024). We held all offspring at this *‘optimal*’ rearing temperature to isolate any effects of temperature variation to the parent generation. Although this created a difference in environments between the parent and offspring generation for the 25.5°C and 30.5°C treatments, there were no effects of the temperature parents experienced on any measured life-history trait (*see results*).

Offspring were inspected weekly for adult emergence or mortality. As per the parent generation, we waited at least 24 hours after adult emergence to measure adult mass. Overall, our sample size consists of 1317 individually housed offspring from 78 parent pairs, while complete life-history data was collected from 1309 offspring. Eight offspring had uncertainty surrounding their adult emergence dates and were excluded from analyses on offspring development time, adult lifespan, and adult mass. A further two animals had missing mass measurements and were thus removed from any size or growth rate analyses.

### 2.5 Offspring reproductive performance

Once the offspring generation reached adulthood, we mated 226 of these individuals (forming 113 offspring pairs) to determine how parental age affected offspring reproductive success. As we mated the offspring within the levels of parental age and parental temperature, we were severely limited by how many unrelated offspring could be coupled. Thus, to increase the diversity of offspring pairs in each temperature treatment, we binned parental age into two broad levels containing a similar number of individuals; offspring from either “early” or “late” aged parents. Offspring from the “early” cohort (n = 58 offspring pairs from 37 parent pairs) descended from the first two parental mating attempts, where the average adult age (mean ± SD) of the parents was 2.06 ± 1.03 weeks. Meanwhile, the “late” cohort of offspring descended from parental mating attempts five through to eight (n = 55 offspring pairs from 33 parent pairs), where the parents average adult age was 7.28 ± 1.28 weeks. The “late” cohort encompassed a broader range of mating attempts to increase the number of offspring that could be paired, as fewer parent pairs successfully produced offspring at these later mating attempts (*Appendix S1: Table S2*).

We collected and incubated offspring eggs from a single mating attempt, at which offspring had an average adult age of 4.60 ± 1.44 weeks (mean ± SD), assuming this window represented an intermediate adult age (e.g., neither young nor old). We selected this intermediate age as it just proceeded the onset of senescence but occurred after the peak age of reproduction (Lorenz, 2007). Therefore, this intermediate age may increase our likelihood of detecting a parental age effect while removing confounding effects arising from the offspring’s own age at reproduction. This was especially important since we were only able to mate offspring a single time to estimate their reproductive performance.

We performed the staged matings, egg collection, photography, incubation, and hatching checks as per the parent generation. Rather than allowing females to lay over three days, due to time limitations, we shortened the egg collection window to two days. Many of the offspring pairs did not successfully mate, with no visual evidence of spermatophore transfer. Any pairs that did not successfully mate were removed from the study. In the “early” aged cohort offspring displayed an 81.03% chance of mating success, with successful copulation observed in 47 offspring pairs descending from 32 parent pairs. In the “late” aged cohort mating success was lower at 63.64%, with spermatophore transfer only observed in 36 offspring pairs descending from 26 parent pairs. We counted eggs and hatchlings using the cell count tool in IMAGEJ, with these counts being conducted by a single observer (MDP). For ease of analysis, we pooled the measures of offspring fecundity and hatching success across two egg batches (mean ± SD = 1.44 ± 0.50 egg batches per female). Our final sample size consists of eggs laid by 83 offspring pairs descending from 45 parent pairs (“early*”* = 47 offspring pairs descending from 32 parent pairs, “late” = 36 offspring pairs descending from 26 parent pairs). In each temperature treatment, our sample size is as follows: 25.5°C – “early” = 17 offspring pairs from 12 parent pairs, “late” = 12 offspring pairs from 9 parent pairs; 28.0°C – “early” = 18 offspring pairs from 11 parent pairs, “late” = 11 offspring pairs from 9 parent pairs; 30.5°C – “early” = 13 offspring pairs from 9 parent pairs, “late” = 12 offspring pairs from 8 parent pairs. A single pair was removed from the hatching success analysis due to unreliable count data (many nymphs had died at the time of the hatchling counts), leaving 82 egg batches for the offspring hatching success analysis.

### 2.6 Data Analysis

We undertook all statistical analysis and data visualisation in *RStudio v.2025.05.1+513*, using *R v.4.3.3* (R Core Team, 2024). We used two different Bayesian statistical workflows, describing average values and variation in life-history traits using *brms (section 2.6.1),* before characterising variation in offspring baseline and age-specific mortality with *BaSTA (section 2.6.2*). Below, we first separately detail our workflow in both frameworks and then introduce the structure and covariates included when modelling each trait (*section 2.6.3*).

#### 2.6.1. General procedure for modelling life-history traits in brms

##### Model workflow

We used the package *brms v.2.21.0* (Bürkner, 2017), implementing *rstan* v.2.32.2 (Gelman *et al*., 2015; Stan Development Team, 2024) as a backend, to create Bayesian generalised linear mixed models (GLMMs). We follow the location-scale approach to regression modelling, where both first-order (i.e., the mean/location; *μ*) and second-order (i.e., the variance/scale; *σ or ϕ*) traits are estimated (O’Dea *et al*., 2022; Nakagawa *et al*., 2025). This location-scale modelling approach permits heteroskedasticity, where differences in error variance exist across the predictors (Barreda & Silbert, 2023). Crucially, these models can tease apart how parental age affected not just the offspring’s average trait value but also the spread of variation in trait values. As the best-fitting model for hatching success was a zero-inflated beta-binomial (*Table 1*), we also placed fixed effects on the zero-inflation parameter (*zi)* to allow parental age to predict the likelihood of complete hatching failure. We included both fixed and random terms in the location submodel, while only fixed effects were included in the scale and zero-inflation components. This single-hierarchical framework ensured that the distributional parameters remained identifiable, given these parameters require large sample sizes to be robustly estimated (Nakagawa *et al*. 2025). We performed model simplification only within the scale and zero-inflation component by removing predictors exhibiting no effect (i.e., posteriors with 95% credible intervals that crossed zero) to reduce the risk of model overparametrisation (Barreda & Silbert, 2023). We always retained parental age in the scale and zero-inflation components, being central to our underlying question (i.e., whether parental age affects the *variability* of offspring traits). In the location component, we did not perform model simplification beyond the removal of non-meaningful interactions (including quadratic terms) to ease the interpretation of single-effect predictors. All continuous covariates were z-transformed (mean = 0, SD = 1), to facilitate prior specification. We used the package *ggplot2 v.4.0.0* (Wickham, 2016) to visualise the raw data and posterior predictions, while we produced model tables using the package *gtsummary v.1.7.2* (Sjoberg *et al*., 2021).

**Table 1.** Likelihoods, parameters, link-functions, and sample sizes used when modelling each life-history trait.

| **Trait**<br>***(response)*** | **Likelihood & Package** | **Parameters & link** | ***n*** | **Corresponding Appendices** |
|---|---|---|---|---|
| Egg length<br>(*mm*) | Lognormal<br>*(brms)* | *μ (identity)*<br>*σ (log)* | 4216 eggs<br>77 parent pairs | ***Section S6***<br>*Model selection:* Table S4<br>*Estimates:* Table S5 |
| Early-life survival<br>(*survived =yes/no*) | Bernoulli<br>*(brms)* | *μ (logit)* | 1317 offspring<br>78 parent pairs | ***Section S7***<br>*Model selection:* Table S6<br>*Estimates:* Table S7 |
| Juvenile survival<br>*(survived =yes/no)* | Bernoulli<br>*(brms)* | *μ (logit)* | 987 offspring<br>77 parent pairs | ***Section S8***<br>*Model selection:* Table S8<br>*Estimates:* Table S9 |
| Total lifespan<br>*(weeks)* | Weibull<br>*(brms)* | *μ (log)*<br>*k (identity)* | 987 offspring<br>77 parent pairs<br>sexed subset: 947 | ***Section 9.1***<br>*Model selection:* Table S10<br>*Estimates:* Tables S11 & S12 |
| Total mortality<br>*(c & b<sub>1</sub> = years*<br>*b<sub>0</sub> = unitless)* | Weibull-Makeham<br>*(BaSTA)* | *c, b<sub>0</sub>, b<sub>1</sub>*<br>*(identity)* | 987 offspring<br>77 parent pairs<br>sexed subset: 947 | ***Section S9.2***<br>*Model selection:* Table S13<br>*Estimates:* Table S14 |
| Development time<br>*(weeks)* | Lognormal<br>*(brms)* | *μ (identity)*<br>*σ (log)* | 939 offspring<br>77 parent pairs | ***Section S10***<br>*Model selection:* Table S15<br>*Estimates:* Table S16 |
| Adult mass<br>*(grams)* | Gaussian<br>*(brms)* | *μ (identity)*<br>*σ (log)* | 937 offspring<br>77 parent pairs | ***Section S11***<br>*Model selection:* Table S17<br>*Estimates:* Table S18 |
| Relative growth rate<br>*(RGR; week-1)* | Gaussian<br>*(brms)* | *μ (identity)*<br>*σ (log)* | 937 offspring<br>77 parent pairs | ***Section S12***<br>*Model selection:* Table S19<br>*Estimates:* Table S20 |
| Fecundity<br>*(No. eggs laid)* | Negative binomial<br>*(brms)* | *μ (log)*<br>*ϕ (log)* | 83 offspring<br>45 parent pairs | ***Section S13***<br>*Model selection:* Table S21<br>*Estimates:* Table S22 |
| Hatching success<br>(*No. hatched eggs per trial)* | Zero-inflated beta-binomial<br>*(brms)* | *μ (logit)*<br>*ϕ (log)*<br>*z (logit)* | 82 offspring<br>45 parent pairs | ***Section S14***<br>*Model selection:* Table S23<br>*Estimates:* Table S24 |
| Adult lifespan<br>*(weeks)* | Weibull<br>*(brms)* | *μ (log)*<br>*k (identity)* | 938 offspring<br>77 parent pairs | ***Section S15.1***<br>*Model selection:* Table S25<br>*Estimates:* Table S26 |
| Adult mortality<br>*(c & b1 = years*<br>*b0 = unitless)* | Weibull-Makeham<br>*(BaSTA)* | *c, b0, b1*<br>*(identity)* | 939 offspring<br>77 parent pairs | ***Section S15.2***<br>*Model selection:* Table S27<br>*Estimates:* Table S28 |

*Note: Sample sizes describe the number of offspring from our starting pool (n = 1317 individuals) included in each model. Corresponding supplements refer to appendix sections detailing the model selection process (e.g., choice of likelihood, evidence for interactions) and parameter estimates.*
*Abbreviations: μ = location parameter; k = shape parameter (Weibull model in brms); c = Makeham term; b0, b1 = age-specific mortality parameters (scale, and shape parameter, respectively); σ = residual standard deviation; ϕ = shape/dispersion parameter (Negative binomial/Zero-inflated beta-binomial); z = zero-inflation parameter (zero-inflated beta-binomial).*

##### Model settings

Unless otherwise stated, we used the following settings for our fitted models: four Hamiltonian Monte Carlo (HMC) parallel chains, 5000 iterations each (2500 warmup chains which were discarded), and a thinning of one. This resulted in 10,000 post-warmup posterior samples per parameter. We followed standard practice and monitored model convergence and the effective sample sizes as recommended by Gelman and Shalizi (2013). We used the potential scale reduction statistic (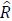) supplied by the model output to compare between and within-chain estimates for the model parameters. We assumed that chains had sufficiently mixed when 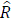 <1.01 (Gelman & Rubin, 1992; Vehtari *et al*., 2021). Model convergence was said to be achieved when the effective sample sizes (both bulk and tail) for model parameters were >1,000 (Gelman & Rubin, 1992; Vats & Knudson, 2021).

##### Model diagnostics

To assess goodness-of-fit, we inspected the Leave-One-Out Probability Integral Transform (LOO-PIT) plots from *bayesplot v.1.11.1* (Gabry & Mahr, 2025), ensuring the LOO-PIT integrals were uniformly distributed (Vehtari *et al*., 2017; Säilynoja *et al*., 2022; Nguyen *et al*., 2025). Additionally, we used the *ppc_loo_intervals()* function to determine how well our model predicted each observation after it was blinded to the original data. We also visualised how well the empirical distribution of the data (*y*) fitted to the distributions of simulated data (*yrep*) from the posterior predictive distribution, using the *pp_check()* function (Gabry *et al*., 2019; Winter and Bürkner, 2021). All model diagnostics for each trait are presented in the supplementary materials (*Table 1*). We checked for any multicollinearity between predictors (VIF) using the package *car v.3.1.2* (Fox & Weisberg, 2019) and by inspecting correlations between pairs of predictors with the *mcmc (type= “pairs”)* function.

##### Model predictive performance

To compare the out-of-sample predictive performance of each model, we used the expected log pointwise predictive density (*elpd*), calculated with Pareto smoothed importance sampling Leave-One-Out cross-validation (PSIS-LOO; Vehtari et al. 2015, 2017). For a series of models fitted to the same data, a higher *elpd* value indicates greater predictive performance (Vehtari *et al*., 2016). We assumed that models had a similar predictive ability if the difference in between their *elpd* (*Δelpd*) was < four (Sivula *et al*., 2020). In instances where the predictive ability of the compared models was similar, we selected the simplest model with the fewest predictors and no interactions.

##### Choice of prior distribution

For each trait, we fitted the same model with progressively tighter priors. These ranged from the default flat priors provided by *brms* to normally distributed constraining priors that assumed no effect of the fixed effects on the response (Gelman *et al*., 2008; Gelman *et al*., 2017). Unless otherwise stated, we fitted our final models with weakly informative priors, using *β_1_∼normal (mean = 0, SD = 1 SD_y_)* priors for fixed effects, accounting for the scale of the response and the model’s link function (Lemoine *et al*., 2016; Lemoine, 2019). This weakly informative prior assumes a one standard deviation increase in the predictor leads to an approximately one standard deviation change in the response, and rarely any larger scale changes (Møller and Jennions, 2002; Lemoine, 2019). For a detailed overview of our prior specification process, including exact prior values for each model parameter and their justification, see *Appendix S1: Section S4*.

#### 2.6.2 General procedure for mortality models in BaSTA

##### Choice of mortality distribution

We used Bayesian survival trajectory analysis (*BaSTA v.2.0.1*; Colchero *et al*., 2012) to build parametric survival models explaining variation in offspring mortality. *BaSTA* uses a Monte Carlo Markov chains (MCMC) algorithm and describes mortality as the instantaneous risk of death given survival to the specified age (Colchero and Clark, 2012). We fitted all possible mortality distributions parameterized by *BaSTA* which included the exponential (Cox & Oakes, 1984), Gompertz (Gompertz, 1825; Pletcher, 1999), Weibull (Pinder *et al*., 1978), and logistic (Vaupel *et al*., 1979) distributions which we extended with the Makeham (*c;* Makeham, 1867) and bathtub (Siler, 1979) terms. The predictive performance of all the models was compared using their deviance information criterion (DIC) value. DIC describes the predictive power of each model by approximating the expected predictive deviance and penalises for model complexity, with a lower DIC indicating that the model has improved predictive ability (Spiegelhalter *et al*., 2002; Millar, 2009). We selected the model with the lowest DIC, with a model said to be the best-performing when its ΔDIC < 3 from alternative candidate models (Spiegelhalter *et al*., 2002). If models had comparable DIC values, we selected the simplest model (with the fewest predictors/no interactions).

Out of all mortality distributions, we found the Weibull-Makeham model had the best predictive performance (*Appendix S1: Tables S13 and S27*). We explain the Weibull-Makeham model in detail within the supplementary materials (*Appendix S1: Section S5.1*). Briefly put, the Makeham term (*c*) describes baseline mortality, the shape parameter (*b_0_*) describes the pattern of the mortality curve (with values >2 indicating an accelerating increase in mortality risk), while *b_1_* describes the characteristic failure time (i.e., when 63.2% of individuals have died; Gómez *et al*., 2023), which *BaSTA* then inverts. Taken together, *c* sets the mortality intercept, while *b_0_* and *b_1_* describe the age-specific pattern of mortality; *b_1_* scales the relationship between the age of the individual and the shape of the curve (*b_0_*), setting the inflection point for when the mortality increase occurs.

##### Structure and model settings

Since *BaSTA* does not provide Kullback-Leibler discrepancy calibration values (KLDC; (Kullback & Leibler, 1951) for continuous predictors, we only included categorical predictors in our mortality models (using the *“fused”* covariate structure). Between a pair of predictors, KLDC provides a measure of the distance between their posterior distributions for each parameter. A value of 0.5 indicates that posterior distributions overlap completely, while a KLDC of one indicates that the posterior distributions exhibit no overlap (McCulloch, 1989). When KLDC >0.8, we concluded that there was a substantial difference between the compared predictors in their effect on the mortality parameter, assuming this indicated a genuine biological effect (Karabatsos, 2006; Rodríguez-Muñoz *et al*., 2019b).

Our final posterior distributions were based on four MCMC chains with 80,000 iterations (burn-in of 1001), and a thinning interval of 50, resulting in n = 6320 post-burn-in posterior samples per parameter. We assessed the convergence of the parameter estimates using 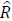. Despite reducing the length of the burn-in sequence, the simulations in all our models rapidly converged, mixing well when stabilising towards the target distribution (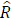 <1.002), and serial autocorrelation was low (<5%). We assessed model fit visually by overlaying the parametric survival curves over the Kaplan-Meier plots of observed survival (*Appendix S1: Figures S16 and S31*). We used *ggplot2* to visualise parameter posterior distributions, alongside the offspring’s cumulative survival curves and instantaneous hazard rates.

##### Choice of prior distribution

To assess the sensitivity of the mortality parameters to our choice of prior structure, we conducted the analysis using three prior distributions (default, weakly-informative, and moderately-informative), before selecting normally distributed weakly informative priors for covariates, as recommended by Lemoine (2019). Specifically, we set the prior for *c* and *b_0_* to the weakly-informative *BaSTA* defaults (*c*∼normal [0,1], *b_0_*∼normal [1.5, 1]). For *b_1_,* we adapted the default prior (normal [0, 1]) to a scale that was appropriate for the short lifespan of *G. bimaculatus*. Given *BaSTA* was developed for long-lived wild vertebrates, the default *b_1_* prior is centred incorrectly for our study species. We therefore scaled the *b_1_* prior to the raw population mean for total or adult lifespan (*details below*). All priors were truncated to be ≥ 0, given the Weibull model’s parameterisation. We also evaluated tighter moderately informative priors, with the same mean, but ½ the SD of the weakly informative priors (*Appendix S1: Tables S13 & S27*).

#### 2.6.3. Statistical models

##### Decomposition of parental age

We used within-subject mean centring to separate parental age into two terms: *average age (age)* and *Δage.* This method controls for between-individual heterogeneity in reproductive lifespan and removes the confounding bias arising from the selective disappearance of parent pairs (Van De Pol & Wright, 2009). The first term describes the between-parent effect of age *(age)*, which was calculated as the average age of the mother across mating attempts with successfully hatched eggs. The second term captures the within-parent age effect *(Δage*) and describes the difference between the parents’ average age and the mother’s age at the current mating attempt. Both terms were included in our statistical framework, with *age* controlling for selective disappearance, while *Δage* captured the true within-parental age effect. Across all models, a unit increase in *Δage* corresponds to a single standard deviation increase in the parents’ age at reproduction, or 2.06 weeks on the original scale. Meanwhile, a single SD increase in *age* corresponds to an increase in the parents’ average reproductive age of 0.92 weeks. As a test for selective disappearance, we also calculated the difference between the slopes of these two age terms. If the slopes for *age* and *Δage* matched (i.e., did not statistically differ from zero), then we assumed there was little evidence that selective disappearance masked the true parental age effect (Van De Pol & Wright, 2009). The parents’ *Δage* was initially included as a quadratic in all models (*Δage^2^*), before this term was removed from the majority of models to ease interpretation of the linear slope and simplify tests of selective disappearance (Fay *et al*., 2022; Sanghvi *et al*., 2026), as this term did not improve out-of-sample predictive performance. The only trait in which *Δage^2^* was retained was egg length, with the quadratic term receiving substantial statistical support (*Appendix S1: Table S4*). Subsequently, we then estimated the ‘*peak age’* of egg length as – *Δage/2*Δage^2^* (Bronshtein *et al*., 2015; Dingemanse *et al*., 2020), ensuring that the uncertainty of this derived parameter was estimated by carrying forward the posterior distributions of the component age terms. We initially aimed to include the parents’ cumulative number of successful mating attempts (i.e., mating attempts from which eggs hatched) as an additional covariate, to tease apart variation in offspring phenotypes explained by accumulated reproductive effort. However, this term was tightly correlated with *Δage* owing to our longitudinal study design (cumulative number of mating attempts ∼ *Δage*, Pearson’s correlation coefficient *r* [95% CI] = 0.94 [0.92, 0.95]; *Appendix S1: Figure S2*) and was not included in our modelling framework.

##### Relative growth rate

We calculated the relative growth rate (RGR; *equation 2* in Pommerening and Muszta, 2015) of each offspring as *ln(adult mass/initial mass at hatching; in mg)/development time; in weeks*, which expresses biomass gain as the rate of increase in log (mass) per week (*week^-1^*). As we did not measure initial mass at hatching, our calculation assumes that all offspring shared the same starting value. To set this initial value, we used the mean mass of a first instar nymph (0.750 mg) from a previous study on *G. bimaculatus* (Behrens *et al*., 1983), applying this value as a fixed constant across all individuals. Given our RGR calculation does not permit variation in initial mass, it should not be interpreted as a true standardised growth rate. Rather, it describes the rate of mass accumulation over the time taken to reach adulthood. We suggest that any bias arising from this shared starting value may be minimal; *G. bimaculatus* adults weigh between 500-1500 mg (Behrens *et al*., 1983), over 600 times their mass at hatching. Therefore, initial hatchling mass will only have a comparatively small influence over the final RGR estimate.

##### Approach to modelling offspring survival

Due to high early-life mortality, we could not include all offspring within a single Weibull-Makeham model that explained variation in both frailty and age-specific mortality. Even when we extended this model with a bathtub term, which introduced a declining Gompertz function to describe the more complex shape of the survival curve (*Appendix S1: Section S5*), this could only weakly capture the steep early-life mortality of our study population (*Appendix S1: Figure S3*). We therefore abandoned using a single parametric survival model with all the offspring included and instead used a mixed modelling approach to describe variation in longevity and mortality.

##### Pre-adult survival

First, we estimated the initial frailty of all the offspring during early-life with a logistic regression (n = 1317 offspring from 78 parent pairs; Table 1). As the Kaplan-Meier curve of lifetime survival (and raw weekly survival plots) suggested that early-life mortality plateaued after week four (*Appendix S1: Figures S3 & S6*), we selected this as the upper boundary for assessing initial frailty after hatching (*Figure 3*). The binary response indicates whether an individual offspring survived their first four weeks of life (i.e., survived = 1, died = 0). To capture survival in the remainder of the pre-adult period, we also estimated juvenile survival, describing if an individual offspring survived from week four post-hatching until adulthood (survived =1, died = 0). This analysis only included the 987 offspring that survived through the early-life window. We note that both these models cannot capture any time-dependence in offspring early-life survival (i.e., differences in longevity and ageing rates in each frailty window). However, from the raw data, the offspring’s weekly survival probabilities as estimated *within* this early-life frailty window (i.e., survival through weeks one, two, three, & four) appeared independent of parental age (*Appendix S1: Figure S6*). Thus, it is unlikely that our cumulative survival estimate over these first four weeks is biased by any individual weekly differences in early-life survival.

##### Total lifespan

We then used the Weibull models in *brms* and *BaSTA* to estimate the total lifespan and mortality of the 987 offspring that survived their first four weeks of life. Entry into the Weibull model was conditional on survival through the early-life window, with this analysis selectively removing frail offspring who died in this period. We first described variation in offspring longevity (and adult longevity) with the *brms* Weibull model, which allowed for the inclusion of multiple covariates and random terms (*all covariates detailed below*). However, under *brms,* we could not place covariates on the shape parameter (*k; analogous to b_0_*) as this inflated model complexity, with the model failing to converge as indicated by low ESS values (<100) and 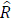 values > 1.5. Therefore, we only estimated the effect of the model covariates on mean lifespan (*µ*) in *brms*. We used the Weibull-Makeham model in *BaSTA* to describe variation in the offspring’s baseline (*c*) and age-specific mortality parameters (*b_1_* and *b_0_*), across either the offspring’s total or adult lifespan.

##### Covariates included when modelling traits in brms

Models on each offspring trait used different likelihoods and subsets of available data (*Table 1*), but the covariate structure remained comparable across the following traits: egg length, early-life survival, juvenile survival, total lifespan, development time, adult mass, RGR, and adult lifespan. We first explained variation in the offspring’s average trait values (i.e., the location component) with the following fixed effects: *Δage* (i.e., the within-individual parental age effect), *age* (i.e., the parents’ average reproductive age, to control for selective disappearance), and *Temp* (the parental temperature treatment, a three-level factor; “25.5”, “28”, and “30.5°C”). For traits where the offspring were sexed (development time, adult mass, RGR, and adult lifespan), we also included *offspring sex* (two-level factor: “Male”, “Female”) as a fixed effect predictor. Meanwhile, for egg length, offspring adult mass, and RGR, the *mother’s adult mass* (*in grams)* was fitted as a fixed effect. Our global model on offspring total lifespan did not include offspring sex, to allow for the inclusion of unsexed nymphs that died before adulthood. Therefore, we also conducted a subset analysis on the sexed adults (n = 947; *Table 1, Appendix S1: Table S12*), enabling offspring sex to predict variation in their total lifespan. For egg length, we also included the quadratic parabola for parental age (*Δage^2^)* and *Incubation day* (two-level factor; *“day 0”* or *“day 5”*) as fixed effects. In all models (except offspring fecundity and hatching success; *details below*), we included *parent pair ID* as a random intercept and fitted *Δage* as a random slope, allowing the effect of *Δage* to deviate between parent pair*s.* For egg length, we also included *eggID* as an additional random intercept, to control for variation explained by a shared batch identity (e.g., differences in sand quality/petri dish effects). Additionally, we slightly change the language when presenting the results for egg length, referring to the parental age effect as a *‘maternal age’* effect, as we assume egg length to be a maternal trait.

For those traits where offspring were sexed, we always tested support for the three-way interaction between *offspring sex x Temp x Δage*, including the lower-order interactions between these terms. Meanwhile, for egg length, we tested evidence for the three-way interaction between *Incubation day x Temp x Δage*. We then fitted additional models with either the two-way interactions between *Temp x Δage*, *offspring sex x Δage,* or *Incubation day x Δage*. However, these interactive terms did not improve predictive performance and were dropped from our final inference. Despite being removed from our final models, we always report the statistical evidence for the *Temp x Δage* (and for egg length, the additional *Temp x Δage^2^* interaction) for all traits, as this directly addresses whether the parental age effect varied between temperature treatments.

For likelihoods containing a residual error or dispersion term (i.e., the *scale component*; *Table 1*), we always included *Δage* as a fixed effect on this distributional parameter. We initially included *Temp* and *offspring sex* as additional fixed effects but these terms were later removed (except in the egg length model) as they did not improve predictive performance, and their effects were only weakly identifiable (i.e., displayed flat posteriors with 95% credible intervals that crossed zero). For egg length, the best-supported model included *Temp*, *Incubation day*, and *Δage* in the scale component. We also tested the two-way interactions between *Temp x Δage*, *offspring sex x Δage*, and *Incubation day x Δage* in the scale component, with these interactions removed from our final model if they did not improve predictive performance (*Table 1*). For egg length, the best-supported model included the two-way interaction between *Incubation day x Δage* in the scale component.

For models on offspring fecundity and hatching success, which used a smaller subset of the available data (*Table 1*), we only included the *parents’ age category* (two-level factor: “Early-aged”, “Late-aged”) and *Temp* as fixed effects in the location component, and *parent pair ID* as a random intercept. Additionally, we tested the statistical support for the two-way interaction between the *parent’s age category x Temp*, before this was dropped from the final model to ease interpretation of single effect predictors. We only added the *parents’ age category* as a fixed effect on the distributional parameters (*ϕ or z*) to retain model identifiability, as these models only used a small subset of individuals.

##### Covariates included in the mortality models in BaSTA

We used Weibull-Makeham models specified by *BaSTA* to describe variation in offspring mortality across either the offsprings’ total lifespan (as estimated for those offspring that survived their first four weeks of life: n = 987 offspring) and adult lifespan (mortality across the *adult lifespan*, from adult eclosion until death: n = 939 offspring, *Table 1*). As we only included categorical covariates in the *BaSTA* model (to obtain KLDC estimates), we binned parental age into three categories: “Early-aged” (mating attempts 1 & 2), “Middle-aged” (mating attempts three to five), and “Late-aged” (mating attempts six to eight). We first included the isolated effects of *parental age* (three-level factor: “Early-aged”, “Middle-aged”, or “Late-aged”) or *Temp* as single-effect predictors on each mortality parameter (*c*, *b_1_*, and *b_0_*), before assessing their joint effect (two-way *parental age* x *parental temperature* interaction). For the subset of sexed offspring (947 offspring, *Table 1*), we also tested support for the interaction between *the parents’ age x offspring sex*. We then compared each candidate model to the null model, which contained no predictors. If the DIC of the null model was lower than the alternative candidate models with covariates, we concluded that the predictors did not explain variation in offspring mortality, although we always report the effects of *parental age*.

We used a weakly-informative prior structure, placing the default priors on *b_0_* and *c*, while we scaled the *b_1_* prior. For total lifespan, (raw mean ± SD lifespan of 18.31 ± 3.97 weeks), this conversion resulted in a truncated *b_1_ ∼* half-normal (2.57, 0.5) prior, permitting characteristic lifespans of between 10 and 40 weeks. Meanwhile, for adult lifespan, (raw mean ± SD adult lifespan = 9.53 ± 3.16 weeks), we used a truncated *b_1_*∼half-normal (4.94, 1) prior, permitting characteristic adult lifespans of between five and 24 weeks. These weakly-informative priors did not change our posterior inferences when compared to the default prior structure (*Appendix S1: Table S13* and *Table S27*).

##### Section 2.6.4. Summary of the approach to the presentation of results

For brevity, in the main manuscript we report the probability of direction (*pd*) and Region of Practical Equivalence (*ROPE*) for each fixed effect estimate, as estimated from *bayestestR v.0.17.0* (Makowski *et al*., 2019a, 2019b). We used *pd* to describe the proportion of posterior estimates in the same direction as the MAP (maximum a posteriori probability estimate). We assumed that parameter estimates with a *pd* ≥ 98% received statistical support, with this corresponding to a two-sided *p* value threshold of 0.05 (Makowski *et al*., 2019b). We then used the ROPE to determine whether the effect was non-negligible (i.e., the percentage of the posterior distribution in the null region). ROPE defines an area around the null value that is practically equivalent to zero effect and potentially of negligible interest (Kruschke, 2018). In this study, we set ROPE to the equivalent of ±0.1*SD*_y_* (Kruschke, 2018). This range was appropriately log-transformed for likelihoods using a log-link, while for families with the logit link (i.e., Bernoulli & beta-binomial), we set ROPE to the suggested default (± 0.18 log-odds; Kruschke, 2018). Together these two statistics provide information on the statistical evidence (*pd*) and likelihood of a true biological effect (ROPE). When reporting our results we use a language of evidence (based on the ROPE and *pd*), as suggested by Muff *et al*. (2022) and detailed in *Figure 2*, following guidelines for Bayesian reporting outlined by Makowski *et al*. (2019a, 2019b) and Schwaferts and Augustin (2020). Full model summaries are provided in the supplementary results of each trait (*detailed in Table 1*), including the MAP, 95% Highest Density Intervals (HDI’s), and effective sample sizes (ESS). We only applied the *pd* to the scale and zero-inflation components, as defining a ROPE on these distributional parameters was not clearly interpretable.

**Figure 2.**
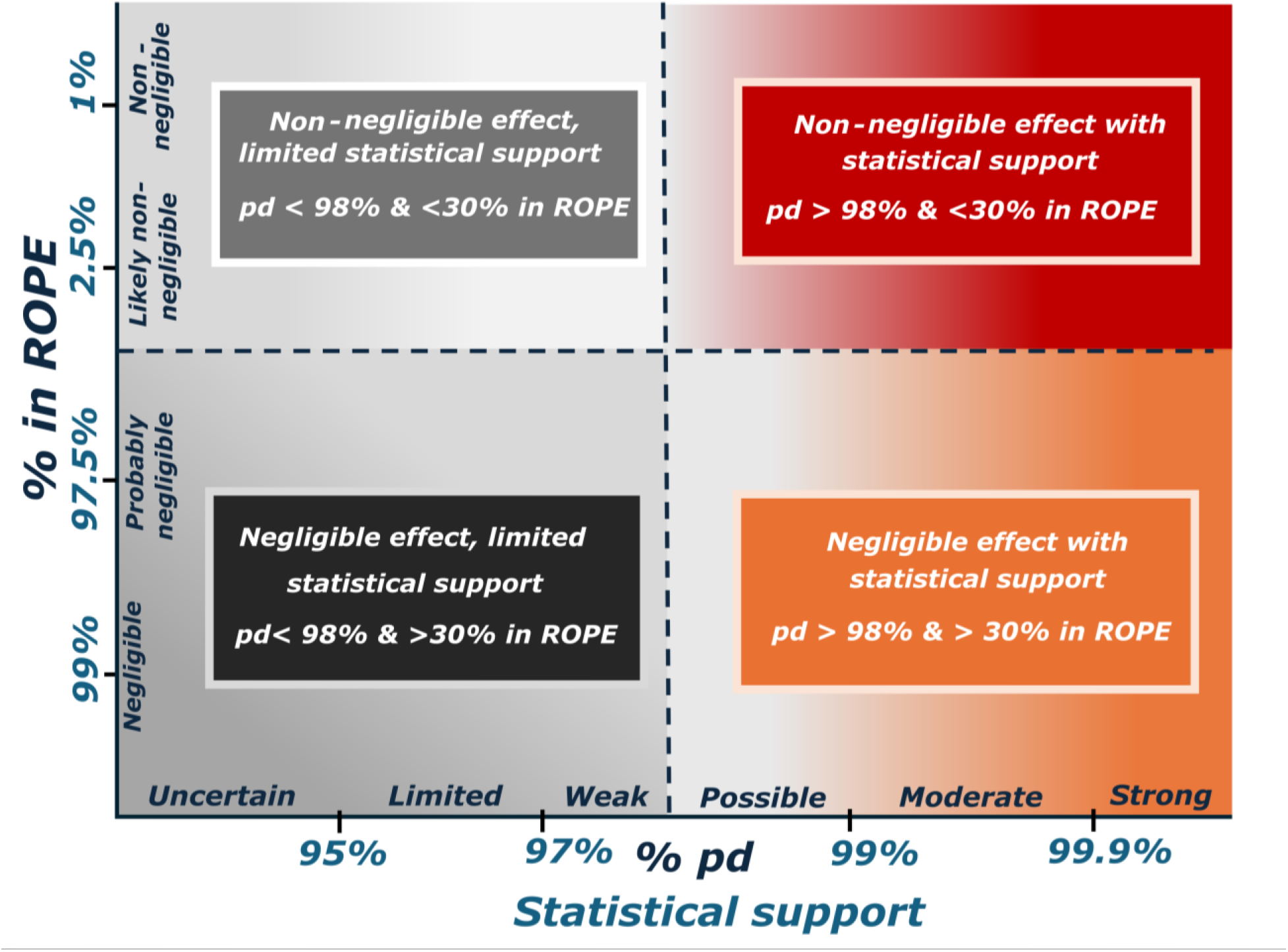
Overview of our language used for statistical reporting. Displayed is how we described the statistical evidence for fixed effects, based on the percentage of posterior estimates falling within the region of practical equivalence (*ROPE*; y-axis) and their probability of direction (*pd*; x-axis). Shown along each axis is our standard language of evidence used for reporting the statistical support of Bayesian effects as outlined by Makowski *et al*. (2019a, 2019b).

For post-hoc inferences, we used the package *emmeans v.1.10.1* (Lenth, 2024) to report posterior medians and 95% HDI’s on the response scale. We first describe the posterior predicted trait values at the lower (2.5^th^ percentile; or –1.96 standard deviations (σ) from the mean *Δage*) and upper (97.5^th^ percentile; or +1.96 standard deviations (σ) from the mean *Δage*) tails of *Δage,* as these represent the lower and upper tails of a normally distributed parameter (Campbell, 2017). We infer these tails to represent the average trait values of offspring from young (–1.96σ) and old (+1.96σ) parents as in our study these tails differ in parental age by eight weeks in total, coinciding with the total age range over which parents mated. Since egg length followed a non-linear relationship with *Δage (see results),* in this instance we base our post-hoc comparisons between the peak age (*Δage* = 0.43*; details in section 3.2)* and old age (+1.96σ). Additionally, we then plotted the effect size of each parental age effect. Here, we used *emmeans* to derive the post-hoc average trait values for offspring from young and old parents (or for egg length, between peak aged and old-aged mothers), calculated the pairwise difference between these trait values, and then divided this difference by the raw population standard deviation of the trait. This then gave a post-hoc standardised mean difference (SMD) for the parental age effect. When presenting the SMD of the post-hoc estimates (i.e., effect size of *Δage*), we indicate which effects are considered small (± 0.2 – 0.5), moderate (± 0.5 – 0.8), or large (> ± 0.8) following thresholds suggested by Andrade (2020).

## 3. Results

### 3.1 Summary of the observed parental age effects

Overall, increasing parental age had heterogeneous effects on the measured life-history traits (*Figure 3*). Parental age had negligible effect on offspring early-life and juvenile survival (i.e., displayed 95% HDI’s that crossed zero and a SMD < 0.2), while the positive effects of parental age on offspring reproductive performance (fecundity and hatching success) carried substantial posterior uncertainty (95% HDI’s that crossed zero but a SMD > 0.5). We observed a small negative effect of parental age on offspring adult mass, whereas parental age had a small positive effect on the offspring’s total lifespan (SMD = ± 0.2 – 0.5). Egg length displayed a moderate (SMD = ± 0.5 – 0.8) negative relationship with parental age while, contrary to The Lansing Effect, we found a moderate positive effect on offspring adult lifespan. Finally, we found large effects (SMD > ± 0.8) of parental age on offspring development time and growth rate, but in opposite directions. Those offspring from old parents were quicker to develop yet increased their growth rates in the time taken to reach adulthood.

**Figure 3.**
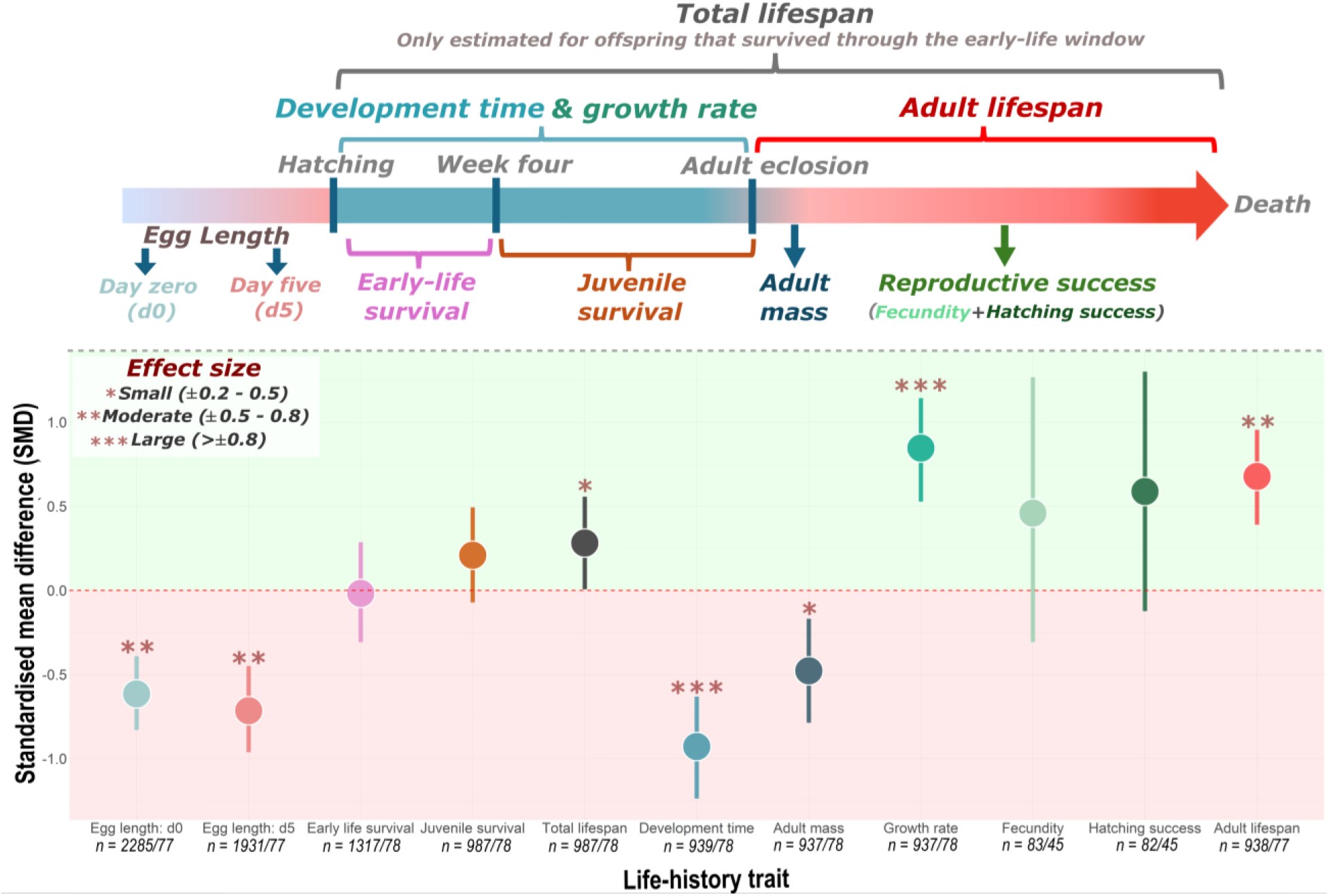
Overview of measured life-history traits and the standardised mean difference (i.e., effect size) of the parental *Δage* effect. Points represent the standardised mean difference in offspring trait values between old and young parents, while the bars describe the 95% HDI’s. Any effects crossing zero lack statistical support. Samples sizes are shown as the number of offspring/number of parent pairs. For all traits (except egg length), pairwise comparisons are shown as the post-hoc mean difference in offspring trait values between young (–1.96σ) and old (+1.96σ) parents. As egg length follows a quadratic relationship with age, the predicted post-hoc difference is instead calculated between the peak age (*Δage = 0.43 weeks; details in section 3.2*) and old age (+1.96σ). For fecundity and hatching success, pairwise comparisons are based on the mean difference between the two categorical parental age groups (“early-aged” versus “late-aged”). Asterisks denote the size of effect based on Andrade (2020).

### 3.2 Egg length

There was strong evidence that egg length displayed a non-negligible quadratic relationship with increasing *Δage* (*Δage^2^*: *pd* = 100% & 0% *in ROPE* (*± 0.01 log mm per SD_Δage_*) *Appendix S1: Figure S4 & Table S5*). Across both incubation days (*day 0* and *day 5*), egg length followed a curvilinear pattern (*Figure 4A*), increasing to a peak when *Δage* equaled 0.43 weeks (95% HDI [0.15, 0.71 weeks]), corresponding to roughly just after the mother’s own mean reproductive age (i.e., when *Δage = 0*). On the day of laying, post-hoc comparisons revealed that the average egg length of young mothers (–1.96 σ) was 2.34 mm (95% HDI [2.29, 2.38 mm]), with this increasing to 2.55 mm (95% HDI [2.53, 2.56 mm]) in peak aged mothers (*Δage = 0.43 weeks*), before deteriorating again in old-age mothers (+1.96σ) to 2.39 mm (95% HDI [2.36, 2.43 mm]). This pattern with *Δage* persisted through the incubation period to *day 5,* despite the overall increase in egg size (*Incubation day*: *pd* = 100% & 0% *in ROPE*; *Figure 4A*). We found uncertain evidence that either the linear or quadratic *Δage* effect differed across incubation days (*Incubation day* x *Δage/Δage^2^*: *pd* < 95% & > 98% *in ROPE*; *Appendix S1: Figure S4 & Table S5*).

**Figure 4.**
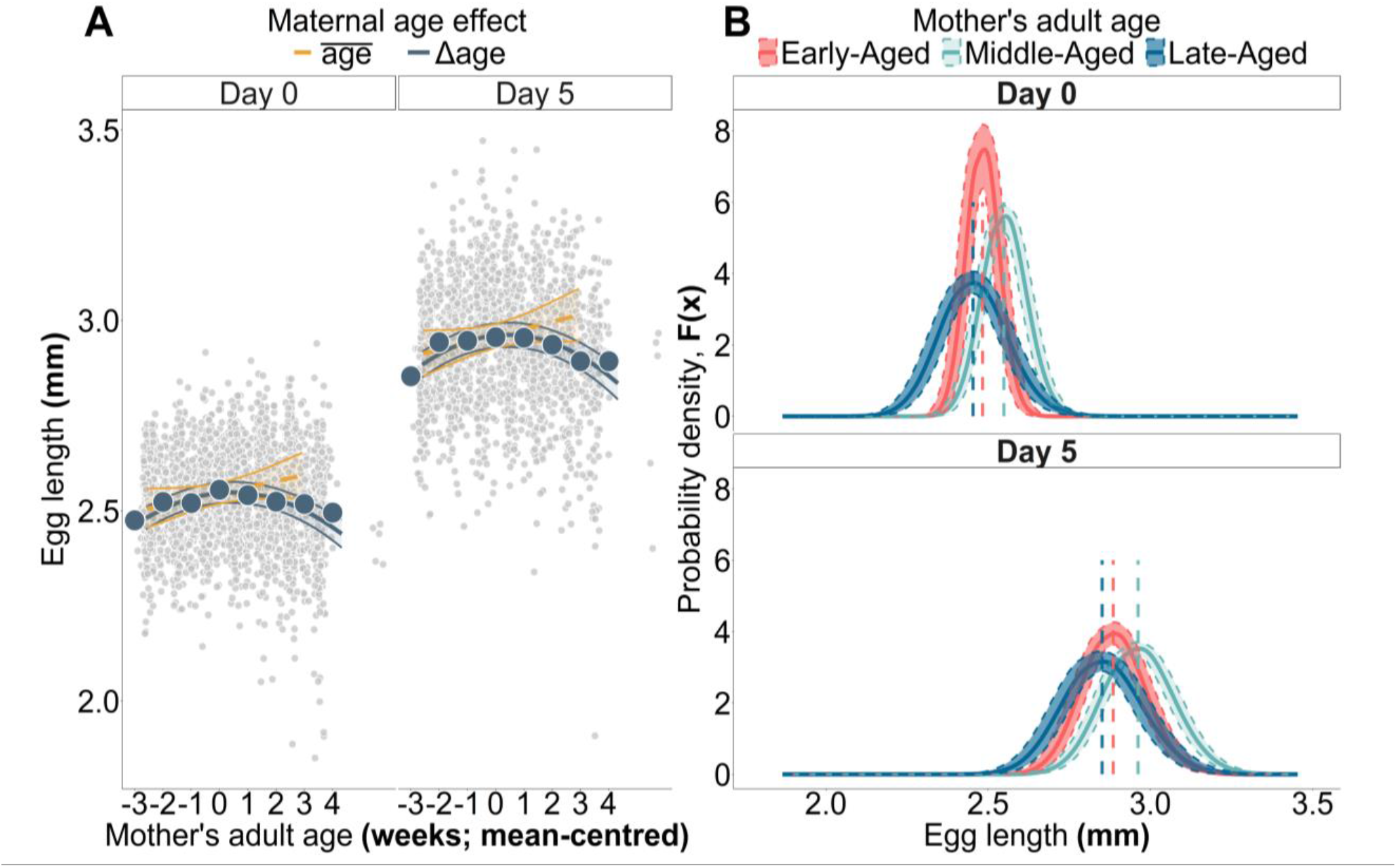
A) The effect of maternal age on egg length. On the x-axis is the mother’s age of reproduction (mean-centred) while the y-axis describes the egg length (in mm). The small dots are the raw data points, while the large dots are the raw means for the effect of *Δage* ± SE. Model predictions for the trait mean are shown as the posterior mean ± 95% CI. The colour describes the type of parental age effect **B) The effect of maternal age on distribution of egg length (i.e., probability density).** The x-axis depicts egg length while the y-axis shows the probability density function, *f(x).* The lines and ribbons display the posterior predictions (median ± 95% CI) for *f(x)*. The colours describe within-parent deviations in age relative to the parent’s mean age: “early-aged” = –1.96σ, “middle-aged” = *Δage = 0.43 weeks*, “late-aged” = +1.96σ. Dashed lines denote the posterior median estimate for egg length in each maternal age category. N = 4216 eggs from 77 parent pairs.

The rate of reproductive senescence in egg length may have differed between temperatures. From the dropped *Δage^2^ x Temp* term, there was strong statistical evidence that the *Δage* curve was more concave at 30.5°C compared to 28.0°C (*pairwise comparison Δage^2^_28.0°C vs. 30.5°C_*: *pd* = 100% & 1.99% *in ROPE*; *Appendix S1: Figure S4 & Table S5*), but not to 25.5°C (*pairwise comparison Δage^2^_30.5°C vs. 25.5°C_*: *pd* = 61.49% & 26.97% *in ROPE*; *Appendix S1: Figure S4 & Table S5*). This increased curvature suggests that the rate of reproductive senescence after the peak *Δage* was similar at the sub-optimal temperatures. There was, however, uncertain evidence for any difference in slope curvature between 25.5 and 28.0°C (*pairwise comparison Δage^2^_25.5°C vs. 28.0°C_*: *pd* = 94.28% & 58.68% *in ROPE*; *Appendix S1: Figure S4 & Table S5*). We suggest that any difference in reproductive senescence between temperatures was slight, as the posterior pairwise comparisons appeared closely situated to zero (*Appendix S1: Table S5*) and this interaction term did not improve out-of-sample predictive performance over the model only containing single-effect predictors (*Appendix S1: Table S4*).

On the day of laying, there was strong evidence that the residual variability (i.e., *σ*) of egg length increased with maternal age (*Δage*: *pd* = 100%, *Figure 4B*, *Appendix S1: Figure S4 & Table S5*). However, this effect did not persist until the fifth day of incubation, with strong statistical support for an *Incubation day x Δage* interaction (*pd* = 100%; *Appendix S1: Figure S4 & Table S5*). On day five, the residual variability in egg length appeared largely independent of maternal age (*Figure 4B*).

### 3.3 Pre-adult survival

We found that increasing parental age had a negligible effect on offspring early-life survival, with this term receiving uncertain statistical support (*Δage*: *pd* = 54.95% *&* 96.51% *in ROPE* (*± 0.18 log-odds per SD_Δage_*)*; Figure 3, Appendix S1: Figure S7 & Table S7*). Parental age had a non-negligible effect on offspring juvenile survival, but this received limited statistical support (*Δage*: *pd* = 95.74% & 23.06% *in ROPE (± 0.18 log-odds per SD_Δage_)*; *Figure 3, Appendix S1: Figure S9 & Table S9*). We found that this parental age effect on pre-adult survival (i.e., both early-life and juvenile survival) did not depend on the parents’ temperature treatment, with uncertain statistical evidence for this interaction (*Temp x Δage*: *pd* < 80% & > 24% *in ROPE*; *Appendix S1: Tables S7 & S9*)

### 3.4 Total lifespan

We found possible evidence of a positive, but negligibly sized, effect of increasing parental age on the offspring’s total lifespan (*Δage*: *pd* = 98.84% & 93.88% *in ROPE* (*± 0.03 log-weeks per SD_Δage_*); *Figure 3*, *Appendix S1*: *Figure S11 & Table S11*). On the response scale, this represented a slight increase in offspring longevity with increasing parental age (*post-hoc slope*: 0.28 weeks/SD*_Δage_*, 95% HDI [0.05, 0.53 weeks]). However, post-hoc comparisons revealed little difference in offspring total lifespan at the extreme ends of *Δage*, with offspring from old parents surviving for a median 18.80 weeks (95% HDI [18.30, 19.40 weeks]), compared with 17.70 weeks (95% HDI [17.20, 18.30 weeks]) for offspring from young parents (*Figure 3*). We found some evidence suggesting this parental age effect was temperature dependent. Here, the conditional slopes revealed moderate evidence for a non-negligible positive effect of *Δage* only at 25.5°C (*Δage_25.5°C_*: *pd* = 99.82% & 26.63% *in ROPE*). However, the coefficient describing the *Δage* by temperature interaction received limited statistical support (*Temp x Δage*: *pd* < 97% & > 40% *in ROPE; Appendix S1: Figure S11 & Table S11*), suggesting substantial posterior overlap in the *Δage* effect across temperatures. These findings were further supported by our mortality models in *BaSTA*, which found only weak positive effects of parental age on both offspring baseline (*c*) and age-specific mortality (*b_0_* and *b_1_; Appendix S1: Section S9.2)*.

### 3.5 Development time

We found strong evidence that parental age had a non-negligible effect on offspring development time (*Δage*: *pd* = 100% & 0.01% *in ROPE* (*± 0.01 log weeks per SD_Δage_*); *Figure 5*, *Appendix S1*: *Figure S17 & Table S16*). Offspring development time decreased with increasing parental age (*post-hoc slope* = –*0.35 weeks/SD_Δage_*, 95% HDI [-0.45, –0.25 weeks]), with those offspring from old parents eclosing at 8.43 weeks on average (95% HDI [8.23, 8.64 weeks]) compared with 9.81 weeks (95% HDI [9.58, 10.07 weeks]) for offspring of young parents (*Figures 3 & 5B*). There was uncertain statistical evidence to suggest that the parental age effect differed between temperature treatments (*Temp x Δage*: *pd* < 80% & > 50% *in ROPE; Appendix S1*: *Figure S17 & Table S16*).

**Figure 5.**
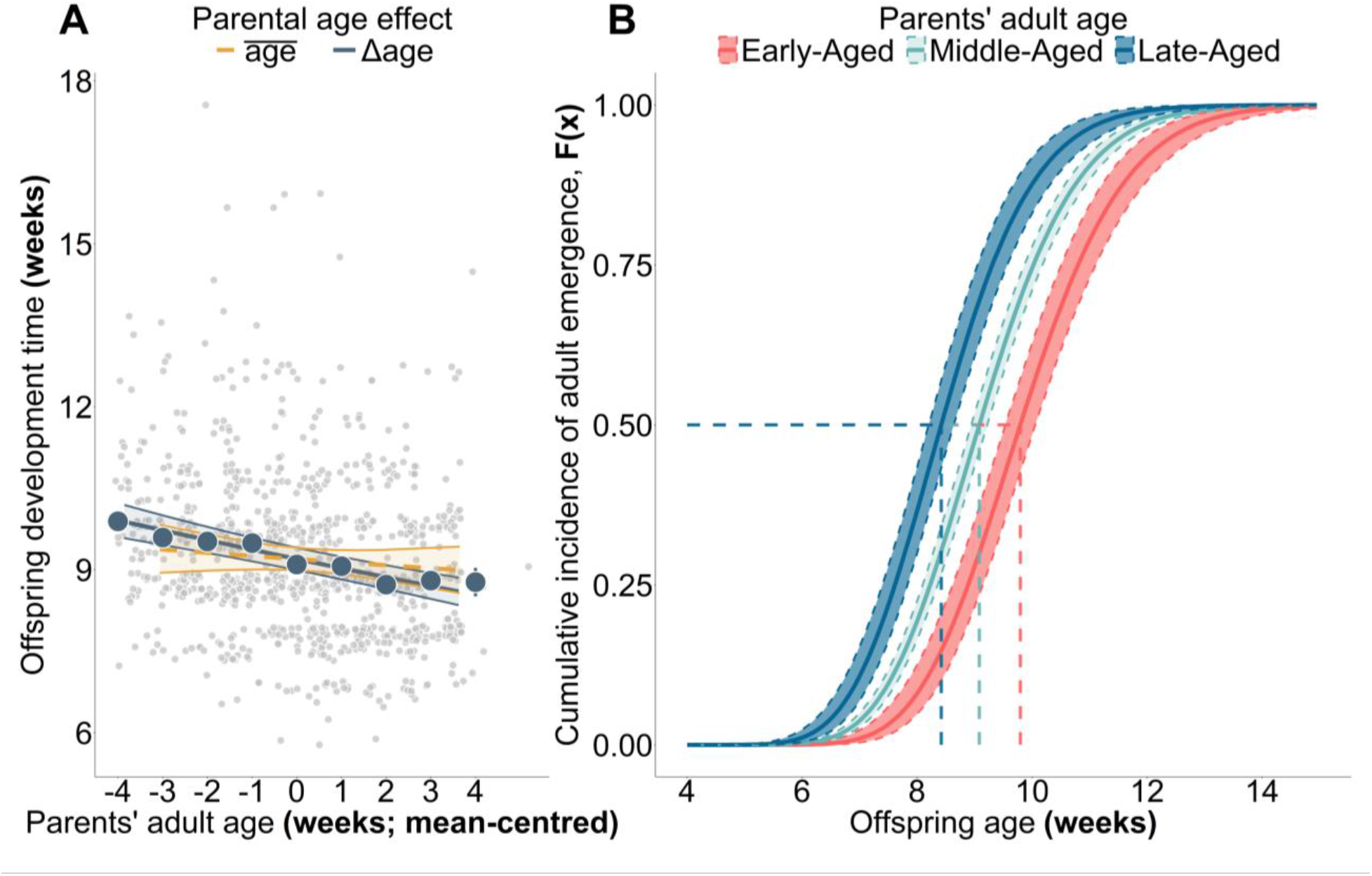
A) The effect of parental age on offspring mean development time. On the x-axis is the parents’ age at reproduction (mean-centred) and on the y-axis is offspring development time (weeks). Small dots are raw data while large dots are raw means for the Δage effect ± SE. The lines and ribbons represent the posterior mean effect ± 95% CI. The colours represent the type of parental age effect. **B) The effect of the parents’ age at reproduction on the cumulative incidence of offspring adult emergence.** The x-axis depicts offspring age (weeks), while the y-axis displays the cumulative incidence of adult emergence, *F(x)*. Lines and ribbons are the posterior predictions (median ± 95% CI) for *F(x)*. The colours represent within-parent deviations in age relative to the parent’s mean age: “early-aged” = –1.96σ, “middle-aged” = mean *Δage*, “late-aged” = +1.96σ. Dashed lines represent the posterior median estimate for development time per parental age category. N = 939 offspring from 77 parent pairs.

### 3.6 Adult mass

Not only did offspring from older parents accelerate development; they may have done so at the expense of adult mass. We found strong evidence that offspring adult mass decreased with increasing parental age (*post-hoc slope* = –0.02 grams/*SD_Δage_*, 95% HDI [-0.03, –0.01 grams]; *Figure 6*), with this effect being non-negligible (*Δage*: *pd* = 99.99% & 25.96% *in ROPE (± 0.02 grams per SD_Δage_*); *Appendix S1: Figure S19 & Table S18*). Post-hoc comparisons revealed the offspring from old parents were the smallest at 0.71 grams (95% HDI [0.69, 0.74 g]), while offspring from young parents weighed, on average, 0.80 grams (95% HDI [0.77, 0.83 g]), a mean difference of 11.77% (95% HDI [5.60, 18.01%]; *Figure 3*). We found uncertain evidence to suggest this parental age effect differed between temperature treatments (*Temp x Δage*: *pd* < 95% & >33% *in ROPE*; *Appendix S1: Figure S19 & Table S18*).

**Figure 6.**
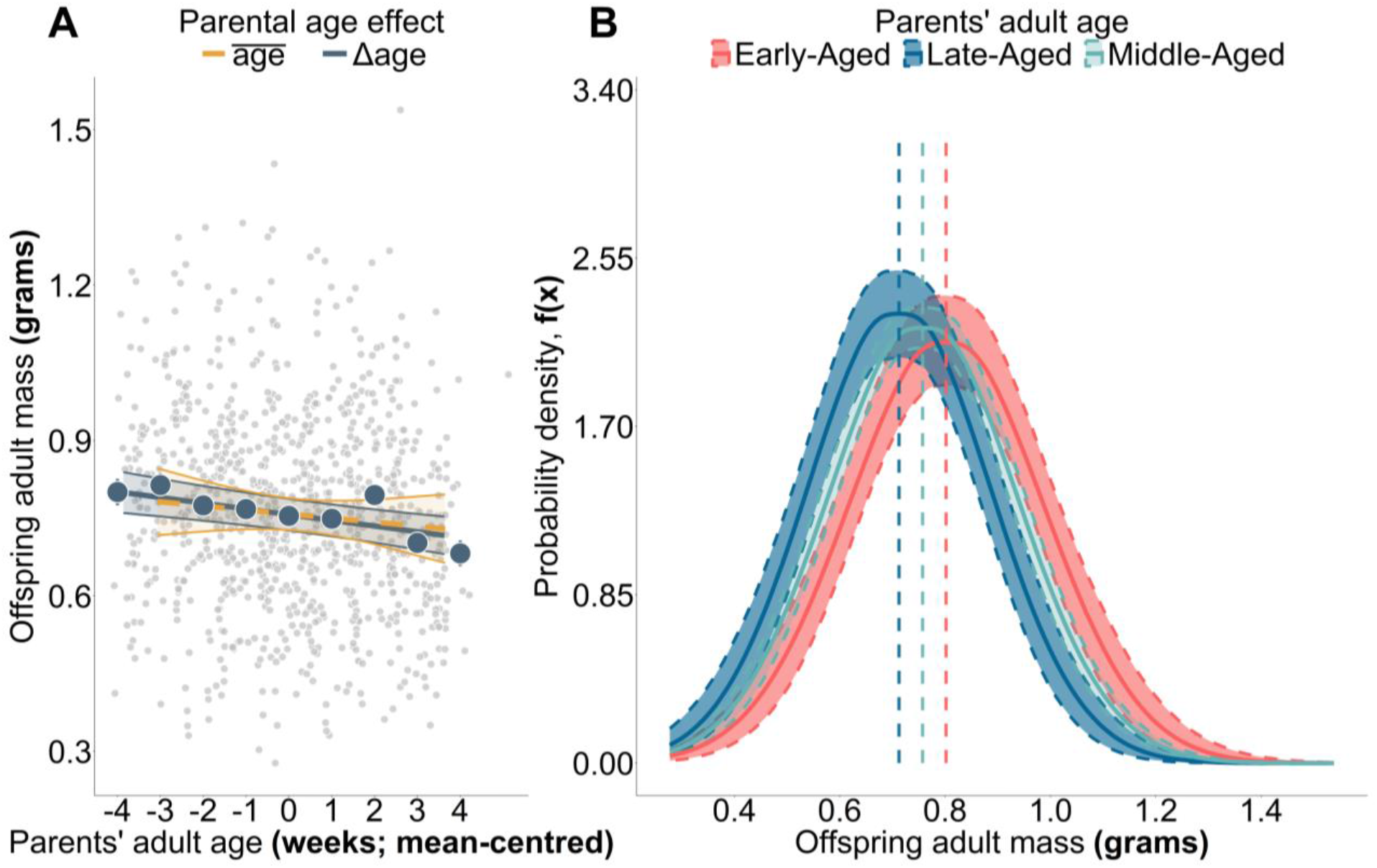
A) The effect of parental age on offspring adult mass. On the x-axis is the parents’ age at reproduction (mean-centred) and on the y-axis is offspring adult mass (g). Small dots represent raw data, large dots are raw means ± SE. Lines and ribbons represent posterior predictions for the mean effect ± 95% CI. The colours reflect the type of parental age effect. **B) The effect of parental age on the distribution of offspring adult mass (probability density).** On the x-axis is offspring adult mass (g), while the y-axis shows probability density, *f(x)*. Lines and ribbons represent posterior predictions for *f(x)*. The colours represent within-parent deviations in age relative to the parent’s mean age: “early-aged” = –1.96σ, “middle-aged” = mean Δage**, “**late-aged” = +1.96σ. The dashed lines represent the posterior median estimate for adult mass per parental age category. N = 937 offspring from 77 parent pairs.

### 3.7 Relative growth rate (RGR)

Given that increasing parental age is negatively associated with both offspring development time and adult mass, it is plausible that offspring from old parents emerged earlier in life and, consequently, at a smaller size, with no resultant change to their growth rate. However, we found no such pattern. Increasing parental age increased the offspring’s RGR in their time taken to reach adulthood (*post-hoc slope* = 0.03 week⁻¹/*SD_Δage_*, 95% HDI [0.02, 0.04 week⁻¹]; *Figure 7*), with this effect being non-negligible and receiving strong statistical support (*Δage*: *pd* = 100% & 0.06% *in ROPE (± 0.01 week⁻¹ per SD_Δage_); Appendix S1: Figure S21 & Table S20*). Despite their smaller mass, offspring from old parents exhibited an RGR of 0.82 week⁻¹ (95% HDI [0.80, 0.84 week⁻¹]), compared to the slower RGR of offspring from young parents at 0.72 week⁻¹ (95% HDI [0.70, 0.74 week⁻¹]). We found uncertain statistical support for an *Δage* by parent temperature interaction (*Temp x Δage*: *pd* < 75% & > 60% *in ROPE*; *Appendix S1: Figure S21 & Table S20)*.

**Figure 7.**
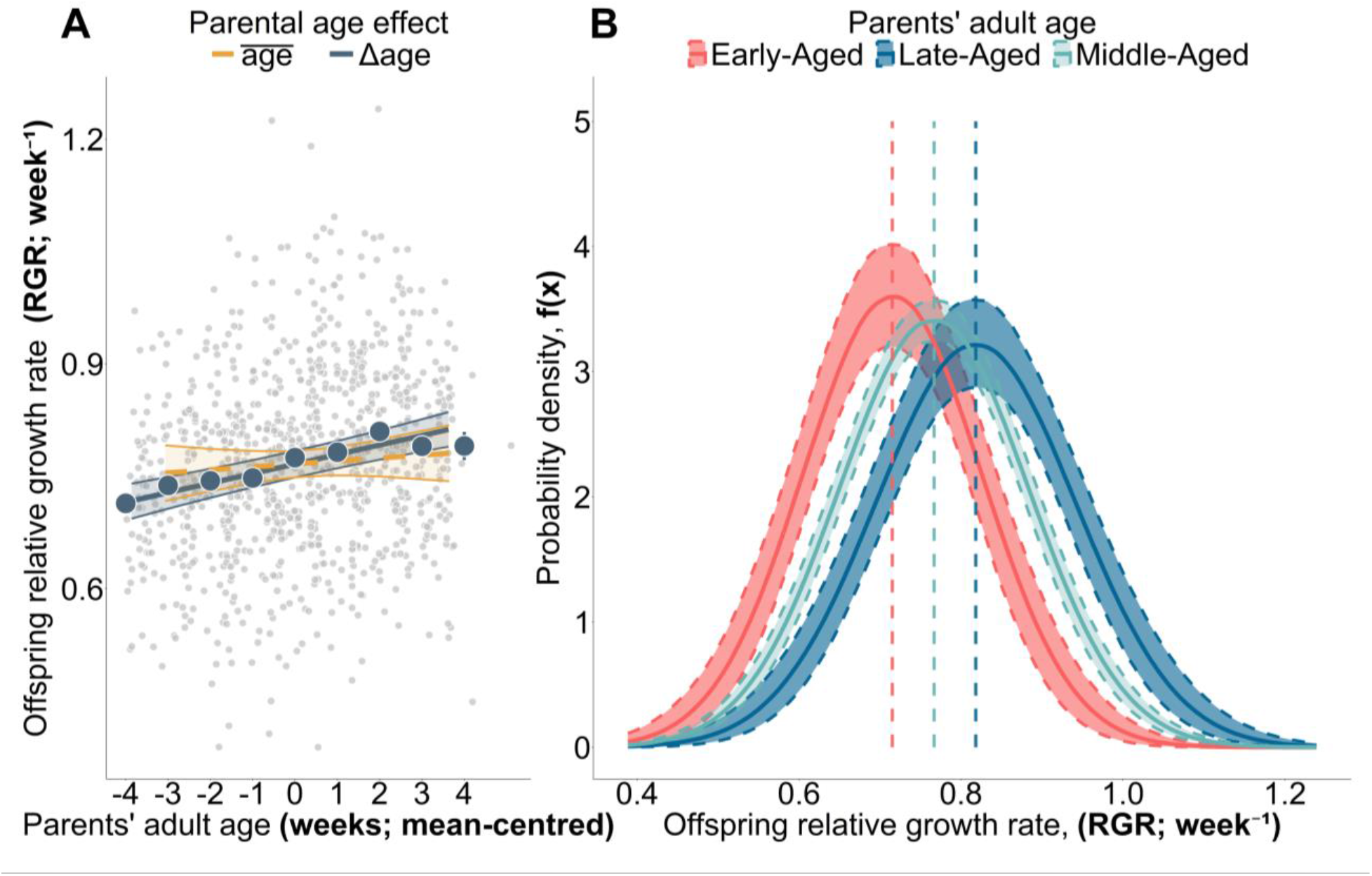
A) The effect of parental age on the relative growth rate of the offspring. On the x-axis we show the parent’s age at reproduction (mean-centred), while the y-axis displays the offspring’s RGR (week⁻¹) during their time taken to reach adulthood. The small dots show the raw data points, while the large dots indicate the raw means ± SE for the *Δage* effect. The lines and ribbons represent the posterior predictions for the mean parental age effect ± 95% CI. The colours indicate the type of parental age effect. **B) The effect of parental age on the distribution of relative mass gain (probability density).** On the x-axis is offspring’s RGR (week⁻¹), while the y-axis displays the probability density, *f(x)*. The lines and ribbons represent posterior predictions for *f(x)*. The colours represent within-parents deviations in age relative to their mean age: “early-aged” = –1.96σ, “middle-aged” = mean *Δage***, “**late-aged” = +1.96σ. The dashed lines describe the posterior median for RMR per parental age category. N = 937 offspring from 77 parent pairs.

### 3.8 Offspring reproductive success

#### 3.8.1 Fecundity

We found that increasing parental age may have a non-negligible effect on offspring fecundity, although this effect received limited statistical support (*Parental age_Late-aged_*: *pd* = 95.79% & 11.99% *in ROPE (± 0.13 log-eggs*); *Appendix S1: Figure S23 & Table S22*). Post-hoc comparisons revealed that offspring from old-aged parents laid, on average, 342 eggs (95% HDI [246, 449 eggs]), compared with the 243 eggs (95% HDI [177, 317 eggs]) produced by offspring descending from young parents (*Figure 8A*). We found uncertain statistical support for any differential effect of parental age across temperature treatments (*Temp x Parental age_Late-aged_*: *pd* < 90% & >10% *in ROPE*; *Appendix S1: Figure S23 & Table S22*).

**Figure 8.**
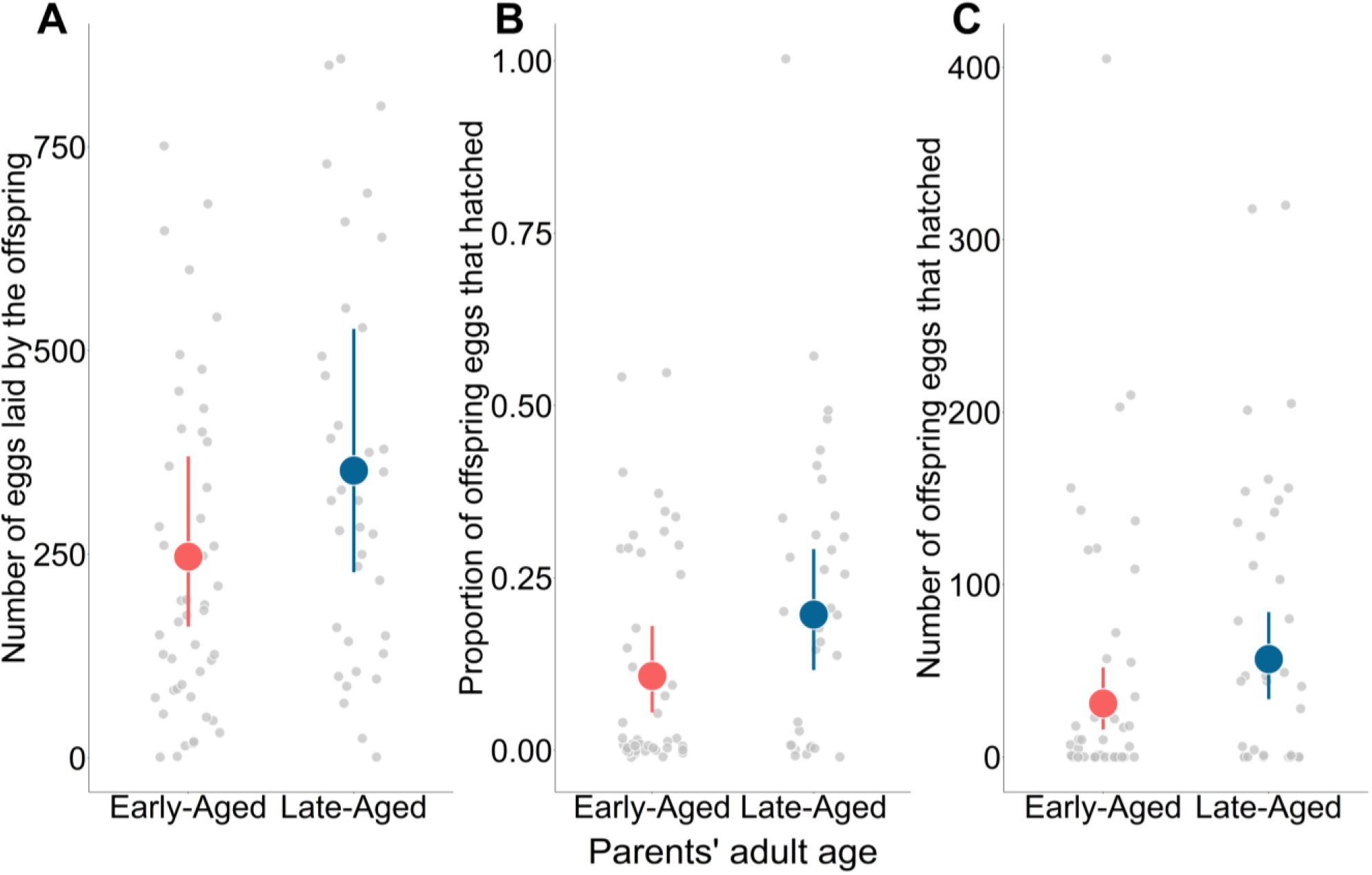
Parental age effects on offspring reproductive performance. A) The effect of parental age on offspring fecundity. On the x-axis is the parents’ age category, and on the y-axis is number of eggs laid by the offspring. Large dots represent model predictions ± 95% CI for mean offspring fecundity. Small dots are raw data. Eggs from n = 83 offspring descending from 45 parent pairs. **B) The effect of parental age on the proportion of eggs hatched by the offspring.** On the x-axis is the parents’ age, and on the y-axis is the proportion of eggs that the offspring hatched (accounting for the zero-inflation component). Large dots represent the posterior mean effect ± 95% CI for hatching success. Small dots are raw data. **C) Parental age and the number of eggs hatched by the offspring**. On the x-axis is the parents’ age category, while on the y-axis is the number of eggs that hatched. Here, we present the posterior mean predictions for number of eggs hatched, estimated as a combination of the total number of eggs laid by offspring in each group and their hatching probability. Eggs from n = 82 offspring descending from 45 parent pairs.

#### 3.8.2 Hatching success

The eggs laid by offspring from old-aged parents exhibited higher hatching success, with this effect being non-negligible and receiving moderate statistical support (*Parental age_Late-aged_*: *pd* = 99.41% & 2.74% *in ROPE* (*± 0.18 log-odds*); *Appendix S1: Figure S25 & Table S24).* Post-hoc comparisons revealed that the eggs laid by offspring from old-aged parents had a hatching probability of 0.25 (95% HDI [0.18, 0.33]), compared with 0.14 (95% HDI [0.08, 0.21]) for offspring from young parents. However, this mean estimate does not account for the zero-inflation component of the model (i.e., complete reproductive failure). Although parental age showed little association with complete hatching failure (*pd* = 56.44%; *Appendix S1: Table S24*), the inclusion of this zero-inflation component in the hatching success estimate reduced the post-hoc predicted difference between the two groups. Here, the average hatching probability dropped to 0.20 (95% HDI [0.12, 0.29]) for eggs laid by offspring from old parents and to 0.11 (95% HDI [0.05, 0.18]) for offspring of young parents (*Figure 8B*). When combining hatching probability with the total number of eggs laid, offspring from old parents may have hatched a greater number of eggs, hatching an average of 56.80 eggs (95% HDI [33.50, 84.10]), compared to the 31.10 eggs (95% HDI [15.90, 52.00 eggs]) hatched by offspring from young parents (*Figure 8C*). We found uncertain statistical evidence to suggest that this parental age effect differed across temperature treatments (*Temp x Parental age_Late-aged_*: *pd* < 80% & >18% *in ROPE*; *Appendix S1: Figure S25 & Table S24*).

### 3.9 Adult lifespan and mortality risk

Given that increasing parental age has limited effect on the offspring’s total lifespan (*Figure 3*) but accelerates their development time (*Figure 5A*), we would expect offspring from old parents to exhibit prolonged adult lifespans, opposite of a Lansing effect. We found strong evidence for this positive relationship between parental age and offspring adult lifespan, with this effect being non-negligible (*Δage*: *pd* = 100% & 5.27% *in ROPE* (*± 0.04 log-weeks per SD_Δage_*); *Appendix S1: Figure S27 & Table S26*). Here, increasing parental age functioned to extend the offspring’s adult lifespan (*post-hoc slope*: 0.55 weeks/SD*_Δage_*, 95% HDI [0.35, 0.74 weeks]), with offspring from old parents surviving for an average 10.64 weeks (95% HDI [10.14, 11.12 weeks]) as adults, compared to 8.51 weeks (95% HDI [8.11, 8.91 weeks]) for offspring from young parents (*Figure 3*). There was uncertain statistical support for a *Δage* by temperature interaction, with the parental age effect being similar across temperature treatments (*Temp x Δage*: *pd* < 94% & >50% *in ROPE*; *Appendix S1: Figure S27 & Table S26*).

For the *BaSTA* Weibull-Makeham model explaining variation in the offspring’s adult baseline and age-specific mortality parameters (i.e., *c, b_0_, & b_1_*), the best-supported model included the two-way interaction between parental age and offspring sex (DIC = –2628.82), with this performing better than the model with isolated effects of parental age (DIC = –2617.67) and the null model (DIC = –2597.37). However, our *brms* model on mean adult lifespan suggested that there was only weak evidence for a sex-specific inverse Lansing effect and that any sex differences were negligible (*Offspring sex x Δage: pd* = 97.84% & 53.03% *in ROPE*; *Appendix S1*: *Figure S27 & Table S26*). Thus, the predictive performance of this *BaSTA* mortality model is likely benefitting from the added inclusion of offspring sex as a fixed effect *(Appendix S1: Figure S30*), rather than from the interaction itself. The null model had improved performance over both the parental temperature model (DIC = –2592.04) and the model including the two-way interaction between *parental age x parental temperature* (DIC = –2595.79; *Appendix S1: Figure S29*). For simplicity, we report mortality estimates from the model with the isolated effect of parental age (*Table 2*). Offspring from young and old-aged parents entered adulthood with a similar baseline mortality (*c*). However, those from old-aged parents displayed both a later onset (*b_1_*) and slowed pace (*b_0_*) of actuarial senescence over their adult lifespan (*Table 2; Figure 9*).

**Figure 9.**
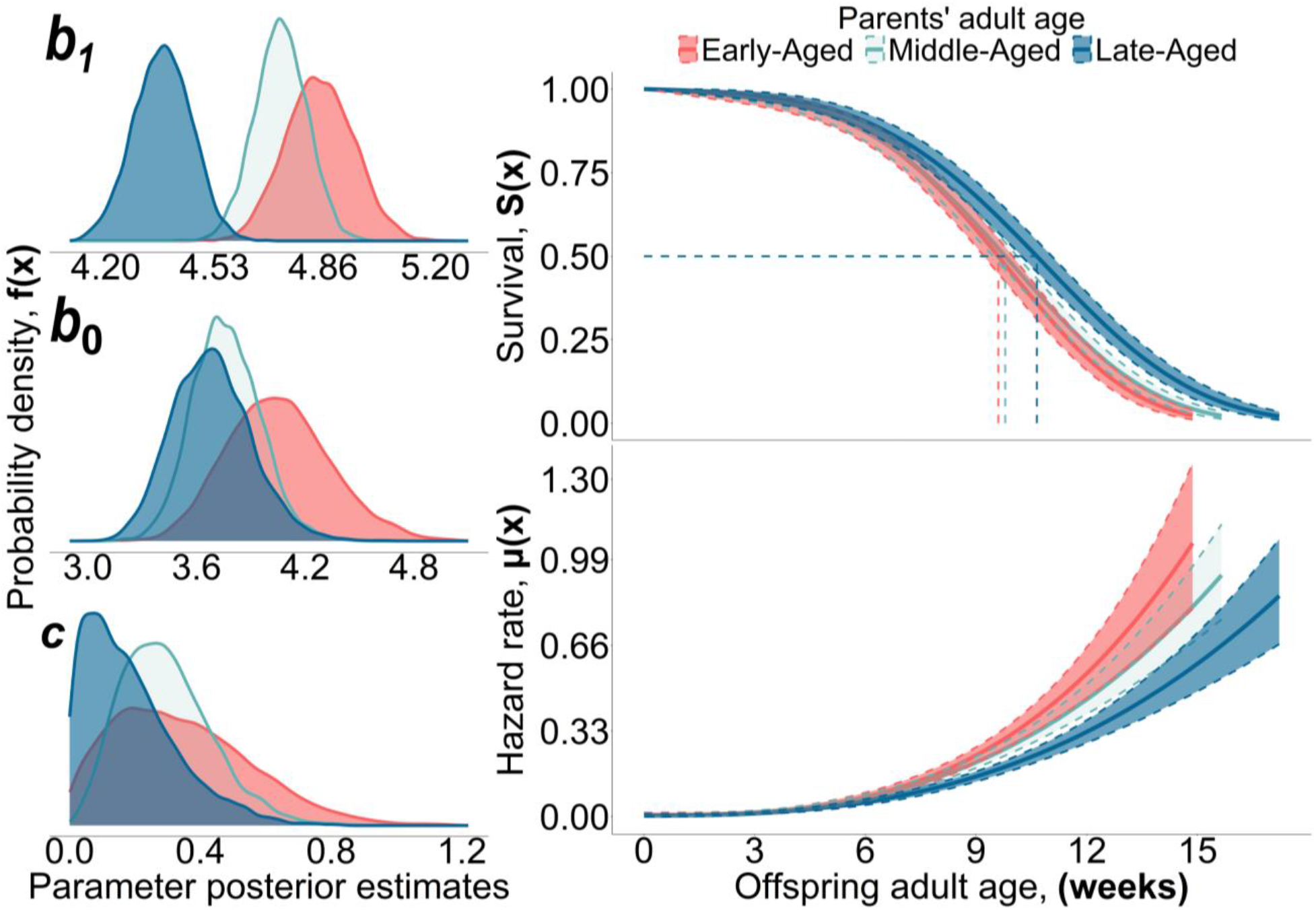
The effect of parental age on offspring adult survival and mortality. Shown on the right is the cumulative adult survival probability (top), with the median adult lifespan for offspring from each parental age group (dashed lines), and the instantaneous adult hazard rate (bottom) against the offspring’s adult age (weeks). Lines and ribbons represent posterior predictions for adult survival and mortality ± 95% CI. For visualisation, we transformed the hazard rate from the units 1/year to 1/week. The panels on the left show the full posterior distributions of the Weibull-Makeham parameters: *b*_1_, *b*_0_, and *c*. Here, *b*_1_ takes units 1/year, *c* is in years, while *b*_0_ is unitless. The colours describe the different parental age groups. N = 939 offspring from 77 parent pairs.

**Table 2.** Posterior estimates for the effect of parental age on adult mortality.

| *Mortality parameters* | *Mean estimate ± SE* | *95% CI*<br>*[lower, upper]* |  |
|---|---|---|---|
| **Makeham parameter (*c*)** |  |  |  |
| *Parental age: Early-Aged* | 0.34 ± 0.21 | [0.03, 0.81] |  |
| *Parental age: Middle-Aged* | 0.30 ± 0.14 | [0.07, 0.60] |  |
| *Parental age: Late-Aged* | 0.18 ± 0.14 | [0.01, 0.53] |  |
| **Scale parameter (*b1*)** |  |  |  |
| *Parental age: Early-Aged* | 4.85 ± 0.10 | [4.65, 5.05] |  |
| *Parental age: Middle-Aged* | 4.73 ± 0.07 | [4.57, 4.88] |  |
| *Parental age: Late-Aged* | 4.37 ± 0.09 | [4.20, 4.54] |  |
| **Shape parameter (*b0*)** |  |  |  |
| *Parental age: Early-Aged* | 4.05 ± 0.28 | [3.54, 4.65] |  |
| *Parental age: Middle-Aged* | 3.76 ± 0.18 | [3.42, 4.14] |  |
| *Parental age: Late-Aged* | 3.68 ± 0.22 | [3.29, 4.13] |  |
| ***KLDC pairwise comparisons*** |  |  |  |
| ***Parental age comparison*** | ***c*** | ***b1*** | ***b0*** |
| *Early-aged vs. Late-aged* | 0.73 | **1.00** | **0.85** |
| *Middle-aged vs. Early-aged* | 0.59 | **0.80** | **0.82** |
| *Late-aged vs. Middle-aged* | 0.61 | **0.99** | 0.55 |

*Note: Shown for each mortality parameter is the estimate (posterior mean), the standard errors, and the 95% CI. We also show the KLDC values for each mortality parameter to describe the distance in the posterior distributions between each parental age category, with estimates > 0.80 indicating genuine biological effects highlighted in bold. N = 939 offspring from 77 parent pairs.*
*Abbreviations: KLDC = Kullback-Leibler discrepancy calibration values; SE = Standard error; 95% CI = 95% Credible intervals*

### 3.10 Variability of offspring traits

For those models that included a variance component (*Table 1*), we found uncertain statistical evidence indicating that parental age affected the variability of the following traits: development time (*σ*, *Δage: pd* = 60.74%; *Figure 5B, Appendix S1*: *Figure S17 & Table S16*), adult mass (*σ*, *Δage: pd* = 76.63%; *Figure 6B, Appendix S1*: *Figure S19 & Table S18)*, growth rate (*σ*: *pd* = 86.32%; *Figure 7B, Appendix S1*: *Figure S21 & Table S20*), and fecundity (*ϕ, Δage*: *pd* = 81.22%; *Appendix S1*: *Figure S23 & Table S22*). There was weak evidence suggesting the variability of offspring hatching success decreased with parental age (*ϕ, Δage*: *pd* = 97.44%; *Appendix S1*: *Figure S25 & Table S24*). Those offspring from old parents may have exhibited improved precision in their hatching success estimates (i.e., the proportion of hatched eggs clustered towards the mean; *Figure 8B & C*).

### 3.11 Selective disappearance of parent phenotypes

We found strong evidence of selective disappearance (i.e., a difference between the *age* and *Δage* slope) for the following offspring traits: juvenile survival (Δslope *_age_*_−*Δage*_: –0.71, 95% HDI [-1.18, –0.19], *pd = 100%*), development time (Δslope *_age_*_−*Δage*_: 0.03, 95% HDI [0.02, 0.05], *pd* = 100%; *Figure 5A*), growth rate (Δslope *_age_*_−*Δage*_: –0.02, 95% HDI [-0.04, –0.01], *pd* = 99.99%; *Figure 7A*), and adult lifespan (Δslope *_age_*_−*Δage*_: –0.04, 95% HDI [-0.06, –0.01], *pd =* 98.89%). Here, the slope for the between-individual age effect progressed in the opposite direction (e.g., juvenile survival; *Appendix S1*: *Table S9*) or was flatter (e.g., development time, growth rate, and adult lifespan; *Appendix S1: Table S16, Table S20, & Table 26*) than the within-individual age estimate. Thus, if we did not control between-individual heterogeneity in the parents mean reproductive age, then this could have masked the within-parental age trajectories we observed.

## 4. Discussion

Our results question the universality of parental effect senescence and contribute to the growing knowledge that increasing parental age can have complex, but not necessarily detrimental, impacts on offspring phenotypes (Bouwhuis *et al*., 2010; Plaistow *et al*., 2015; Kroeger *et al*., 2020; Travers *et al*., 2021; Anderson *et al*., 2022). Irrespective of their parents’ thermal environment, which induced experimental variation in parental mortality rate, offspring from old parents showed no evidence for The Lansing Effect. Rather, when compared to offspring from young parents, those from old parents accelerated their development, eclosed with a smaller adult mass, potentially increased growth rate, and, crucially, displayed an extended adult lifespan, opposite of the expected Lansing Effect. This inverse Lansing Effect arose not from a reduced initial frailty, but from a delayed onset and slowed pace of actuarial senescence. Finally, we found no apparent negative effects of advanced parental age on the offspring’s own reproduction. Below, we first discuss the lack of temperature induced environmental heterogeneity in our parental age effects, before outlining the following alternate explanations for our findings: do our results indicate constraints imposed by parental senescence, or a late life shift in the parents’ reproductive strategy?

### Environmental heterogeneity and parental age effects

Although it was clear that increasing temperatures exacerbated parent mortality, we found no convincing evidence that the parents’ thermal environment mediated our parental age effects. According to the metabolic theory of ecology (Brown *et al*., 2004), warming temperatures accelerate metabolic processes in ectotherms, resulting in the excessive generation of damaging reactive oxygen species (ROS; Angilletta *et al*., 2002; Munch & Salinas, 2009; Burraco *et al*., 2020). Under chronic exposure to high temperatures, this could have increased the parent’s rate of actuarial senescence by exacerbating ROS damage to lipids, proteins, and telomeres (Mair *et al*., 2003; Conti, 2008; Flouris & Piantoni, 2015; Keil *et al*., 2015; Friesen *et al*., 2022).

Exposure to heat stress can also accelerate the progression of reproductive senescence by further reducing fertility and inducing germline damage (Walsh *et al*., 2019; Martin *et al*., 2025; Snook *et al*., 2026). In *G. bimaculatus,* thermal stress is known to impair male gamete quality (Ratz *et al*., 2024), including the quantity and viability of sperm (Gasparini *et al*., 2018), and reduce female fecundity (Behrens *et al*., 1983). Across insects, it has been argued that the thermal fertility limits (i.e., the threshold temperature where individuals become sterile) might also be lower than those for survival (i.e., the critical threshold where physiological function collapses, known as CT_max_; Angilletta *et al*., 2002; Overgaard *et al*., 2012; Walsh *et al*., 2019), so that thermal deteriorations in egg and offspring quality occur before any effects on longevity. We found evidence of the inverse; despite the observed effects on parent mortality (*Appendix S1: Figure S1*), heat stress only slightly accelerated reproductive senescence in egg length (*Appendix S1: Figure S4*) and did not govern parental effect senescence for any offspring trait.

Although we found that parents maintained under the highest temperatures displayed the shortest lifespans, this reduced longevity does not necessarily indicate an earlier onset of ageing (Péron *et al*., 2019; Ronget & Gaillard, 2020). Alternatively, temperature may have directly increased the parents’ baseline mortality, without accelerating their senescence (Bowler & Terblanche, 2008). For example, Vindenes *et al*. (2025) found that, although the longevity of four *D. magna* clones was negatively associated with increasing temperatures (from 5 to 30°C), there were no consistent effects on the *shape* of the ageing curve for mortality and fertility. Temperature instead appeared to have complex effects on both baseline and age-specific mortality, impairing longevity beyond a direct effect on ageing itself. In our study, if temperature accelerated the parents’ ageing rates then we would have expected some component of the senescent physiology of old parents to be transferred to their offspring, inducing detrimental age effects (such as The Lansing Effect). In this scenario, we should have found evidence for a temperature by parental age interaction on offspring adult age-specific mortality, which is not supported by our data. However, if The Lansing Effect is best understood as a manifestation of accelerated senescence (Kern *et al*., 2001), and if increasing temperatures only increased the parents baseline mortality, independently of senescence, then this could explain the lack of environmental variation in our parental age effects. We believe that this latter explanation best reconciles with our findings, but this hypothesis requires further experimental testing.

### Parental age as a constraint: Egg provisioning and resource allocation trade-offs

Our Inverse Lansing Effect may have arisen from late-life improvements to pre-natal provisioning. Older mothers could have displayed a terminal investment strategy (Clutton-Brock, 1984; Velando *et al*., 2006; Jehan *et al*., 2021) or improved in body condition with age (Stahlschmidt *et al*., 2013). Under this scenario, older mothers would provision their eggs with high concentrations of macro and micronutrients (e.g., lipids, carbohydrates, proteins, vitamins), thereby improving offspring survival (Clutton-Brock, 1984; Mousseau *et al*., 2009; Gilbert & Manica, 2010). Additionally, older males may package their spermatophores with high concentrations of accessory proteins and hormones (e.g., as demonstrated in *Teleogryllus oceanicus*; Rutkowski *et al*., 2023), which subsequently increases maternal pre-natal allocation of nutrients to the developing oocytes (Burpee & Sakaluk, 1993; Oberhauser, 1997; Ducatez *et al*., 2012). We suggest that a terminal investment strategy appears unlikely given the substantial evidence for reproductive senescence across insects (Begon & Parker, 1986; Fox & Czesak, 2000; Rivero *et al*., 2001; Fox *et al*., 2003; Bloch Qazi *et al*., 2017; Muller *et al*., 2017).

Likewise, we found no evidence for late-life improvements in pre-natal investment, assuming egg length is a reliable proxy of the mother’s investment (Simmons & García-González, 2007). In our system, egg length peaked at an intermediate age before deteriorating in late life; a pattern indicative of reproductive senescence (*Figure 4A*). Finally, even in instances where maternal provisioning increases with age, The Lansing Effect often remains (Plaistow *et al*., 2015; Anderson *et al*., 2022), suggesting that age-dependent improvements in maternal provisioning do not necessarily alleviate, and may even exacerbate, offspring senescence.

It is entirely possible that The Lansing Effect (or inverse thereof) can arise from the offspring’s response to age-dependent changes in maternal pre-natal provisioning, rather than from the inheritance of their old parents’ senescent phenotype (Monaghan *et al*., 2020). In this scenario, parental provisioning modulates how offspring resolve existing life-history trade-offs between the competing demands of growth, reproduction, and somatic maintenance (Stearns, 2000; Anderson *et al*., 2022). Previous research in *Daphnia pulex* found that offspring from old parents exhibited a truncated lifespan, yet this was correlated with an increased body size, accelerated growth rates, and improved early-life fecundity (Plaistow *et al.,* 2015). Similar parental age effects on body size, development, and reproductive scheduling have been demonstrated in *Caenorhabditis elegans* and *Daphnia magna* (Travers *et al.,* 2021; Anderson *et al*., 2022), being attributed to an offspring response to late-life improvements in the mothers’ egg quality. This could indicate that increased maternal provisioning initiates an alternate offspring growth trajectory in the offspring from old parents, accelerating their growth and increasing their early-life productivity at the consequence of reducing survival (Boersma & Wit, 1997; Metcalfe & Monaghan, 2003; Biro *et al*., 2006; Plaistow *et al*., 2015). Increased investment in growth or reproduction is known to exacerbate senescence, being linked to reduced antioxidant defences (Blount *et al*., 2003; Birch *et al*., 2023) increased telomere attrition (Hall *et al*., 2004; Monaghan & Haussmann, 2006; Houben *et al*., 2008; Monaghan, 2014), and heightened oxidative stress responses (Smith *et al*., 2016; Janssens & Stoks, 2020). However, the existence of these life-history trade-offs between survival and other life-history traits has been increasingly questioned, and they may not be as common as once assumed (Chang *et al*., 2024). Finally, there may be maintenance costs that scale with size, such as an increased physiological cost of moulting (Hessen and Alstad Rukke, 2000). Yet there is a major caveat to interpreting these studies relative to our findings; the species used display indeterministic growth, with maternal age and adult size being tightly correlated (Bolanowski *et al*., 1981; Glazier, 1992; Plaistow *et al*., 2015; Travers *et al*., 2021). Any improvements in pre-natal provisioning could reflect the larger size of older mothers, rather than late-life changes to provisioning. Meanwhile*, G. bimaculatus* displays deterministic growth, removing the confounding effect of *within-individual* age-dependent variation in maternal size in explaining variation in offspring phenotypes.

In our study we found an inverse pattern; offspring from old parents exhibited slowed senescence and a smaller adult mass. Could this reflect reductions in pre-natal provisioning by older mothers? Our results may hint at such a pattern; old-aged mothers appeared to lay smaller eggs when compared to those laid in their prime (*Figure 4A*). Anderson *et al*. (2022) showed that the Lansing effect can operate in both directions between clones of *D. magna* depending on how maternal provisioning changes with age, with the inverse Lansing Effect fully explained by reduced maternal lipid provisioning of the neonates. Brief periods of dietary restriction have also been widely demonstrated to extend lifespan, by slowing metabolic processes, decelerating growth rates, and reducing body size (Masoro, 2005, 2006; Spindler, 2010; Bock *et al*., 2019). Thus, our inverse Lansing Effect may not arise from parental senescence *per se*; meaning we should expect to find similar patterns in younger mothers if their egg provisioning was somehow experimentally reduced (e.g., through caloric restriction; Gribble *et al*., 2014; Hibshman, Hung and Baugh, 2016). Under this explanation, we would also expect offspring from old parents to display both reduced reproductive performance and prolonged development (Ivimey-Cook *et al*., 2023), with this relationship well established across insects (Zhang *et al*., 2019). For example, reduced egg provisioning has been frequently demonstrated to delay maturity in *Callosobruchus maculatus* (Fox, 1994; Fox *et al*., 2003; Lind *et al*., 2015). We found no evidence for this pattern as increasing parental age did not constrain offspring reproduction, and offspring from old parents accelerated, rather than slowed, their development. Finally, this increased caloric restriction does not necessarily explain the ameliorated senescence in the offspring of old parents. For instance, in *Drosphilla melanogaster*, dietary restriction does not slow actuarial senescence, instead extending longevity by lowering their baseline mortality (Mair *et al*., 2003). Alternatively, given we did not account for the offspring’s initial size at hatching, it is possible that offspring from old parents exhibited slowed growth rates and an earlier adult emergence by eclosing at a smaller body size (Vega-Trejo *et al*., 2018). Our results suggest the opposite; our measure of RGR, despite its limitations, revealed that the offspring from old parents accelerated their growth rates (*Figure 7*). This increased investment in growth is unlikely to occur if these offspring hatched from eggs receiving limited resources. Thus, it appears unlikely that reduced pre-natal provisioning explains the accelerated development and slowed senescence we observed in offspring of old parents.

It is possible that our inverse Lansing Effect is partially explained by the increased *variability* in egg length coinciding with an advanced maternal age. This increased variation in egg size may lead to the selective disappearance of offspring phenotypes within the families of old-aged mothers (Fox *et al*., 2003). In this situation, embryonic development may be slowed or completely arrested in those eggs that are smaller than the average in clutches of old females (Noguera, 2021). This selective embryonic mortality then leaves a non-random subset of high-quality offspring phenotypes in the hatchling pool of old-aged mothers, who only descend from the largest eggs (*viability selection hypothesis*; Hadfield, 2008; Mojica & Kelly, 2010). This explanation is partially supported by our finding that the initial difference in the variability of egg length between young and old mothers did not persist to influence the variability of those viable eggs on the fifth day of incubation (*Figure 4B*). It is possible that the only eggs to develop in the clutches of old mothers were a selective subset that received similar levels of investment to the eggs laid earlier in their adult life (*Figure 4B*). However, we believe viability selection is unlikely to explain our findings as the initial mortality in the first four weeks after hatching was independent of parental age (*Figure 3*), suggesting no increased mortality rate in frail individuals conceived by old parents.

### Reproductive parity and parental age

Given the known importance of reproductive history in influencing physiological condition and future reproductive investment (Bell, 1980; Gershman, 2008; Stiver & Alonzo, 2009), it may be that the changes in offspring phenotypes we observed reflect the cost of accumulated reproductive effort rather than an effect of parental age *per se* (Monaghan *et al*., 2020). In our study, we were unable to tease apart the variation explained by repeated reproduction and parental age, which covaried almost completely (*Appendix S1: Figure S2*). As such, the costs of reproduction will have been exacerbated in the oldest females, who could have mated a maximum of eight times. Elevated reproductive investment can be associated with compromised immune function (*reproduction versus immunity trade-off*: Lawniczak *et al*., 2007; Schwenke *et al*., 2016), while in constant nutrient rich environments (e.g., laboratory studies with *ad-libatum* food and water) it could be selectively advantageous to increase investment in early-reproduction, even if this incurs a late-life cost (*current versus future reproduction trade-off*; Sibly & Calow, 1986; Stearns, 2000). In both instances, increased investment in reproduction will divert resources away from somatic maintenance or incur additional damage to the germline (Edward & Chapman, 2011; Travers, Garcia-Gonzalez & Simmons, 2015; Blount *et al*., 2016; Maklakov & Immler, 2016), with these effects operating over and above the direct effects of parental ageing in our study.

Alternatively, these costs might have been offset by the additional acquisition of resources via repeated spermatophore consumption in the oldest females. Previous research in *Gryllus veletis* and *Gryllodes sigillatus* illustrated that females who consumed spermatophores daily exhibited improved longevity relative to those mated singly, displaying similar survival times to virgin females (Burpee & Sakaluk, 1993). We suggest spermatophore consumption is unlikely to offset such costs in our study; the spermatophores of male *G. bimaculatus* are comparatively small (∼0.18% of the male’s mass, while in other orthopterans these can constitute 40% of the males mass; Simmons, 1986, 1988), and the benefits of repeated spermatophore consumption may only arise in females who mate continuously through life (Wickler & Seibt, 1985; Simmons, 1988). Finally, our repeated mating design could have artificially impaired male fertility by depleting old-aged males of sperm fluid (Sanghvi *et al*., 2025). Meanwhile, females may have preferentially fertilised all their eggs with the sperm received only from the earliest mating attempt, as a single mating is thought to be sufficient to maintain lifetime reproduction in this species (Simmons, 1986). To disentangle the variation explained by repeated reproduction from parental age, it would be necessary to assign first time breeders of different ages to a mate of a standardised age (as described in Monaghan *et al*., 2020). Moreover, methods of *in-vitro* fertilisation should also be favoured to bypass confounding effects of differential investment, as arthropods can modulate their investment (i.e., yolk provisioning and content of ejaculates) depending on their partners age or condition (Zuk and Simmons, 1997; Ratz *et al*., 2024). These would be important avenues of future research on parental age effects in multivoltine species.

### Indirect parental age effects

It is plausible to speculate that our inverse Lansing Effect manifests indirectly through causal effects of parental age on adult size or development time. This is supported given numerous studies found that parental age is either positively or negatively correlated with offspring size, yet so far these effects on size seem consistently in the opposite direction to those on adult lifespan (Plaistow *et al*., 2015; Noguera, 2021; Anderson *et al.,* 2022). Likewise, those studies which found that the offspring’s adult lifespan is independent of parental age tend to find the joint absence of any effects on offspring adult size (Tregenza *et al*., 2026). This relationship has previously been described by Gershman *et al*. (2022), who suggested that parental age could either directly affect multiple offspring phenotypic traits (including adult size) independently (*Figure 10A*), or operate indirectly, entirely through a direct parental age effect on offspring development time (*Figure 10B*).

**Figure 10.**
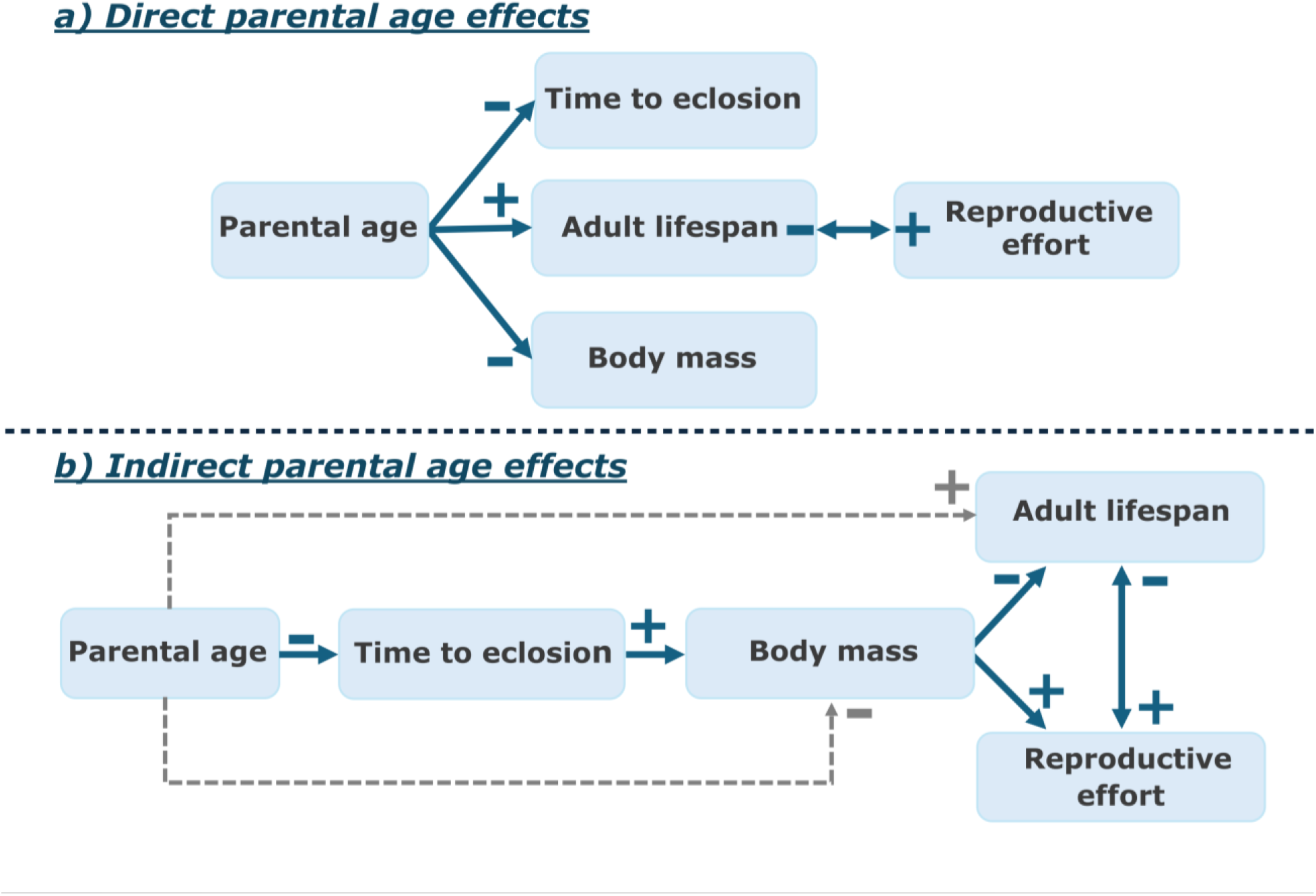
The non-mutually exclusive causal relationships between parental age, offspring development time, and offspring adult lifespan (Cause ◊ Effect). Direct effects are shown as solid blue lines, while indirect effects are shown as grey dashed lines. The sign describes the direction of the relationship: + = positive relationship, – = negative relationship. **A) Direct parental age effects.** Under this scenario, parental age directly affects all offspring phenotypic traits independently. There also exists a trade-off between reproduction and adult lifespan. **B) Indirect parental age effects on offspring adult size and lifespan, arising from causal effects of parental age on offspring development time.** Here, any effect of parental age on offspring adult size and adult lifespan operates entirely indirectly, through the direct parental age effect on the time to eclosion. Diagrams adapted from Gershman *et al*. (2022).

The pathway to causality may have progressed as follows: first, parental age could directly accelerate offspring development times, with it being adaptive for offspring from old parents to exhibit an earlier adult emergence (e.g., due to seasonal time constraints; *discussed below*). Second, this accelerated development limits the time available for somatic growth, with offspring from old parents being smaller on reaching adulthood. Indeed, this positive correlation between development time and adult size has been well-established across insects (Mousseau & Dingle, 1991; Roff, 2000). Finally, as offspring from old parents eclose with a smaller adult mass, they incur less damage from ROS during development which thereby slows senescence (*growth-lifespan trade-off*; Metcalfe and Monaghan, 2001, 2003). In this scenario, our inverse Lansing effect would arise indirectly, through causal parental age effects on offspring development time. Despite their smaller adult size, we found that offspring from old parents may have accelerated their growth rates (*Figure 7*). Under the growth-lifespan trade-off, these offspring would be expected to display the shortest lifespans, so this causal pathway may not fully explain our findings.

It is possible that, alongside their earlier adult emergence, those offspring from old parents improved their early-life fecundity, as is being increasingly evidenced (Bouwhuis *et al*., 2010; Plaistow *et al*., 2015; Travers *et al*., 2021; Dutt & Laird, 2024), but in our case such improvements were likely partially offset by their smaller adult mass. Across invertebrates, a smaller soma constrains reproduction, owing to how the ovarian tissues scale with somatic size (Honěk & Honek, 1993; Berger *et al*., 2012; Sturm, 2016). Although we found little support for a positive effect of parental age on reproduction, there was also no evidence for the inverse; that offspring reproduction was constrained by increasing parental age (*Figure 3*). Crucially, we cannot infer whether the reproductive scheduling of these offspring was shifted to increase early-life productivity, as we only measured reproductive effort from a single mating attempt at an intermediate adult age of ∼5 weeks, e.g. when the offspring were neither young nor old.

Regardless of the causal pathway, our explanation here postulates a direct effect of parental age on offspring development time. This raises the question whether it could be adaptive for the offspring from old parents to exhibit an earlier adult emergence? To answer this question, we should consider the ecology and voltinism of our study species.

### Late-life shifts in the parents’ reproductive strategy

We argue that parental age effects on the timing of adult emergence could be adaptive in seasonal multivoltine insects, who display a definitive environmental end point to reproduction. This explanation may be particularly relevant to the Mediterranean population of *G. bimaculatus*, who, unlike those found in Africa and southern Asia, cannot continuously reproduce year-round; their reproduction is completely arrested when temperatures drop beneath 16.6°C (Hoffmann, 1974; Ferreira & Ferguson, 2010). In this population, it may be beneficial for old parents to produce offspring that can accelerate their development, as older parents are bound to be more prevalent late in the season (Phelan & Frumhoff, 1991; Zehnder *et al*., 2007; Gershman *et al*., 2022). The seasonal deteriorations in environmental conditions and increasing resource scarcity place limits on the time available for older parents to complete reproduction *(“time-stress”* hypothesis; (Gotthard, 2008; Metcalfe *et al*., 2002). As *G. bimaculatus* undergoes uninterrupted development, for every directly developing generation initiated before the autumn, the length of the season suitable for growth and reproduction increasingly narrows (Nylin *et al*., 2009). In response to these time-constraints, old parents may selectively produce offspring that exhibit an earlier adult emergence (Roff, 1980; Phelan & Frumhoff, 1991; Zehnder *et al*., 2007), with these offspring accelerating their growth to not sacrifice adult size, as a larger body size is correlated with fitness in this species (Simmons, 1987a, 1987b). However, even though we found some evidence that the offspring from old parents did accelerate their growth (*Figure 7*), this was not sufficient for them to maintain the same adult mass as the offspring from young parents (*Figure 6*). Thus, the offspring from old parents may be forced to eclose as adults earlier in life at a less-than-optimal size, so they can gain some fitness returns before the season deteriorates (Roff, 1980; Nylin *et al*., 1989; Abrams *et al*., 1996; Pfenning *et al*., 2007; Larsdotter Mellström *et al*., 2010). These findings are not unexpected; previous research in *G. bimaculatus* has revealed that adult size is secondary in its net fitness contributions when compared to development time (Simmons, 1987a, 1987b), due to the greater return on reproduction in being an early emerging adult (Collins, 1980). Alternatively, offspring from old parents may encounter heightened intra-specific competition from their siblings that were produced earlier in their parent’s life, who express an age-related competitive advantage (Plaistow *et al*., 2007; Benton *et al*., 2008; Travers *et al*., 2021). This earlier emergence of offspring from young parents could create a growth race among the remaining offspring from older parents (Pásztor *et al*., 2022). Such time constraints could be avoided in those species that diapause as nymphs as they may synchronise their growth in the following year. For example, recent evidence in *G. campestris* suggests that increasing parental age has no effect on the offspring’s adult size and, subsequently, adult lifespan (Tregenza *et al.,* 2026). Despite being a close relative of *G. bimaculatus* (hybrids between the two remain fertile; Tyler *et al*., 2013), *G. campestris* is univoltine, with the nymphs undergoing an obligate diapause at their penultimate instar (Martínez-Viejo *et al*., 2024). In this species, those offspring produced later in life may not experience the same time limitations on growth as observed in the multivoltine *G. bimaculatus*. Overall, in seasonal insects that cannot arrest development, there is little fitness benefit in being a late emerging adult as this can only reduce residual reproductive value (Simmons, 1987a; Pásztor *et al*., 2022).

If development time is under strong selection, we would expect to see two patterns; not just a reduction in mean development time but reduced variability in development time in the offspring of old parents. However, despite strong effects of increasing parental age on the average development time (*Figure 5A*), there was limited evidence that the variation in offspring development time changed with parental age (*Figure 5B*). Rather, we found substantial variability in development time regardless of parental age (range = 5.86 to 17.57 weeks). This lack of canalisation could suggest there is little selection for the offspring of old parents to accelerate development. It is also conceivable that selection actively maintains variability in development time as a bet-hedging strategy, irrespective of parental age, especially if the seasonal environment varies unpredictably (Philippi & Seger, 1989; Simons & Johnston, 2006; Crean & Marshall, 2009).

### Scope for future research

As explained above, producing offspring with earlier emergence times late in the season could be adaptive given that *G. bimaculatus* is a multivoltine species, enabling another fully functional generation to be fully realized in a single year. From an evolutionary perspective, an increase in the number of generations per year facilitates rapid population growth and a greater potential for adaptation (Simmons, 1987a; Altermatt, 2010). What remains is in understanding whether these observed effects of parental age can accumulate over multiple generations to influence grandoffspring phenotypes. Previous evidence for the transgenerational inheritance of parental age effects remains ambiguous, with some finding immediate parental age effects while, in others, the effects persist across multiple generations (Bloch Qazi *et al*., 2017; Wylde *et al*., 2019; Travers *et al*., 2021; Bleu *et al*., 2022). Evolutionary theory predicts that if parental age effects do accumulate, then they may alter selection gradients on late-life reproduction (Kern *et al*., 2001; Plaistow *et al*., 2007; Moorad & Nussey, 2016; Barks & Laird, 2020). Additionally, we were unable to disentangle how paternal and maternal contributions separately impacted offspring phenotypes. It is widely known that the sexes do not age at the same rate (Bonduriansky *et al*., 2008), and previous research in *G. bimaculatus* has revealed strong sex-dependent heterogeneity in parental age effects, with these effects possibly mediated by different underlying mechanisms (Noguera, 2021). In insects, maternal and paternal age can impact the offspring differently across their ontogeny; increasing maternal age has been shown to impact traits expressed post-maturity, including adult size and adult lifespan (Noguera, 2021).

Meanwhile, paternal age often impacts offspring fitness before maturity, such as pre-adult survival and development time (Gibbs *et al*., 2010; Ducatez *et al*., 2012). Finally, given our emphasis on the importance of parental age effects in life-history evolution, it would be necessary to perform a similar longitudinal study in the relevant natural environment. Our study population was maintained under captive conditions, where no predators were present and food was provided *ad libitum*, so that two of the main drivers of life history variation were controlled for but not directly examined. The resolution of these life history trade-offs might work in different, or even opposite, ways when predators and dietary restriction are introduced into the system (Zajitschek *et al*., 2020; Promislow *et al*., 2022). Currently, little is known on the importance of parental age effects for wild insect populations, and the two studies that have investigated such effects are limited in how much they can conclude (Angell *et al*., 2022; Tregenza *et al*., 2026). Both these studies were unable to control potential sources of viability selection (including effects of parental age on early-life mortality) and only investigated cross-sectional parental age effects over a single generation. Establishing whether demographic parental age effects can accumulate across generations, operate differently between the sexes, or persist under natural environments, will be a necessary next step in establishing their ecological importance and role in the evolution of ageing.

## Author Contributions

Mark D. Pitt (MDP), Brendan O’Connor (BC), Timothy D. Sheen (TS), Tom Tregenza (TT), and Jelle J. Boonekamp (JJB) conceived and designed the study. MDP, BC, TDS, and JJB collected the data. MDP, BC, and JJB conducted the data analysis, while MDP and JJB wrote the first draft of the manuscript. All authors contributed extensively to revise the manuscript.

## Supporting information

Appendix 1: supplementary methods and results

## Acknowledgements

We would like to thank the Scottish Centre for Ecology and the Natural Environment (SCENE) for facilitating our research. Additionally, we thank Tom Ratz and Cristina Tuni for supplying us with the founding animals used to generate our colony population. We are also incredibly grateful to Lucy Winder, Phoebe Kaiser-Wilks, Maddie Inglis, Margot Gibert, Joseph Roy, and Manon Quentin for covering husbandry when additional help was required. Finally, we thank Fernando Colchero for his expertise and welcome advice on how to best utilise the *BaSTA* package for our survival analysis. This study was supported by the Natural Environment Research Council (NE/X012018/1) and by the Leverhulme Trust (RPG-2024-207). Claude Sonnet 5 was used to correct syntax errors in the R-scripts and to help develop code for plotting the model outputs, specifically for extracting and plotting the *BaSTA* outputs. The corresponding author MDP takes full responsibility over the code and has ensured it remains accurate. No AI was ever used in the writing of this manuscript.

## Ethics Statement

All work involving housing, husbandry, and care of the animals involved in this study was performed at the highest standard possible. Animals were cleaned and fed weekly, and housed individually from conception, removing welfare concerns regarding overcrowding.

