## Appendix 1: supplementary methods and results for "Parental age at reproduction accelerates offspring adult emergence with no apparent effects on reproductive success"

#### Appendix S1

### Parental age at reproduction accelerates offspring adult emergence with no apparent effects on reproductive success

Mark D. Pitt, Brendan O'Connor, Timothy D. Sheen, Davide M. Dominoni, Tom Tregenza, Jelle J. Boonekamp

#### Section S1: The effect of temperature on parent survival and mating success

##### *S1.1. Parent longevity*

To validate that our temperature manipulations successfully manipulated parent longevity, we visualised the Kaplan-Meier curves for A) the parent's duration in the study (i.e., their time spent at their set temperatures in the incubators) and B) the parents' adult survival. The time spent in the study was calculated as the difference between the experimental entry date (the date of exposure to the experimental temperatures) and the exit date (either from *death* **or** *early removal*). Not all pairs were maintained at their set temperatures until death, with those that terminated reproduction being removed from the incubators and held in a climate room at 28.5°C, due to capacity constraints. We assumed pairs had terminated reproduction after two consecutive mating attempts with no hatched eggs (which occurred for n = 13 pairs), at which point the pair (both male and female) were moved to the climate room. The termination of reproduction could have either occurred from a decline in sperm or oocyte quality, with it not being possible to disentangle the two processes. Additionally, any remaining parent animals still alive by the 26/03/2024 were removed from their set temperatures to the climate room (n = 34 pairs), to free up space in the incubators for another experiment. Individuals removed from their temperature treatments before death was observed (i.e., due to reproductive cessation or if the animal was still alive ten weeks after first mating) were subsequently right-censored in our Cox proportional hazards survival models and Kaplan-Meier curves (*Figure S1B*).

To generate all survival estimates, we used the package *survival v.3.8.3* (Therneau, 2024) and plotted the Kaplan-Meier survival curves using the package *survminer v.0.5.1* (Kassambara et al. 2025). To estimate how temperature affected the parents' survival (i.e., to assess if its effects are additive or multiplicative), we fitted both a Cox proportional hazard model, to obtain hazard ratios, and an Aalen's additive model (Aalen, 1989) to get the additive incremental increases in the hazard. Here, we fitted lifespan as the response, while temperature was included as a fixed effect. Any animals removed from their set temperatures before death was observed were right censored in both survival models.

We observed a total of 54 deaths and 102 experimental removals out of the starting pool of 156 parent animals. For parent survival, the Cox proportional hazard model revealed that our temperature manipulations successfully created experimental variation in parent longevity (*likelihood ratio test*: $X^2_{df = 2} = 31.27, p < 0.005$). Relative to parents maintained at 25.5°C, those at 28.0°C exhibited, on average, a 264.7% increased risk of death (*HR = 3.65, 95% Confidence intervals (CI) [1.53 – 8.67], p = 0.003)*, while those maintained under 30.5°C displayed a 637.4% increased mortality risk (*HR = 7.37, 95% CI [3.31, 16.43], p<0.005*). Survival also differed between the highest temperatures; parents maintained at 30.5°C had a 102% increased risk of death relative to those maintained at 28.0°C (*HR = 2.02, 95% CI [0.32, 2.20], p = 0.03*). We found no evidence that the assumption of proportional hazards was violated by temperature, with the Schoenfeld residuals being independent of time (*Global likelihood ratio test*: $X^2_{df = 2} = 0.441, p = 0.80$). This temperature effect was likely multiplicative, as the estimated slopes for each treatment group were not evenly spaced out in the Aalen's additive model. Relative to the baseline hazard at 25.5°C (*Intercept = 0.015, z = 2.83, p = 0.005*), parents maintained at 28.0°C displayed an added hazard increment of 0.04 (*z = 2.59, p = 0.01*), while those at 30.5°C exhibited an additive incremental increase in their hazard of 0.1 (z = 4.49, p <0.001).

This pattern was also confirmed based on visual inspection of the Kaplan-Meir curves; when parent animals that were removed from the study were right-censored in the Kaplan-Meier curve (*Figure S1B*), a greater number of animals exited the experiment due to an earlier observed death in the 30.5°C than 25.5°C treatment (*Figure S1B*). Out of the 54 observed deaths in the incubators, only eight occurred at 25.5°C, 15 occurred at 28.0°C, while 31 deaths occurred at 30.5°C. The pattern still held when animals that were removed from the study to the climate room were included. Parents maintained at higher temperatures exhibited evidence for an earlier experimental exit, regardless of whether this was from artificial removal (due to reproductive cessation) or death (*Figure S1A*). Parents at 25.5°C spent, on average, 8.72 ± 1.66 weeks (mean ± SD) in the study, compared to the 7.57 ± 2.10 and 7.28 ± 1.89 weeks for parents maintained at 28.0°C and 30.5°C, respectively.

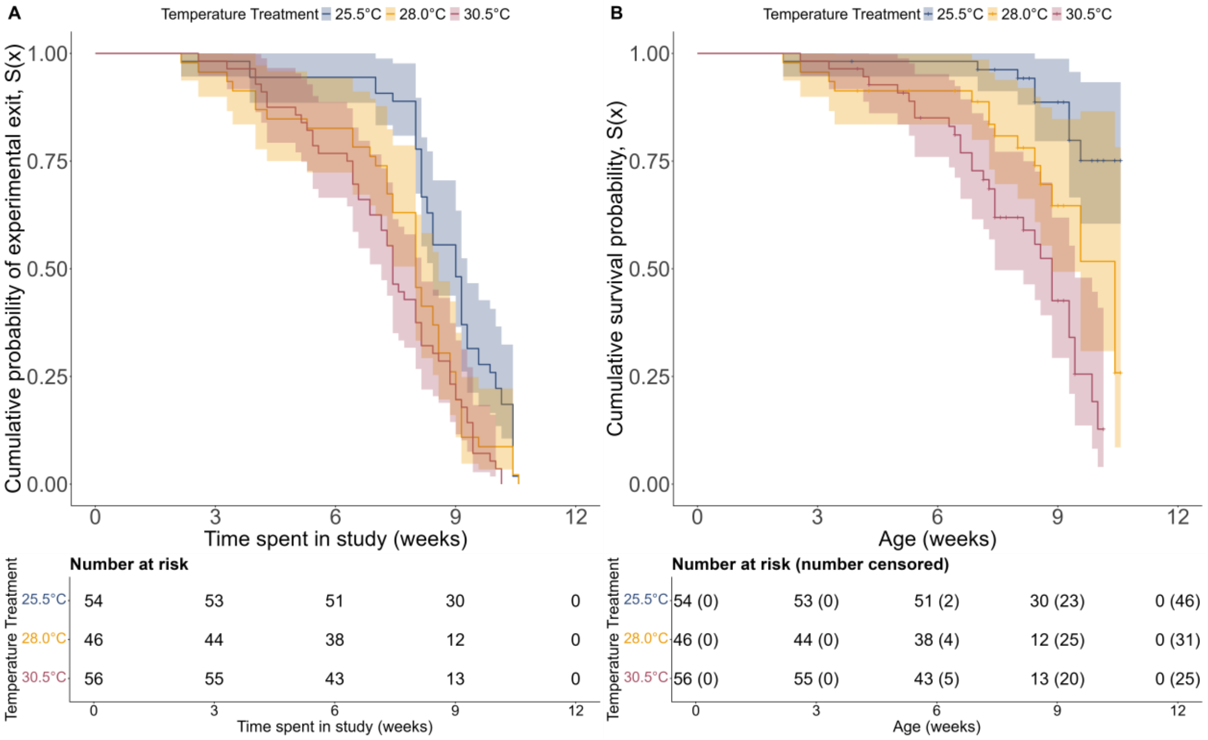

**Figure S1. Kaplan-Meir curve and risk table for parent's time in the study and adult survival under exposure to the three experimental temperatures. A) The effect of temperature on the duration in the study.** Shown is the cumulative probability that parents exited the experiment (from either death or experimental removal) plotted against the time that parent has spent in the study. **B) The effect of temperature on parent's adult survival, with right-censoring for the individuals removed from the temperature treatments before death was observed.** Shown is the cumulative survival probability plotted against the parents' age. The crosses on the Kaplan-Meier curve represent the point at which the artificially removed parents were right-censored (their time of removal). Lines and ribbons show the mean Kaplan-Meier curves ± 95% confidence intervals. The colours here reflect the three temperature treatments: blue= "25.5°C", orange = "28.0°C", and purple = "30.5°C". N = 156 parent animals (of 78 parent pairs). The table below each plot shows the number of individuals alive at a given age, while in **B)**, we also show the cumulative number of individuals censored (i.e., moved to the climate room).

##### *S1.2. Parents maximum number of successful mating attempts*

For those parents which did not die at their set temperatures during the sampling period (i.e., before their eighth mating attempt), we ran a generalised linear model in *brms* to assess whether the temperature treatments affected their number of successful mating attempts. Here, we fitted the pairs maximum number of mating attempts that resulted in hatched offspring as the response, and temperature as a fixed effect. We used the default flat priors specified by *brms* for this model, with the same model settings as outlined in our main manuscript. This analysis was conducted on the pair level (rather than individual parent level, which was used for the survival model in *S1.1*). We then compared the fit of the lognormal, Gamma, and Weibull distributions, with inspection of the posterior predictive plots revealing that the Weibull likelihood best replicated our observed data. We found that increasing temperature had negligible effect on the pairs maximum number of mating attempts, with this effect receiving limited statistical support ($β_{Temp\ x\ 28.0°C}$= -0.02, 95% HDI [-0.16, 0.11]; $β_{Temp\ x\ 30.5°C}$ = -0.13, 95% HDI [-0.26, 0.01]).

##### *S1.3. Total sample sizes*

**Table S1. Overview of sample sizes, including the number of offspring & parent pairs in each temperature treatment**. Here, we show how many offspring were produced from each mating attempt, and the number of parent pairs they descended from (in brackets). Our overall sample size includes 1317 offspring descending from 78 parent pairs.

| Mating attempt | 25.5°C | 28°C | 30.5°C | Total |
|---|---|---|---|---|
| Week 1 | 63 (27 pairs) | 54 (23 pairs) | 65 (28 pairs) | 182 (78 pairs) |
| Week 2 | 68 (24 pairs) | 57 (22 pairs) | 68 (25 pairs) | 193 (71 pairs) |
| Week 3 | 69 (27 pairs) | 58 (20 pairs) | 69 (25 pairs) | 196 (72 pairs) |
| Week 4 | 58 (23 pairs) | 48 (19 pairs) | 63 (23 pairs) | 169 (65 pairs) |
| Week 5 | 62 (21 pairs) | 56 (19 pairs) | 55 (19 pairs) | 173 (59 pairs) |
| Week 6 | 73 (25 pairs) | 47 (16 pairs) | 53 (18 pairs) | 173 (59 pairs) |
| Week 7 | 59 (21 pairs) | 39 (14 pairs) | 38 (13 pairs) | 136 (48 pairs) |
| Week 8 | 43 (16 pairs) | 35 (13 pairs) | 17 (7 pairs) | 95 (36 pairs) |
| **Total No. Offspring** | **495** | **394** | **428** | **1317** |

**Table S2. Overview of the number of offspring that reached adulthood & the number of parent pairs in each temperature treatment**. We show the number of offspring that reached adulthood from each mating attempt, alongside the number of parent pairs they descended from (in brackets). Our overall sample size includes 947 adult offspring descending from 77 unique parent pairs. Only 939 of these offspring had robustly estimated adult emergence dates.

| Mating attempt | 25.5°C | 28°C | 30.5°C | Total |
|---|---|---|---|---|
| Week 1 | 39 (21 pairs) | 38 (21 pairs) | 49 (26 pairs) | 126 (68 pairs) |
| Week 2 | 41 (23 pairs) | 37 (19 pairs) | 47 (23 pairs) | 125 (65 pairs) |
| Week 3 | 57 (26 pairs) | 41 (20 pairs) | 56 (25 pairs) | 154 (71 pairs) |
| Week 4 | 43 (23 pairs) | 37 (19 pairs) | 48 (21 pairs) | 128 (63 pairs) |
| Week 5 | 41 (19 pairs) | 37 (16 pairs) | 41 (18 pairs) | 119 (53 pairs) |
| Week 6 | 48 (21 pairs) | 39 (16 pairs) | 43 (17 pairs) | 130 (54 pairs) |
| Week 7 | 47 (21 pairs) | 27 (13 pairs) | 28 (13 pairs) | 102 (47 pairs) |
| Week 8 | 26 (14 pairs) | 23 (10 pairs) | 14 (7 pairs) | 63 (31 pairs) |
| **Total No. Offspring** | **342** | **279** | **326** | **947** |

#### Section S2: Additional experimental details

##### *2.1. Additional egg collection details*

After each mating attempt, we provided females with a small petri dish (35mm x 10mm) filled with fine sand (Children's play sand; ≤ 250 μm) as an oviposition substrate. We covered this substrate with a cotton pad dampened using tap water. Females can then insert their ovipositors through this cotton layer and oviposit their eggs into the damp sand below. To control for variable laying effort, the sand was provided to females for 15 hours across three consecutive nights, with females producing up to three batches of eggs from a single mating attempt. We provided females with sand at 17:00 on the day of mating, and eggs were collected the following morning at 09:00. To isolate the eggs from the substrate, we discarded the damp cotton layer, ensuring that any eggs attached to this cotton were returned to the sand. The oviposition substrate was then washed through a fine metal tea strainer (*HAUSPROFI Tea strainer, 304 stainless steel*), and the eggs were isolated by gently centrifuging (by hand) the egg-laden sand for ~15 seconds through tap water. This was sufficient to remove all the sand grains, while ensuring all the eggs remained in the tea strainer. We then removed the eggs from the tea strainer using a Pasteur pipette with the narrow tip cut away with scissors, which allowed the eggs to travel up to the bulb of the pipette. All eggs from a single batch were first placed into a petri dish (Greiner 6-point 60mm x 15mm) filled with tap water and photographed as a group using a digital camera mounted on a tripod (Panasonic LUMIX G7, 12-60mm lens), with a 10 x 10mm strip of graph paper for scale. These eggs were then plated onto damp cotton and incubated in the climate room (28.5 °C, 40–70% RH)). Five days into incubation, we then photographed the eggs once more, using the same setup as on the day of laying. At this point, we also rehydrated the cotton to prevent the eggs from desiccating, and the eggs remained undisturbed until we initiated the hatchling checks.

##### *2.2. Egg measurement details*

Egg length was measured by a single observer (Timothy Sheen; *TS*) using IMAGEJ. Here, using an ImageJ Macro, five eggs were randomly selected from each clutch of eggs. From each mating attempt, due to time constraints, we only photographed a single batch of eggs per female from each mating attempt, rather than measuring all eggs across the three egg batches. Egg length (*in mm*) was then measured with the line tool, using the 10mm strip of graph paper for scale. Egg length was measured on both the day of laying (hereafter *"day 0"*) and five days into incubation (hereafter, *"day five"). G. bimaculatus* eggs sharply increase in size at a single point during embryonic development, with the greatest rate of change occurring three-to-five days after laying (Donoughe and Extavour, 2016). We therefore photographed the eggs five days into incubation, assuming this represented the point where egg length peaked. We assumed that after this point that there was unlikely to be any subsequent change in eggs size until hatching. Unfortunately, it was not possible to mark the eggs without causing damage to the egg casing, and we were unable to obtain repeated measurements from a single egg on both days. As eggs were selected randomly from a clutch, it is unlikely that the same egg was measured twice during the incubation period. In total, 4216 eggs were measured from 77 parent pairs. These represent 2285 eggs that were measured on day 0, and 1931 eggs that were measured on day five.

#### Section S3: Correlation between reproductive history and age at reproduction

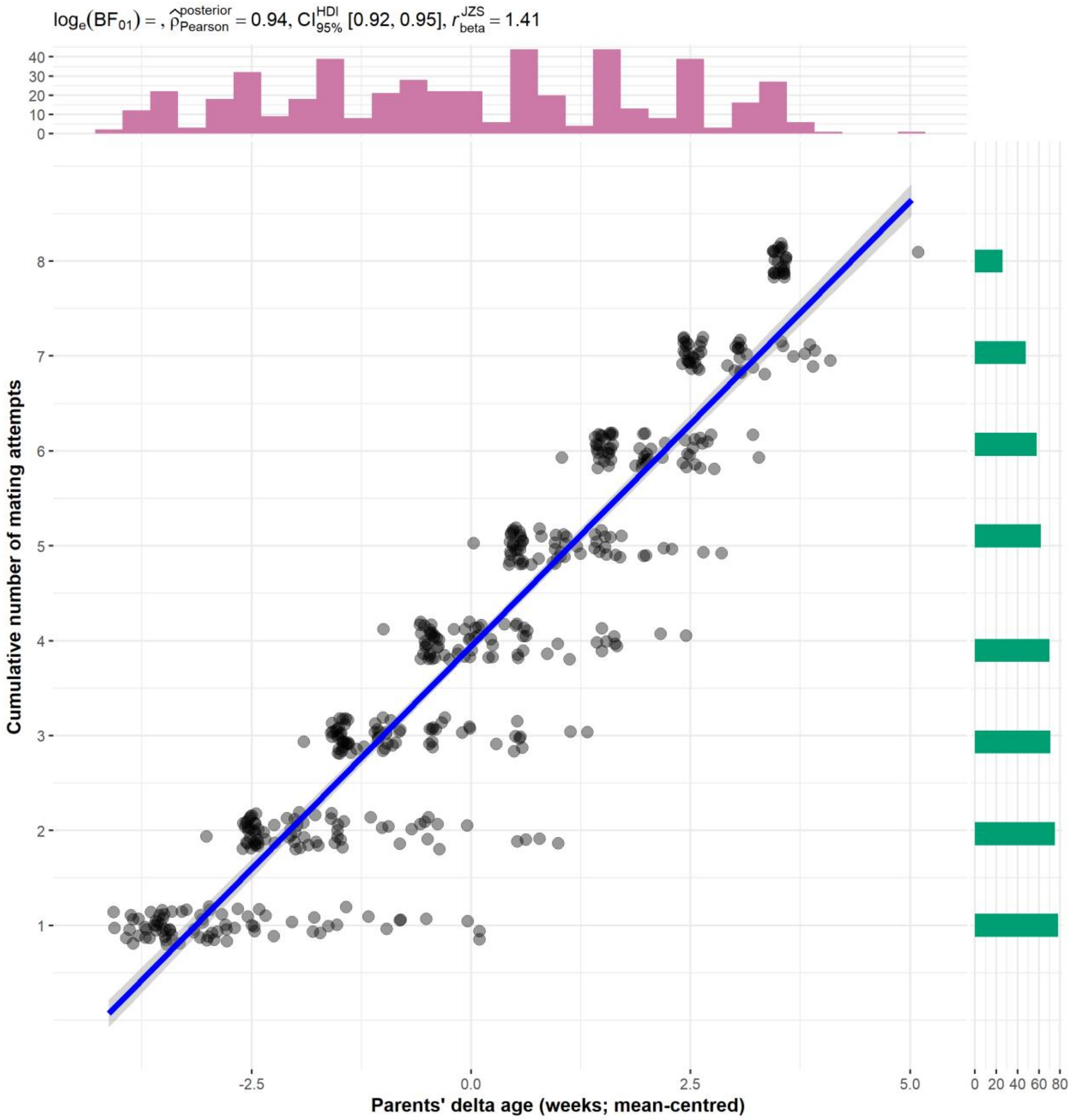

**Figure S2. Relationship between reproductive history and parents' age at reproduction.** The y-axis represents the cumulative number of mating attempts, while the x-axis shows the parent's delta age at reproduction. Each point corresponds to a single offspring collected from a single mating attempt. The solid line + ribbon shows the Bayesian Pearson's correlation between the two variables, with the estimate ± Highest Density Interval (HDI) displayed above the plot, alongside the Bayes factor. Marginal histograms illustrate the distributions of delta age at reproduction (*pink/top*) and cumulative number of successful mating attempts (*green/right*). N = 488 offspring. We filtered the data so that each pair contributed only one offspring per mating attempt, as to not skew the estimated correlation coefficient. Both the x- and y-axis are jittered by 0.1 for visualisation. Scatterplot and Pearson's correlation estimate generated with *ggscatterstats v.0.13.0*.

#### Section S4: Prior specification

We tested a range of priors for each modelled trait, from default uniform *brms* priors to mildly constraining normally distributed priors. Our approach to prior choice is outlined in the associated *R* scripts for each trait. In our final models, we selected the weakly informative normal priors (mean = 0, SD = 1) for fixed-effects ($\beta_1$) in the location component, as we had little prior assumption regarding their possible parameter values. These priors introduce mild scepticism into the model, shrinking effect sizes towards zero when sample sizes are small or when the data is noisy (Gelman et al. 2017; Lemoine, 2019). Thus, the use of these weakly informative priors makes our posterior inferences more conservative than compared to a noninformative uniform prior (i.e., the *brms* defaults), reducing the likelihood of type 1 errors (Møller & Jennions, 2002; Winter & Bürkner, 2021). The weakly informative prior assumes most ecological effects to be small and constrains any large effects arising from the data unless they are supported by high statistical power (Jennions, 2003).

Despite its widespread use, the values predicted by the $\beta_1$ ~ normal (0, 1) depend on both the scale of the response and the predictor, alongside the link-function used by the model family (Lemoine *et al.*, 2016; Lemoine, 2019). Thus, when creating GLMMS in *brms*, to ensure this prior did not become too constraining or diffuse, we z-transformed our continuous predictors, while leaving our response on the original scale. Although the safest way to use this prior is to z-transform both the predictors and response, we wanted our model inferences to be on the response, not standard deviation, scale. We therefore adapted the prior to a $\beta_1$~ normal (mean = 0, SD = 1 $SD_y$), where the $SD_y$ is the raw standard deviation of the response. This prior now assumes that a one standard deviation increase in the predictor leads to approximately one standard deviation change in the response, and rarely any larger scale changes, thus preventing the model from overestimating effect sizes (Lemoine, 2019). Priors for intercepts ($\beta_0$) of the location component were data informed, being set as $\beta_0$~ normal (mean = raw population mean, SD = raw population standard deviation). This bound the intercept towards the population mean but also permitted the intercept to explore the entire relevant biological range of the response. For random intercepts ($u_{0j}$) and slopes ($u_{1j}$) we used exponential ($\lambda$) priors (i.e., regularising priors) to stabilise the estimation of group-level variances (McElreath, 2016). These exponential priors are monotonically decreasing, placing most of the prior mass on zero, and assume that large random effect SDs are unlikely. Like the normal (0,1) prior used for fixed effects, the effect of the rate value ($\lambda$) in the exponential prior depends on the scale of the response and the link used by the model family. Generally, we set the rate in the exponential prior to allow group-level variation of up to 40% of the trait mean, though the 95% prior intervals for each prior are outlined below. We used the *r ~ LKJcorr (2)* prior for correlations between the random intercept and random slope (McElreath, 2016). In all cases, the specified priors were transformed with respect to the link-function used by the model likelihood (log-link: *Weibull, lognormal, and negative binomial*; logit-link: *Bernoulli, Zero-inflated beta-binomial*), accounting for the non-linear transformation of the log- and logit link.

For fixed effects in either the scale ($\gamma_1$) and zero-inflation ($\delta_1$) components, we used less-specific normal priors for fixed effects (*Table S1*; usually $\gamma_1/\delta_1$~ normal [0,1] or ~ normal [0,0.5]), allowing for broad effects of predictors on the variance of the trait. For models using a log-or logit link, we transformed the priors to account for the link function used (*Table S1*). Priors for the intercepts of the scale ($\gamma_0$) or zero-inflation ($\delta_0$) component were separately set for each trait, being estimated from the raw data and transformed in line with the given link function. Under the Weibull distribution, the prior for the shape parameter ($k$) was set as a half-normal prior (>0), which did not allow for any negative shape values (which are impossible under the Weibull distribution). The shape parameter was set to have a mean of 1 (constant hazard across time, assuming no senescence), but the broad standard deviation allowed for both decreasing hazards (k <1) and accelerating increases in the hazard (k>2).

**Table S3. Descriptions of the prior distributions used for models on each offspring trait.** Shown are the model likelihood and link-function used for each trait, the model parameters (left), and the prior distributions. For models using a log-link, we transformed the intercept to account for the link function, before using a normal prior distribution (equivalent to using a lognormal prior with untransformed values). Normal priors are specified by their mean and standard deviation. Exponential priors are specified by their scale and rate, while Lewandowski-Kurowicka-Joe priors (LKJ) are specified by their shape parameter. Priors were separately set for the location ($\mu$), scale ($\sigma$ or $\phi$), and zero-inflation (z) components of the model.

| ***Prior values for brms models*** | | | |
|---|---|---|---|
| ***Parameter*** | **Prior distribution** | ***Mean/scale/ shape*** | ***SD/Rate*** |
| ***Egg length: Lognormal*** *Location component ($\mu$): Identity link (of a lognormal distribution)/ Scale component ($\sigma$): log-link (on top of the lognormal distribution)* *The scale prior intercept was placed on the log of the lognormal distribution (i.e., the log of the log-transformed response SD)* | | | |
| *$\beta_0$ (Location Intercept)* | Normal | 0.99 | 0.09 |
| *$\beta_1$ (Fixed effects in the location component)* | Normal | 0 | 0.09 |
| *$\beta_{1Day: day5}$ (for egg length we expected a 10% increase in egg size at day 5 (Donoughe and Extavour, 2016), so increased the mean to 0.1)* | Normal | 0.1 | 0.09 |
| *$u_{0j}$ (random intercept)* | Exponential | 0 | 10 |
| *$u_{1j}$ (random slope)* | Exponential | 0 | 10 |
| *$r_{(u0j \sim u1j)}$ (correlation between random intercept ~ slope)* | LKJcorr | 2 | - |
| *$\gamma_0$ (Scale intercept)* | Normal | -2.42 | 0.5 |
| *$\gamma_1$ (Fixed effects in scale component)* | Normal | 0 | 0.5 |
| ***Early-life survival: Bernoulli*** *Location component ($\mu$): logit link/No scale component* | | | |
| *$\beta_0$ (Location Intercept)* | Normal | 1.10 | 1 |
| *$\beta_1$ (Fixed effects in location component)* | Normal | 0 | 1 |
| *$u_{0j}$ (random intercept)* | Exponential | 0 | 1 |
| *$u_{1j}$ (random slope)* | Exponential | 0 | 1 |
| *$r_{(u0j \sim u1j)}$ (correlation between random intercept ~ slope)* | LKJcorr | 2 | - |
| ***Juvenile survival: Bernoulli*** *Location component ($\mu$): logit link/No scale component* | | | |

| | | | |
|---|---|---|---|
| $\beta_0$ *(Location Intercept)* | Normal | 3.15 | 1 |
| $\beta_1$ *(Fixed effects in location component)* | Normal | 0 | 1 |
| $u_{0j}$ *(random intercept)* | Exponential | 0 | 1 |
| $u_{1j}$ *(random slope)* | Exponential | 0 | 1 |
| $r_{(u0j \sim u1j)}$ *(correlation between random intercept ~ slope)* | LKJcorr | 2 | - |
| ***Total lifespan: Weibull*** *Location component ($\mu$): log-link/Scale component (k): Identity link* | | | |
| $\beta_0$ *(Location Intercept)* | Normal | 2.88 | 0.21 |
| $\beta_1$ *(Fixed effects in the location component)* | Normal | 0 | 0.21 |
| $u_{0j}$ *(random intercept)* | Exponential | 0 | 10 |
| $u_{1j}$ *(random slope)* | Exponential | 0 | 10 |
| $r_{(u0j \sim u1j)}$ *(correlation between random intercept and slope)* | LKJcorr | 2 | - |
| *K (Weibull shape parameter)* | Half-normal | 1 | 2.5 |
| ***Development time: Lognormal*** *Location component ($\mu$): Identity link (of the lognormal distribution)/ Scale component ($\sigma$): log-link (of the lognormal distribution) The scale prior intercept was placed on the log of the lognormal distribution (i.e., the log of the log-transformed response SD)* | | | |
| $\beta_0$ *(Location Intercept)* | Normal | 2.21 | 0.16 |
| $\beta_1$ *(Fixed effects in the location component)* | Normal | 0 | 0.16 |
| $u_{0j}$ *(random intercept)* | Exponential | 0 | 10 |
| $u_{1j}$ *(random slope)* | Exponential | 0 | 10 |
| $r_{(u0j \sim u1j)}$ *(correlation between random intercept ~ slope)* | LKJcorr | 2 | - |
| $\gamma_0$ *(Scale intercept)* | Normal | -1.87 | 0.5 |
| $\gamma_1$ *(Fixed effects in scale component)* | Normal | 0 | 0.5 |
| ***Adult mass: Gaussian*** *Location component ($\mu$): Identity link/Scale component ($\sigma$): log-link* | | | |
| $\beta_0$ *(Location Intercept)* | Normal | 0.76 | 0.19 |
| $\beta_1$ *(Fixed effects in the location component)* | Normal | 0 | 0.19 |
| $u_{0j}$ *(random intercept)* | Exponential | 0 | 10 |
| $u_{1j}$ *(random slope)* | Exponential | 0 | 10 |
| $r_{(u0j \sim u1j)}$ *(correlation between random intercept ~ slope)* | LKJcorr | 2 | - |
| $\gamma_0$ *(Scale intercept)* | Normal | -1.66 | 0.5 |
| $\gamma_1$ *(Fixed effects in the scale component)* | Normal | 0 | 0.5 |
| ***Relative growth rate: Gaussian*** *Location component ($\mu$): Identity link/Scale component ($\sigma$): log-link* | | | |
| $\beta_0$ *(Location Intercept)* | Normal | 0.77 | 0.12 |
| $\beta_1$ *(Fixed effects in the location component)* | Normal | 0 | 0.12 |
| $u_{0j}$ *(random intercept)* | Exponential | 0 | 10 |
| $u_{1j}$ *(random slope)* | Exponential | 0 | 10 |
| $r_{(u0j \sim u1j)}$ | LKJcorr | 2 | - |

| | | | |
|---|---|---|---|
| *(correlation between random intercept ~ slope)* | | | |
| $\gamma_0$ *(Scale intercept)* | Normal | -2.10 | 0.5 |
| $\gamma_1$ *(Fixed effects in the scale component)* | Normal | 0 | 0.5 |
| ***Fecundity: Negative binomial*** | | | |
| *Location component ($\mu$): log-link/Scale component ($\phi$): log-link* | | | |
| *The intercept for the* ***scale component ($\gamma_0$)*** *was set at the empirical dispersion parameter, calculated from the raw data using the following equation from Zhang et al. (2017), solved for phi:* $\phi = \frac{\mu^2}{Var\ (Y) - \mu}$ *Where μ is the raw mean fecundity, while Var (y) is the raw variance of offspring fecundity. This estimate was then log-transformed to account for the log-link.* | | | |
| $\beta_0$ *(Location Intercept)* | Normal | 5.43 | 0.69 |
| $\beta_1$ *(Fixed effects)* | Normal | 0 | 0.69 |
| $u_{0j}$ *(random intercept)* | Exponential | 0 | 1 |
| $u_{1j}$ *(random slope)* | Exponential | 0 | 1 |
| $r_{(u0j\ \sim\ u1j)}$ *(correlation between random intercept ~ slope)* | LKJcorr | 2 | - |
| $\gamma_0$ *(Scale intercept)* | Normal | 0.503 | 1 |
| $\gamma_1$ *(Fixed effects in the scale component)* | Normal | 0 | 1 |
| ***Hatching success: Zero-inflated beta-binomial*** | | | |
| *Location component: logit-link/Scale component: log-link/Zero-inflation component: logit-link* | | | |
| *For the* ***scale component****, we used a very broad prior that centred φ on the empirical estimated dispersion parameter using the following formula (from Hardcastle & Thomas (2013), solved for phi):* $\phi = \frac{n(p(1-p) - Var\ (Y))}{n(Var(Y)\ ) - p(1-p)}$ *Where n is the mean population clutch size, p is the mean proportion of hatched eggs, and Var(y) is the variance of the proportion of hatched eggs. This empirical phi estimate was then log-transformed to set the mean estimate and given a broad SD of 1. This allowed φ to take values that were very small (substantial overdispersion) or large (little variability; close to the binomial limit where variance = mean).* | | | |
| $\beta_0$ *(Location Intercept)* | Normal | -1.64 | 1 |
| $\beta_1$ *(Fixed effects)* | Normal | 0 | 1 |
| $u_{0j}$ *(random intercept)* | Exponential | 0 | 1 |
| $u_{1j}$ *(random slope)* | Exponential | 0 | 1 |
| $r_{(u0j\ \sim\ u1j)}$ *(correlation between random intercept ~ slope)* | LKJcorr | 2 | - |
| $\gamma_0$ *(Scale intercept)* | Normal | 1 | 1 |
| $\gamma_1$ *(Fixed effects in the scale component)* | Normal | 0 | 1 |
| $\delta_0$ *(Zero-inflation intercept)* | Normal | -0.663 | 1 |
| $\delta_1$ *(Fixed effects in the zero-inflation component)* | Normal | 0 | 1 |
| ***Adult lifespan: Weibull*** | | | |
| *Location component: log-link/Scale component: Identity link* | | | |
| $\beta_0$ *(Location Intercept)* | Normal | 2.20 | 0.32 |
| $\beta_1$ *(Fixed effects)* | Normal | 0 | 0.32 |
| $u_{0j}$ *(random intercept)* | Exponential | 0 | 10 |
| $u_{1j}$ *(random slope)* | Exponential | 0 | 10 |
| $r_{(u0j\ \sim\ u1j)}$ *(correlation between random intercept ~ slope)* | LKJcorr | 2 | - |
| *K (Weibull shape parameter)* | Half-normal | 1 | 2.5 |

#### *Section S5: Additional details on statistical models*

##### *S5.1. Weibull Makeham model parameters*

The Weibull-Makeham model (Pinder, Wiener and Smith, 1978) assumes that mortality risk changes as a power function of age, describing mortality as:

$$\mu_b(x|c, b_0, b_1) = c + b_0 b_1 \,(b_1 x)^{b_0 - 1} \; (1),$$

and survival probability as:

$$S_b(x|c, b_0, b_1) = \; exp^{-cx-(b_1 x)^{b_0}} \; (2).$$

Under the three-parameter Weibull-Makeham model, $b_0$ represents the shape parameter, $b_1$ is the scale parameter, and *c* is the Makeham parameter, while $x$ denotes the individual's age. The shape parameter ($b_0$) describes the pattern mortality follows through time. When $b_0$ equals one, mortality is held constant, while when $b_0$ equals two, mortality risk increases linearly with age, and when $b_0$ is greater than two, mortality exhibits an accelerating increase over time. Meanwhile, $b_1$ determines the characteristic scale of failure times (i.e., when 63.2% of individuals have died, which we interpret as the age of onset for senescence) of the offspring (Gómez et al. 2023). *BaSTA* inverses the scale parameter ($1/b_1$), with larger values of $b_1$ corresponding to an earlier characteristic failure time. Taken together, $b_0$ and $b_1$ describe age-specific mortality; $b_1$ scales the relationship between the age of the individual ($x$) and the shape of the curve ($b_0$), setting the inflection point for when the mortality increase occurs. Finally, *c* is the age-independent baseline mortality risk (i.e., background mortality). Here, we estimated the effect of our predictors on each mortality parameter. In *BaSTA*, both *c* and $b_1$ have units in years, while $b_0$ is unitless. For further details on the parametrisation of the *BaSTA* Weibull-Makeham model, see the supplementary materials in Colchero, Jones and Rebke, 2012)

##### *S5.2. The Weibull-bathtub model*

We initially aimed to include all offspring within a single parametric survival model, which would allow us to describe how parental age impacted both offspring frailty and age-specific mortality in a single statistical framework. To this end, we fitted a Weibull-bathtub model in *BaSTA*. The bathtub term (Siler, 1979) extends the Weibull-Makeham model (*equation 1*) by adding a declining Gompertz function (Gompertz, 1825), so that mortality is expressed as:

$$\mu_b(x|c,\; a_0,\; a_1, b_0, b_1) = e^{a_0 - a_1 x} + c + b_0 b_1 \,(b_1 x)^{b_0 - 1} \; (3),$$

while the survival probability is described with:

$$S_b(x|c,\; a_0,\; a_1, b_0, b_1) = \exp[\frac{exp^{\,a_0}}{a_1}(exp^{-a_1 x} - 1) - cx] - \left(b_1 x\right)^{b_0}] \; (4),$$

where *c* is the Makeham constant, $b_0$ is the shape parameter, and $b_1$ is the scale parameter, as described in *equation 1*. Meanwhile, the additional terms $a_0$ and $a_1$ describe the declining Gompertz function used to capture early-life mortality: $a_0$ is the initial mortality risk at birth, while $a_1$ is the declining rate parameter, specified to be < 0 to allow early-life mortality to decrease with age. In this model, we included all 1317 offspring and no covariates, with this null model first being used to inspect model fit. The Weibull-bathtub model was not used for our final inference of offspring frailty or age-specific mortality, as the declining Gompertz function did not effectively capture the early-life mortality of our study population (*Figure S3*). Although the overall model performed well (Coefficient of determination $r^2 = 0.99$), this was largely driven by Weibull-Makeham component reliably capturing senescent mortality (*Figure S3*). When limiting our inspection of model fit to the early-life period (i.e., a comparison of the Kaplan Maeir and parametric survival curve in the first four weeks), the model's predictive ability deteriorated markedly ($r^2 = 0.88$).

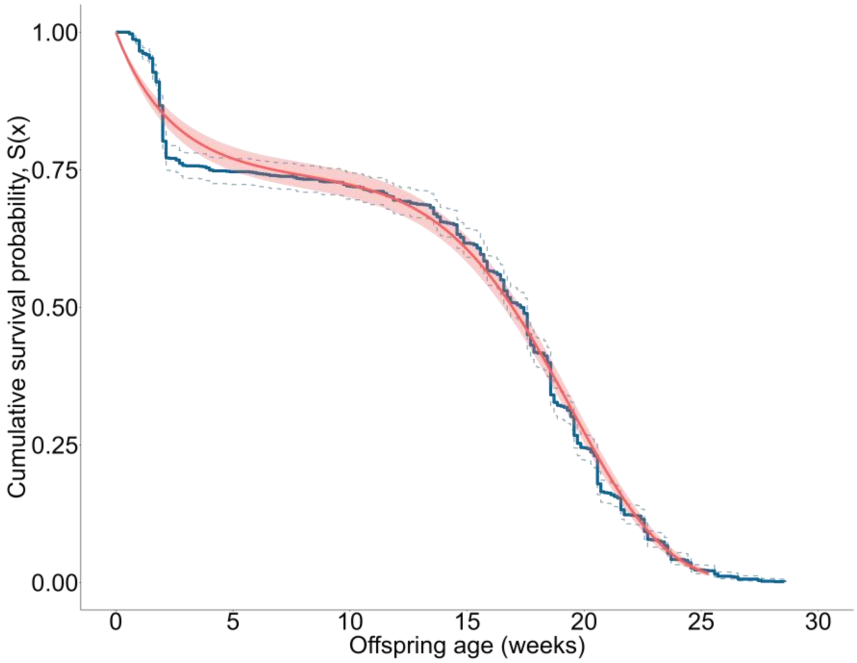

**Figure S3. Model fit for the Weibull bathtub model parameterised by *BaSTA* when we included *all* offspring in the analysis**. Shown in blue is the observed Kaplan-Meier survival curve ± 95% confidence intervals, while the parametric survival curve ± 95% credible intervals are shown in red. Based on this plot, we concluded that the Weibull bathtub parametric curve did not capture the steep early-life mortality of our study population (<4 weeks), although it robustly captured senescent mortality (the period > 4 weeks). Thus, we decided to use a mixed-modelling approach, where a Bernoulli model captured mortality occurring in the first four weeks of life. While, for animals surviving past week four and crossed the hurdle imposed by our analysis, we used a Weibull Makeham model to capture mortality across their entire lifespan (from birth until death). N = 1317 offspring from 78 parent pairs.

#### Section S6: Supplementary results for egg length

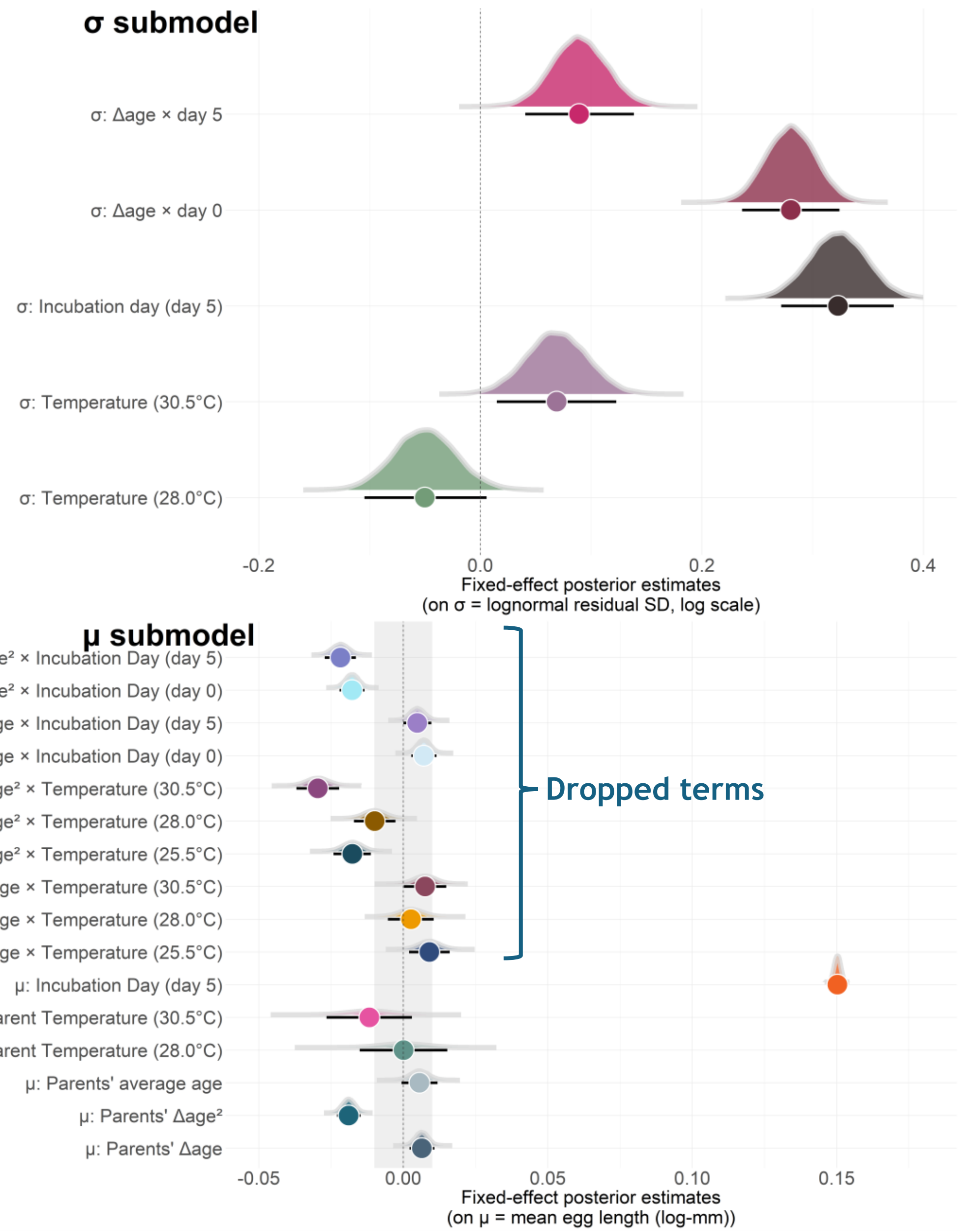

**Figure S4. Posterior distributions for fixed effects from the lognormal model explaining variation in egg length.** Here, we show fixed effects from the location (bottom) and scale (top) component separately. For each fixed effect, the circles are the posterior median while the error bars describe the 95% highest density intervals. The probability densities shown are the full posterior distributions for each fixed effect parameter. The dotted line in each model represents the null value, or the models statistical intercept (i.e., the average egg length at 25.5 °C on the day of laying). For dropped model terms we show the conditional *Δage* slopes (both the linear (*Δage)* and quadratic ($\Delta age^2$) components*)* for egg length under each temperature treatment and across incubation days. The shaded area describes the region of practical equivalence (ROPE) which represents the area of the posterior distribution practically equivalent to the null value (i.e., zero; ± 0.01 log-mm). N = 4216 eggs from 77 parent pairs.

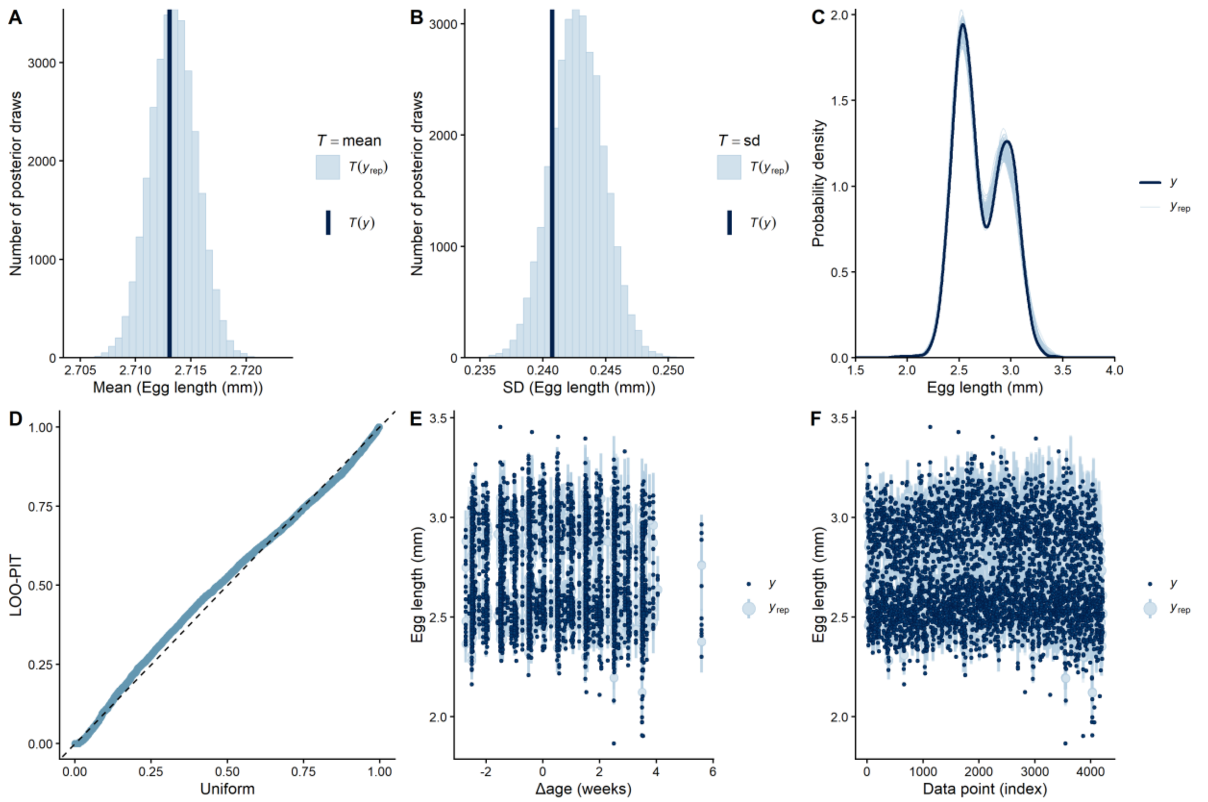

**Figure S5. Model fit plots for egg length, based on posterior simulated predictions and out-of-sample predictive performance from the selected model (*Model 13*; *Table S4*). A) and B) Histogram of means and standard deviations from the posterior predictive and observed distribution.** On the y-axis is the number of posterior draws, while the x-axis describes the mean or standard deviation of egg length. The estimated means/SD's from the posterior simulated data sets are shown in light blue ($T(Y_{rep})$), while the dark vertical line is the empirical mean/SD ($T(y)$). **C)** Probability density of the empirical and simulated data sets. On the y-axis is the probability density while the x-axis describes egg length. The light thin blue lines describe the posterior simulations of the entire distribution of the response as estimated from 100 posterior draws. The dark blue line is the observed empirical distribution of egg length. **D) Quantile-Quantile (Q-Q plot) plotting the distribution of the Leave-One-Out Probability Integrals (LOO-PIT).** This functions as a marginal predictive check *of each observation*, so that a single data point is compared only with its corresponding posterior predictive samples. To calculate the PIT of a single observation ($y_i$), the model is fitted with all the data *except* this observation ($y_{-i}$). The dashed diagonal line represents the expected uniform distribution of the LOO-PIT integrals, while the blue points represent the observed distribution of the LOO-PIT integrals. **E) Posterior predictive interval plot.** On the y-axis is egg length, while the x-axis shows the effect of *Δage*. The dark dots describe the observed data, while the light blue points and error bars show the posterior predictive estimate for the data point and the 95% credible intervals. This plot differs from **F) The LOO interval plot**. This plot describes the ability of the model to predict each observation from the LOO predictive distribution. This plot avoids "*double-dipping*" from the posterior distribution and functions as a test of the model's ability to generate each observation after being blinded to the original data point. Shown on the x-axis is each observation while on the y-axis is the estimate of the response variable (egg length). The dark points represent the observed value of each observation, while the light green points and bars represent the LOO predicted data and credible intervals (95% CI's). The observed data should fall within the credible intervals of the LOO estimates if the model is well specified. N = 421 offspring from 77 parent pairs.

**Table S4. Comparison of the out-of-sample predictive performance of models explaining variation in egg length (mm).** Highlighted in bold is the model selected from each stage of the analysis, that we then carried forward to the next stage of analysis. In this model, the following predictors were included as fixed effects: The parents' delta age at reproduction (*ΔAge*), the quadratic age term (*$\Delta Age^2$*), the parents' average age ($\overline{Age}$), the parental temperature treatment (*Temp*), the mother's adult mass (*Mothermass*), and incubation day *("day")*. We also included the mother's identity as a random intercept (*1| Female ID*) and allowed the *ΔAge* slope to vary across parent pairs by including this term as a random slope (*1 + ΔAge |Female ID)*. In the following models we show the expected log pointwise predictive density ("*elpd*"), the difference in *elpd* relative to the best-performing model ("*Δelpd*"), alongside their standard errors ("*SE*"). Models with a *Δelpd* <±4 were inferred to contain equal predictive performance to the best-supported model. **a)** Evidence for random slopes and quadratic *ΔAge* effects. **b)** Fixed effects in the scale component. **c)** Evidence for interactions between fixed effects in the *location* component. d) Evidence for interactions between fixed effects in the *scale* component. In **a)** we *always* selected the model with the random slope unless there was substantial evidence against its inclusion (i.e., if it harmed predictive performance). This final selected model is outlined in bold (*Model 13*). Across all models, n = 4216 eggs from 77 parent pairs.

| **Model selection based on predictive performance** | | |
|---|---|---|
| ***Model*** | ***elpd (± SE)*** | ***Δelpd (± SE)*** |
| ***a)* Testing support for random slopes and quadratic age effects**<br>*Using **a Lognormal** likelihood* | | |
| Model 1<br>*Egg length ~ $\overline{Age}$ + ΔAge + Temp + Mothermass + day + (1\|PairID)* | 3407.9 ± 73.2 | -237.7 ± 23.4 |
| Model 2<br>*Egg length ~ $\overline{Age}$ + ΔAge + $\Delta Age^2$ + Temp + Mothermass + day + (1\|PairID)* | 3508 ± 72.4 | -137.1 ± 17.6 |
| Model 3<br>*Egg length ~ $\overline{Age}$ + ΔAge + Temp + Mothermass + day + (1+ ΔAge\|PairID)* | 3536.1 ± 71.3 | -109.5 ± 14.6 |
| **Model 4**<br>***Egg length ~ $\overline{Age}$ + ΔAge + $\Delta Age^2$ + Temp + day + Mothermass + (1 + ΔAge\|PairID)*** | **3645.6 ± 70.9** | **0.0 ± 0.0** |
| ***b)* Evidence for fixed effects in the scale component**<br>*Using **Model 4**: Lognormal model with a quadratic term for ΔAge* | | |
| Model 5<br>σ (e*gg length*) ~1 (No fixed effects on σ) | 3645.6 ± 70.9 | -228.2 ± 26.8 |
| **Model 6**<br>**σ (*egg length*) ~ *ΔAge + Temp + day*** | **3764.3 ± 67.3** | **0.0 ± 0.0** |
| Model 7<br>σ (e*gg length*) ~ *ΔAge* | 3680.6 ± 66.2 | -83.7 ± 18.6 |
| ***c)* Evidence for interactions between fixed effects (location component)**<br>*Using **Model 6** as a reference* | | |
| **Model 6: No interactions**<br>**Location: *Egg length ~ $\overline{Age}$ + ΔAge + $\Delta Age^2$ + Temp + day + Mothermass + (1+ ΔAge\|PairID) + (1\| eggID)***<br>**Scale: σ (*egg length*) ~ *ΔAge + Temp + day*** | 3984.9 ± 67.0 | -1.1 ± 2.6 |
| Model 8: All location component interactions<br>Location: *Egg length ~ $\overline{Age}$ + ΔAge + $\Delta Age^2$ + Temp + day + Mothermass + ΔAge*Temp*day + $\Delta Age^2$*Temp*day+(1+ ΔAge\|PairID) + (1\|eggID)*<br>Scale: σ (*egg length*) ~ *ΔAge + Temp + day* | 3983.4 ± 67.0 | -2.5 ± 4.0 |
| Model 9: *Temp x ΔAge (linear slope only)*<br>Location: *Egg length ~ $\overline{Age}$ + ΔAge + Temp + day + Mothermass+ ΔAge*Temp + (1+ ΔAge\|PairID) + (1\| eggID)*<br>Scale: σ (*egg length*) ~ *ΔAge + Temp + day* | 3984.4 ± 66.9 | -1.6 ± 2.4 |
| Model 10: *Temp x ΔAge (linear + quadratic slope)* | 3986.0 ± 66.7 | 0.0 ± 0.0 |

<table>
<tr><td>Location: Egg length ~ $\overline{Age}$ + ΔAge + Temp + day + Mothermass + ΔAge*Temp + ΔAge$^2$*Temp + (1+ ΔAge|PairID) + (1| eggID)<br>Scale: σ (egg length) ~ ΔAge + Temp + day</td><td></td><td></td></tr>
<tr><td>Model 11: <u>Day x ΔAge (linear slope only)</u><br>Location: Egg length ~ $\overline{Age}$ + ΔAge + Temp + day + Mothermass + ΔAge*Temp + (1+ ΔAge|PairID) + (1| eggID)<br>Scale: σ (egg length) ~ ΔAge + day + Temp</td><td>3985.7 ± 67.3</td><td>-0.3 ± 3.3</td></tr>
<tr><td>Model 12: <u>Day x ΔAge (linear + quadratic slope slope)</u><br>Location: Egg length ~ $\overline{Age}$ + ΔAge + Temp + day + Mothermass + ΔAge*Day + ΔAge$^2$*Day + (1+ ΔAge|PairID) + (1| eggID)<br>Scale: σ (egg length) ~ ΔAge + day + Temp</td><td>3985.4 ± 67.3</td><td>-0.5 ± 3.8</td></tr>
<tr><td colspan="3">d) Evidence for interactions between fixed effects (scale component)<br>Using Model 6 as a reference for location component interactions (i.e., no interactions)</td></tr>
<tr><td>Model 11: <u>No interactions in scale component</u><br>Location: Egg length ~ $\overline{Age}$ + ΔAge + Temp + day + ΔAge$^2$ + Mothermass + (1+ ΔAge|PairID) + (1| eggID)<br>Scale: σ (egg length) ~ ΔAge + Temp + day</td><td>3984.9 ± 67.0</td><td>-14.3 ± 9.1</td></tr>
<tr><td>Model 13: Day x <u>ΔAge (in scale component)</u><br>Location: Egg length ~ $\overline{Age}$ + ΔAge + ΔAge$^2$ + Temp + day + Mothermass + (1+ ΔAge|PairID) + (1| eggID)<br>Scale: σ (egg length) ~ ΔAge + day + Temp + ΔAge*day</td><td>3999.2 ± 65.2</td><td>0.0 ± 0.0</td></tr>
<tr><td>Model 14: Temperature x <u>ΔAge (in scale component)</u><br>Location: Egg length ~ $\overline{Age}$ + ΔAge + Temp + ΔAge$^2$ + day + Mothermass + (1+ ΔAge|PairID) + (1| eggID)<br>Scale: σ (egg length) ~ ΔAge + day + Temp + ΔAge*Temp</td><td>3990.4 ± 65.9</td><td>-8.8 ± 10.2</td></tr>
</table>

**Table S5. Posterior parameter estimates from the selected model explaining variation in egg length, from the best-fitting lognormal model (*Model 13*).** The following were included as fixed effect predictors in the location component: *ΔAge, ΔAge*$^2$, $\overline{Age}$, *Temp, incubation day,* and the *mother's mass.* We also included the *Δslope* between the parents' average and delta age, as a test of selective disappearance. Female ID and Egg ID were included as random intercepts, while we also included a random slope for the effect of delta age. We additionally report the Pearson's correlation coefficient (*r*) to assess the correlation between the random intercept and random slope. In the scale component (σ), we included *ΔAge, Temp,* and *Incubation Day* as fixed effects. Additionally, in this scale submodel, there was statistical support for the interaction between *Incubation Day* and *ΔAge*. We show for each parameter the maximum a posteriori probability estimate ("*MAP*"), the 95% highest density intervals ("*95% HDI*"), along with the effective sample size ("*ESS [bulk/tail]*"), the probability of direction ("*pd*"), and the percentage of posterior estimates in the ROPE ("*% in ROPE*"). Any fixed effects with a pd > 98% are highlighted in bold, assuming these represent genuine biological effects. Finally, we also show the estimates for the dropped interaction between the parents' delta age and temperature treatment, and between delta age and incubation day. n = 4216 eggs from 77 parent pairs.

| Coefficients from the selected model (*Model 13*) | | | | | |
|---|---|---|---|---|---|
| ***Predictors*** | ***MAP*** | ***HDI (95%)*** | ***% in ROPE***[1] | ***Pd***[2] | **ESS (Bulk/Tail)** |
| ***a) Location (μ) Model*** | | | | | |
| **Intercept** *Log (Egg length [mm])* | 0.94 | [0.93, 0.95] | 0% | 100% | 9741/15634 |
| **Parents' Δage (*weeks; z-scaled*)** | **0.01** | **[0.00, 0.01]** | **87.02%** | **99.81%** | **31188/23937** |
| **Parents' Δage$^2$** | **-0.02** | **[-0.03, -0.02]** | **0%** | **100%** | **35754/25737** |

| | | | | | |
|---|---|---|---|---|---|
| **Parents' average age (*weeks; z-scaled*)** | **0.01** | **[0.00, 0.01]** | **84.88%** | **100%** | **13505/18735** |
| **$\Delta$slope$_{average\ age\ -\Delta age}$** | **0.00** | **[-0.01, 0.01]** | **-** | **58.25%** | **-** |
| Mother's adult mass (*g; z-scaled*) | 0.01 | [0.00, 0.01] | 85.03% | 97.31% | 13915/18697 |
| Incubation day | | | | | |
| *Day 0* | - | - | - | - | - |
| ***Day 5*** | **0.15** | **[0.15, 0.15]** | **0%** | **100%** | **29759/21264** |
| Parents' temperature treatment | | | | | |
| *25.5°C* | - | - | - | - | - |
| *28.0°C* | 0.00 | [-0.01, 0.02] | 74.89% | 50.74% | 10223/14335 |
| *30.5°C* | -0.01 | [-0.03, 0.01] | 34.72% | 93.89% | 9290/14588 |
| **Random Effects** | | | | | |
| σ Intercept$_{Parent\ pair\ ID}$ | 0.02 | [0.02, 0.03] | - | - | 12640/18898 |
| σ Intercept$_{EggID}$ | 0.02 | [0.01, 0.02] | - | - | 12582/20775 |
| σ Parents' $\Delta$age$_{Parent\ pair\ ID}$ | 0.01 | [0.01, 0.02] | - | - | 11750/15061 |
| *r* (Intercept ~ slope Δage) | 0.22 | [-0.13, 0.50] | - | 88.7% | 17850/19784 |
| ***Temperature Δage slopes and pairwise comparisons (from the dropped Parents' Δage x Temperature interactions on the linear and quadratic age slope)*** | | | | | |
| ***Δage*: *25.5°C*** | **0.01** | **[0.00, 0.02]** | **47.94%** | **99.43%** | **-** |
| *Δage*: *28.0°C* | 0.00 | [-0.01, 0.01] | 93.49% | 74.79% | - |
| *Δage*: *30.5°C* | 0.01 | [0.00, 0.02] | 63.71% | 97.8% | - |
| *Δage*: *25.5°C vs. 28.0°C* | -0.01 | [-0.02, 0.00] | 67.54% | 88.34% | - |
| *Δage*: *28.0°C vs. 30.5°C* | -0.01 | [-0.02, 0.01] | 75.87% | 81.44% | - |
| *Δage*: *30.5°C vs. 25.5°C* | 0.00 | [-0.01, 0.01] | 89.96% | 61.49% | - |
| **$\Delta age^2$: *25.5°C*** | **-0.02** | **[-0.02, -0.01]** | **0.37%** | **100%** | **-** |
| **$\Delta age^2$: *28.0°C*** | **-0.01** | **[-0.02, -0.00]** | **38.68%** | **99.61%** | **-** |
| **$\Delta age^2$: *30.5°C*** | **-0.03** | **[-0.04, -0.02]** | **0%** | **100%** | **-** |
| $\Delta age^2$: *25.5°C vs. 28.0°C* | -0.01 | [-0.04, 0.02] | 58.68% | 94.28% | - |
| **$\Delta age^2$: *28.0°C vs. 30.5°C*** | **0.02** | **[0.01, 0.03]** | **1.99%** | **100%** | **-** |
| $\Delta age^2$: *30.5°C vs. 25.5°C* | 0.00 | [-0.01, 0.01] | 26.97% | 61.49% | - |
| ***Incubation day pairwise comparisons for the Δage slope (from the dropped Parents' Δage x Incubation day interactions on the linear and quadratic age slope)*** | | | | | |
| ***Δage*: *day 0*** | **0.01** | **[0.00, 0.01]** | **78.65%** | **99.93%** | **-** |

| | | | | | |
|---|---|---|---|---|---|
| *Δage*: *day 5* | 0.00 | [0.00, 0.01] | 94.56% | 97.62% | - |
| *Δage*: *day 0 vs. day 5* | 0.00 | [-0.01, 0.01] | 99.98% | 90.96% | - |
| **Δage$^2$: *day 0*** | -0.02 | [-0.02, -0.01] | 0.01% | 100% | - |
| **Δage$^2$: *day 5*** | -0.02 | [-0.03, -0.02] | 0% | 100% | - |
| **Δage$^2$:** *day 0 vs. day 5* | 0.00 | [-0.01, 0.00] | 97.49% | 95.00% | - |
| ***b) Scale (σ) Model*** | | | | | |
| **Intercept** *Log (residual SD)* | -3.60 | [-3.64, -3.56] | - | 100% | 9741/15634 |
| Incubation day | | | | | |
| *Day 0* | - | - | - | - | - |
| ***Day 5*** | **0.33** | **[0.27, 0.37]** | **-** | **100%** | **-** |
| Temp | | | | | |
| *25.5°C* | - | - | - | - | - |
| *28.0°C* | -0.05 | [-0.11, 0.01] | - | 96.12% | 48112/24580 |
| ***30.5°C*** | **0.06** | **[0.02, 0.12]** | **-** | **99.39%** | **44819/24075** |
| ***Δage*** **Incubation day ~ day 0** | **0.28** | **[0.24, 0.32]** | **-** | **100%** | **29759/21264** |
| ***Δage*** **Incubation day~ day 5** | **0.09** | **[0.04, 0.14]** | **-** | **99.98%** | **60328/25606** |
| ***Δage: day 0 vs. day 5*** | **-0.19** | **[-0.26. -0.13]** | **-** | **100%** | **44845/23819** |
| **Marginal $R^2$** | 0.73 | [0.73, 0.74] | - | - | - |
| **Conditional $R^2$** | 0.85 | [0.84, 0.85] | - | - | - |

[1]ROPE = [−0.01, 0.01] on log scale; applied to location fixed effects only.

[2]pd = probability of direction.

#### Section S7: Supplementary results for offspring early-life survival

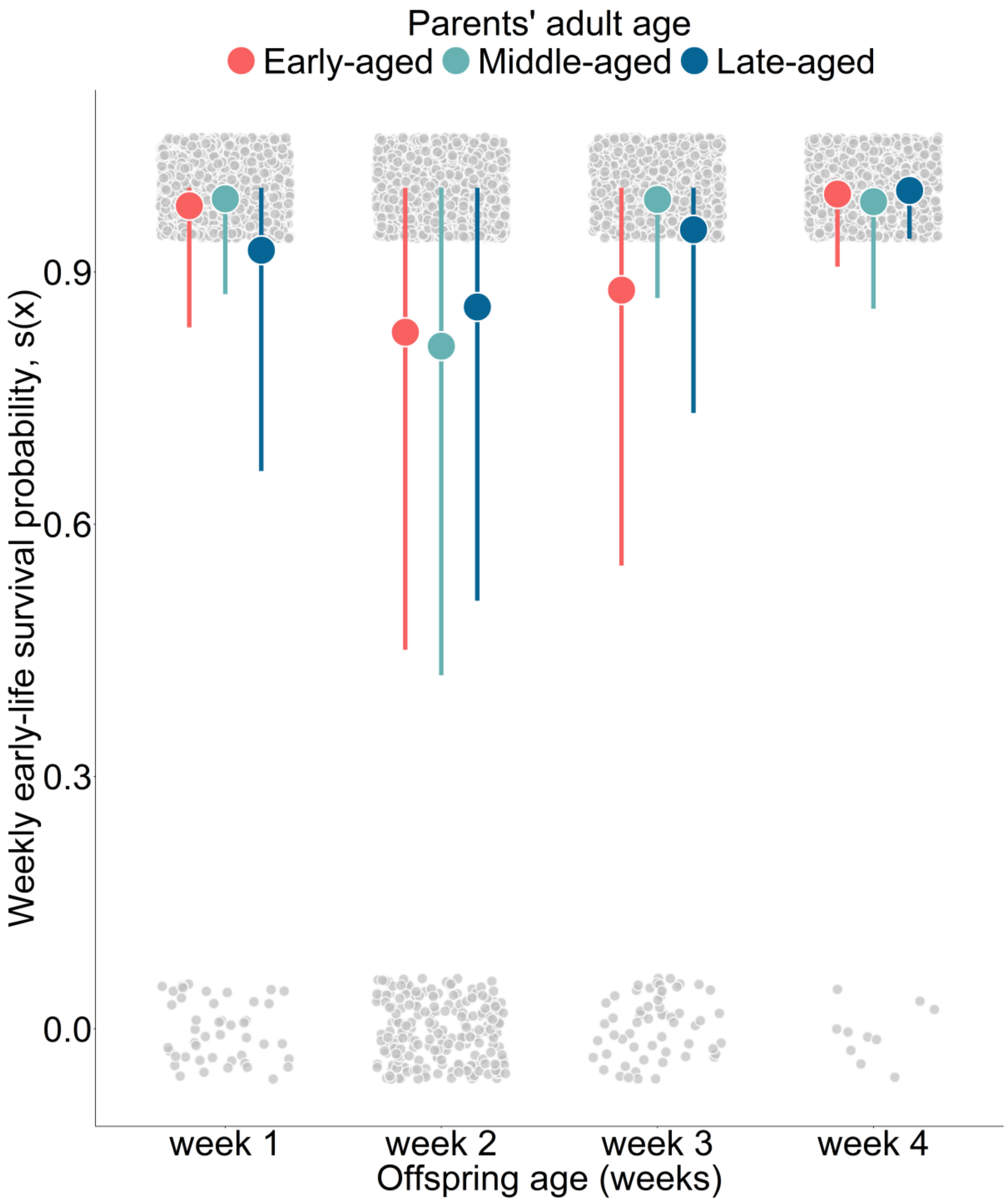

**Figure S6. Empirical weekly survival probabilities in the early-life window (weeks 0-4 after hatching).** The circles describe the raw mean survival probabilities (per week, not cumulative), while the error bars describe the empirical standard deviations. Small points describe the observed survival for the individual (binary response: survived = 1, died = 0). The colours describe the different parental age categories: red = "early-aged" (parent mating attempts 1 & 2), green = "middle-aged" (parent mating attempts 3 – 5), and blue = "late-aged" (parent mating attempts 6 – 8). From this plot we inferred that there was little difference in offspring weekly survival with parental age *within* this early-life frailty window. Our logistic regression on early-life survival is thus based on the cumulative number of deaths over these first four weeks. N = 1317 offspring from 78 parent pairs

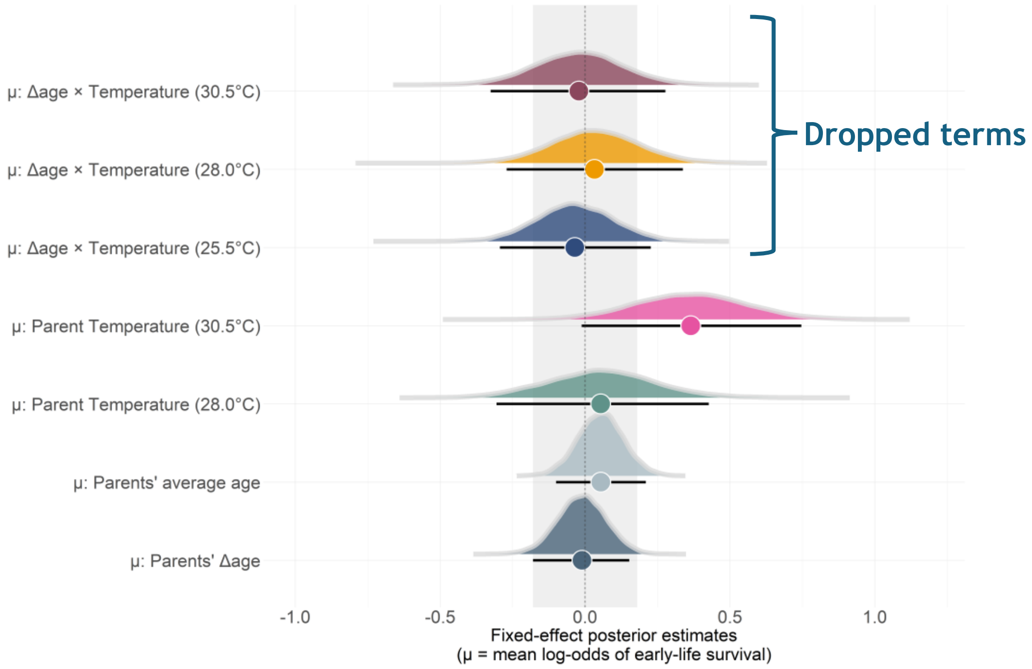

**Figure S7. Posterior distributions for fixed effects from the Bernoulli model explaining variation in offspring early-life survival.** For each fixed effect, circles represent the posterior median, while error bars are the 95% Highest density intervals. The probability densities represent the entire posterior distribution for the effect. The dotted line represents the line of no effect (i.e., the statistical intercept, which here is average early-life survival when Δage = 0 at the model's reference temperature (25.5°C)). Meanwhile, the shaded area is the region of practical equivalence (ROPE), which represents the area of negligible effect (± 0.18 log-odds). Although the *parents' temperature x Δage* interaction was removed from our final model, we show the conditional slopes for the Δage under the three temperature treatments. N = 1317 offspring from 78 pairs.

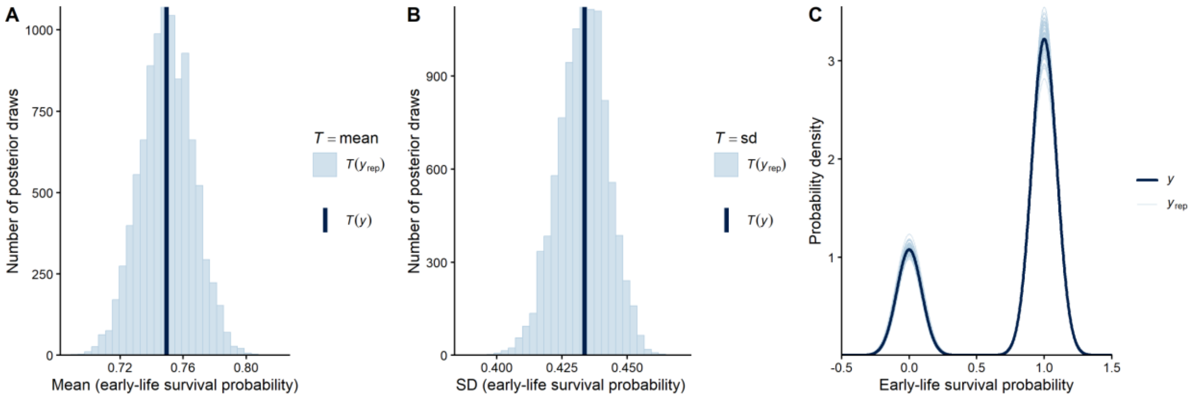

**Figure S8. Model fit plots for early-life survival.** Unlike the model fit plots for the other offspring traits (which used more complex LOO QQ & predictive interval plots), for the Bernoulli models of offspring early-life and juvenile survival, we only based inspection of model fit on the posterior predictive distribution plots. **A)** and **B)** show the posterior predictive and empirical mean and standard deviation, respectively. On the y-axis is the number of posterior draws, while the x-axis is either the mean or standard deviation of early-life survival. The dark line represents the observed mean or SD, while the light blue lines denote the posterior predictive test statistic. **C)** Describes the probability density of early-life survival. The dark blue lines represent the observed distribution of early-life failures (i.e., 0) or successes (i.e., 1), while the light blue lines describe estimates from 100 posterior draws.

**Table S6. Comparison of the predictive performance of the Bernoulli models explaining variation in offspring early-life survival (*EarlySurv*).** Here, models included the following fixed effects: The parents' delta age at reproduction (*ΔAge*), the parents' average age at reproduction $\overline{(Age)}$, the parents' temperature treatment (*Temp*), alongside the quadratic effect of delta age ($\Delta Age^2$).

Additionally, we included parent pair ID as a random intercept (*1|Pair ID*) and *ΔAge* as a random slope (*1 + ΔAge |Pair ID*). The model selected from each stage of the analysis is highlighted in bold, which was then carried forward as a reference model for the next stage of the analysis. For each model, shown are the expected log pointwise predictive densities ("*elpd*"), the Δelpd relative to the best-fitting model, along with their standard errors *("SE")*. Any models with a Δelpd <±4 were said to have identical predictive performance to the best-fitting model. If models had similar predictive performance, then we selected the simplest model to explain variation in the response (*i.e, the model with no interactions)*. **a)** Testing the evidence for random slopes and quadratic *ΔAge* effects. **b)** Evidence for interactions between fixed effects. In ***a)*** we chose the model with the random slope unless there existed substantial evidence that this harmed predictive performance. In ***b)*** If models had comparable performance, then we selected the simplest model to explain variation in the response (*i.e, the model with no interactions)*. The final model used to explain variation in early life survival was **model 5,** which only included single effect predictors. In all models, n = 1317 offspring from 78 parent pairs.

| Model selection | | |
|---|---|---|
| ***Model*** | ***elpd (± SE)*** | ***Δelpd ± SE*** |
| ***a) Support for random slopes and quadratic age effects*** | | |
| Model 1<br>*EarlySurv~* $\overline{Age}$ *+ ΔAge + Temp + (1\|PairID)* | -741.4 ± 17.6 | -3.2 ± 3.9 |
| Model 2<br>*EarlySurv~* $\overline{Age}$ *+ ΔAge +* $\Delta Age^2$ *+ Temp + (1\|PairID)* | -740.1 ± 17.7 | -2.9 ± 3.2 |
| **Model 3**<br>***EarlySurv~*** $\overline{Age}$ ***+ ΔAge + Temp + (1 + ΔAge\|PairID)*** | **-737.7 ± 17.9** | **-0.4 ± 1.8** |
| Model 4<br>*EarlySurv~* $\overline{Age}$*+ ΔAge +* $\Delta Age^2$ *+ Temp + (1 + ΔAge\|PairID)* | -737.2 ± 17.9 | 0.0 ± 0.0 |
| ***b)*** **Evidence for interactions between fixed effects**<br>*Using Model 4: Bernoulli model with single-effect predictors* | | |
| ***Model 5***<br>***EarlySurv~*** $\overline{Age}$ ***+ ΔAge + Temp + (1 + ΔAge\|PairID)*** | **-737.4 ± 17.9** | **0.0 ± 0.0** |
| Model 6<br>*EarlySurv~* $\overline{Age}$ *+ ΔAge + Temp + (ΔAge* Temp) + (1 + ΔAge\|PairID)* | -738.7 ± 18.0 | -1.4 ± 0.4 |

**Table S7. Coefficients explaining variation in offspring early-life survival from the selected model *(Model 5 from Table S6)*.** The following predictors were included as fixed effects: the parents' delta age at reproduction, the parents' average age at reproduction, and the parents' temperature treatment. Additionally, we also included the Δslope between the parents' average and delta age, which was used as a test for selective disappearance. Parent pair ID was included as a random intercept, while we fitted a random slope for the parents' delta age. We also report the Pearson's correlation coefficient (*r*) to assess correlations between the random intercept and random slope. Shown is the maximum a posteriori probability estimate ("*MAP*"), the 95% highest density intervals ("*HDI*"), along with the effective sample size ("*ESS*" [bulk/tail]), the probability of direction ("*pd*") and the percentage of posterior estimates within ROPE ("*% in ROPE*"). Any fixed effects with a pd > 98% are highlighted in bold, assuming these represent genuine biological effects. Finally, we show the estimates for the dropped interaction between the parents' temperature treatment and parent's delta age. n = 1317 offspring from 78 parent pairs.

| Coefficients from the selected model (*Model 5*) | | | | | |
|---|---|---|---|---|---|
| ***Predictors*** | ***MAP*** | ***HDI (95%)*** | ***% in ROPE***[1] | ***pd***[2] | **ESS (Bulk/Tail)** |
| ***a) Location (μ) Model*** | | | | | |
| **Intercept**<br>*Log odds of early-life survival* | 1.01 | [0.77, 1.27] | 0% | 100% | 6368/7135 |

| | | | | | |
|---|---|---|---|---|---|
| Parents' Δage *(weeks; z-scaled)* | 0.00 | [-0.17, 0.16] | 96.51% | 54.95% | 7413/6172 |
| Parents' average age *(weeks; z-scaled)* | 0.06 | [-0.10, 0.21] | 94.09% | 75.40% | 7413/6172 |
| $\Delta slope_{average\ age\ -\Delta age}$ | 0.06 | [-0.15, 0.31] | - | 71.73% | - |
| Parents' temperature treatment | | | | | |
| *25.5°C* | - | - | - | - | - |
| *28.0°C* | 0.05 | [-0.31, 0.42] | 64.60% | 62.26% | 7372/7436 |
| *30.5°C* | 0.39 | [-0.01, 0.75] | 16.88% | 96.97% | 7270/7603 |
| **Random Effects** | | | | | |
| $\sigma$ Intercept $_{Parent\ pair\ ID}$ | 0.36 | [0.04, 0.58] | - | 100% | 1624/1385 |
| $\sigma$ Parents' $\Delta age_{Parent\ pair\ ID}$ | 0.41 | [0.11, 0.64] | - | 100% | 1696/1125 |
| *r* (Intercept ~ slope Δage) | -0.09 | [-0.73, 0.60] | - | 60.17% | 2239/3070 |
| ***Temperature Δage slopes and pairwise comparisons (from the dropped Parents' Δage x Temperature interaction)*** | | | | | |
| *Δage*: *25.5°C* | -0.04 | [-0.28, 0.24] | 85.24% | 60.76% | 6180/6825 |
| *Δage*: *28.0°C* | 0.02 | [-0.27, 0.34] | 75.32% | 58.56% | - |
| *Δage*: *30.5°C* | -0.02 | [-0.32, 0.29] | 76.13% | 55.68% | - |
| *Δage*: *28.0°C vs. 25.5°C* | 0.09 | [-0.33, 0.47] | 60.06% | 62.47% | 6530/6258 |
| *Δage*: *28.0°C vs. 30.5°C* | 0.04 | [-0.38, 0.47] | 59.21% | 60.5% | - |
| *Δage*: *30.5°C vs. 25.5°C* | 0.05 | [-0.38, 0.42] | 62.81% | 52.81% | 6988/5970 |
| **Marginal $R^2$** | 0.005 | [0, 0.02] | - | - | - |
| **Conditional $R^2$** | 0.05 | [0.02, 0.09] | - | - | - |

[1]ROPE = [−0.18, 0.18] on log-odds scale; applied to location fixed effects only.

[2]pd = probability of direction.

#### Section S8: Supplementary results for offspring juvenile survival

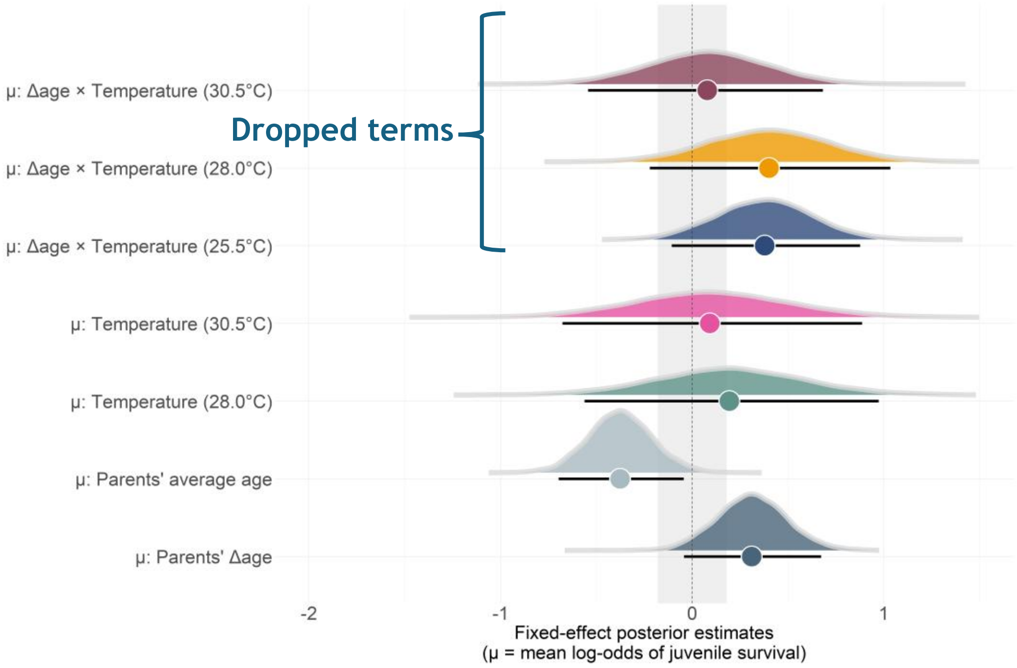

**Figure S9. Posterior estimates of the fixed effects from the Bernoulli model explaining variation in offspring juvenile survival.** For each fixed effect we show the posterior median (circles), and the 95% HDI'S. The dotted line shows the line of no effect (i.e., the population intercept, representing the average juvenile survival when Δage = 0 at the reference temperature (25.5°C)). The shaded area shows the region of practical equivalence (ROPE), representing the area of negligible effect (± 0.18 log-odds). Although the *parents' temperature x Δage* interaction was removed from our final model, we show the conditional slopes for the Δage effect under each of the three temperature treatments. N = 987 offspring from 77 pairs.

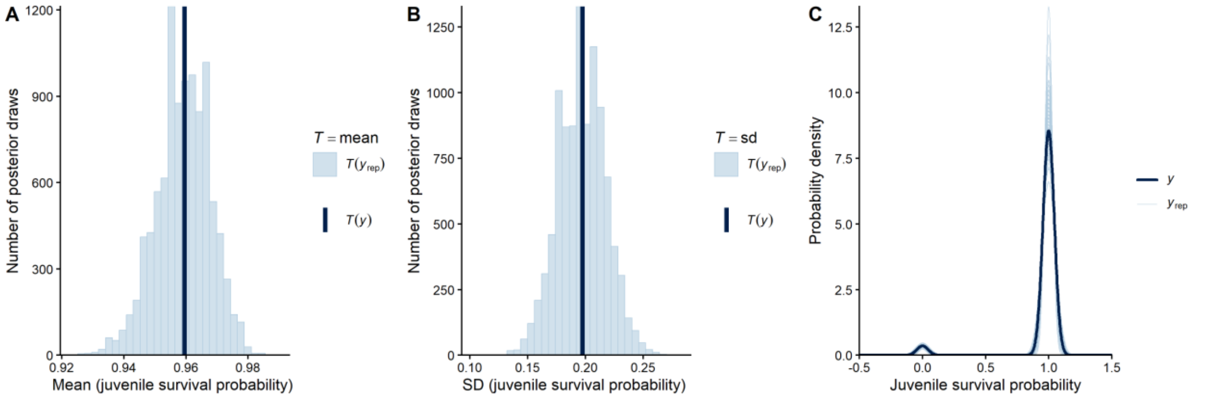

**Figure S10. Inspection of model fit for juvenile survival.** Unlike the model fit plots for the other offspring traits (which used more complex LOO QQ & predictive interval plots), for the Bernoulli models of offspring early-life and juvenile survival, we based our inspection of model fit solely on the posterior predictive distribution plots. **A)** and **B)** denote the posterior predictive and empirical means and standard deviations, respectively. On the y-axis is the number of posterior draws, while the x-axis is either the mean or standard deviation of offspring juvenile survival. The dark line represents the observed mean or SD, while the light blue lines describe the posterior predictive test statistic. **C)** The probability density of juvenile survival. The dark blue lines represent the observed distribution of juvenile failures (i.e., 0) or successes (i.e., 1), while the light blue lines describe estimates from 100 posterior draws.

**Table S8. Comparison of the predictive performance of the Bernoulli models explaining variation in offspring juvenile survival (*JuvSurv*).** We included the following as fixed effects: The parents' delta age at reproduction (*ΔAge*), the parents' average age at reproduction $\overline{(Age)}$, the parents' temperature treatment (*Temp*), and a quadratic effect for delta age (*ΔAge*$^2$). Additionally, we included parent pair ID as a random intercept (*1|Pair ID*) and *ΔAge* as a random slope (*1 + ΔAge |Pair ID*). We highlight the selected from each stage of the analysis in bold, which we then carried as a reference model for the next stage of the analysis. Shown for each model is the expected log pointwise predictive densities ("*elpd*"), the Δelpd relative to the best-fitting model, along with the standard errors *("SE")*. Models with a Δelpd <±4 are said to have identical predictive performance to the best-fitting model. **a)** Testing the evidence for random slopes and quadratic *ΔAge* effects. **b)** Evidence for interactions between fixed effects. In ***a)*** we selected the model with the random slope unless there was substantial evidence its inclusion harmed predictive performance. In ***b)*** If models had similar predictive performance, then we selected the simplest model to explain variation in the response (*i.e, the model with no interactions)*. We selected **model 5** as the final model to explain variation in juvenile survival, which only included single-effect predictors. In all models, n = 987 offspring from 77 parent pairs.

| Model selection | | |
|---|---|---|
| ***Model*** | ***elpd (± SE)*** | ***Δelpd ± SE*** |
| ***a) Support for random slopes and quadratic age effects*** | | |
| Model 1<br>*JuvSurv~* $\overline{Age}$ *+ ΔAge + Temp + (1\|PairID)* | -168.4 ± 20.0 | 0.0 ± 0.0 |
| Model 2<br>*JuvSurv~* $\overline{Age}$ *+ ΔAge + ΔAge*$^2$ *+ Temp + (1\|PairID)* | -169.5 ± 20.2 | -1.1 ± 0.2 |
| **Model 3**<br>***JuvSurv~*** $\overline{Age}$ ***+ ΔAge + Temp + (1 + ΔAge\|PairID)*** | **-171.3 ± 20.3** | **-2.9 ± 1.2** |
| Model 4<br>*JuvSurv~* $\overline{Age}$ *+ ΔAge + ΔAge*$^2$ *+ Temp + (1 + ΔAge\|PairID)* | -170.6 ± 20.3 | -2.2 ± 0.8 |
| ***b)*** **Evidence for interactions between fixed effects**<br>*Using Model 3: Bernoulli model with single-effect predictors as a reference* | | |
| **Model 5**<br>***JuvSurv~*** $\overline{Age}$ ***+ ΔAge + Temp + (1 + ΔAge\|PairID)*** | **-167.8 ± 20.2** | **0.0 ± 0.0** |
| Model 6<br>*JuvSurv~* $\overline{Age}$ *+ ΔAge + Temp + ΔAge* Temp + (1 + ΔAge\|PairID)* | -169.1 ± 19.8 | -1.3 ± 0.8 |

**Table S9. Coefficients explaining variation in offspring juvenile survival from the selected model *(Model 5 from Table S8)*.** The following were included as fixed effects: the parents' delta age at reproduction, the parents' average age at reproduction, and the parents' temperature treatment. Additionally, we included the *Δslope* between the parents' average and delta age, used as a test for selective disappearance. Parent pair ID was included as a random intercept, while a random slope was fitted for the parents' delta age. We also report the Pearson's correlation coefficient (*r*) to assess correlations between the random intercept and random slope. Shown is the maximum a posteriori probability estimate ("*MAP*"), the 95% highest density intervals ("*95% HDI*"), along with the effective sample size ("ESS" [bulk/tail]), the probability of direction ("*pd*"), and the percentage of posterior estimates within ROPE ("*% in ROPE*"). Any fixed effects with a pd > 98% are highlighted in bold, assuming these represented genuine biological effects. Finally, we show the estimates for the dropped interaction between the parents' temperature treatment and parent's delta age. n = 987 offspring from 77 parent pairs.

| Coefficients from the selected model (*Model 5*) | | | | | |
|---|---|---|---|---|---|
| ***Predictors*** | ***MAP*** | ***HDI (95%)*** | ***% in ROPE*** [1] | ***pd*** [2] | **ESS (Bulk/Tail)** |
| ***a) Location (μ) Model*** | | | | | |

| | | | | | |
|---|---|---|---|---|---|
| **Intercept**<br>*Log odds of juvenile survival* | 3.31 | [2.77, 3.89] | 0% | 100% | 7959/7271 |
| Parents' Δage<br>*(weeks; z-scaled)* | 0.31 | [-0.06, 0.64] | 23.06% | 95.74% | 10983/8124 |
| **Parents' average age**<br>*(weeks; z-scaled)* | **-0.38** | **[-0.70, -0.04]** | **11.86%** | **98.66%** | **9036/7883** |
| **Δslope**$_{\text{average age -}\Delta\text{age}}$ | **-0.71** | **[-1.18, -0.19]** | **-** | **100%** | **-** |
| Parents' temperature treatment | | | | | |
| *25.5°C* | - | - | - | - | - |
| *28.0°C* | 0.15 | [-0.53, 0.95] | 31.62% | 69.69% | 10983/8124 |
| *30.5°C* | 0.08 | [-0.68, 0.89] | 34.00% | 59.33% | 8814/7521 |
| **Random Effects** | | | | | |
| σ Intercept$_{\text{Parent pair ID}}$ | 0.22 | [0.00, 0.66] | - | 100% | 3352/4205 |
| σ Parents' Δage$_{\text{Parent pair ID}}$ | 0.25 | [0.00, 0.71] | - | 100% | 2947/4407 |
| *r* (Intercept ~ slope Δage) | -0.10 | [-0.85, 0.76] | - | 54.24% | 6374/6769 |
| ***Temperature Δage slopes and pairwise comparisons (from the dropped interaction Parents' Temperature x Δage interaction)*** | | | | | |
| *Δage*: *25.5°C* | 0.40 | [-0.13, 0.85] | 20.37% | 93.26% | 5544/6239 |
| *Δage*: *28.0°C* | 0.41 | [-0.25, 1.01] | 42.25% | 89.97% | - |
| *Δage*: *30.5°C* | 0.09 | [-0.55, 0.67] | 43.35% | 60.54% | - |
| *Δage*: *25.5°C vs. 28.0°C* | -0.01 | [-0.75, 0.76] | 35.88% | 52.22% | 6694/7272 |
| *Δage*: *28.0°C vs. 30.5°C* | 0.33 | [-0.56, 1.21] | 24.31% | 76.81% | - |
| *Δage*: *30.5°C vs. 25.5°C* | -0.20 | [-1.07, 0.43] | 28.52% | 78.03% | 6631/7326 |
| **Marginal $R^2$** | **0.01** | [0.00, 0.04] | - | - | - |
| **Conditional $R^2$** | **0.02** | [0.00, 0.07] | - | - | - |

[1]ROPE = [−0.18, 0.18] on log-odds scale; applied to location fixed effects only.

[2]pd = probability of direction.

#### Section S9: Supplementary results for offspring total lifespan and mortality

##### *S9.1. Supplementary results for offspring mean total lifespan*

In the subset analysis that only included sexed adults, we found uncertain evidence that the three-way interaction between the *parents' delta age x temperature treatment x offspring sex* improved model predictive performance (Table S10). There was some limited evidence favouring a two-way interaction between the *parents' delta age x offspring sex*, but any difference between the sexes was likely negligible (*pd* = 96.74% & 68.8% in ROPE; Table S12). We found that the slight positive effect of parental age on offspring longevity may have been weaker in sons than in daughters (Table S12). We found strong evidence that sons exhibited longer total lifespans than daughters, with this sex-specific effect being independent of parental age (*pd* = 99.93% & 6.36% *in ROPE*; Table S12).

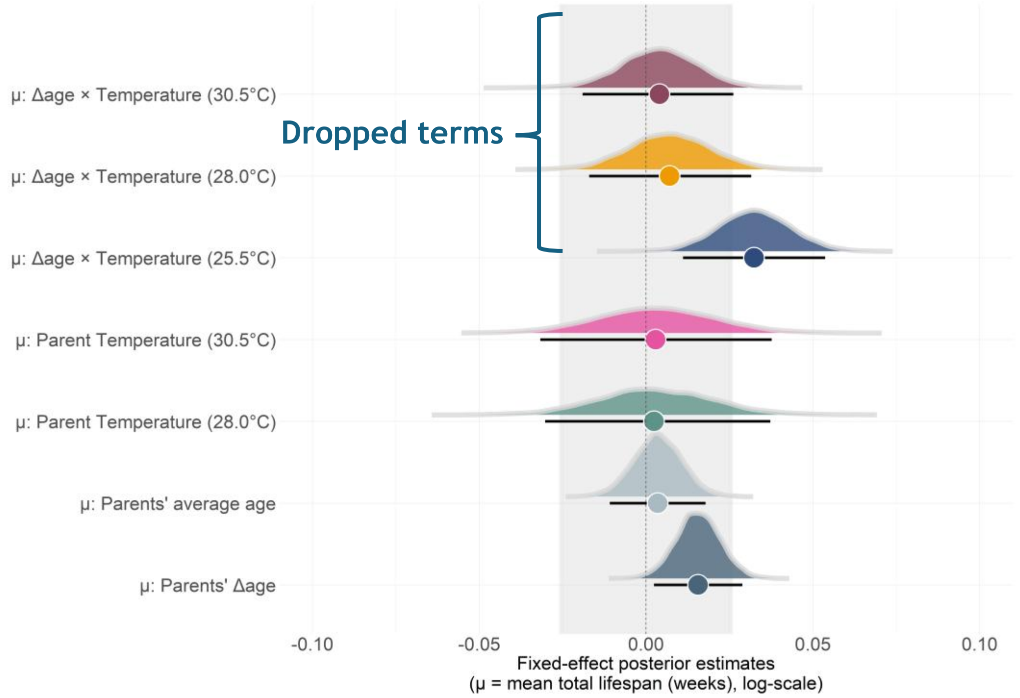

**Figure S11. Posterior distributions for the fixed effects from the Weibull model explaining variation in offspring longevity.** For each fixed effect we show the posterior median (circles) and the 95% Highest density intervals (error bars). The dotted line at zero represents the line of no effect, which describes the population intercept. Here, this represents the log of the mean total lifespan, when *Δage = 0* and at the reference temperature (25.5°C). The shaded area represents the ROPE, or region of negligible effect (±0.03 log weeks). Although the *parents' temperature x Δage* interaction was removed from our final model, we show the conditional slopes for Δage under the three temperature treatments. N = 987 offspring from 77 pairs.

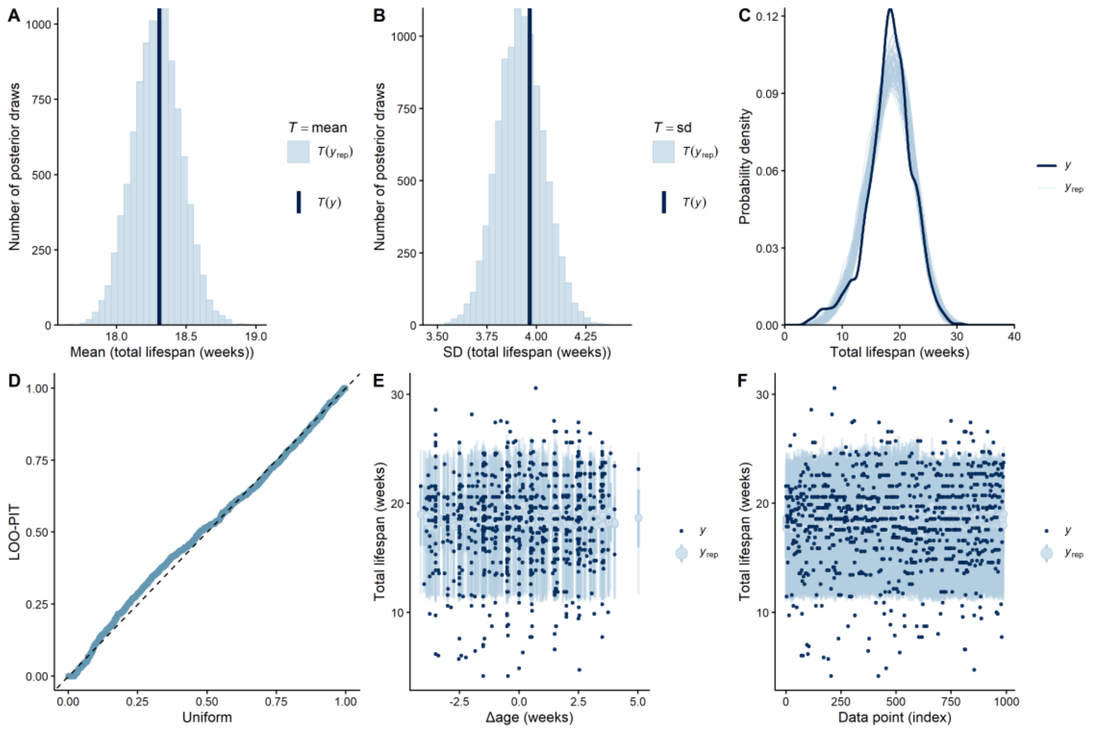

**Figure S12. Model fit plots for total lifespan, showing posterior predictive and out-of-sample predictive performance plots from the selected model (Model 7; Table S10). A) & B) Histogram of means and standard deviations for total lifespan from the posterior predictive and observed distribution.** On the y-axis is the number of posterior draws, while the x-axis describes the test-statistic (either mean or standard deviation). Shown are the estimated means and SD's from the posterior replicated data sets (*T*($Y_{rep}$); light blue), while the dark vertical line is the empirical mean or SD (*T(y)*). **C) Probability density of the empirical and posterior simulated data sets.** Light blue lines describe the distribution of the response (lifespan) as estimated from 100 simulated posterior draws. The dark blue line is the observed distribution of offspring total lifespan. **D) Quantile-Quantile (Q-Q plot) plotting the distribution of the Leave-One-Out Probability Integrals (LOO-PIT).** The dashed diagonal line represents the expected uniform distribution of the LOO-PIT integrals, while the blue points represent the observed distribution of the LOO-PIT integrals. **E) Posterior predictive interval plot.** On the y-axis is the offspring's total lifespan while the x-axis describes the effect of *Δage*. Small dots represent the observed data, while light blue points and error bars describe the posterior predictive value and credible intervals (50% and 95%) for each observation. **F) LOO interval plot.** This plot describes the ability of the model to predict each observation when blinded to the original data. Shown on the x-axis is each data point, while on the y-axis is the estimate of the response variable (total lifespan). The dark points represent the observed value of each observation, while the light green points and bars represent the LOO predicted data and credible intervals (95% CI's). Both plots **E)** and **F)** show that although the model is well-calibrated, it struggles to directly predict individual observations (i.e., the model contains considerable unexplained variation), although most observations fall within the posterior estimates 95% CI's. N = 987 offspring from 77 parent pairs.

**Table S10. Comparison of the predictive performance of *brms* models explaining variation in offspring longevity (*TotalLife*).** We included the following as fixed effects: The parents' delta age at reproduction (*ΔAge*), the parents' average age at reproduction $\overline{(Age)}$, the parents' temperature treatment (*Temp*), and the quadratic effect of delta age (*ΔAge*$^2$). Additionally, we included parent pair ID as a random intercept (*1|Pair ID*) and *ΔAge* as a random slope (*1 + ΔAge |Pair ID*). We highlight the selected from each stage of the analysis in bold, which we then carried as a reference model for the next stage of the analysis. For each model, we show the expected log pointwise predictive densities ("*elpd*"), the Δelpd relative to the best-fitting model, along with the standard errors *("SE")*. **a)** Comparison of model likelihoods. **b)** Testing the evidence for random slopes and quadratic *ΔAge* effects. **c)** Evidence for interactions between fixed effects. In **b)** we selected the model with the random slope unless there was substantial evidence its inclusion harmed predictive performance. In **c)** If models had similar predictive performance, then we selected the simplest model to explain variation in the response (*i.e, the model with no interactions)*. We selected **model 7**, which only included single-effect predictors, as the final model to explain variation in total lifespan. In all models, n = 987 offspring from 77 parent pairs.

| Model selection based on predictive performance | | |
|---|---|---|
| ***Model*** | ***elpd (± SE)*** | ***Δelpd (± SE)*** |
| ***a) Comparison of model likelihoods*** | | |
| **Model 1** ***Weibull*** | **-2748.0 ± 26.7** | **0.0 ± 0.0** |
| Model 2 *Exponential* | -3859.4 ± 6.8 | -1111.5 ± 29.5 |
| Model 3 *Lognormal* | -2912.5 ± 38.9 | -164.5 ± 29.5 |
| ***b)*** **Testing support for random slopes and quadratic age effects** | | |
| *Using **Model 1**: Weibull model as a reference* | | |
| Model 1 *TotalLife~* $\overline{Age}$ *+ ΔAge + Temp + (1\|PairID)* | -2748.0 ± 26.7 | -1.1 ± 2.4 |
| Model 4 *TotalLife~* $\overline{Age}$ *+ ΔAge + ΔAge*$^2$ *+ Temp + (1\|PairID)* | -2747.1 ± 27.0 | -0.8 ± 1.6 |
| **Model 5** ***TotalLife~*** $\overline{Age}$ ***+ ΔAge + Temp + (1+ ΔAge\|PairID)*** | **-2748.5 ± 27.0** | **-0.3 ± 1.7** |
| Model 6 *TotalLife~* $\overline{Age}$ *+ ΔAge + ΔAge*$^2$ *+ Temp + (1 + ΔAge\|PairID)* | -2746.8 ± 27.2 | 0.0 ± 0.0 |
| ***c)*** **Evidence for interactions between fixed effects** | | |
| *Using **Model 5**: Weibull model with single-effect predictors only as a reference* | | |
| **Model 7** ***TotalLife~ ~*** $\overline{Age}$ ***+ ΔAge + Temp + (1+ ΔAge\|PairID)*** | **-2747.5 ± 26.9** | **0.0 ± 0.0** |
| Model 8 *TotalLife~ ~* $\overline{Age}$ *+ ΔAge + Temp + ΔAge*Temp + (1+ ΔAge\|PairID)* | -2747.0 ± 26.9 | -0.4 ± 1.9 |

**Table S11. Coefficients explaining variation in offspring total longevity from the selected model *(Model 7 from Table S10)*.** We included the following as fixed effects: the parents' delta age at reproduction, the parents' average age at reproduction, and the parents' temperature treatment. Additionally, we also report the *Δslope* between the parents' average and delta age, used to test for selective disappearance. Parent pair ID was included as a random intercept, while we also included a random slope for the parents' delta age. We additionally report the Pearson's correlation coefficient (*r)* to assess the covariance between the random intercept and random slope. Shown is the maximum a posteriori probability estimate ("*MAP*"), 95% highest density intervals ("*95% HDI*"), the effective sample size ("*ESS [bulk/tail]*"), the probability of direction ("*pd*"), and the percentage of posterior estimates within ROPE ("*% in ROPE*"). Any fixed effects with a pd > 98% highlighted in bold, assuming these represented genuine biological effects. Finally, we show the estimates for the dropped interaction between the parents' temperature treatment and parent's delta age. n = 987 offspring from 77 parent pairs.

| Coefficients from the selected model (*Model 7*) | | | | | |
|---|---|---|---|---|---|
| ***Predictors*** | ***MAP*** | ***HDI (95%)*** | ***% in ROPE***[1] | ***pd***[2] | **ESS (Bulk/Tail)** |
| ***a) Location (μ) Model*** | | | | | |
| **Intercept** *Log (total lifespan [weeks])* | 2.91 | [2.88, 2.93] | 0% | 100% | 12090/7350 |
| **Parents' Δage** *(weeks; z-scaled)* | **0.01** | **[0.00, 0.03]** | **93.88%** | **98.84%** | **14437/8177d** |
| Parents' average age | 0.00 | [-0.01, 0.02] | 99.87% | 69.03% | 14437/8177 |

| | | | | | |
|---|---|---|---|---|---|
| *(weeks; z-scaled)* | | | | | |
| $\Delta slope_{average\ age - \Delta age}$ | -0.01 | [-0.03, 0.01] | - | 88.74% | - |
| Parents' temperature treatment | | | | | |
| *25.5°C* | - | - | | - | - |
| *28.0°C* | 0.00 | [-0.03, 0.04] | 86.72% | 55.46% | 11527/7628 |
| *30.5°C* | 0.00 | [-0.03, 0.04] | 85.41% | 56.27% | 10892/7830 |
| **Random Effects** | | | | | |
| σ $Intercept_{Parent\ pair\ ID}$ | 0.03 | [0.00, 0.05] | - | 100% | 2077/2596 |
| σ Parents' $\Delta age_{Parent\ pair\ ID}$ | 0.02 | [0.00, 0.04] | - | 100% | 2351/4750 |
| *r* (Intercept ~ slope Δage) | 0.29 | [-0.54, 0.89] | - | 71.66% | 5814/6606 |
| ***Temperature Δage slopes and pairwise comparisons (from the dropped parents' Δage x temperature interaction)*** | | | | | |
| ***Δage: 25.5°C*** | **0.03** | **[0.01, 0.05]** | **26.63%** | **99.82%** | **7102/7570** |
| *Δage: 28.0°C* | 0.01 | [-0.02, 0.03] | 93.00% | 71.48% | - |
| *Δage: 30.5°C* | 0.00 | [-0.19, 0.03] | 96.84% | 63.61% | - |
| *Δage: 25.5°C vs. 28.0°C* | -0.03 | [-0.06, 0.01] | 51.19% | 93.60% | 8363/7734 |
| *Δage: 28.0°C vs. 30.5°C* | 0.00 | [-0.03, 0.03] | 87.32% | 57.44% | - |
| *Δage: 30.5°C vs. 25.5°C* | -0.03 | [-0.06, 0.00] | 44.12% | 96.47% | 8572/7332 |
| **Shape parameter (*k*)** | 5.49 | [5.20, 5.79] | - | - | 7897/7830 |
| **Marginal $R^2$** | 0.01 | [0.00 – 0.02] | - | - | - |
| **Conditional $R^2$** | 0.04 | [0.01, 0.07] | - | - | - |

[1]ROPE = [−0.026, 0.026] on log scale; applied to location fixed effects only.

[2]pd = probability of direction.

**Table S12. Subset analysis for offspring total lifespan where only sexed adults were included (to allow sex to explain variation in longevity).** ***1. Comparison of the predictive performance of models including interactions.*** For each model we show the expected log pointwise predictive densities ("*elpd*"), the *Δelpd*, along with their standard errors ("*SE*"). We inferred models with an Δelpd <±4 had identical predictive performance to the best-fitting model. Each of the following models include the following: Parents' average age ($\overline{Age}$), the Parents' delta age (*ΔAge*), the parent's temperature treatment (*Temp*), and offspring sex (*Sex*) as fixed effects and parent pair ID (1|*PairID*) as a random intercept, while *ΔAge* was also fitted as a random slope (1+ *ΔAge*|*PairID*). We selected **model 1**, the model with single-effect predictors only, to explain variation in offspring longevity. ***2. Coefficients explaining variation in offspring longevity (from Model 1).*** Shown is the mean estimate ("*Estimate*"), the 95% credible intervals ("*95% CI*"), the effective sample size ("*ESS [bulk/tail]*"), the probability of direction ("*pd*"), and the percentage of posterior estimates within ROPE ("*% in ROPE*"). Any fixed effects with a pd > 98% are highlighted in bold, assuming these represented genuine biological effects. n = 947 offspring from 77 parent pairs.

**1. Evidence for interactions based on predictive performance**

| Model | elpd (± SE) | Δelpd (± SE) |
|---|---|---|
| **Model 1** | | |
| ***TotalLife~ $\overline{Age}$ + ΔAge + Temp + Sex + (1+ ΔAge\|PairID)*** | *-2513.3 ± 23.7* | *-1.4 ± 2.0* |
| **Model 2** | | |
| *TotalLife~ $\overline{Age}$ + ΔAge + Temp + ΔAge*Sex*Temp + (1+ ΔAge\|PairID)* | *-2515.4 ± 23.9* | *-1.4 ± 2.0* |
| **Model 3** | | |
| *TotalLife~ $\overline{Age}$ + ΔAge + Temp + ΔAge*Sex + (1+ ΔAge\|PairID)* | *-2511.8 ± 23.7* | *0.00 ± 0.0* |

#### 2. Model coefficients: From Model 1

| Predictors | MAP | HDI (95%) | % in ROPE[1] | pd[2] | ESS (Bulk/tail) |
|---|---|---|---|---|---|
| ***a) Location (μ) Model*** | | | | | |
| **Intercept** *Log (total lifespan [weeks])* | 2.91 | [2.88, 2.94] | 0% | 100% | 5680/7410 |
| **Parents' Δage** *(weeks; z-scaled)* | **0.02** | **[0.00, 0.03]** | **75.56%** | **98.56%** | **8153/7332** |
| Parents' average age *(weeks; z-scaled)* | 0.01 | [-0.01, 0.02] | 95.67% | 88.34% | 7322/8076 |
| $\Delta slope_{average\ age\ -\ delta\ age}$ | -0.01 | [-0.02, 0.00] | - | 75% | - |
| Parents' temperature treatment | | | | | |
| *25.5°C* | - | - | - | - | - |
| *28.0°C* | 0.00 | [-0.03, 0.04] | 75.64% | 59.50% | 4978/6454 |
| *30.5°C* | 0.01 | [-0.03, 0.04] | 75.25% | 61.15% | 4978/6454 |
| Offspring sex | | | | | |
| *Female* | - | - | - | - | - |
| ***Male*** | **0.04** | **[0.01, 0.06]** | **6.36%** | **99.93%** | **13993/7400** |
| **Random Effects (*u*)** | | | | | |
| $\sigma$ $Intercept_{Parent\ pair\ ID}$ | 0.04 | [0.02, 0.06] | - | 100% | 3750/4652 |
| $\sigma$ Parents' $\Delta age_{Parent\ pair\ ID}$ | 0.03 | [0.01, 0.05] | - | 100% | 1852/1248 |
| *r* (Intercept ~ slope Δage) | 0.34 | [-0.22, 0.82] | - | 85.87% | 2998/4473 |
| ***Offspring Sex pairwise comparisons for the Δage slope (from the dropped interaction Parents' Δage x offspring sex)*** | | | | | |
| **Δage: Female** | **0.02** | **[-0.01, 0.02]** | **55.51%** | **99.84%** | **7511/7889** |
| *Δage*: Male | 0.01 | [-0.04, 0.00] | 99.11% | 68.72% | - |
| *Δage: Female vs. Male* | -0.02 | [-0.04, 0.00] | 68.8% | 96.74% | 8894/7622 |
| **Shape parameter (*k*)** | 6.35 ± 0.18 | [6.02, 6.70] | - | - | 13993/7400 |
| **Marginal $R^2$** | 0.03 | [0.01 – 0.04] | - | - | - |

| | | | | | |
|---|---|---|---|---|---|
| **Conditional $R^2$** | 0.09 | [0.05, 0.14] | - | - | - |

[1]ROPE = [−0.02, 0.02] on log scale; applied to location fixed effects only.

[2]pd = probability of direction.

##### *S9.2: Supplementary results for offspring total mortality*

We found the model with best predictive performance was the null model (DIC = -2311.98; *Table S13*), suggesting both parental age (DIC = -2304.66; *Figure S13*) and parental temperature (DIC = -2309.59), alongside their interaction (DIC = -2247.708; *Figure S14*), did not substantially influence offspring mortality. Additionally, we found no convincing evidence for a parental age by offspring sex interaction (*Figure S15*). We provide the exact mortality parameter estimates in *Table S14.*

Focusing on the isolated effect of parental age, baseline mortality (*c*) did not differ between offspring from late- and middle-aged parents (KLDC comparisons: *middle-aged* vs. *late-aged* = 0.50; *Figure S13: bottom-left*), although offspring from early-aged parents had the highest baseline mortality, albeit with broad posterior uncertainty (KLDC comparisons: *early-aged* vs. *late-aged* = 0.87, *early-aged* vs. *middle-aged* = 0.89; *Figure S13: bottom-left*). The posteriors for *b0* also overlapped (KLDC comparisons: *early-aged* vs. *late-aged* = 0.68, *early-aged* vs. *middle-aged* = 0.64, *middle-aged* vs. *late-aged* = 0.53; *Figure S13: centre-left*). In contrast, *b1* was smallest among offspring from late-aged parents (KLDC comparisons: *early-aged* vs. *late-aged* = 0.93, *middle-aged* vs. *late-aged* = 0.99), while there was little difference in *b1* between offspring from early- and middle-aged parents (KLDC comparisons: *early-aged* vs. *middle-aged* = 0.69; *Figure S13: top-left*). Overall, we concluded that both offspring baseline (*c*) and age-specific mortality (*b0* and *b1*) were independent of parental age, as, visually, there were only marginal differences in the mortality and survival trajectories (*Figure S13: right*).

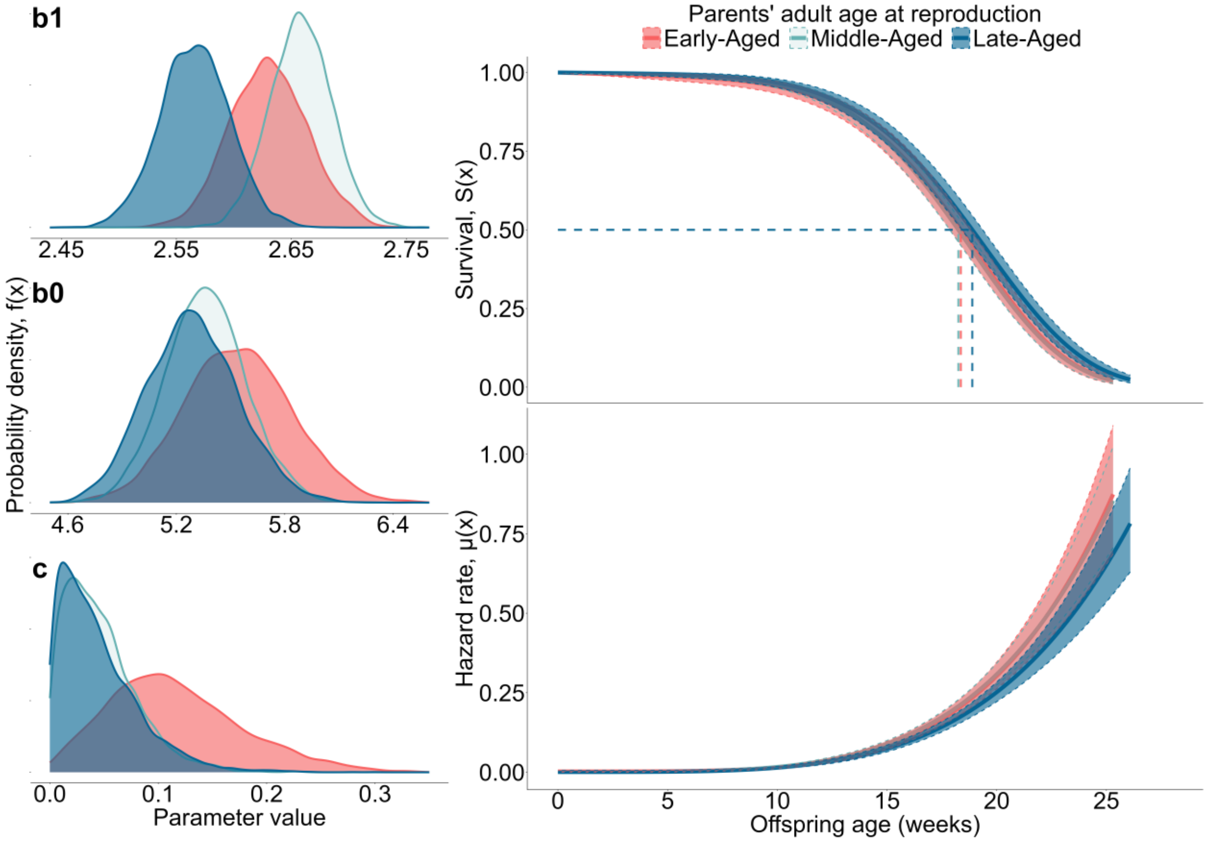

**Figure S13. The effect of parental age on the offspring's survival and mortality.** Shown on the right is the cumulative survival probability (top), with the median lifespan of offspring from each parental age group (dashed lines), and the instantaneous hazard rate (bottom) against the offspring's age (weeks). Lines and ribbons represent model predictions for survival and mortality ± 95% CI, estimated from the posterior draws. For visualisation, we transformed the hazard from the units 1/year (*BaSTA's* internal parameterisation) to the units 1/week. The panels on the left show the full posterior distributions of the Weibull mortality parameters *b1*, *b0* and *c*. Here, *b1* is in 1/year, *c* is in years, while *b0* is unitless. The colours describe the different parental age categories: red = "early-aged", green = "middle-aged", and blue = "late-aged". N = 987 offspring from 77 parent pairs.

We found no statistical support for the interaction between parental age and parental temperature (DIC = -2247.708; *Figure S14*), with this model having a higher DIC than both the null model (DIC = -2311.98) and the models including the isolated effects of parental age (DIC = -2304.66) and parental temperature (DIC = -2309.59). Therefore, we concluded that the effect of parental age on offspring longevity was not conditional on the parents' temperature treatment. This was despite visual inspection of the mortality and survival trajectories, which indicated slight positive effects of increasing parental age on offspring mortality when parents were maintained at 25.5°C, but not 28.0°C or 30.5°C (*Figure S14: bottom*). Any positive effect of parental age at 25.5°C may have arose through joint effects on *b1*, *b0*, and *c,* although the *b1* parameter exhibited the steepest separation of posteriors for early- and late-aged parents (*Figure S14: top-left*).

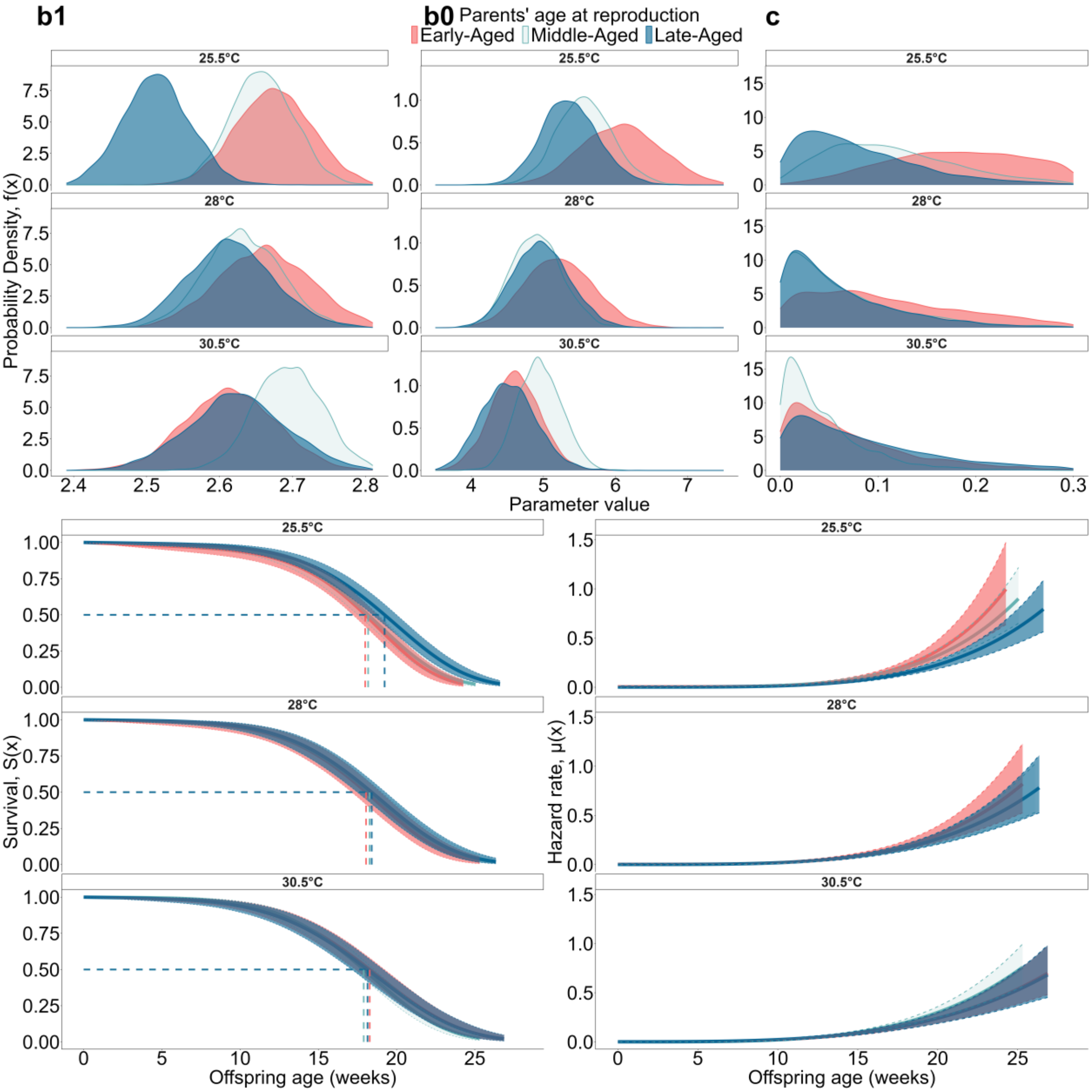

**Figure S14. The effect of the two-way interaction between the parents' age at reproduction and the parents' temperature treatment on offspring mortality and survival.** Shown in the top row are the posterior distributions for the *b1*, *b0*, and *c* parameters. Along the bottom row, the lines and ribbons are the model posterior predictions ± 95% CI for offspring mortality and survival. Model estimates for the Weibull parameters and survival/mortality curves are faceted by parental temperature. We transformed the hazard to take the units 1/week. The colours describe the different parental age categories: red = "early-aged", green = "middle-aged", and blue = "late-aged". N = 987 offspring from 77 parent pairs.

The model including the two-way interaction between offspring sex and parental age could not be directly compared using DIC with the other mortality models, as it used a different subset of available data (947 sexed offspring from 77 parent pairs). In Table S14, we only provide within-sex KLDC parental age comparisons to determine whether the direction, and magnitude, of the parental age effect was sex-dependent. The sexes exhibited noticeable differences in mean total longevity (*Table S12*), and we aimed to isolate our inferences to sex-dependent parental age effects on mortality, not baseline differences between each sex's mortality trajectory. Visual inspection of the survival and mortality curves revealed substantial overlap between the posterior estimates and credible intervals for offspring from early- and late-aged parents, irrespective of offspring sex (*Figure S14: bottom*).

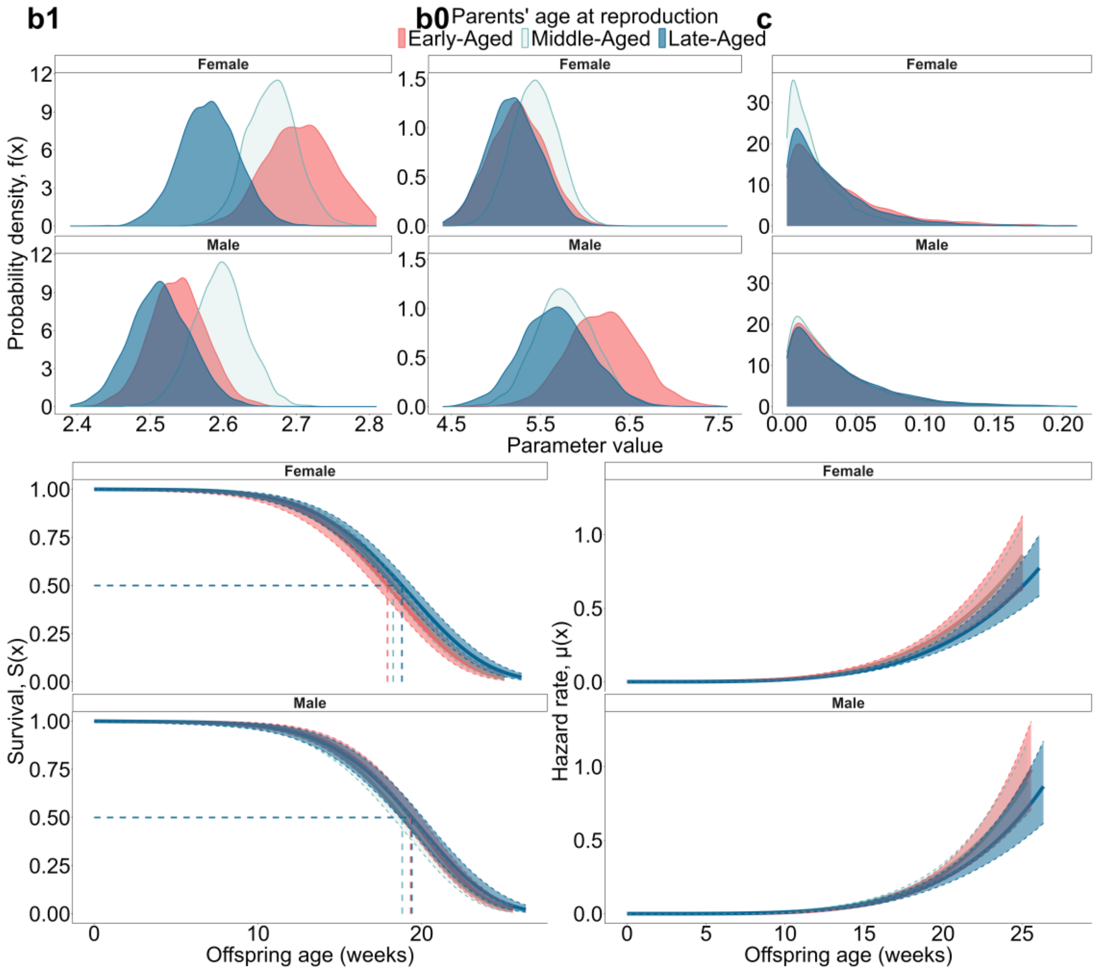

**Figure S15. The effect of the two-way interaction between offspring sex and the parents' adult age at reproduction on both the Weibull-Makeham mortality parameters (top) and the offspring's survival and mortality curves (bottom).** The full posterior distributions for *b1*, *b0*, and *c* are shown along the top-row (from left-right, respectively). In the bottom row, the lines and ribbons represent model predictions ± 95% CI for survival (left) and mortality (right). Model estimates for the Weibull parameters, alongside the survival/mortality curves, are faceted by the offspring's sex. The colours represent the different parental age categories: red = "early-aged", green = "middle-aged", and blue = "late-aged". N = 947 sexed offspring from 77 parent pairs.

**Table S13. Comparison of the predictive performance of models explaining variation in offspring mortality. A) Ranked list of all mortality distributions available in *BaSTA*.** For each model, we give their deviance information criterion (DIC), as well as their difference in DIC when compared to the best-performing model (the model with the lowest DIC value). We concluded that the best-fitting model was the most suitable when its ΔDIC < 3 from the next-best candidate model. If models had comparable predictive performance, then we selected the simplest model (the model with the fewest number of parameters). When testing the given mortality distributions for our study population, we included no covariates in the model. **B) Comparison of the predictive performance of different prior structures** (default, weakly informative, and moderately informative) fitted to models with each set of covariates: null model, parent's adult age at reproduction and the parents' temperature treatment. Moderately informative priors have the same mean, but ½ the SD of the weakly informative priors. For models with the *parental age x parental temperature interaction*, along with the *parental age x offspring sex* interaction, we based our posterior inferences on the weakly informative prior structure. N = 987 offspring from 77 parent pairs.

| Model selection based on predictive performance | | | |
|---|---|---|---|
| ***Model*** | ***Shape*** | ***DIC*** | ***ΔDIC*** |
| ***a)*** **Comparison of *BaSTA* mortality distributions** | | | |
| **Weibull** | **Makeham** | **-2311.93** | **0** |
| Weibull | bathtub | -2310.60 | 1.33 |
| Weibull | simple | -2307.14 | 4.79 |
| Gompertz | simple | -2213.62 | 98.31 |
| Gompertz | Makeham | -2208.02 | 103.91 |
| Gompertz | bathtub | -2203.64 | 108.29 |
| Logistic | simple | -2194.59 | 117.34 |
| Logistic | Makeham | -2193.82 | 118.11 |
| Logistic | bathtub | -2185.23 | 126.70 |
| Exponential | simple | -91.53 | 2220.40 |
| ***b)*** **Comparison of different prior structures on posterior predictive performance** *Using the Weibull-Makeham mortality distribution* | | | |
| **Model** | **Prior** | **DIC** | ***ΔDIC*** |
| Null | Weakly-informative | -2311.97 | 0 |
| Null | Default | -2311.93 | 0.04 |
| Null | Moderately-informative | -2300.36 | 11.61 |
| Parental temperature | Weakly-informative | -2309.60 | 2.37 |
| Parental temperature | Default | -2309.42 | 2.55 |
| Parental temperature | Moderately-informative | -2236.96 | 75.01 |
| Parental age | Weakly-informative | -2304.66 | 7.31 |
| Parental age | Default | -2304.41 | 7.56 |
| Parental age | Moderately-informative | -2241.18 | 70.79 |

**Table S14. Posterior parameter estimates for covariates explaining variation in the offspring mortality parameters.** Shown is the null model (with no covariates) (**1**), the independent effects of parental age (**2**) and parental temperature (**3**). For each parameter, we show the posterior mean ("*Estimate*"), Standard errors ("*SE*"), the 95% credible intervals ("*95% CI*"), along with the $\hat{R}$ and serial autocorrelation ("*S.A.corr*") estimates. We also include the KLDC estimates to determine the distance in the posterior distributions between categorical covariates. KLDC estimates > 0.8 are highlighted in bold, assuming these represent genuine biological effects. We also provide the DIC values to estimate which model had the best predictive performance to explain variation in offspring mortality. For the combined effect of *parental age x parental temperature* (**4**), we only report the DIC value, as this model had substantially poorer predictive performance than the null model. N = 987 offspring from 77 parent pairs. We also show the parameter estimates for the model including the combined effect of *parental age* x *offspring sex* (**5**). Note that we do not report the DIC from this analysis. This model uses a different subset of data and cannot be directly compared to the other models based on information criterion (N = 947 sexed offspring from 77 parent pairs).

| Posterior mortality parameter estimates | | | | |
|---|---|---|---|---|
| *Mortality Parameters* | ***Estimate ± SE*** | ***CI (95%)*** | $\hat{R}$ | **S.A.corr** |
| ***1) Null Model (no covariates): DIC = -2311.98*** | | | | |
| **Makeham parameter (*c*)** | 0.05 ± 0.02 | 0.01 - 0.11 | 1.00 | 0.00 |
| **Scale parameter (*b1*)** | 2.62 ± 0.02 | 2.58 - 2.65 | 1.00 | -0.01 |
| **Shape parameter (b0)** | 5.50 ± 0.16 | 5.19 - 5.82 | 1.00 | 0.00 |
| ***2) Parents' adult age at reproduction: DIC = -2304.66*** | | | | |
| **Makeham parameter (*c*)** | | | | |
| *Parental age: Early-Aged* | 0.12 ± 0.06 | 0.02 - 0.26 | 1.00 | 0.01 |
| *Parental age: Middle-Aged* | 0.05 ± 0.03 | 0.00 - 0.13 | 1.00 | 0.00 |
| *Parental age: Late-Aged* | 0.04 ± 0.04 | 0.00 - 0.14 | 1.00 | 0.00 |
| **Scale parameter (*b1*)** | | | | |
| *Parental age: Early-Aged* | 2.63 ± 0.03 | 2.57 - 2.70 | 1.00 | -0.02 |
| *Parental age: Middle-Aged* | 2.66 ± 0.03 | 2.61 - 2.71 | 1.00 | -0.01 |
| *Parental age: Late-Aged* | 2.57 ± 0.03 | 2.51 - 2.63 | 1.00 | -0.01 |
| **Shape parameter (*b0*)** | | | | |
| *Parental age: Early-Aged* | 5.54 ± 0.30 | 4.97 - 6.13 | 1.00 | 0.02 |
| *Parental age: Middle-Aged* | 5.36 ± 0.22 | 4.93 - 5.81 | 1.00 | 0.01 |
| *Parental age: Late-Aged* | 5.29 ± 0.25 | 4.81 - 5.81 | 1.00 | 0.02 |

| *KLDC pairwise comparisons* | | | |
|---|---|---|---|
| ***Parental age comparison*** | ***c*** | ***b1*** | ***b0*** |
| *Early-aged vs. Late-aged* | **0.87** | **0.93** | 0.68 |
| *Middle-aged vs. Early-aged* | **0.89** | 0.69 | 0.64 |
| *Late-aged vs. Middle-aged* | 0.50 | **0.99** | 0.53 |

**3) Parents' temperature treatment: *DIC = -2309.59***

| *Mortality Parameters* | ***Estimate ± SE*** | ***CI (95%)*** | $\hat{R}$ | **S.A.corr** |
|---|---|---|---|---|
| **Makeham parameter (*c*)** | | | | |
| *Parental temperature: 25.5°*C | 0.13 ± 0.05 | 0.05 - 0.24 | 1.00 | -0.01 |
| *Parental temperature: 28.0°*C | 0.04 ± 0.04 | 0.00 - 0.15 | 1.00 | 0.01 |
| *Parental temperature: 30.5°*C | 0.03 ± 0.03 | 0.00 - 0.10 | 1.00 | 0.02 |
| **Scale parameter (*b1*)** | | | | |
| *Parental temperature: 25.5°*C | 2.59 ± 0.03 | 2.54 - 2.64 | 1.00 | 0.01 |

| | | | | |
|---|---|---|---|---|
| *Parental temperature: 28.0°C* | 2.63 ± 0.03 | 2.56 – 2.67 | 1.00 | -0.01 |
| *Parental temperature: 30.5°C* | 2.65 ± 0.03 | 2.59 – 2.71 | 1.00 | -0.01 |
| **Shape parameter (*b0*)** | | | | |
| *Parental temperature: 25.5°C* | 6.04 ± 0.28 | 5.49 – 6.60 | 1.00 | 0.00 |
| *Parental temperature: 28.0°C* | 5.37 ± 0.26 | 4.88 – 5.89 | 1.00 | 0.00 |
| *Parental temperature: 30.5°C* | 4.93 ± 0.20 | 4.54 – 5.35 | 1.00 | 0.00 |

***KLDC pairwise comparisons***

| ***Parental temperature comparison*** | ***c*** | ***b1*** | ***b0*** |
|---|---|---|---|
| *25.5°C vs. 28.0°C* | **0.92** | **0.80** | **0.98** |
| *25.5°C vs. 30.5°C* | **0.98** | **0.94** | **0.99** |
| *28.0°C vs. 30.5°C* | 0.65 | 0.61 | **0.92** |

**4) *Parents' temperature treatment x Parents' adult age at reproduction: DIC = -2247.708***

**5) *Parents' age at reproduction x Offspring sex***

| ***Mortality Parameters*** | ***Estimate ± SE*** | ***CI (95%)*** | $\hat{R}$ | **S.A.corr** |
|---|---|---|---|---|
| **Makeham parameter (*c*)** | | | | |
| ***Parental age: early-aged,*** *Offspring sex: male* | 0.04 ± 0.04 | 0.00 - 0.14 | 1.00 | 0.01 |
| *Offspring sex: female* | 0.04 ± 0.04 | 0.00 - 0.14 | 1.00 | -0.00 |
| ***Parental age: middle-aged,*** *Offspring sex: male* | 0.03 ± 0.03 | 0.00 – 0.12 | 1.00 | -0.00 |
| *Offspring sex: female* | 0.02 ± 0.02 | 0.00 – 0.08 | 1.00 | 0.01 |
| ***Parental age: late-aged,*** *Offspring sex: male* | 0.04 ± 0.04 | 0.00 – 0.14 | 1.00 | -0.00 |
| *Offspring sex: female* | 0.03 ± 0.03 | 0.00 – 0.11 | 1.00 | 0.01 |
| **Scale parameter (*b1*)** | | | | |
| ***Parental age: early-aged,*** *Offspring sex: male* | 2.54 ± 0.04 | 2.46 – 2.62 | 1.00 | -0.01 |
| *Offspring sex: female* | 2.70 ± 0.05 | 2.62 – 2.81 | 1.00 | 0.01 |
| ***Parental age: middle-aged,*** *Offspring sex: male* | 2.60 ± 0.04 | 2.53 – 2.67 | 1.00 | -0.01 |
| *Offspring sex: female* | 2.67 ± 0.03 | 2.60 – 2.74 | 1.00 | -0.00 |
| ***Parental age: late-aged,*** *Offspring sex: male* | 2.51 ± 0.04 | 2.43 – 2.60 | 1.00 | -0.02 |
| *Offspring sex: female* | 2.58 ± 0.04 | 2.50 – 2.66 | 1.00 | -0.02 |
| **Shape parameter (*b0*)** | | | | |

| | | | | |
|---|---|---|---|---|
| ***Parental age: early-aged,*** *Offspring sex: male* | 6.21 ± 0.40 | 5.46 - 7.01 | 1.00 | 0.00 |
| *Offspring sex: female* | 5.24 ± 0.32 | 4.62 - 5.87 | 1.00 | 0.00 |
| ***Parental age: middle-aged,*** *Offspring sex: male* | 5.78 ± 0.33 | 5.14 - 6.42 | 1.00 | 0.02 |
| *Offspring sex: female* | 5.44 ± 0.26 | 4.93 - 5.95 | 1.00 | 0.00 |
| ***Parental age: late-aged,*** *Offspring sex: male* | 5.67 ± 0.39 | 4.91 - 6.46 | 1.00 | 0.00 |
| *Offspring sex: female* | 5.20 ± 0.30 | 4.61 - 5.79 | 1.00 | 0.01 |

***KLDC pairwise comparisons → Only providing within-sex comparisons***

| *Offspring sex comparison* | *c* | *b1* | *b0* |
|---|---|---|---|
| ***Offspring sex: Female*** *Parental age: early vs. late* | 0.52 | **0.99** | 0.51 |
| *Parental age: early vs. middle* | 0.71 | 0.74 | 0.62 |
| *Parental age: middle vs. late* | 0.61 | **0.97** | 0.66 |
| ***Offspring sex: male*** *Parental age: early vs. late* | 0.50 | 0.59 | **0.80** |
| *Parental age: early vs. middle* | 0.51 | **0.85** | 0.76 |
| *Parental age: middle vs. late* | 0.52 | **0.95** | 0.53 |

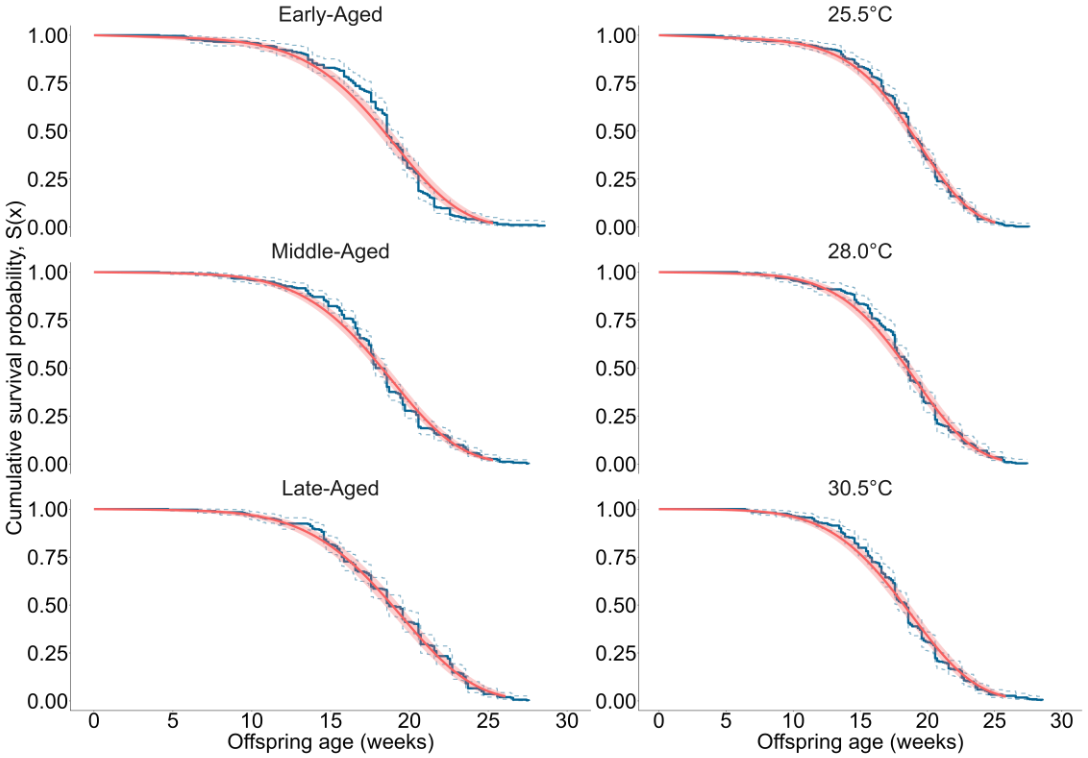

**Figure S16. Model fit for Weibull-Makeham models explaining variation in the offspring mortality (after removing early-life mortality).** Shown on the left are the survival curves for offspring from each parental age category, while the, on the right, the offspring's survival curves for each parental temperature treatment are shown. Kaplan-Meier plots of the offspring's observed survival probabilities are coloured in blue, while the parametric Weibull-Makeham survival curves are coloured in red. N = 987 offspring from 77 parent pairs.

#### Section S10: Supplementary results for offspring development time

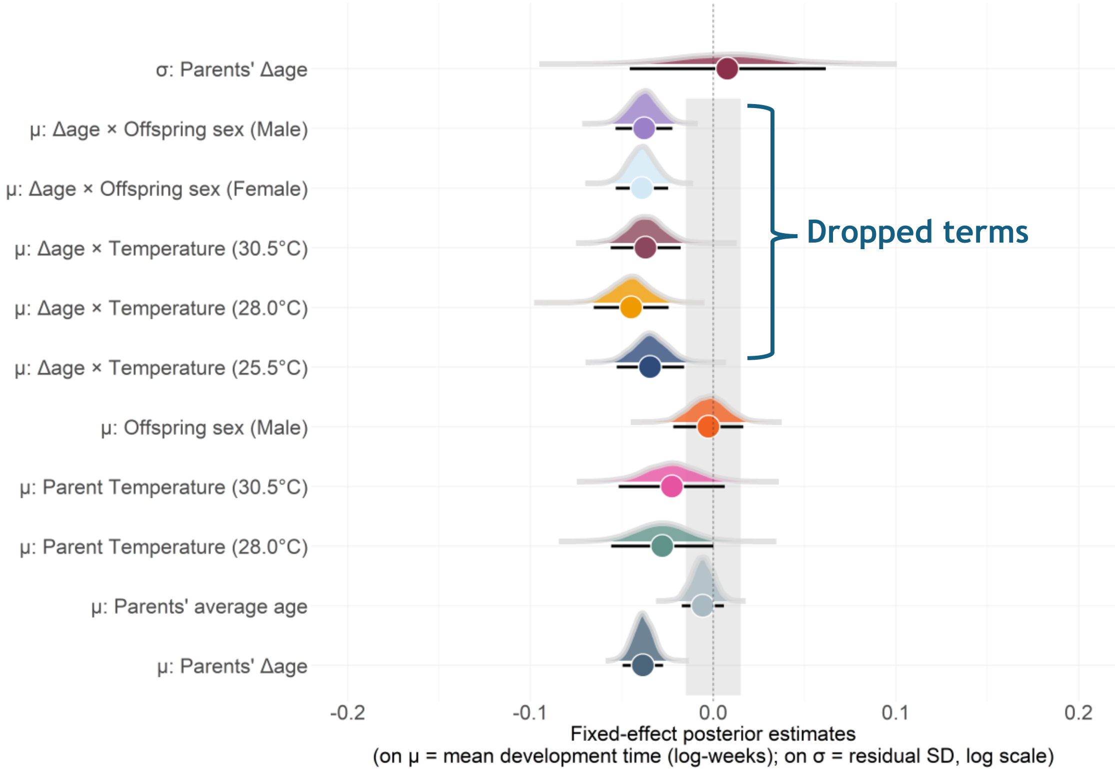

**Figure S17. Posterior distributions of the fixed effects from the lognormal model explaining variation in offspring development time.** Shown are the fixed effects from both the location ($\mu$) and scale ($\sigma$) component. For each fixed effect, shown is the posterior median (circles) and the 95% HDI's (error bars). The probability density curves describe the entire posterior distribution for each fixed effect. The dotted line represents the line of no effect, while the shaded area depicts the region of practical equivalence (i.e., the ROPE), describing the area of negligible effect (± 0.01 log weeks). For $\mu$ parameters, the intercept represents the log (mean development time) for daughters of parents maintained at 25.5°C. For σ, the intercept is the log (residual standard deviation for development time). Although not included in our final model, we show the conditional *Δage* slopes under each temperature treatment, and for each offspring sex. N = 939 offspring from 77 parent pairs.

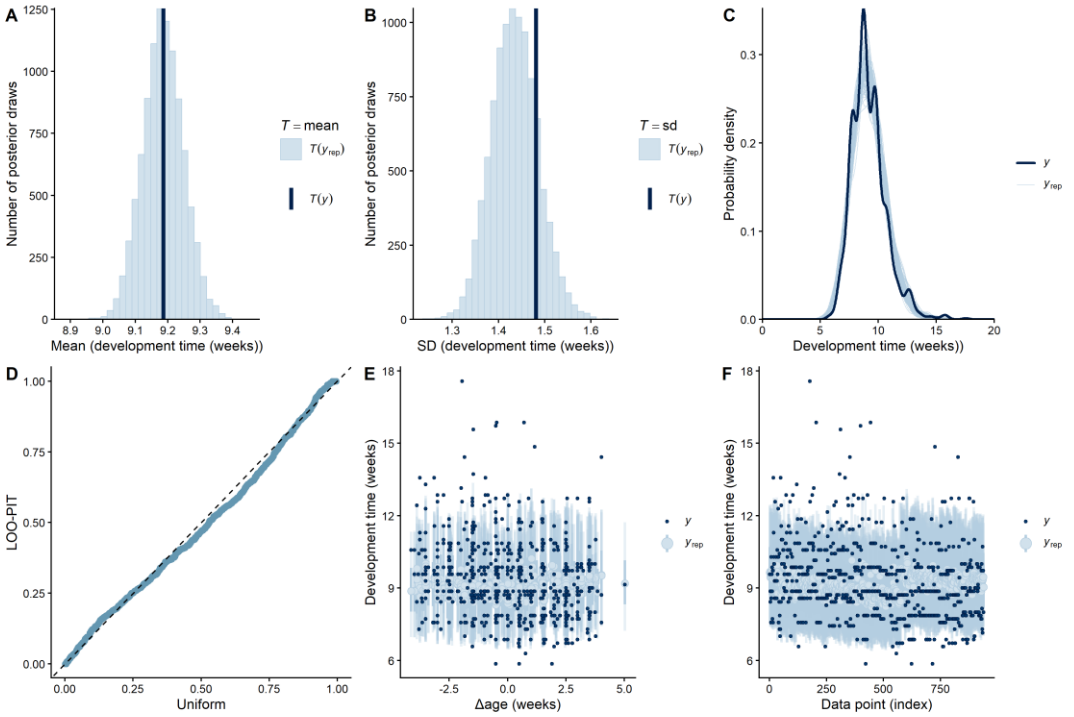

**Figure S18. Model fit plots for development time, showing posterior predictive and out-of-sample predictive performance plots from the selected model (Model 10, *Table S15*). A) & B) Histogram of means and standard deviations (SD's) from the posterior predicted and observed distribution.** We show the estimated means/SD's from the posterior simulated data (*$T(Y_{rep})$*; light blue), and the raw mean/SD of the observed data (*T(y);* dark blue). **C) Probability density of the empirical and simulated data sets.** Light blue lines display the distribution of the response (development time), estimated from 100 simulated posterior draws. The dark blue line describes the observed distribution of development time. Note that although the observed distribution has several peaks, this is likely an artifact of our sampling, since we only inspected animals weekly for adult emergence. **D) Quantile-Quantile (Q-Q plot) plotting the distribution of the Leave-One-Out Probability Integrals (LOO-PIT).** The dashed diagonal line represents the expected uniform distribution of the LOO-PIT integrals, while the blue points represent the observed distribution of each of the LOO-PIT integrals. **E) Posterior predictive interval plot.** On the y-axis is the offspring's development time while the x-axis describes the effect of *Δage*. Small dots represent the empirical data, while light blue points and error bars describe the posterior predictive value and credible intervals (50% and 95%) for each observation. **F) LOO interval plot.** This plot is used to show the ability of the model to predict each observation. On the x-axis is each data point, while on the y-axis is development time. The dark points represent the observed value of each observation, while the light blue points and bars represent the LOO predicted data and credible intervals (both 50% and 95% CI's). Both plots **E)** and **F)** show that although the model is well-calibrated, it struggles to directly predict individual observations (i.e., the model contains considerable unexplained variation), although most observations fall within the posterior estimates 95% CI's. N = 939 offspring from 77 parent pairs.

**Table S15. Comparison of the predictive performance of models explaining variation in offspring development time (*Devtime*).** Highlighted in bold is the model selected from each stage of the analysis, which was then carried forward as a reference model to the next stage. We include the following as fixed effects: The parents' delta age at reproduction (*ΔAge*), the parents' average age at reproduction $(\overline{Age})$, the parents' temperature treatment (*Temp*), the quadratic effect of delta age (*$\Delta Age^2$*), and offspring sex (*Sex*). We also included parent pair ID as a random intercept (*1|Pair ID*) and *ΔAge* as a random slope (*1 + ΔAge |Pair ID*). For each model, shown is the expected log pointwise predictive density ("*elpd*"), the Δelpd relative to the best-fitting model, along with the standard errors *("SE")*. Models with a Δelpd <±4 were said to have identical predictive performance to the best-fitting model. **a)** Comparison of model likelihoods. **b)** Testing the evidence for random slopes and quadratic *ΔAge* effects. **c)** Support for fixed effects in the scale component. **d)** Evidence for interactions between fixed effects. In **b)** we selected the model with the random slope unless there was substantial evidence against its inclusion (i.e., if it harmed predictive performance). In **d)**

If models had similar predictive performance, then we selected the simplest model to explain variation in the response (*i.e, the model with no interactions)*. We selected ***model 10*** as the final model to explain variation in offspring development time, which only included single-effect predictors. In all models, n = 939 offspring from 77 parent pairs.

**Model selection based on predictive performance**

| ***Model*** | ***elpd (± SE)*** | ***Δelpd (± SE)*** |
|---|---|---|
| ***a) Comparison of model likelihoods*** | | |
| Model 1<br>*Weibull* | -1768.2 ± 32.90 | -148.1 ± 13.8 |
| Model 2<br>*Exponential* | -3024.4 ± 4.9 | -1404.2 ± 25.1 |
| **Model 3**<br>***Lognormal*** | **-1620.1 ± 27.4** | **0.0 ± 0.0** |
| ***b)* Testing support for random slopes and quadratic age effects**<br>*Using **Model 3**: Lognormal model as a reference* | | |
| Model 3<br>*Devtime ~* $\overline{Age}$ *+ ΔAge + Temp + Sex+ (1\|PairID)* | -1620.1 ± 27.4 | -0.4 ± 1.4 |
| Model 4<br>*Devtime ~* $\overline{Age}$ *+ ΔAge +* $\Delta Age^2$ *+ Temp + Sex+ (1\|PairID)* | -1620.0 ± 27.5 | -0.3 ± 1.9 |
| **Model 5**<br>***Devtime ~*** $\overline{Age}$ ***+ ΔAge + Temp + Sex+ (1+ ΔAge\|PairID)*** | **-1619.7 ± 27.5** | **0.0 ± 0.0** |
| Model 6<br>*Devtime ~* $\overline{Age}$ *+ ΔAge +* $\Delta Age^2$ *+ Temp + Sex+ (1 + ΔAge\|PairID)* | -1619.7 ± 27.6 | 0.0 ± 1.4 |
| **c) Evidence for fixed effects in the scale component**<br>*Using **Model 5**: Lognormal model with single-effect predictors* | | |
| Model 7<br>σ (*Devtime*) ~1 (No fixed effects on σ) | -1619.7 ± 27.5 | 0.0 ± 0.0 |
| Model 8<br>σ (*Devtime*) ~ *ΔAge* + *Temp* | -1620.9 ± 27.6 | -0.7 ± 1.4 |
| **Model 9**<br>**σ (*Devtime*) ~ *ΔAge*** | **-1623.5 ± 27.8** | **-3.3 ± 1.6** |
| ***d)* Evidence for interactions between fixed effects**<br>*Using **Model 9** as a reference* | | |
| **Model 10**<br>**Location: *Devtime ~*** $\overline{Age}$ ***+ ΔAge + Temp +Sex+ (1+ ΔAge\|PairID)***<br>**Scale: σ (*Devtime*) ~ *ΔAge*** | **-1621.0 ± 27.6** | **0.0 ± 0.0** |
| Model 11<br>Location: *Devtime ~* $\overline{Age}$ *+ ΔAge + Temp +Sex+ ΔAge*Temp*Sex + (1+ ΔAge\|PairID)*<br>Scale: σ (*Devtime*) ~ *ΔAge* | -1623.4 ± 27.5 | -2.4 ± 2.5 |
| Model 12<br>Location: *Devtime~* $\overline{Age}$ *+ ΔAge + Temp + Sex + ΔAge*Temp + (1+ ΔAge\|PairID)*<br>Scale: σ (*Devtime*) ~ *ΔAge* | -1621.8 ± 27.6 | -0.8 ± 0.7 |
| Model 13<br>Location: *Devtime~* $\overline{Age}$ *+ ΔAge + Temp + Sex + ΔAge*Sex + (1+ ΔAge\|PairID)*<br>Scale: σ (*Devtime*) ~ *ΔAge* | -1621.5 ± 27.6 | -0.5 ± 0.2 |
| Model 14<br>Location: *Devtime ~* $\overline{Age}$ *+ ΔAge + Temp + Sex + ΔAge*Temp + (1+ ΔAge\|PairID)*<br>Scale: σ (Devtime) ~ *ΔAge + Temp + ΔAge*Temp* | -1625.9 ± 27.9 | -4.9 ± 1.3 |

<table>
<tr><td>Model 15<br>Location: Devtime ~ $\overline{Age}$ + ΔAge + Temp + Sex + ΔAge*Sex + (1+ ΔAge|PairID)<br>Scale: σ (Devtime) ~ ΔAge + Sex + ΔAge*Sex</td><td>-1621.6 ± 27.3</td><td>-0.6 ± 2.4</td></tr>
</table>

**Table S16. Parameters explaining variation in offspring development time from the selected model *(Model 10)*.** In the location component (*μ*), the model included the following fixed effects: the parents' delta age, the parents' average age, the parents' temperature treatment, and offspring sex. Additionally, we also include the *Δslope* between the parents' average and delta age, as a test of selective disappearance. Parent pair ID was included as a random intercept, while we also included a random slope for the parents' delta age. We also report the Pearson's correlation coefficient (*r)* to assess the covariance between the random intercept and random slope. In the scale component (σ), only parental age was included as a fixed effect. Shown for each parameter is the maximum a posteriori probability estimate ("*MAP*"), the 95% highest density intervals ("*95% HDI*"), along with the effective sample size ("*ESS [bulk/tail]*"), the probability of direction ("*pd*"), and the percentage of posterior estimates in the ROPE ("*% in ROPE*"). Any fixed effects with a pd > 98% are highlighted in bold, assuming these represent genuine biological effects. Finally, we also show the estimates for the dropped interaction between the parents' delta age and temperature treatment (and between the parents' delta age and offspring sex). n = 939 offspring from 77 parent pairs.

| **Coefficients from the selected model (*Model 10*)** | | | | | |
|---|---|---|---|---|---|
| ***Predictors*** | ***MAP*** | ***HDI (95%)*** | ***% in ROPE[1]*** | ***Pd[2]*** | **ESS (Bulk/Tail)** |
| ***a) Location (μ) Model*** | | | | | |
| **Intercept** *Log (development time [weeks])* | 2.23 | [2.21, 2.25] | 0% | 100% | 8921/7257 |
| **Parents' Δage** | **-0.04** | **[-0.05, -0.03]** | **0.01%** | **100%** | **12347/7642** |
| Parents' average age | -0.01 | [-0.02, 0.01] | 96.21% | 83.63% | 10367/7491 |
| **$\Delta slope_{average\ age\ - \Delta age}$** | **0.03** | **[0.02, 0.05]** | **-** | **100%** | **-** |
| Parents' temperature treatment | | | | | |
| *25.5°C* | - | - | - | - | - |
| *28.0°C* | -0.03 | [-0.06, 0.00] | 17.80% | 97.46% | 8082/7786 |
| *30.5°C* | -0.02 | [-0.05, 0.01] | 29.07% | 93.89% | 8320/7419 |
| Offspring sex | | | | | |
| *Female* | - | - | - | - | - |
| *Male* | -0.00 | [-0.02, 0.02] | 85.12% | 60.42% | 16300/7514 |
| **Random Effects** | | | | | |
| σ $Intercept_{Parent\ pair\ ID}$ | 0.03 | [0.01, 0.04] | - | 100% | 1998/1820 |
| σ Parents' $\Delta age_{Parent\ pair\ ID}$ | 0.02 | [0.00, 0.04] | - | 100% | 1709/2639 |
| *r* (Intercept ~ slope Δage) | -0.04 | [-0.77, 0.70] | - | 55.08% | 4505/5654 |
| ***Temperature Δage slopes and pairwise comparisons (from the dropped Parents' Δage x Temperature interaction)*** | | | | | |

| | | | | | |
|---|---|---|---|---|---|
| ***Δage*: 25.5°C** | **-0.03** | **[-0.05, -0.02]** | **2.01%** | **100%** | **4345/5857** |
| ***Δage*: 28.0°C** | **-0.04** | **[-0.07, -0.03]** | **0.37%** | **100%** | **-** |
| ***Δage*: 30.5°C** | **-0.04** | **[-0.06, -0.02]** | **1.32%** | **100%** | **-** |
| *Δage*: 25.5°C vs. 28.0°C | -0.01 | [-0.04, 0.02] | 59.98% | 77.67% | 5254/6245 |
| *Δage*: 28.0°C vs. 30.5°C | -0.01 | [-0.04, 0.02] | 63.65% | 70.54% | - |
| *Δage*: 30.5°C vs. 25.5°C | 0.00 | [-0.03, 0.02] | 72.78% | 57.75% | 4753/6234 |
| ***Offspring sex pairwise comparisons for the Δage slope (from the dropped Parents' Δage x Offspring sex interaction)*** | | | | | |
| ***Δage*: Female** | **-0.04** | **[-0.05, 0.02]** | **0.02%** | **100%** | **9473/7451** |
| ***Δage*: Male** | **-0.04** | **[-0.05, -0.02]** | **0.23%** | **100%** | **-** |
| *Δage: Female vs. Male* | 0.00 | [-0.02, 0.02] | 85.56% | 54.63% | 5254/6245 |
| ***b) Scale (σ) Model*** | | | | | |
| **Intercept** *Log (residual SD)* | -1.92 | [-1.97, -1.87] | - | 100% | 6259/6461 |
| Parents' *Δage* | 0.01 | [-0.04, 0.06] | - | 60.74% | 12919/7520 |
| **Marginal $R^2$** | 0.07 | [0.03, 0.10] | - | - | - |
| **Conditional $R^2$** | 0.10 | [0.06, 0.15] | - | - | - |

[1]ROPE = [−0.015, 0.015] on log scale; applied to location fixed effects only.

[2]pd = probability of direction.

#### Section S11: Supplementary results for offspring adult mass

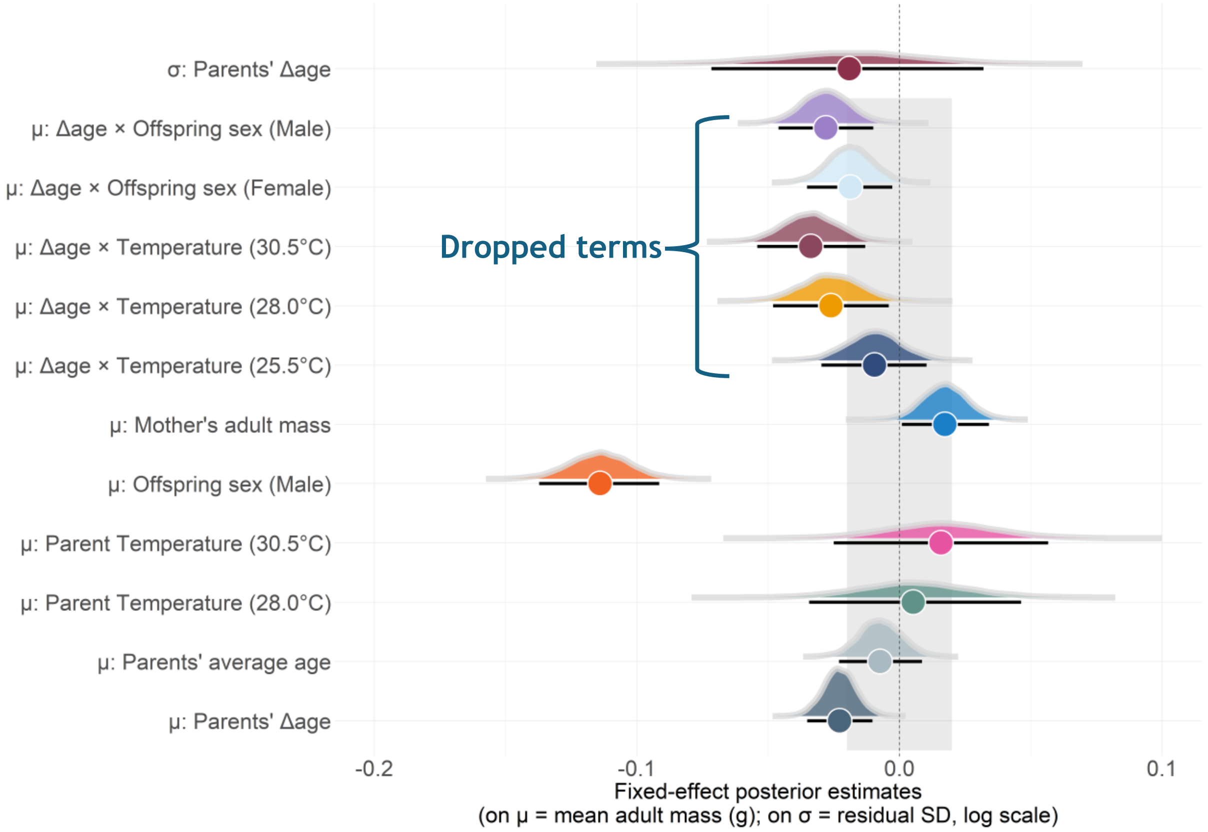

**Figure S19. Posterior distributions of the fixed effects from the Gaussian model explaining variation in offspring adult mass.** Shown are the fixed effects from both the location ($\mu$) and scale ($\sigma$) component. For each fixed effect, the circles display the posterior median, while the error bars describe the 95% HDI's. The dotted line is the line of no effect, representing the models statistical intercept. The shaded area represents the region of practical equivalence (i.e., the ROPE), describing the area of negligible effect (± 0.02g). For μ parameters, the intercept is the mean adult mass for daughters of parents kept at 25.5°C and when *Δage* = *0*. Additionally, we show the conditional slopes for the *Δage* effects in each of the temperature treatment and for each offspring sex. All interaction terms were removed from our final models, given they did not receive statistical support. For σ, the intercept is the log (residual standard deviation of offspring adult mass). N = 937 offspring from 77 parent pairs.

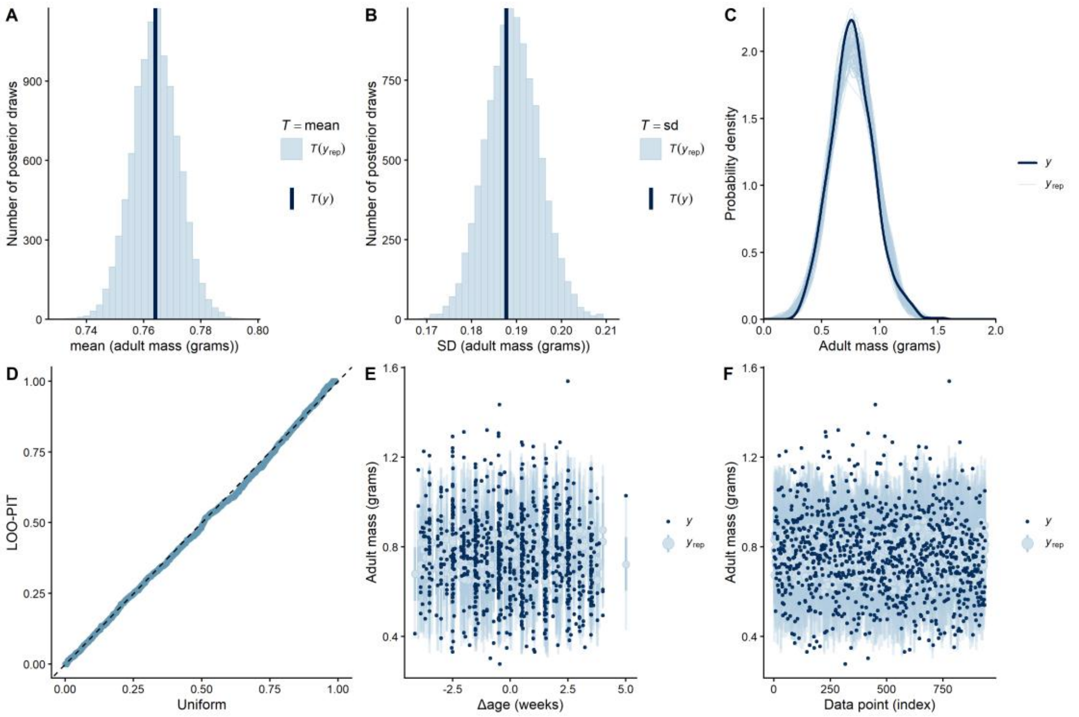

**Figure S20. Model fit plots for adult mass (from Model 8, *Table S17*). A) & B) Histogram of means and standard deviations (SD's) from the posterior predictive and observed distribution.** Here we show the simulated means/SD's from the posterior distribution (*$T(Y_{rep})$*; light blue), and the empirical means/SD's from the observed distribution (*T(y);* dark blue). **C) Probability density of the empirical and simulated data sets.** The thin light blue lines represent the simulated distribution of adult mass, as estimated from 100 posterior draws. The dark blue line describes the observed distribution of offspring adult mass. **D) Quantile-Quantile (Q-Q plot) plotting of the LOO-PIT values.** The dashed diagonal line denotes the expected uniform distribution of LOO-PIT integrals, while the blue points are observed distribution of the LOO-PIT integrals. **E) Posterior predictive interval plot.** On the y-axis is the offspring's adult mass while the x-axis describes the *Δage* effect. Small dots represent the observed data, while light blue points and error bars describe the posterior predicted value and credible intervals (50% and 95%) for each observation. **F) LOO interval plot.** We used this plot to describe the models ability to predict each observation after it was blinded to that data point. On the x-axis is each observed value, while on the y-axis is the estimate of the response variable (adult mass). The dark points represent the observed value of each observation, while the light blue points and bars represent the LOO predicted data and credible intervals (50% and 95% CI's). N = 937 offspring from 77 parent pairs.

**Table S17. Model selection for offspring adult mass, based on the comparison of out-of-sample predictive performance.** Highlighted in bold is the model selected from each stage of the analysis, which was then carried forward as a reference model to the next stage. We included the following as fixed effects: The parents' delta age at reproduction (*ΔAge*), the parents' average age at reproduction $\overline{(Age)}$, the parents' temperature treatment (*Temp*), the quadratic effect of delta age (*$\Delta Age^2$*), offspring sex (*Sex*), and the mother's adult mass (*Mothermass*). Here, we also included parent pair ID as a random intercept (*1|Pair ID*) and *ΔAge* as a random slope (*1 + ΔAge |Pair ID*). For each model, we show the expected log pointwise predictive density ("*elpd*"), the Δelpd relative to the best-fitting model, along with the standard errors *("SE")*. Models with a Δelpd <±4 were said to have identical predictive performance to the best-fitting model. **a)** Testing the evidence for random slopes and quadratic *ΔAge* effects. **b)** Support for fixed effects in the scale component. **c)** Evidence for interactions between fixed effects. In **a)** we selected the model with the random slope unless there was substantial evidence against its inclusion (i.e., if it harmed predictive performance). In **c)** If models had similar predictive performance, then we selected the simplest model to explain variation in the response (*i.e, the model with no interactions)*. We selected **model 8** as the final model to explain variation in offspring adult mass, which only included single-effect predictors. In all models, n = 937 offspring from 77 parent pairs.

| **Model selection based on predictive performance** |
| --- |

| *Model* | *elpd (± SE)* | *Δelpd (± SE)* |
|---|---|---|
| ***a)* Testing support for random slopes and quadratic age effects**<br>*Using **a Gaussian** likelihood* | | |
| Model 1<br>*Mass ~ $\overline{Age}$ + ΔAge + Temp + Sex + Mothermass + (1\|PairID)* | 300.2 ± 24.1 | 0.0 ± 0.0 |
| Model 2<br>*Mass ~ $\overline{Age}$ + ΔAge + $ΔAge^2$ + Temp + Sex + Mothermass + (1\|PairID)* | 299.2 ± 24.1 | -1.0 ± 0.3 |
| **Model 3**<br>***Mass ~ $\overline{Age}$ + ΔAge + Temp + Sex + Mothermass + (1+ ΔAge\|PairID)*** | **300.0 ± 24.1** | **-0.2 ± 1.3** |
| Model 4<br>*Mass ~ $\overline{Age}$ + ΔAge + $ΔAge^2$ + Temp + Sex+ Mothermass + (1 + ΔAge\|PairID)* | 299.0 ± 24.1 | -1.2 ±1.3 |
| ***b)* Evidence for fixed effects in the scale component**<br>*Using **Model 3**: Gaussian model with single-effect predictors* | | |
| Model 5<br>σ (*Mass*) ~1 (No fixed effects on σ) | 300.0 ± 24.1 | -0.6 ± 3.0 |
| Model 6<br>σ (*Mass*) ~ *ΔAge* + *Temp* | 300.5 ± 23.7 | 0.0 ± 0.0 |
| **Model 7**<br>**σ (*Mass*) ~ *ΔAge*** | **299.3 ± 24.2** | **-1.2 ± 3.0** |
| ***c)* Evidence for interactions between fixed effects**<br>*Using **Model 7** as a reference* | | |
| **Model 8**<br>**Location: *Mass ~ $\overline{Age}$ + ΔAge + Temp +Sex+ Mothermass + (1+ ΔAge\|PairID)***<br>**Scale: σ (*Mass*) ~ *ΔAge*** | **298.7 ± 24.2** | **-0.1 ± 2.7** |
| Model 9<br>Location: *Mass ~ $\overline{Age}$ + ΔAge + Temp +Sex+ Mothermass + ΔAge*Temp*Sex + (1+ ΔAge\|PairID)*<br>Scale: σ (*Mass*) ~ *ΔAge* | -294.7 ± 24.0 | -4.1 ± 3.4 |
| Model 10<br>Location: *Mass ~ $\overline{Age}$ + ΔAge + Temp + Sex + Mothermass+ ΔAge*Temp + (1+ ΔAge\|PairID)*<br>Scale: σ (*Mass*) ~ *ΔAge* | 298.6 ± 24.1 | -0.2 ± 3.1 |
| Model 11<br>Location: *Mass ~ $\overline{Age}$ + ΔAge + Temp + Sex + Mothermass + ΔAge*Sex + (1+ ΔAge\|PairID)*<br>Scale: σ (*Mass*) ~ *ΔAge* | 298.2 ± 24.2 | -0.6 ± 2.6 |
| Model 12<br>Location: *Mass ~ $\overline{Age}$ + ΔAge + Temp + Sex + Mothermass + ΔAge*Temp + (1+ ΔAge\|PairID)*<br>Scale: σ (*Mass*) ~ *ΔAge + Temp + ΔAge*Temp* | 297.9 ± 23.7 | -0.9 ± 4.1 |
| Model 13<br>Location: *Mass ~ $\overline{Age}$ + ΔAge + Temp + Sex + Mothermass + ΔAge*Sex + (1+ ΔAge\|PairID)*<br>Scale: σ (*Mass*) ~ *ΔAge + Sex + ΔAge*Sex* | 298.8 ± 24.1 | -4.1 ± 3.4 |

**Table S18. Parameters explaining variation in offspring adult body mass from the selected model *(Model 8)*.** In the location component ($\mu$), we included the following as fixed effects: the parents' delta age, the parents' average age, the parents' temperature treatment, offspring sex, and the mother's adult body mass. Additionally, we also include the *Δslope* between the parents' average and delta age, as a test of selective disappearance. Parent pair ID was set as a random intercept, while we also included a random slope for the parents' delta age. We also report *r* to assess correlations between the random intercept and random slope. In the scale component (σ), parental age was included as a fixed effect. Shown for each parameter is the posterior maximum a posteriori probability estimate ("*MAP*"), the 95% highest density intervals ("*95% HDI*"), along with the effective sample size ("*ESS [bulk/tail]*"), the probability of direction ("*pd*"), and the percentage of posterior estimates in ROPE *("% in ROPE")*. We highlight in bold any fixed effects with a pd > 98%. Finally, we also show the estimates for the dropped interaction between the parents' delta age and temperature treatment, and for the *parents' delta age x offspring sex* interaction. n = 937 offspring from 77 parent pairs.

| Coefficients from the selected model (*Model 8*) | | | | | |
|---|---|---|---|---|---|
| ***Predictors*** | ***MAP*** | ***HDI (95%)*** | ***% in ROPE[1]*** | ***pd[2]*** | **ESS (Bulk/Tail)** |
| ***a) Location (μ) Model*** | | | | | |
| **Intercept** *Offspring adult mass (g)* | 0.81 | [0.78, 0.84] | 0% | 100% | 5268/6143 |
| **Parents' Δage** | **-0.02** | **[-0.04, -0.01]** | **25.96%** | **99.99%** | **13971/7707** |
| Parents' average age | -0.01 | [-0.02, 0.01] | 92.36% | 81.28% | 6827/6594 |
| $\Delta slope_{average\ age\ -\Delta age}$ | 0.02 | [0.00, 0.04] | - | 93.96% | - |
| Mother's adult mass (g) | 0.02 | [0.00, 0.03] | 56.86% | 98.05% | 5612/6999 |
| Parents' temperature treatment | | | | | |
| *25.5°C* | - | - | - | - | - |
| *28.0°C* | 0.00 | [-0.04, 0.05] | 63.15% | 60.79% | 4510/6536 |
| *30.5°C* | 0.02 | [-0.03, 0.06] | 51.20% | 77.75% | 4781/6057 |
| Offspring sex | | | | | |
| *Female* | - | - | - | - | - |
| ***Male*** | **-0.11** | **[-0.14, -0.09]** | **0%** | **100%** | **13722/8248** |
| **Random Effects** | | | | | |
| σ $Intercept_{Parent\ pair\ ID}$ | 0.05 | [0.03, 0.07] | - | 100% | 3667/5100 |
| σ Parents' $\Delta age_{Parent\ pair\ ID}$ | 0.01 | [0.00, 0.03] | - | 100% | 2739/3447 |
| *r* (Intercept ~ slope Δage) | -0.55 | [-0.94, 0.46] | - | 77.91% | 6303/6082 |
| ***Temperature Δage slopes and pairwise comparisons (from the dropped Parent's Δage x Temperature interaction)*** | | | | | |
| *Δage: 25.5°C* | -0.01 | [-0.03, 0.01] | 81.16% | 83.04% | 5938/7269 |
| Δage: 28.0°C | -0.03 | [-0.05, -0.003] | 26.34% | 99.01% | - |
| Δage: 30.5°C | -0.03 | [-0.05, -0.01] | 7.8% | 99.91% | - |
| *Δage: 25.5°C vs. 28.0°C* | -0.02 | [-0.05, 0.01] | 54.42% | 85.90% | 8061/7624 |
| *Δage: 28.0°C vs. 30.5°C* | 0.01 | [-0.02, 0.039] | 72.01% | 69.41% | - |

| | | | | | |
|---|---|---|---|---|---|
| *Δage: 30.5°C vs. 25.5°C* | -0.02 | [-0.05, 0.00] | 34.87% | 94.64% | 8170/7814 |
| ***Offspring sex pairwise comparisons for the Δage slope (from the dropped Parent's* Δage x Offspring sex interaction)** | | | | | |
| ***Δage*: Female** | **-0.02** | **[-0.04, -0.01]** | **50.17%** | **98.87%** | **9312/7844** |
| ***Δage*: Male** | **-0.03** | **[-0.05, -0.01]** | **15.29%** | **99.84%** | **-** |
| *Δage: Female vs. Male* | -0.01 | [-0.03, 0.01] | 77.32% | 78.12% | 9813/8766 |
| ***b) Scale (σ) Model*** | | | | | |
| **Intercept** *Log (residual SD)* | -1.76 | [-1.81, -1.71] | - | 100% | 10079/8490 |
| Parents' Δage | -0.02 | [-0.07, 0.03] | - | 76.63% | 19117/ 8038 |
| **Marginal $R^2$** | 0.11 | [0.08, 0.15] | - | - | - |
| **Conditional $R^2$** | 0.18 | [0.13, 0.23] | - | - | - |

[1]ROPE = [−0.019, 0.019 grams]; applied to location fixed effects only.

[2]pd = probability of direction.

#### Section S12: Supplementary results for offspring relative growth rate (RGR)

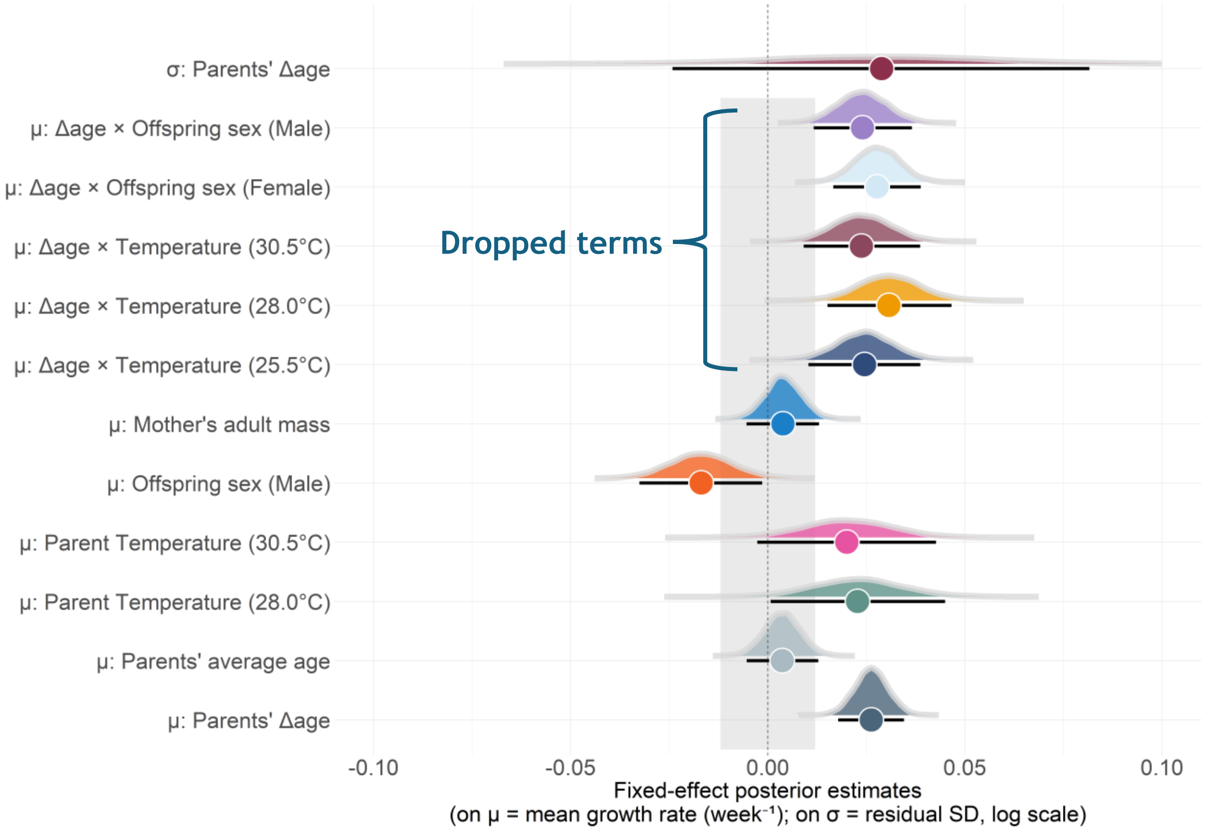

**Figure S21. Posterior distributions of the fixed effects from the Gaussian model, explaining variation in the offspring's relative growth rates (RGR).** Here, we show the fixed effects on both the average trait value (i.e., the location component) and on the residual error term (the scale component; *σ)*. The circles describe the posterior median for each fixed effect, while the error bars display the 95% HDI's. The dotted line at zero is the line of no effect, which represents the population-level intercept. The shaded area is the region of practical equivalence (i.e., ROPE), which denotes the region of negligible effect (± 0.01). For the μ parameters, the intercept is the mean RGR of daughters from parents maintained at 25.5˚C and when *Δage* = *0*. In this component, we also show the conditional temperature slopes for the effect of *Δage* in each temperature treatment, and for each offspring sex. Meanwhile, for σ the intercept is the log of the residual standard deviation of RGR. N = 937 offspring from 77 parent pairs.

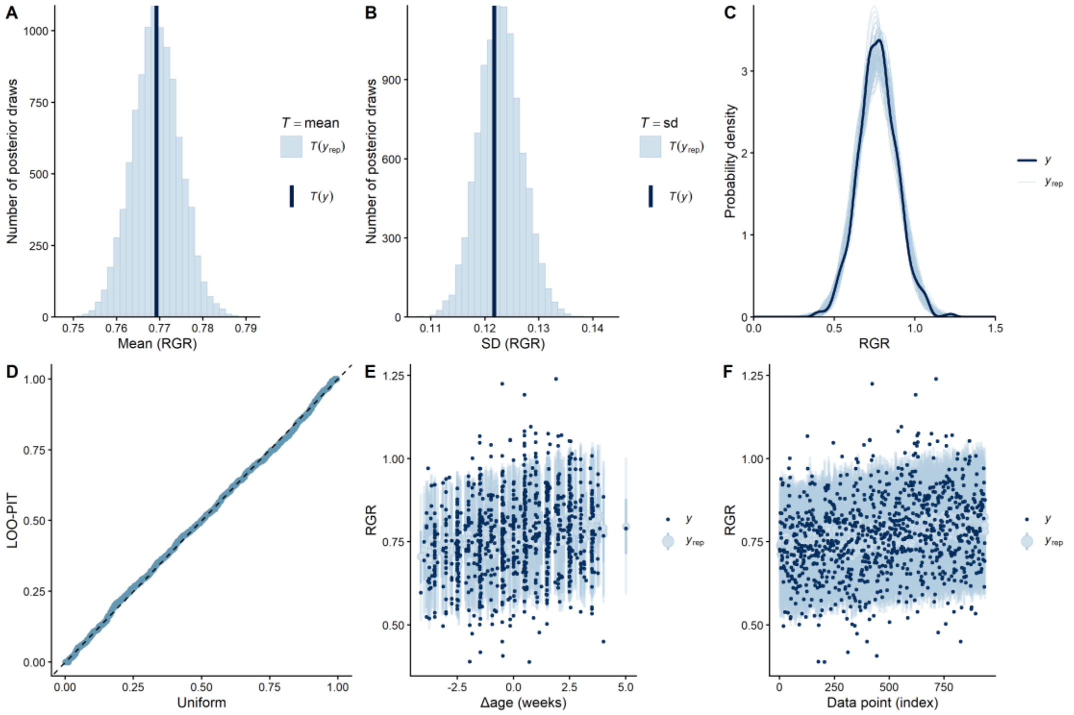

**Figure S22. Model fit plots for RGR (from Model 8, *Table S19*). A) and B) Histogram displaying the means and standard deviations as estimated from the posterior predictive and empirical distribution.** The posterior simulated means/SD's from the posterior distribution are shown in light blue (*T($Y_{rep}$)*), while the empirical mean/SD from the observed distribution is shown in dark blue *T(y)*). **C) Probability density of the empirical and posterior simulated data.** The thin light blue lines represent the simulated distribution of growth rate, as estimated from 100 posterior draws. The dark line represents the distribution of the observed data. **D) Quantile-Quantile (Q-Q plot) of the LOO-PIT values.** The dashed diagonal line denotes the expected uniform distribution of LOO-PIT integrals, while each individual point represents the observed value of the LOO-PIT integral. **E) Posterior predictive interval plot.** On the y-axis is the offspring's relative growth rate, while the x-axis describes the *Δage* effect. Small dots represent the observed data, while light blue points and error bars describe the posterior predicted value and credible intervals (50% & 95% CI's) for each observation. **F) LOO interval plot.** We used this plot to describe the model's ability to predict each observation after it was blinded to that data point. On the x-axis is each observed value, while on the y-axis is the estimate of the response variable (RGR). The dark points represent the observed value of each observation, while the light blue points and bars represent the LOO predicted data and credible intervals (50% and 95% CI's). N = 937 offspring from 77 parent pairs.

**Table S19. Comparison of models explaining variation in the offspring's RGR, based on their out-of-sample predictive performance.** Highlighted in bold is the model selected from each stage of the analysis, which was then carried forward as a reference model to the next stage. We included the following as fixed effects: The parents' delta age at reproduction (*ΔAge*), the parents' average age at reproduction $\overline{(Age)}$, the parents' temperature treatment (*Temp*), the quadratic effect of delta age (*ΔAge*$^2$), offspring sex (*Sex*), and the mother's adult mass (*Mothermass*). Here, we also included parent pair ID as a random intercept (*1|Pair ID*) and *ΔAge* as a random slope (*1 + ΔAge |Pair ID*). For each model, we show the expected log pointwise predictive density ("*elpd*"), the Δelpd relative to the best-fitting model, along with the standard errors *("SE")*. Models with a Δelpd <±4 were said to have identical predictive performance to the best-fitting model. **a)** Testing the evidence for random slopes and quadratic *ΔAge* effects. **b)** Support for fixed effects in the scale component. **c)** Evidence for interactions between fixed effects. In **a)** we selected the model with the random slope unless there was substantial evidence against its inclusion (i.e., if it harmed predictive performance). In **c)** If models had similar predictive performance, then we selected the simplest model to explain variation in the response (*i.e, the model with no interactions)*. We selected **model 8** as the final model to explain variation in offspring growth rates, which only included single-effect predictors. In all models, n = 937 offspring from 77 parent pairs.

| Model selection based on predictive performance | | |
|---|---|---|
| ***Model*** | ***elpd (± SE)*** | ***Δelpd (± SE)*** |
| ***a)* Testing support for random slopes and quadratic age effects** *Using **a Gaussian** likelihood* | | |
| Model 1 | | |
| *Growth rate ~ $\overline{Age}$ + ΔAge + Temp + Sex + Mothermass + (1\|PairID)* | 665.6 ± 24.3 | -0.4 ± 1.5 |
| Model 2 | | |
| *Growth rate ~ $\overline{Age}$ + ΔAge + ΔAge$^2$ + Temp + Sex + Mothermass + (1\|PairID)* | 665.4 ± 24.3 | 0.0 ± 0.0 |
| **Model 3** | | |
| ***Growth rate ~ $\overline{Age}$ + ΔAge + Temp + Sex + Mothermass + (1+ ΔAge\|PairID)*** | **666.0 ± 24.4** | **-0.6 ± 1.5** |
| Model 4 | | |
| *Growth rate ~ $\overline{Age}$ + ΔAge + ΔAge$^2$ + Temp + Sex+ Mothermass + (1 + ΔAge\|PairID)* | 665.7 ± 24.3 | -0.3 ± 0.9 |
| ***b)* Evidence for fixed effects in the scale component** *Using **Model 3**: Gaussian model with single-effect predictors as a reference for the location component* | | |
| Model 5 σ (*Growth rate*) ~1 (No fixed effects on σ) | 665.4 ± 24.4 | 0.0 ± 0.0 |
| Model 6 σ (*Growth rate*) ~ *ΔAge* + *Temp* | 663.6 ± 24.5 | -1.9 ± 1.8 |
| **Model 7 σ (*Growth rate*) ~ *ΔAge*** | **665.2 ± 24.5** | **-0.2± 1.1** |
| ***c)* Evidence for interactions between fixed effects** *Using **Model 7** as a reference* | | |
| **Model 8** | | |
| **Location: *Growth rate ~ $\overline{Age}$ + ΔAge + Temp +Sex+ Mothermass + (1+ ΔAge\|PairID)* Scale: σ (*Growth rate*) ~ *ΔAge*** | **665.2 ± 24.4** | **0.0 ± 0.0** |
| Model 9 | | |
| Location: *Growth rate ~ $\overline{Age}$ + ΔAge + Temp +Sex+ Mothermass + ΔAge*Temp*Sex + (1+ ΔAge\|PairID)* Scale: σ (*Growth rate*) ~ *ΔAge* | 662.3 ± 24.4 | -2.9 ± 2.5 |
| Model 10 | | |
| Location: *Growth rate ~ $\overline{Age}$ + ΔAge + Temp + Sex + Mothermass+ ΔAge*Temp + (1+ ΔAge\|PairID)* Scale: σ (*Growth rate*) ~ *ΔAge* | 663.7 ± 24.4 | -1.5 ± 0.6 |
| Model 11 | | |
| Location: *Growth rate ~ $\overline{Age}$ + ΔAge + Temp + Sex + Mothermass + ΔAge*Sex + (1+ ΔAge\|PairID)* Scale: σ (*Growth rate*) ~ *ΔAge* | 664.5 ± 24.4 | -0.6 ± 0.5 |
| Model 12 | | |
| Location: *Growth rate ~ $\overline{Age}$ + ΔAge + Temp + Sex + Mothermass + ΔAge*Temp + (1+ ΔAge\|PairID)* Scale: σ (*Growth rate*) ~ *ΔAge* + *Temp* + *ΔAge*Temp* | 660.6 ± 24.5 | -4.6 ± 1.8 |
| Model 13 | | |
| Location: *Growth rate ~ $\overline{Age}$ + ΔAge + Temp + Sex + Mothermass + ΔAge*Sex + (1+ ΔAge\|PairID)* Scale: σ (*Growth rate*) ~ *ΔAge* + *Sex* + *ΔAge*Sex* | 664.4 ± 24.3 | -0.8 ± 2.2 |

**Table S20. Parameters explaining variation in offspring relative growth rate (RGR) from the selected model *(Model 8)*.** In the location component ($\mu$), we included the following as fixed effects: the parents' delta age, the parents' average age, the parents' temperature treatment, offspring sex, and the mother's adult body mass. Additionally, we also include the *Δslope* between the parents' average and delta age, as a test of selective disappearance. We set parent pair ID as a random intercept and the effect of *Δage* as a random slope for each parent pair. We also report the correlation between the random intercept and random slope (*r*). In the scale component, only *Δage* was included as a fixed effect predictor. Although dropped from the final model, we show the estimates for the effect of *Δage* in each temperature treatment and between each of the offspring sexes and report the pairwise comparisons for these interactions. Shown for each parameter is the posterior maximum a posteriori probability estimate ("*MAP*"), the 95% highest density intervals ("*95% HDI*"), along with the effective sample size ("*ESS [bulk/tail]*"), the probability of direction ("*pd*"), and the percentage of posterior estimates in ROPE *("% in ROPE"*). We highlight in bold any fixed effects with a pd > 98%, assuming these are genuine biological effects. n = 937 offspring from 77 parent pairs.

| Coefficients from the selected model (*Model 8*) | | | | | |
|---|---|---|---|---|---|
| ***Predictors*** | ***MAP*** | ***HDI (95%)*** | ***% in ROPE*[1]** | ***pd*[2]** | **ESS (Bulk/Tail)** |
| ***a) Location (μ) Model*** | | | | | |
| **Intercept** *Offspring RGR (week⁻¹)* | 0.76 | [0.74, 0.78] | 0% | 100% | 7724/7388 |
| **Parents' Δage** | **0.03** | **[0.02, 0.035]** | **0.06%** | **100%** | **13324/8286** |
| Parents' average age | 0.00 | [-0.01, 0.013] | 96.53 | 79.16% | 8675/7646 |
| $\Delta slope_{average\ age\ -\Delta age}$ | -0.02 | [-0.04, -0.01] | - | 99.99% | - |
| Mother's adult mass (g) | 0.00 | [-0.01, 0.01] | 96.21% | 80.34% | 7556/7627 |
| Parents' temperature treatment | | | | | |
| *25.5°C* | - | - | - | - | - |
| *28.0°C* | 0.00 | [0.00, 0.045] | 17.76% | 97.87% | 7614/7394 |
| *30.5°C* | 0.02 | [0.00, 0.042] | 23.48% | 96.04% | 7117/7241 |
| Offspring sex | | | | | |
| *Female* | - | - | - | - | - |
| ***Male*** | **-0.02** | **[-0.03, -0.002]** | **27.46%** | **98.45%** | **13722/8248** |
| **Random Effects** | | | | | |
| σ $Intercept_{Parent\ pair\ ID}$ | 0.02 | [0.00, 0.03] | - | 100% | 2141/1477 |
| σ Parents' $\Delta age_{Parent\ pair\ ID}$ | 0.01 | [0.00, 0.02] | - | 100% | 2183/3705 |
| *r* (Intercept ~ slope Δage) | -0.55 | [-0.75, 0.78] | - | 86.32% | 5218/5440 |
| ***Temperature Δage slopes and pairwise comparisons (from the dropped Parent's Δage x Temperature interaction)*** | | | | | |
| ***Δage*: *25.5°C*** | **0.02** | **[0.01, 0.04]** | **4.24%** | **99.94%** | **6207/6459** |
| ***Δage*: *28.0°C*** | **0.03** | **[0.02, 0.05]** | **1.05%** | **99.98%** | **-** |
| ***Δage*: *30.5°C*** | **0.02** | **[0.01, 0.04]** | **5.71%** | **99.93%** | **-** |
| *Δage: 25.5°C vs. 28.0°C* | 0.01 | [-0.01, 0.03] | 67.93% | 72.32% | 6971/6592 |

| | | | | | |
|---|---|---|---|---|---|
| *Δage: 28.0°C vs. 30.5°C* | 0.01 | [-0.01, 0.03] | 64.81% | 73.91% | - |
| *Δage: 30.5°C vs. 25.5°C* | 0.00 | [-0.02, 0.02] | 76.98% | 53.37% | 6775/6641 |
| ***Sex pairwise comparisons for the Δage slope (from the dropped Parent's* Δage x Offspring sex interaction)** | | | | | |
| ***Δage*: Female** | **0.03** | **[0.02, 0.04]** | **0.37%** | **100%** | **11333/8140** |
| ***Δage*: Male** | **0.02** | **[0.01, 0.04]** | **2.93%** | **100%** | **-** |
| *Δage: Female vs. Male* | 0.00 | [-0.02, 0.01] | 82.29% | 68.08% | 10857/8096 |
| ***b) Scale (σ) Model*** | | | | | |
| **Intercept** *Log (residual SD)* | -2.15 | [-2.21, -2.10] | - | 100% | 7724/7388 |
| Parents' Δage | 0.03 | [-0.02, 0.08] | - | 86.32% | 10652/7986 |
| **Marginal $R^2$** | 0.06 | [0.03, 0.09] | - | - | - |
| **Conditional $R^2$** | 0.09 | [0.05, 0.14] | - | - | - |

[1]ROPE = [−0.012, 0.012] on unit scale (RGR; week$^{-1}$); applied to location fixed effects only.

[2]pd = probability of direction.

#### Section S13: Supplementary results for offspring fecundity

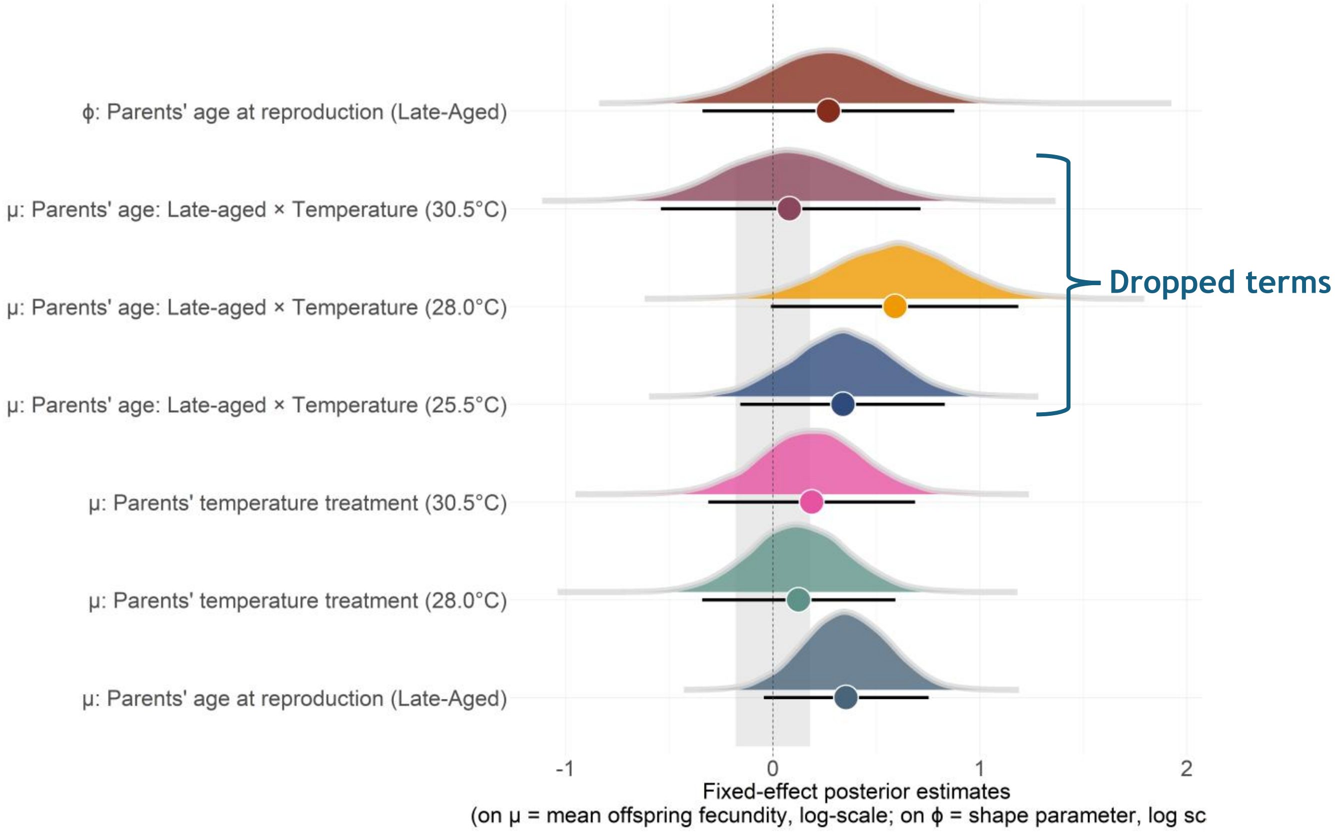

**Figure S23. Posterior distributions of fixed effects from the negative binomial model explaining variation in offspring fecundity.** Shown are the fixed effects from both the location ($\mu$) and scale ($\phi$) component. For each fixed effect, circles show the posterior median, while error bars describe the 95% HDI's. The dotted line represents the line of no effect, or the statistical intercept of the model. The shaded area denotes the region of practical equivalence (ROPE), describing the region of negligible effect (± 0.13 log-eggs). For $\mu$ parameters, the intercept represents the log (mean number of eggs) laid by offspring from early-aged parents kept at 25.5 °C. For $\phi$, the intercept is the log ($\phi$) for offspring from early-aged parents. Although not included in our final model, we show the posterior estimates for the conditional effect of parental age under each temperature treatment. N = clutches from 83 offspring descending from 45 parent pairs.

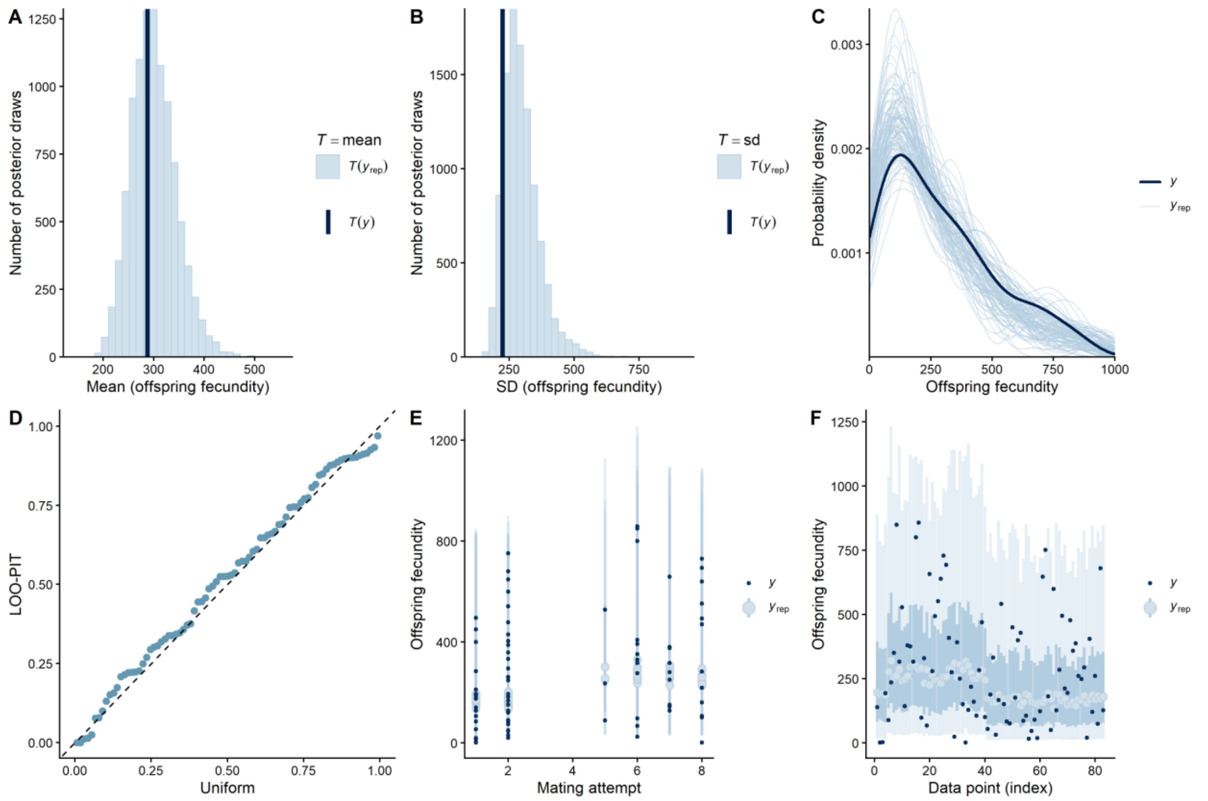

**Figure S24. Model fit plots for offspring fecundity (from Model 6, *Table S21*). A) & B) Histogram of the means & standard deviations (SD) from the posterior predictive and observed distribution.** The posterior simulated means/SD's from the posterior distribution are shown in light blue (*$T(Y_{rep})$*), while the empirical mean from the observed distribution is shown in dark blue *T(y)*). **C) Probability density of the empirical and posterior simulated data.** The thin light blue lines describe the simulated range of values for offspring fecundity, estimated from 100 posterior draws. The dark blue line represents the observed distribution of offspring fecundity. **D) Quantile-Quantile (Q-Q plot) of the LOO-PIT values.** The dashed diagonal line denotes the expected uniform distribution of the LOO-PIT integrals, while each individual point represents the observed value of the LOO-PIT integral. **E) Posterior predictive interval plot.** On the y-axis is the offspring's fecundity, while the x-axis describes the parents' mating attempt (here, as two categories: "early" – mating attempts 1 & 2, "late": mating attempts 5 through to 8). Small dots represent the observed data, while light blue points and error bars describe the posterior predicted value and credible intervals (50% & 95% CI's) for each observation. **F) LOO interval plot.** On the x-axis is each observed value, while on the y-axis is the estimate of the response variable (fecundity). The dark points represent the observed value of each observation, while the light blue points and bars represent the LOO predicted data and credible intervals (50% and 95% CI's). N = 83 offspring from 45 parent pairs.

**Table S21. Comparison of the predictive performance of models explaining variation in offspring fecundity (*Eggslaid*).** Highlighted in bold is the model selected from each stage of the analysis, which we then carried forward as a reference model to the next stage. The following were included as fixed effects: The parents' age category (*ParentalAge*) and the parents' temperature treatment (*Temp*). We also included parent pair ID as a random intercept (*1|Pair ID*). For each model, shown is the expected log pointwise predictive density ("*elpd*"), the Δelpd relative to the best-fitting model, along with the standard errors *("SE")*. Models with a Δelpd <±4 were said to have identical predictive performance to the best-fitting model. **a)** Comparison of model likelihoods. **b)** Support for fixed effects in the scale component ($\phi$). **c)** Evidence for interactions between fixed effects. In ***c)*** If models had similar predictive performance, then we selected the simplest model to explain variation in the response (*i.e, the model with no interactions)*. We selected **Model 6** as our final model to explain variation in offspring fecundity, which only included single-effect predictors. In all models, egg batches from n = 83 offspring descending from 45 parent pairs.

| Model selection based on predictive performance | | |
|---|---|---|
| ***Model*** | ***elpd (± SE)*** | ***Δelpd (± SE)*** |

| | | |
|---|---|---|
| ***a) Comparison of model likelihoods and calculated using 10-fold cross-validation*** | | |
| Model 1<br>*Negative binomial* | **-561.2 ± 7.4** | **0.0 ± 0.0** |
| Model 2<br>*Poisson* | -6649.3 ± 1530.2 | -6088.1 ± 1528.4 |
| **b) Evidence for fixed effects in the scale component - *calculated with LOO-CV***<br>*Using **Model 1**: Negative binomial model with single-effect predictors as a reference* | | |
| Model 3<br>Location*: Eggcounts ~ ParentalAge* + *Temp* + (*1/PairID*)<br>Scale*: ϕ* (*Eggcounts*) *~1 (No fixed effects on ϕ)* | -555.5 ± 7.2 | 0.0 ± 0.0 |
| Model 4<br>Location: Eggcounts ~ ParentalAge + *Temp* + (*1/PairID*)<br>Scale: *ϕ* (*Eggcounts*) ~ *ParentalAge+ Temp* | -556.8 ± 7.4 | -1.2 ± 3.6 |
| **Model 5**<br>**Location: *Eggcounts ~ ParentalAge* + *Temp* + (*1/PairID*)**<br>**Scale: *ϕ* (*Eggcounts*) ~ *ParentalAge*** | **-556.8 ± 7.4** | **-1.3 ± 1.8** |
| ***d)* Evidence for interactions between fixed effects**<br>*Using **Model 5** as a reference* | | |
| Model 6<br>**Location: *Eggcounts* ~ *ParentalAge* + *Temp* + (*1/PairID*)**<br>**Scale: *ϕ* (*Eggcounts*) ~ *ParentalAge*** | **-556.1 ± 7.3** | **-0.3 ± 0.9** |
| Model 7<br>Location: *Eggcounts ~ ParentalAge + Temp + ParentalAge*Temp + (1\|PairID)*<br>Scale: *ϕ* (Eggcounts) ~ *ParentalAge* | -555.8 ± 7.4 | 0.0 ± 0.0 |

**Table S22. Parameters explaining variation in offspring fecundity from the selected model *(Model 6)***. In the location component (*μ*), the model included the following as fixed effect predictors: parental age (*"parents' age category")* and parental temperature. Parent pair ID was included as a random intercept. In the scale component (*ϕ*), we only included parental age as a fixed effect. Shown for each parameter is the maximum a posteriori probability estimate ("*MAP*"), the 95% highest density intervals ("*95% HDI*"), along with the effective sample size ("*ESS [bulk/tail]*"), probability of direction ("*pd*"), and the percentage of posterior estimates in ROPE ("*% in ROPE*"). We highlight in bold any predictors with pd values > 98%, assuming these represent real biological effects. Additionally, we show the estimates for the dropped interaction between parental age and parental temperature. n = clutches from 83 offspring descending from 45 parent pairs.

| **Coefficients from the selected model (Model 6)** | | | | | |
|---|---|---|---|---|---|
| ***Predictors*** | ***MAP* ± SE** | ***HDI (95%)*** | ***% in ROPE*[1]** | ***pd*[2]** | **ESS (Bulk/Tail)** |
| ***a) Location (μ) Model*** | | | | | |
| **Intercept**<br>*Log (number eggs laid)* | 5.39 | [4.99, 5.80] | 0% | 100% | 9185/7021 |
| Parents' age category | | | | | |
| *Early-Aged* | - | - | - | - | - |
| *Late-Aged* | 0.36 | [-0.04, 0.75] | 11.99% | 95.79% | 12412/6772 |
| Parents' temperature treatment | | | | | |
| *25.5°C* | - | - | - | - | - |

| | | | | | |
|---|---|---|---|---|---|
| *28.0°C* | 0.13 | [-0.34, 0.59] | 40.16% | 70.05% | 9534/7257 |
| *30.5°C* | 0.19 | [-0.31, 0.69] | 33.24% | 77.72% | 9977/7003 |
| **Random Effects** | | | | | |
| σ $\text{Intercept}_{\text{Parent pair ID}}$ | 0.18 | [0.01, 0.56] | - | - | 2745/3247 |
| ***Temperature specific parental age effects and pairwise comparisons (from the dropped Parent's age x Temperature interaction)*** | | | | | |
| *Parents' Age*: *Late-aged (25.5°C)* | 0.34 | [-0.16, 0.83] | 18.55% | 90.64% | 11168/8171 |
| *Parents' Age*: *Late-aged (28.0°C)* | 0.61 | [-0.04, 1.15] | 5.67% | 97.32% | - |
| *Parents' Age*: *Late-aged (30.5°C)* | 0.07 | [-0.52, 0.73] | 31.65% | 60.07% | - |
| *25.5°C vs. 28.0°C* | 0.23 | [-0.43, 0.91] | 23.59% | 76.87% | 12461/8531 |
| *28.0°C vs. 30.5°C* | 0.48 | [-0.30, 1.35] | 11.99% | 88.61% | - |
| *30.5°C vs. 25.5°C* | -0.25 | [-0.93, 0.44] | 22.61% | 76.27% | 11354/7610 |
| ***b) Scale (ϕ) Model*** | | | | | |
| **Intercept** *Log (ϕ)* | 0.09 | [-0.31, 0.47] | - | 67.00% | 12028/6772 |
| Parents' age category | | | | | |
| *Early-Aged (reference category)* | - | - | - | - | - |
| *Late-Aged* | 0.27 | [-0.34, 0.88] | - | 81.22% | 11228/6787 |
| **Marginal $R^2$** | 0.10 | [0.08, 0.153] | - | - | - |
| **Conditional $R^2$** | 0.18 | [0.13, 0.23] | - | - | - |

[1]ROPE = [−0.134, 0.134] on log scale; applied to location fixed effects only.

[2]pd = probability of direction.

#### Section S14: Supplementary results for offspring hatching success

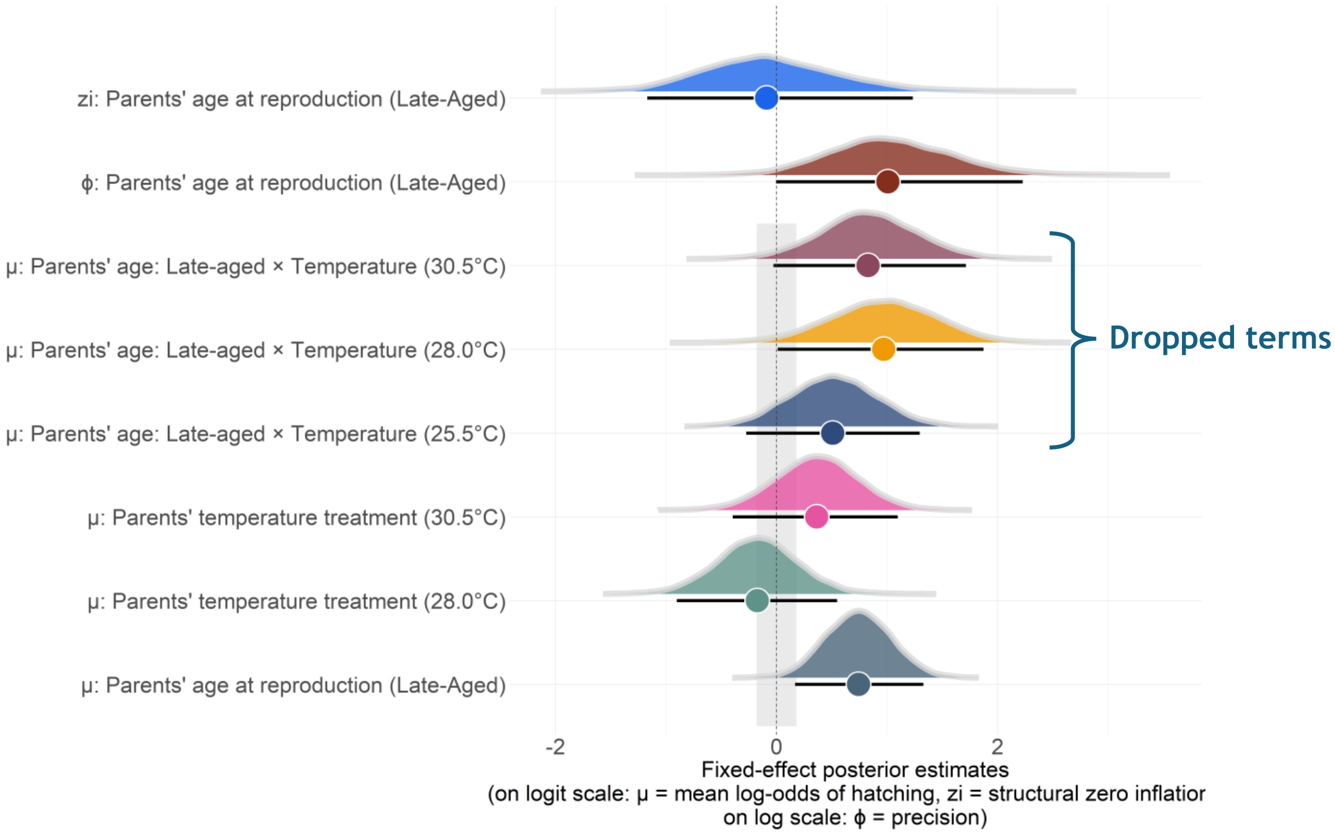

**Figure S25. Posterior distributions of fixed effects from the zero-inflated beta binomial model explaining variation in offspring hatching success.** Shown are the fixed effects from the location ($\mu$), scale ($\phi$), and zero-inflation ($z$) submodels. For each fixed effect, circles depict the posterior median, while the error bars describe the 95% HDI's. The dotted line describes the line of no effect, or the models statistical intercept. The shaded area is the region of practical equivalence (i.e., the ROPE) used for fixed effects in the location submodel, describing the area of negligible effect (± 0.18 log-odds). For $\mu$ parameters, the intercept is the log-odds of hatching for eggs laid by offspring from early-aged parents kept at 25.5 °C. For $\phi$, the intercept is the log ($\phi$) in hatching success for offspring from early-aged parents. For *zi*, the intercept is the log odds of structural zero hatchlings (i.e., complete hatching failure) for offspring from early-aged parents. Additionally, we show the conditional estimates of the parental age effects in each temperature treatment (although this interaction was removed from our final model). Eggs from n = Eggs from 82 offspring descending from 45 parent pairs.

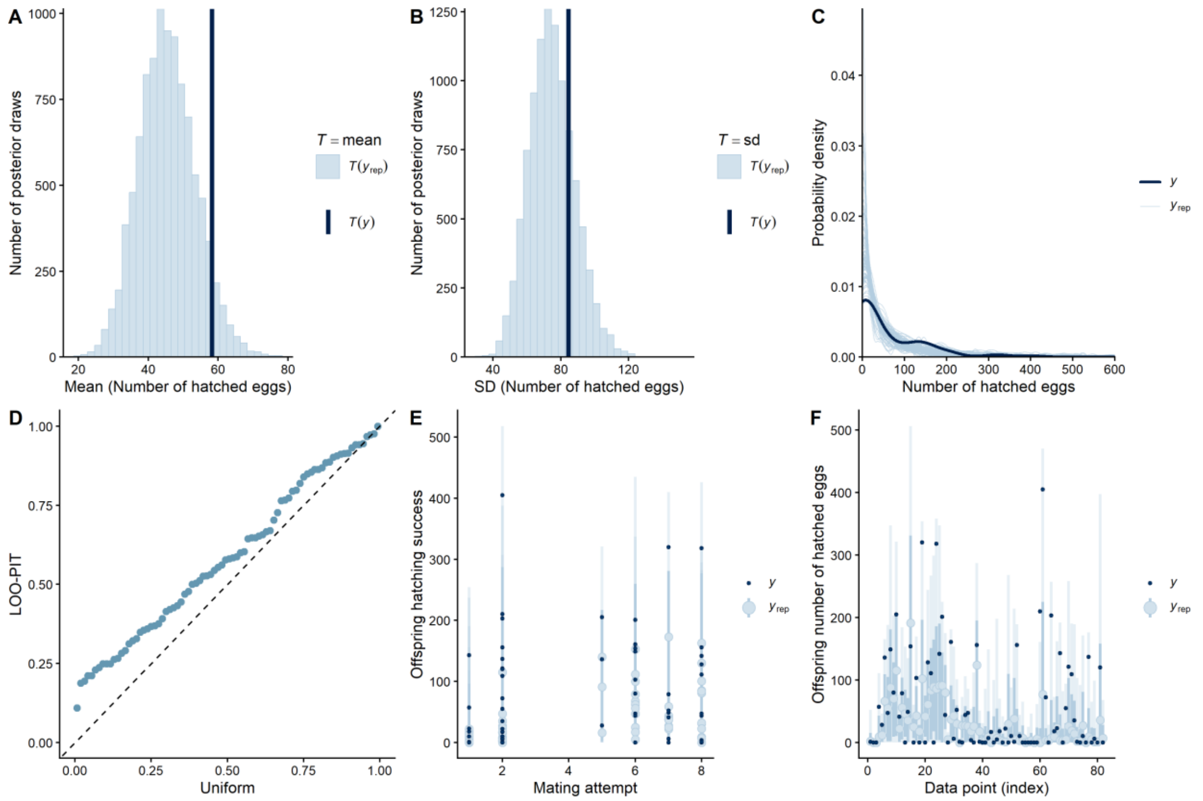

**Figure S26. Model fit plots for offspring hatching success (from Model 7, *Table S23*).** Note that although this model is not a perfect fit, the zero-inflated beta-binomial was the best fitting model of all possible likelihoods that could be fitted to this trial data in *brms*. **A) and B) Histogram of the means and standard deviations from the posterior simulated and empirical distribution.** The light blue lines show the simulated means/SD's from the posterior distribution (*T*($Y_{rep}$)), while the empirical mean from the observed distribution is shown in dark blue *T(y)*). **B) The probability density of the empirical and posterior simulated data.** The thin light blue lines describe the simulated range of values for offspring hatching success, as estimated from 100 posterior draws. The dark blue line describes the observed distribution of the number of hatched eggs. **D) Quantile-Quantile (Q-Q plot) of the LOO-PIT values.** The dashed diagonal line describes the uniform distribution of LOO-PIT integrals, while the individual points represent the observed values of each LOO-PIT integral. This model is heavily underdispersed in some regions, expecting more variation than what is observed. **E) Posterior predictive interval plot.** On the y-axis is the number of hatched eggs, while the x-axis describes the parents' mating attempt (here, as two categories: "early" – mating attempts 1 & 2, "late": mating attempts 5 through to 8). Small dots represent the observed data, while light blue points and error bars describe the posterior predicted value and credible intervals (50% & 95% CI's) for each observation. **F) LOO interval plot.** On the x-axis is each observed value, while on the y-axis is the estimate of the response variable (number of hatched eggs). The dark points represent the observed value of each observation, while the light blue points and bars represent the LOO predicted data and credible intervals (50% and 95% CI's). N = Eggs from 82 offspring pairs descending from 45 parent pairs.

**Table S23. Comparison of the predictive performance of models explaining variation in the hatching success of eggs laid by the offspring (*HatchSuccess*).** Highlighted in bold is the model selected from each stage of the analysis, which we then carried forward as a reference model to the next stage. We included the following as fixed effects: The parents' age category (*ParentalAge*) and the parents' temperature treatment (*Temp*). We also included parent pair ID as a random intercept (*1|Pair ID*). For each model, shown is the expected log pointwise predictive density ("*elpd*"), the Δelpd relative to the best-fitting model, along with the standard errors *("SE"*). Models with a Δelpd <±4 were said to have identical predictive performance to the best-fitting model. **a)** Comparison of model likelihoods. **b)** Support for fixed effects in the scale ($\phi$) and zero-inflation ($z$) component. **c)** Evidence for interactions between fixed effects. In **c)** If models had similar predictive performance, then we selected the simplest model to explain variation in the response (*i.e, the model with no* *interactions)*. We selected **Model 7** as our final model to explain variation in offspring hatching success, which only included single-effect predictors. In all models, egg batches from n = 82 offspring descending from 45 parent pairs.

| Model selection based on predictive performance | | |
|---|---|---|
| ***Model*** | ***elpd (± SE)*** | ***Δelpd (± SE)*** |
| ***a) Comparison of model likelihoods*** | | |
| Model 1<br>*Binomial* | -2321.0 ± 433.9 | -1998.8 ± 426.0 |
| Model 2<br>*Beta-binomial* | -330.8 ± 22.0 | -8.6 ± 4.1 |
| **Model 3**<br>***Zero-inflated beta-binomial (ZIBB)*** | **-322.2 ± 21.2** | **0.0 ± 0.0** |
| **b) Evidence for fixed effects in the scale and zero-inflation component (*dpars*)**<br>*Using **Model 3**: ZIBB model with single-effect predictors as a reference* | | |
| Model 4<br>Location*: HatchSuccess ~ ParentalAge* + *Temp* + (*1\|PairID*)<br>dpars*: ϕ/z* (*HatchSuccess*) *~1 (No fixed effects on ϕ/z)* | -323.5 ± 21.3 | -3.4 ± 2.5 |
| Model 5<br>Location: *HatchSuccess* ~ ParentalAge + *Temp* + (*1\|PairID*)<br>dpars: *ϕ/z* (*HatchSuccess*) ~ *ParentalAge*+ *Temp* | -325.5 ± 21.1 | -5.3 ± 1.1 |
| **Model 6**<br>**Location: *HatchSuccess ~ ParentalAge* + *Temp* + (*1\|PairID*)**<br>**dpars: *ϕ/z* (*HatchSuccess*) ~ *ParentalAge*** | **-320.1 ± 21.2** | **0.0 ± 0.0** |
| ***c)* Evidence for interactions between fixed effects**<br>*Using **Model 6** as a reference* | | |
| **Model 7**<br>**Location: *HatchSuccess* ~ *ParentalAge* + *Temp* + (*1\|PairID*)**<br>**dpars: *ϕ/z* (*HatchSuccess*) ~ *ParentalAge*** | **-320.2 ± 21.0** | **0.0 ± 0.0** |
| Model 8<br>Location: *HatchSuccess* ~ *ParentalAge* + *Temp* + *ParentalAge*Temp* + *(1\|PairID)*<br>dpars: *ϕ/z* (*HatchSuccess*) ~ *ParentalAge* | -321.9 ± 21.1 | -1.7 ± 1.1 |

**Table S24. Coefficients explaining variation in the hatching success of the eggs laid by the offspring from the selected model *(Model 7)*.** In the location component ($\mu$), the model included the following as fixed effects: parental age and parental temperature. Parent pair ID was included as a random intercept. In both the scale component ($\phi$) and zero-inflation ($z$) component, only parental age was included as a fixed effect. For each parameter, we show the posteriori probability estimate ("*MAP*"), the standard errors ("*SE*"), the 95% highest density intervals ("*95% HDI*"), along with the effective sample size ("*ESS [bulk/tail]*"), the probability of direction ("*pd*") and the percentage of posterior estimates in ROPE ("*% in ROPE*"). We highlight in bold any fixed effects with pd values >98%, assuming these reflect genuine biological effects. Additionally, we show the estimates for the dropped interaction between parental age and parental temperature. Eggs from n = 82 offspring descending from 45 parent pairs.

| Coefficients from the selected model (Model 7) | | | | | |
|---|---|---|---|---|---|
| ***Predictors*** | ***MAP*** | ***HDI (95%)*** | ***% in ROPE[1]*** | ***pd[2]*** | **ESS (Bulk/Tail)** |
| ***a) Location (μ) Model*** | | | | | |
| **Intercept** | -1.84 | [-2.61, -1.22] | 0% | 100% | 5962/6855 |

| | | | | | |
|---|---|---|---|---|---|
| *Log-odds of successful hatching* | | | | | |
| Parents' age category | | | | | |
| *Early-Aged* | - | - | - | - | - |
| *Late-Aged* | 0.74 | [0.17, 1.33] | 2.74% | 99.41% | 9538/7987 |
| Parents' temperature treatment | | | | | |
| *25.5°C* | - | - | - | - | - |
| *28.0°C* | -0.16 | [-0.91, 0.53] | 34.45% | 69.05% | 7421/5908 |
| *30.5°C* | 0.36 | [-0.36, 1.12] | 23.35% | 83.17% | 6025/6327 |
| **Random Effects** | | | | | |
| σ Intercept$_{\text{Parent pair ID}}$ | 0.65 | [0.05, 1.11] | - | 100% | 2745/3247 |
| ***Temperature specific parental age effects and pairwise comparisons (from the dropped Parent's age x Temperature interaction)*** | | | | | |
| *Parents' Age: Late-aged (25.5°C)* | 0.51 | [-0.27, 1.30] | 17.29% | 89.69% | 4764/6736 |
| *Parents' Age: Late-aged (28.0°C)* | 1.03 | [0.02, 1.878] | 4.19% | 97.61% | - |
| *Parents' Age: Late-aged (30.5°C)* | 0.78 | [0.00, 1.743] | 5.75% | 97.09% | - |
| *25.5°C vs. 28.0°C* | 0.53 | [-0.56, 1.525] | 18.41% | 79.96% | 5267/6733 |
| *28.0°C vs. 30.5°C* | 0.12 | [-1.08, 1.315] | 22.78% | 58.29% | - |
| *30.5°C vs. 25.5°C* | 0.41 | [-0.72, 1.304] | 22.43% | 73.58% | 5790/6220 |
| ***b) Scale (ϕ) Model*** | | | | | |
| **Intercept** *Log (ϕ)* | 1.37 | [0.71, 2.10] | - | 100% | 4970/7236 |
| Parents' age category | | | | | |
| *Early-Aged* | - | - | - | - | - |
| *Late-Aged* | 0.92 | [0.01, 2.24] | - | 97.44% | 2888/3229 |
| ***c) Zero-inflation (z) Model*** | | | | | |
| ***Intercept*** *Log odds of structural zeroes* | -1.07 | [-2.44, -0.30] | - | 99.71% | 7094/5288 |
| *Parents' age category* | | | | | |
| *Early-Aged* | - | - | - | - | - |
| *Late-Aged* | -0.11 | [-1.23, 1.17] | - | 56.44% | 8972/6971 |

| | | | | | |
|---|---|---|---|---|---|
| **Marginal $R^2$** | 0.28 | [0.15, 0.416] | - | - | - |
| **Conditional $R^2$** | 0.42 | [0.23, 0.621] | - | - | - |

[1]ROPE = [−0.18, 0.18] on log-odds scale; applied to location fixed effects only.

[2]pd = probability of direction.

#### Section S15: Supplementary results for offspring adult lifespan and mortality

##### *S15.1 Supplementary results for offspring mean adult lifespan*

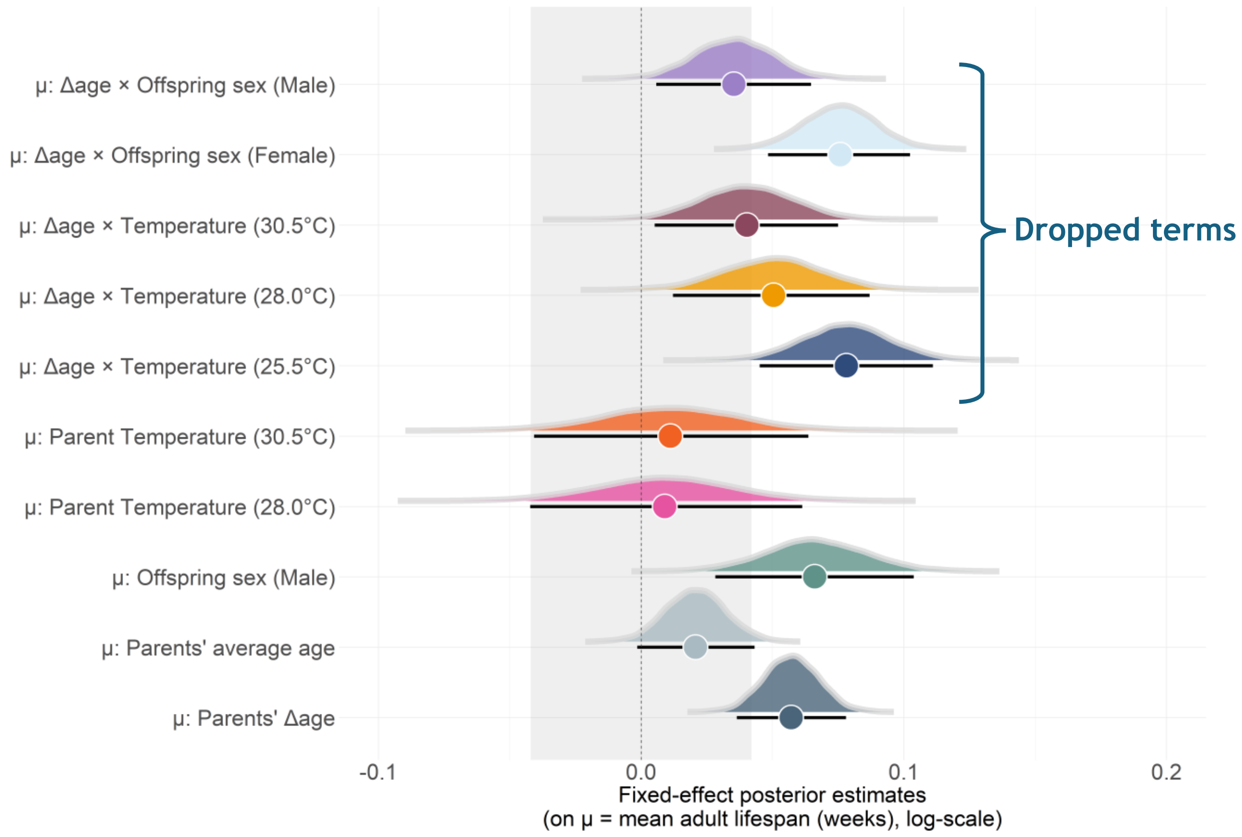

**Figure S27. Posterior distributions for the fixed effects from the Weibull model explaining variation in offspring adult lifespan.** For each fixed effect, the circles describe the posterior median, while error bars denote the 95% HDI's. The dotted line is the line of no effect, representing the models statistical intercept. Here, the intercept is the log (mean adult lifespan) for daughters of parents maintained at 25.5 °C and when *Δage* = *0*. The shaded area describes the region of practical equivalence (i.e., ROPE), indicating the area of negligible effect (± 0.04 log weeks). Although removed from our final model, we show the conditional *Δage* slopes for each temperature treatment, and for each offspring sex. N = 938 offspring from 77 parent pairs.

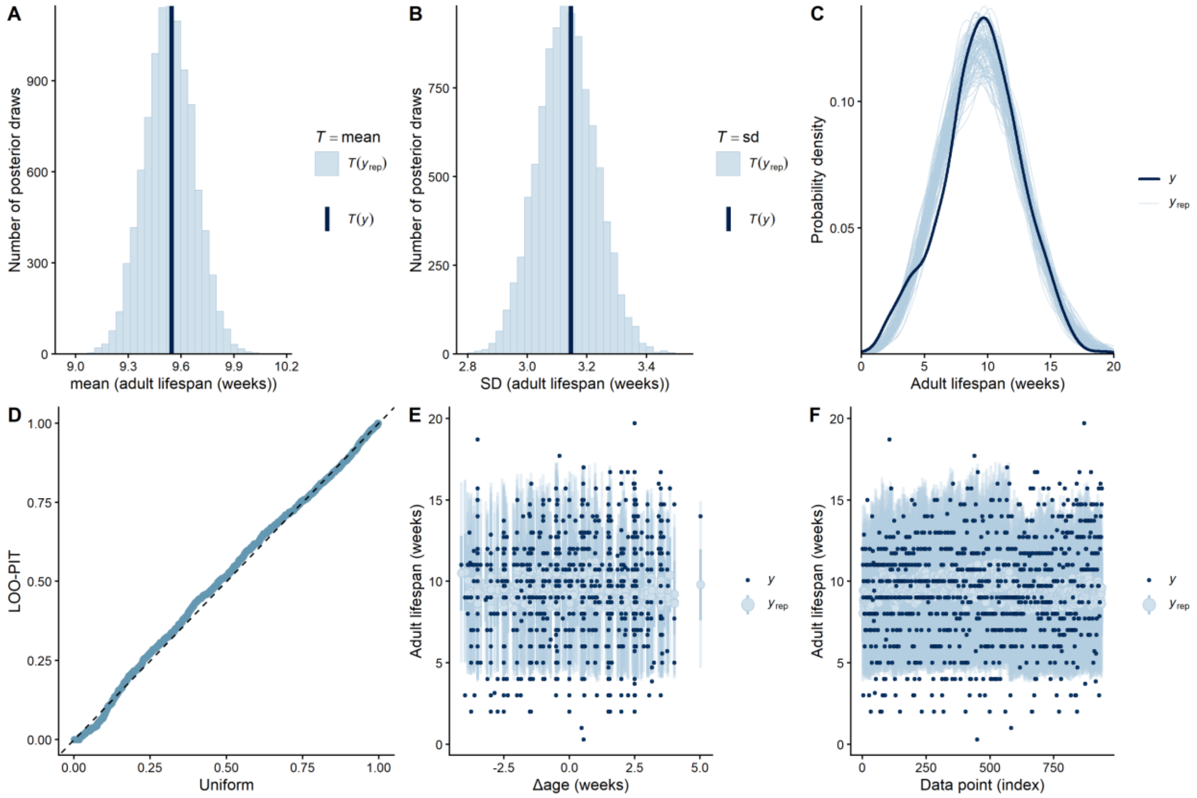

**Figure S28. Model fit plots for adult lifespan (Model 7, Table S25). A) & B) Histogram of means and standard deviations from the posterior predictive and observed distribution.** Shown are the simulated means/SD's for adult longevity from the posterior replicated data sets (*T(*$Y_{rep}$*)*; light blue), while the dark vertical line is the empirical mean (*T(y)*). **C) Probability density of the empirical and posterior simulated data sets.** Light blue lines describe the distribution of the response (lifespan) as estimated from 100 simulated posterior draws. The dark blue line is the observed distribution of offspring total lifespan. **D) Quantile-Quantile (Q-Q plot) describing the distribution of the Leave-One-Out Probability Integrals (LOO-PIT).** The dashed diagonal line is the expected uniform distribution under a well-specified model, while the blue points represent the observed distribution of the LOO-PIT integrals. **E) Posterior predictive interval plot.** On the y-axis is the offspring's adult lifespan, while the x-axis describes the *Δage* effect. Small dots represent the observed data, while light blue points and error bars describe the posterior predicted value and credible intervals (50% & 95% CI's) for each observation. **F) LOO interval plot.** This plot is used to describe the ability of the model to predict each individual observation. Shown on the x-axis is each observation, while on the y-axis is the estimate of the response variable (adult lifespan). The dark points represent the observed value of each observation, while the light green points and bars represent the LOO predicted data and credible intervals (both 50% and 95% CI's). N = 938 offspring from 77 parent pairs.

**Table S25. Comparison of the predictive performance of *brms* models explaining variation in offspring adult lifespan (*AdultLife*).** We highlight in bold the model that was selected from each stage of the analysis, which we then carried forward to use as a reference for the next stage. We included the following as fixed effects: The parents' delta age at reproduction (*ΔAge*), the parents' average age at reproduction $\overline{(Age)}$, the parents' temperature treatment (*Temp*), Offspring sex (*Sex*), and the quadratic effect of delta age ($\Delta Age^2$). Additionally, we included parent pair ID as a random intercept (*1|Pair ID*) and *ΔAge* as a random slope (*1 + ΔAge |Pair ID*). For each model, we show the expected log pointwise predictive densities ("*elpd*"), the Δelpd relative to the best-fitting model, along with the standard errors *("SE")*. Models with a Δelpd <±4 were said to have identical predictive performance to the best-fitting model. **a)** Comparison of model likelihoods. **b)** Testing the evidence for random slopes and quadratic *ΔAge* effects. **c)** Evidence for interactions between fixed effects. In **b)** we selected the model with the random slope unless there was substantial evidence its inclusion harmed predictive performance. In **c)** If models had similar predictive performance, then we selected the simplest model to explain variation in the response (*i.e, the model with no interactions)*. We selected **model 7**, which only included single-effect predictors, as the final model to explain variation in adult lifespan. In all models, n = 938 offspring from 77 parent pairs.

| Model selection based on predictive performance | | |
|---|---|---|
| ***Model*** | ***elpd (± SE)*** | ***Δelpd (± SE)*** |
| ***a) Comparison of model likelihoods (using default brms priors)*** | | |
| **Model 1** ***Weibull*** | **-2392.2± 23.1** | **0.0 ± 0.0** |
| Model 2 *Exponential* | -2557.6 ± 41.5 | -663.3 ± 26.3 |
| Model 3 *Lognormal* | -3055.5 ± 10.2 | -165.4 ± 28.7 |
| ***b)* Testing support for random slopes and quadratic age effects** *Using **Model 1**: Weibull model as a reference* | | |
| Model 1 *AdultLife~* $\overline{Age}$ *+ ΔAge + Temp + Sex + (1\|PairID)* | -2392.2 ± 23.1 | -0.8 ± 1.9 |
| Model 4 *AdultLife~* $\overline{Age}$ *+ ΔAge + ΔAge*$^2$ *+ Temp + Sex+ (1\|PairID)* | -2391.4 ± 23.1 | 0.0 ± 0.0 |
| **Model 5** ***AdultLife~*** $\overline{Age}$ ***+ ΔAge + Temp + Sex + (1+ ΔAge\|PairID)*** | **-2392.4± 23.2** | **-1.0 ± 2.0** |
| Model 6 *AdultLife~* $\overline{Age}$ *+ ΔAge + ΔAge*$^2$ *+ Temp + Sex + (1 + ΔAge\|PairID)* | -2391.8 ± 23.1 | -0.4 ± 0.8 |
| ***c)* Evidence for interactions between fixed effects** *Using **Model 5**: Weibull model with single-effect predictors only* | | |
| **Model 7** ***AdultLife ~ ~*** $\overline{Age}$ ***+ ΔAge + Temp +Sex + (1+ ΔAge\|PairID)*** | **-2392.4 ± 23.1** | **-1.1 ± 2.1** |
| Model 8 *AdultLife~ ~* $\overline{Age}$ *+ ΔAge + Temp + Sex+ ΔAge*Temp*Sex + (1+ ΔAge\|PairID)* | -2394.2 ± 23.1 | -2.9 ± 2.2 |
| Model 9 *AdultLife~ ~* $\overline{Age}$ *+ ΔAge + Temp + Sex+ ΔAge*Temp + (1+ ΔAge\|PairID)* | -2393.0 ± 23.1 | -1.6 ± 2.9 |
| Model 10 *AdultLife~ ~* $\overline{Age}$ *+ ΔAge + Temp + Sex+ ΔAge*Sex + (1+ ΔAge\|PairID)* | -2391.3 ± 23.4 | 0.0 ± 0.0 |

**Table S26. Parameters explaining variation in the adult lifespan of offspring from the selected model *(Model 7)*.** The following were included as fixed effects: the parents' delta age, the parents' average age, the parents' temperature treatment, and offspring sex. We additionally report the *Δslope* between the parents' average and delta age, used as a test of selective disappearance. Parent pair ID was included as random intercept, while we also included a random slope for the parents' delta age. Covariances between the random intercept and random slope are described with the Pearsons's correlation coefficient, *r*. Shown for each parameter is the maximum a posteriori probability estimate ("*MAP*"), the 95% highest density intervals ("*95% HDI*"), along with the ESS (bulk/tail), the probability of direction ("*pd*"), and the percentage of the posterior estimates in ROPE ("*% in ROPE*"). Additionally, we show the estimates for the dropped interaction between parental age and parental temperature, and between parental age and offspring sex. Any fixed effects with a pd > 98% are highlighted in bold, assuming these represent genuine biological effects. n = 938 offspring from 77 parent pairs.

| Coefficients from the selected model (*Model 7*) | | | | | |
|---|---|---|---|---|---|
| ***Predictors*** | ***MAP*** | ***HDI (95%)*** | ***% in ROPE***[1] | ***pd***[2] | **ESS (Bulk/Tail)** |
| ***a) Location (μ) Model*** | | | | | |

| | | | | | |
|---|---|---|---|---|---|
| **Intercept**<br>*Log (adult lifespan [weeks])* | 2.21 | [2.17, 2.25] | 0% | 100% | 8446/7484 |
| **Parents' Δage** | **0.06** | **[0.04, 0.08]** | **5.27%** | **100%** | **1230/7849** |
| Parents' average age | 0.02 | [-0.00, 0.04] | 96.83% | 96.68% | 10945/8007 |
| **$\Delta slope_{average\ age\ -\Delta age}$** | **-0.04** | **[-0.06, -0.01]** | **-** | **98.89%** | **-** |
| Parents' temperature treatment | | | | | |
| *25.5°C* | - | - | | - | - |
| *28.0°C* | 0.01 | [-0.04, 0.06] | 87.01% | 63.85% | 8908/6990 |
| *30.5°C* | 0.01 | [-0.04, 0.06] | 86.12% | 66.28% | 7985/7872 |
| *Offspring sex* | | | | | |
| *Female* | - | - | - | - | - |
| ***Male*** | **0.07** | **[0.03, 0.10]** | **10.19%** | **99.97%** | **13677/7872** |
| **Random Effects** | | | | | |
| σ $Intercept_{Parent\ pair\ ID}$ | 0.05 | [0.00, 0.07] | - | 100% | 1804/2925 |
| σ Parents' $\Delta age_{Parent\ pair\ ID}$ | 0.02 | [0.00, 0.05] | - | 100% | 2525/3611 |
| *r* (*Intercept ~ slope Δage)* | 0.26 | [-0.72, 0.83] | - | 58.73% | 5711/6325 |
| ***Temperature Δage slopes and pairwise comparisons (from the dropped Parent's Δage x Temperature interaction)*** | | | | | |
| ***Δage*: 25.5°C** | **0.08** | **[0.05, 0.11]** | **1.6%** | **100%** | **5704/7014** |
| ***Δage*: 28.0°C** | **0.05** | **[0.01, 0.09]** | **33.04%** | **99.49%** | **-** |
| ***Δage*: 30.5°C** | **0.04** | **[0.01, 0.08]** | **53.81%** | **98.64%** | **-** |
| *Δage*: *25.5°C vs. 28.0°C* | -0.03 | [-0.08, 0.02] | 71.14% | 86.27% | 6073/7132 |
| *Δage*: *28.0°C vs. 30.5°C* | 0.01 | [-0.04, 0.06] | 86.68% | 64.75% | - |
| *Δage*: *30.5°C vs. 25.5°C* | -0.03 | [-0.09, 0.01] | 56.74% | 93.82% | 6595/7040 |
| ***Sex Δage slopes and pairwise comparisons (from the dropped Parent's* Δage x Offspring sex interaction)** | | | | | |
| ***Δage*: Female** | **0.08** | **[0.05, 0.10]** | **50.17%** | **98.87%** | **8190/7477** |
| ***Δage*: Male** | **0.04** | **[0.01, 0.07]** | **67.54%** | **98.87%** | **-** |
| *Δage*: *Female vs. Male* | -0.04 | [-0.08, 0.00] | 53.03% | 97.84% | 8442/7351 |
| **Shape parameter (*k*)** | 3.50 | [3.31, 3.68] | - | - | 8902/8001 |
| **Marginal $R^2$** | 0.04 | [0.02, 0.07] | - | - | - |
| **Conditional $R^2$** | 0.06 | [0.03, 0.10] | - | - | - |

[1]ROPE = [−0.042, 0.042] on log scale; applied to location fixed effects only.

[2]pd = probability of direction.

##### *S15.2 Supplementary results for offspring adult mortality*

We report the exact estimates for mortality parameters in *Table S28*. The model including the interaction between parental age and parental temperature received little statistical support, with the DIC value for this model (DIC = -2595.79) being higher than both the isolated effect of parental age (DIC = -2617.67). Thus, we concluded that there was no meaningful evidence that the effect of parental age was temperature dependent. However, there appeared to be visual evidence for an interactive effect between the two terms (*Figure S29: bottom*). Parental age appeared to have a slight positive effect on offspring adult mortality when the parents were maintained under 25.5°C, but not at 28.0°C or 30.5°C. Despite this, visual inspection of the posteriors for each mortality parameter (*Figure S29: top*) revealed that the direction of the effect was comparable for *b1*, *b0*, and *c*, though the posteriors for each parental age class appeared to increasingly overlap under warming temperatures (*Figure S29: top centre*).

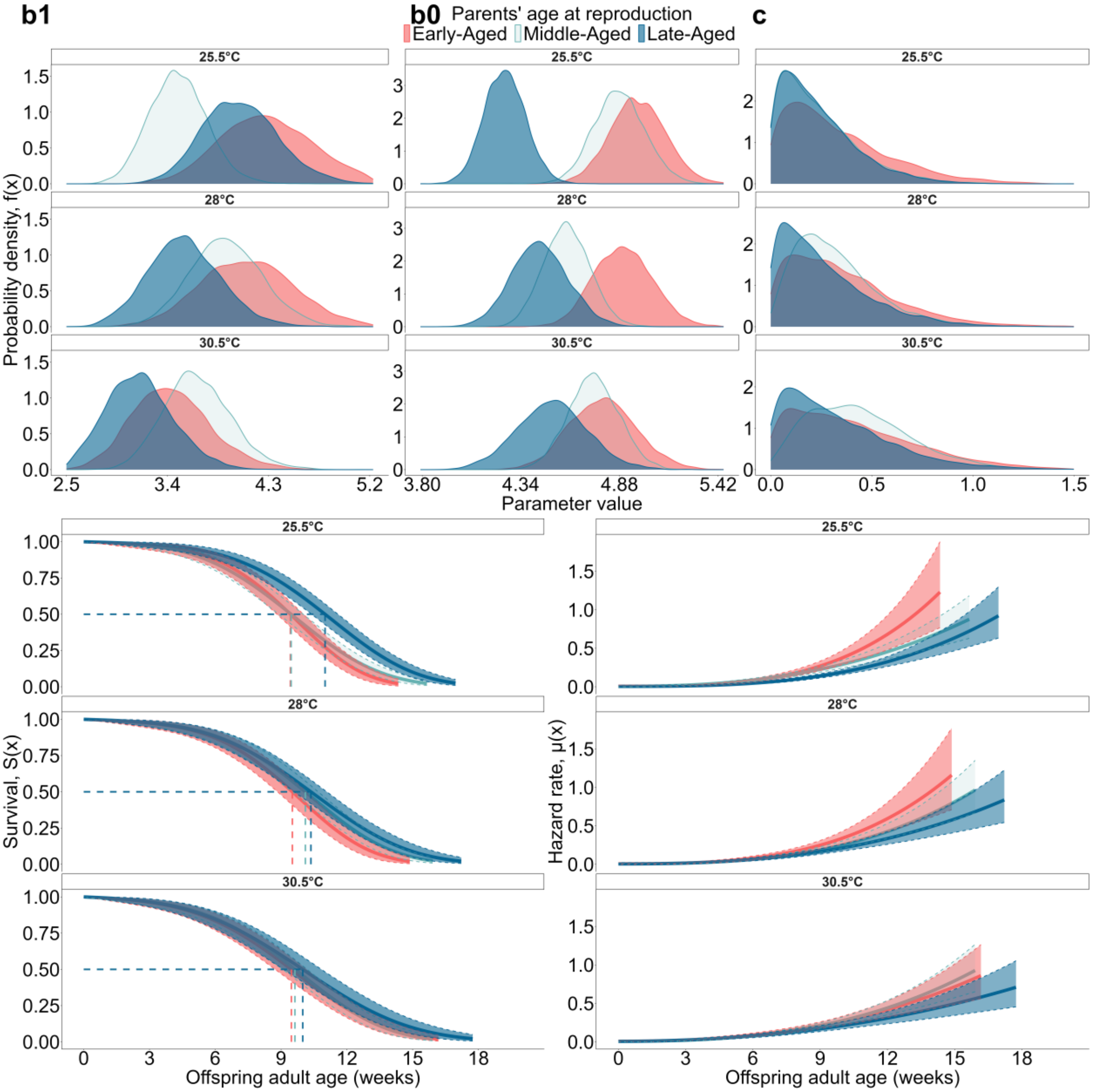

**Figure S29. The effect of the two-way interaction between the parents' adult age at reproduction and the parents' temperature treatment on the offspring's adult survival and mortality.** Shown along the top row are the full posterior distributions for the *b1*, *b0*, and *c* parameters. On the bottom row, the lines and ribbons reflect the model predictions ± 95% CI as estimated from the posterior draws (for offspring adult mortality and survival). Model estimates for the Weibull parameters and survival/mortality curves are faceted by parental temperature. We transformed the hazard to take the units 1/week. The colours describe the different parental age categories: red = “early-aged”, green = “middle-aged”, and blue = “late-aged”. N = 939 offspring from 77 parent pairs.

In *Table S28*, we do not report KLDC comparisons between the offspring sexes as to isolate our inferences to sex-specific parental age effects, not baseline differences in each sex’s mortality trajectories (although full tables are available to generate from our model scripts). The model including the interaction between offspring sex and parental temperature had the lowest DIC (DIC = -2628.82) of all models explaining variation in adult mortality. Thus, how parental age effects manifest could differ between the two offspring sexes. We suggest that support for this interaction is weak, as we have previously demonstrated in *brms*, on mean adult lifespan, that there was uncertain evidence for this interaction (*Table S26*). Rather, the fit of the adult mortality model could be benefitting from the added inclusion of offspring sex, not the interaction. Visual inspection of the mortality and survival trajectories revealed that the direction of the parental age effect was comparable between the offspring sexes (*Figure S30: bottom*). Here, offspring from late-aged parents exhibited reduced mortality over their adult lifespan, irrespective of offspring sex.

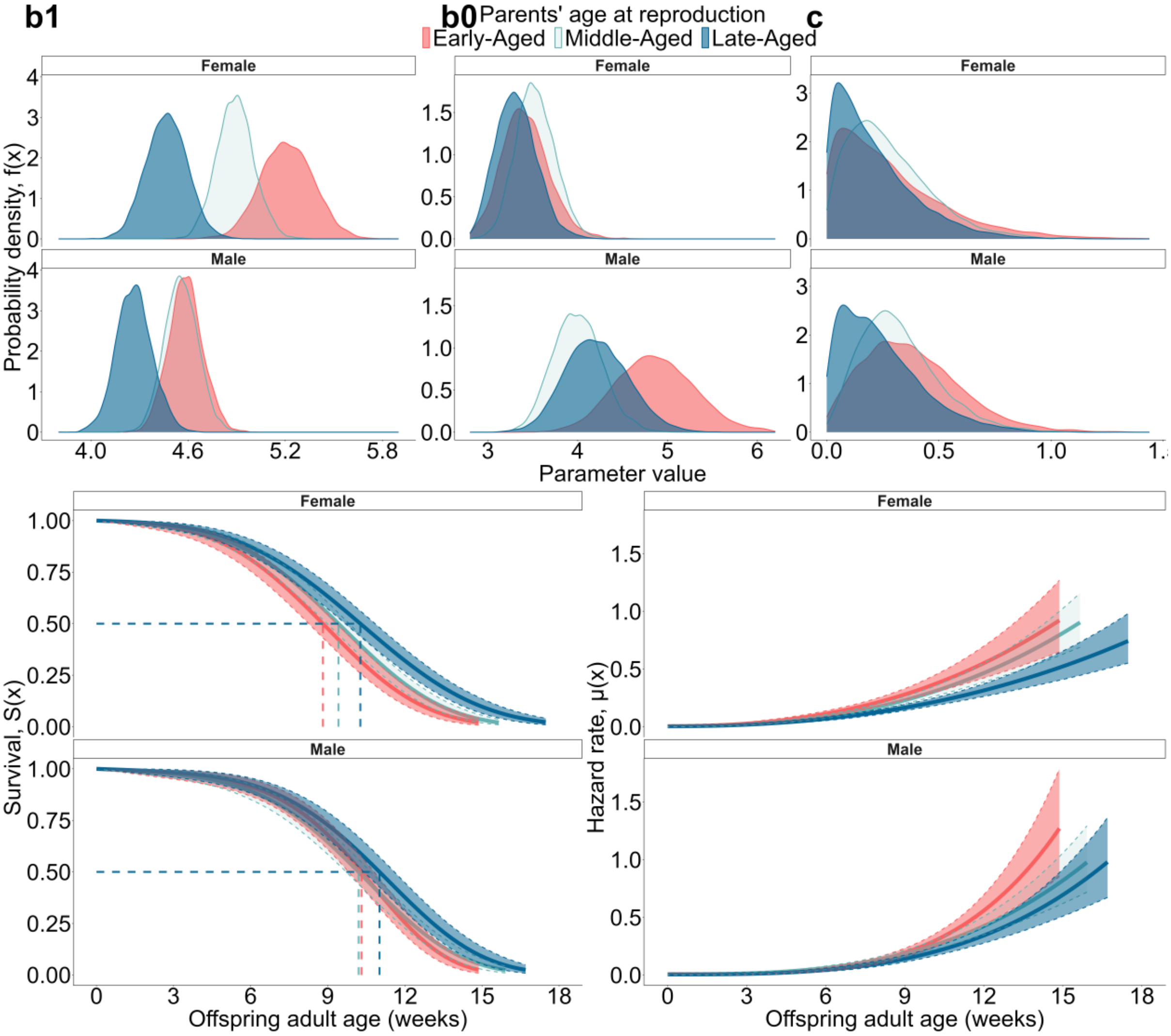

**Figure S30. The effect of the two-way interaction between the parents’ adult age at reproduction and the offspring’s sex on the offspring’s adult survival and mortality.** Shown along the top are the posterior distributions for the *b1*, *b0*, and *c* parameters. On the bottom panel, the lines and ribbons are the model predictions ± 95% CI as estimated from the posterior draws (for offspring adult mortality and survival). Model estimates for the Weibull parameters and survival/mortality curves are faceted by offspring sex. We transformed the hazard to take the units 1/week. The colours describe the different parental age categories: red = "early-aged", green = "middle-aged", and blue = "late-aged". N = 939 offspring from 77 parent pairs.

**Table S27. Comparison of the predictive performance of models explaining variation in offspring adult mortality. A) Ranked comparison of all mortality distributions available in BaSTA.** For each model, we provide their deviance information criterion (DIC), as well as their difference in DIC when compared to the best-performing model (the model with the lowest DIC). We selected the model with the lowest DIC, with the model said to be the best-performing when its ΔDIC < 3 from alternative candidate models. If models had comparable predictive performance, then we selected the simplest model (the model with the fewest number of parameters). We included no covariates in the model when testing which mortality distribution was most suitable given our data. **B) Comparison of the predictive performance of different prior distributions** (default, weakly informative, and moderately informative) fitted to models with each set of covariates: null model, parent's adult age at reproduction and the parents' temperature treatment. Moderately informative priors have the same mean, but ½ the SD of the weakly informative priors. For models with the *parental age* x *parental temperature* interaction, and the *parental age* x *offspring sex* interaction, we based our posterior inferences on the weakly informative prior structure. N = 939 offspring from 77 parent pairs.

| **Model selection based on predictive performance** | | | |
|---|---|---|---|
| ***Model*** | ***Shape*** | ***DIC*** | ***ΔDIC*** |
| ***a)*** **Comparison of BaSTA mortality distributions** | | | |
| **Weibull** | **Makeham** | **-2597.46** | **0** |
| Weibull | bathtub | -2597.12 | 0.34 |
| Weibull | simple | -2587.30 | 10.16 |
| Logistic | Makeham | -2454.29 | 143.17 |
| Logistic | simple | -2454.14 | 143.32 |
| Logistic | bathtub | -2444.76 | 152.70 |
| Gompertz | simple | -2451.72 | 145.74 |
| Gompertz | bathtub | -2447.06 | 150.40 |
| Gompertz | Makeham | -2446.74 | 150.72 |
| Exponential | simple | -1311.66 | 1285.8 |
| ***b)*** **Comparison of different prior structures on posterior predictive performance** *Using the Weibull-Makeham mortality distribution* | | | |
| **Model** | **Prior** | **DIC** | ***ΔDIC*** |
| Null | Weakly-informative | -2597.37 | 0 |
| Null | Default | -2597.46 | -0.26 |
| Null | Moderately-informative | -2595.32 | 1.89 |
| Parental age | Weakly-informative | -2617.67 | -20.47 |
| Parental age | Default | -2615.79 | 18.59 |
| Parental age | Moderately-informative | -2602.30 | -5.10 |
| Parental temperature | Weakly-informative | -2592.04 | 5.16 |
| Parental temperature | Default | -2590.73 | 6.47 |
| Parental temperature | Moderately-informative | -2578.17 | 19.03 |

**Table S28. Posterior parameter estimates for covariates explaining variation in offspring adult mortality parameters.** Here, we show the null model (with no covariates) (**1**), alongside the independent effects of the parents' adult age at reproduction (**2**), the parents' temperature treatment (**3**), and the combined effect of *parental age x offspring sex* (**5**). Shown for each parameter is the posterior mean ("*Estimate*"), the standard errors ("*SE*"), the 95% credible intervals ("*95% CI*"), along with the $\hat{R}$ and serial autocorrelation ("*S.A.corr*") estimates. We include the Kullback-Leibler discrepancy calibration ("*KLDC*") to estimate the distance between the posterior distributions among a pair of categorical covariates, with KLDC estimates > 0.8 being highlighted in bold. We also provide the DIC values to estimate which model (and set of variables) have the highest predictive performance in explaining variation in offspring adult mortality. For the two-way interaction between *parental age x parental temperature* (**4**), we only report the DIC value, as this model had substantially poorer predictive performance than the null model. N = 939 offspring from 77 parent pairs.

| Posterior mortality parameter estimates | | | | |
|---|---|---|---|---|
| ***Mortality Parameters*** | ***Estimate ± SE*** | ***CI (95%)*** | $\hat{R}$ | **S.A.corr** |
| ***1) Null Model (no covariates): DIC = -2597.37*** | | | | |
| **Makeham parameter (*c*)** | 0.22 ± 0.09 | 0.06 – 0.42 | 1.00 | 0.02 |
| **Scale parameter (*b1*)** | 4.64 ± 0.05 | 4.54 – 4.75 | 1.00 | 0.01 |
| **Shape parameter (b0)** | 3.74 ± 0.13 | 3.51– 4.00 | 1.00 | 0.01 |
| ***2) Parents' adult age at reproduction: DIC = -2617.67*** | | | | |
| **Makeham parameter (*c*)** | | | | |
| *Parental age: Early-Aged* | 0.34 ± 0.21 | 0.03 - 0.81 | 1.00 | 0.01 |
| *Parental age: Middle-Aged* | 0.30 ± 0.14 | 0.07 – 0.60 | 1.00 | 0.01 |
| *Parental age: Late-Aged* | 0.18 ± 0.14 | 0.01 – 0.53 | 1.00 | 0.02 |
| **Scale parameter (*b1*)** | | | | |
| *Parental age: Early-Aged* | 4.85 ± 0.10 | 4.65 – 5.05 | 1.00 | 0.00 |
| *Parental age: Middle-Aged* | 4.73 ± 0.07 | 4.57 – 4.88 | 1.00 | 0.00 |
| *Parental age: Late-Aged* | 4.37 ± 0.09 | 4.20 – 4.54 | 1.00 | 0.02 |
| **Shape parameter (*b0*)** | | | | |
| *Parental age: Early-Aged* | 4.05 ± 0.28 | 3.54 – 4.65 | 1.00 | 0.02 |
| *Parental age: Middle-Aged* | 3.76 ± 0.18 | 3.42– 4.14 | 1.00 | 0.01 |
| *Parental age: Late-Aged* | 3.68 ± 0.22 | 3.29 – 4.13 | 1.00 | 0.02 |
| ***KLDC pairwise comparisons*** | | | | |
| ***Parental age comparison*** | ***c*** | ***b1*** | ***b0*** | |
| *Early-aged vs. Late-aged* | 0.73 | **1.00** | **0.85** | |
| *Middle-aged vs. Early-aged* | 0.59 | **0.80** | **0.82** | |
| *Late-aged vs. Middle-aged* | 0.61 | **0.99** | 0.55 | |
| **3) Parents' temperature treatment: *DIC = -2592.04*** | | | | |
| **Makeham parameter (*c*)** | | | | |
| *Parental temperature: 25.5°*C | 0.16 ± 0.12 | 0.01 - 0.44 | 1.00 | 0.00 |
| *Parental temperature: 28.0°*C | 0.24 ± 0.14 | 0.04 – 0.56 | 1.00 | -0.02 |

| Posterior mortality parameter estimates | | | | |
|---|---|---|---|---|
| ***Mortality Parameters*** | ***Estimate ± SE*** | ***CI (95%)*** | $\widehat{R}$ | **S.A.corr** |
| *Parental temperature: 30.5°C* | 0.33 ± 0.19 | 0.03 – 0.77 | 1.00 | 0.00 |
| **Scale parameter (*b1*)** | | | | |
| *Parental temperature: 25.5°C* | 4.64 ± 0.08 | 4.49 – 4.80 | 1.00 | -0.01 |
| *Parental temperature: 28.0°C* | 4.60 ± 0.09 | 4.44 – 4.77 | 1.00 | -0.01 |
| *Parental temperature: 30.5°C* | 4.69 ± 0.10 | 4.48 – 4.89 | 1.00 | 0.01 |
| **Shape parameter (*b0*)** | | | | |
| *Parental temperature: 25.5°C* | 3.76 ± 0.19 | 3.40 – 4.16 | 1.00 | 0.00 |
| *Parental temperature: 28.0°C* | 3.91 ± 0.21 | 3.51 – 4.35 | 1.00 | 0.00 |
| *Parental temperature: 30.5°C* | 3.49 ± 0.21 | 3.10 – 3.92 | 1.00 | 0.02 |

| KLDC pairwise comparisons | | | |
|---|---|---|---|
| ***Parental temperature comparison*** | ***c*** | ***b1*** | ***b0*** |
| *25.5°C vs. 28.0°C* | 0.60 | 0.62 | 0.56 |
| *25.5°C vs. 30.5°C* | **0.81** | 0.59 | **0.81** |
| *28.0°C vs. 30.5°C* | 0.62 | 0.69 | **0.93** |

***4) Parents' temperature treatment x Parents' adult age at reproduction: DIC = -2595.79***

***5) Parents' adult age at reproduction x Offspring sex: DIC = -2628.82***

| ***Mortality Parameters*** | ***Estimate ± SE*** | ***CI (95%)*** | $\widehat{R}$ | **S.A.corr** |
|---|---|---|---|---|
| **Makeham parameter (*c*)** | | | | |
| ***Parental age: early aged,*** *Offspring sex: male* | 0.38 ± 0.22 | 0.04 – 0.87 | 1.00 | -0.01 |
| *Offspring sex: female* | 0.28 ± 0.24 | 0.01 - 0.91 | 1.00 | -0.01 |
| ***Parental age: middle-aged,*** *Offspring sex: male* | 0.32 ± 0.18 | 0.06 – 0.73 | 1.00 | 0.00 |
| *Offspring sex: female* | 0.28 ± 0.18 | 0.03 – 0.72 | 1.00 | -0.00 |
| ***Parental age: late-aged,*** *Offspring sex: male* | 0.24 ± 0.18 | 0.01 – 0.68 | 1.00 | 0.00 |
| *Offspring sex: female* | 0.21 ± 0.18 | 0.01 – 0.66 | 1.00 | 0.00 |
| **Scale parameter (*b1*)** | | | | |
| ***Parental age: early-aged,*** *Offspring sex: male* | 4.58 ± 0.10 | 4.38 – 4.80 | 1.00 | -0.01 |

| Posterior mortality parameter estimates | | | | |
|---|---|---|---|---|
| ***Mortality Parameters*** | ***Estimate ± SE*** | ***CI (95%)*** | $\hat{R}$ | **S.A.corr** |
| *Offspring sex: female* | 5.21 ± 0.16 | 4.90 – 5.53 | 1.00 | 0.01 |
| ***Parental age: middle-aged,*** *Offspring sex: male* | 4.56 ± 0.10 | 4.36 – 4.76 | 1.00 | 0.02 |
| *Offspring sex: female* | 4.89 ± 0.11 | 4.67 – 5.12 | 1.00 | -0.01 |
| ***Parental age: late-aged,*** *Offspring sex: male* | 4.27 ± 0.11 | 4.05 – 4.49 | 1.00 | -0.00 |
| *Offspring sex: female* | 4.47 ± 0.13 | 4.23 – 4.72 | 1.00 | -0.01 |
| **Shape parameter (*b0*)** | | | | |
| ***Parental age: early-aged,*** *Offspring sex: male* | 4.87 ± 0.44 | 4.05 – 5.76 | 1.00 | 0.00 |
| *Offspring sex: female* | 3.42 ± 0.27 | 2.92 – 3.98 | 1.00 | 0.00 |
| ***Parental age: middle-aged,*** *Offspring sex: male* | 4.00 ± 0.27 | 3.49 – 4.57 | 1.00 | 0.01 |
| *Offspring sex: female* | 3.53 ± 0.22 | 3.13 – 3.97 | 1.00 | 0.00 |
| ***Parental age: late-aged,*** *Offspring sex: male* | 4.21 ± 0.35 | 3.57 – 4.93 | 1.00 | 0.00 |
| *Offspring sex: female* | 3.31 ± 0.23 | 2.90 – 3.79 | 1.00 | 0.00 |

| ***KLDC pairwise comparisons → Only providing within-sex comparisons*** | | | |
|---|---|---|---|
| ***Offspring sex comparison*** | ***c*** | ***b1*** | ***b0*** |
| ***Offspring sex: female*** *Parental age: early vs. late* | 0.57 | **1.00** | 0.55 |
| *Parental age: early vs. middle* | 0.52 | **0.96** | 0.57 |
| *Parental age: middle vs. late* | 0.53 | **1.00** | 0.68 |
| ***Offspring sex: male*** *Parental age: early vs. late* | 0.61 | **0.99** | **0.88** |
| *Parental age: early vs. middle* | 0.54 | 0.51 | **0.97** |
| *Parental age: middle vs. late* | 0.54 | **0.99** | 0.63 |

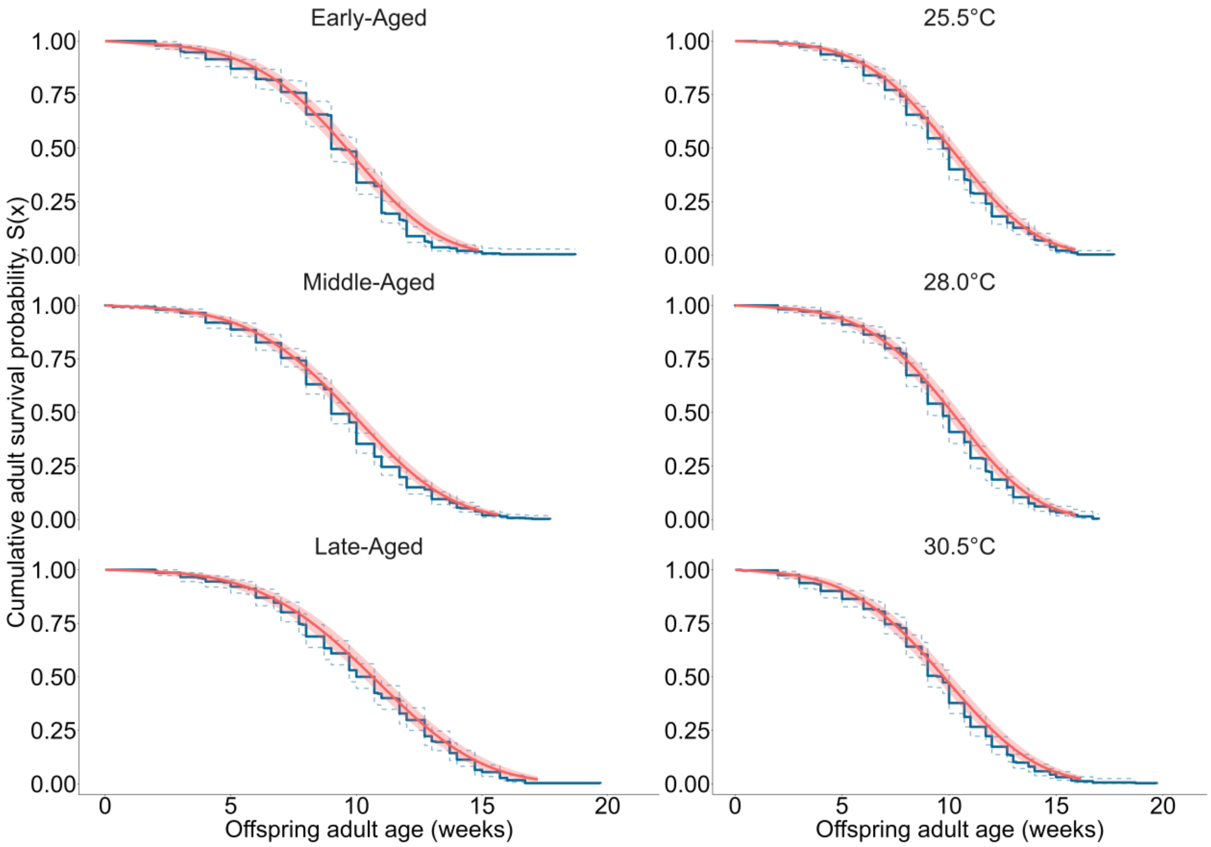

**Figure S31. Model fit diagnostic plots for Weibull-Makeham models explaining variation in offspring adult mortality.** Shown on the left are the adult survival curves for offspring from each parental age category, while, on the right, the offspring's adult survival curves for each parental temperature treatment are shown. The blue lines are the Kaplan-Meier plots of the offspring's observed survival probabilities, while the red lines represent the parametric Weibull-Makeham survival curves. N = 939 offspring from 77 parent pairs.

#### *References*

Kassambara, Alboukadel, Marcin Kosinski, and Przemyslaw Biecek. 2025. 'survminer: Drawing Survival Curves using ggplot2'. *CRAN: Contributed Packages (2016). R package version 0.5.1.*

Therneau, Terry M. 2024. 'A Package for Survival Analysis in R'. *R package version 3.8-3*. https://CRAN.R-project.org/package=survival.

Aalen, O.O. (1989) 'A linear regression model for the analysis of life times', *Statistics in Medicine*, 8(8), pp. 907–925. Available at: https://doi.org/10.1002/sim.4780080803.

Colchero, F., Jones, O.R. and Rebke, M. (2012) 'BaSTA: an R package for Bayesian estimation of age-specific survival from incomplete mark-recapture/recovery data with covariates', *Methods in Ecology and Evolution*, 3(3), pp. 466–470. Available at: https://doi.org/10.1111/j.2041-210X.2012.00186.x.

Donoughe, S. and Extavour, C.G. (2016) 'Embryonic development of the cricket Gryllus bimaculatus', *Developmental Biology*, 411(1), pp. 140–156. Available at: https://doi.org/10.1016/j.ydbio.2015.04.009.

Gelman, A., Simpson, D. and Betancourt, M. (2017) 'The Prior Can Often Only Be Understood in the Context of the Likelihood', *Entropy*, 19(10), p. 555. Available at: https://doi.org/10.3390/e19100555.

Gompertz, Benjamin. 1825. 'XXIV. On the Nature of the Function Expressive of the Law of Human Mortality, and on a New Mode of Determining the Value of Life Contingencies. In a Letter to Francis Baily, Esq. F. R. S. &c'. *Philosophical Transactions of the Royal Society of London*, no. 115 (December): 513–83. https://doi.org/10.1098/rstl.1825.0026.

Jennions, M.D. (2003) 'A survey of the statistical power of research in behavioral ecology and animal behavior', *Behavioral Ecology*, 14(3), pp. 438–445. Available at: https://doi.org/10.1093/beheco/14.3.438.

Lemoine, N.P. *et al.* (2016) 'Underappreciated problems of low replication in ecological field studies', *Ecology*, 97(10), pp. 2554–2561. Available at: https://doi.org/10.1002/ecy.1506.

Lemoine, N.P. (2019) 'Moving beyond noninformative priors: why and how to choose weakly informative priors in Bayesian analyses', *Oikos*, 128(7), pp. 912–928. Available at: https://doi.org/10.1111/oik.05985.

Møller, A. and Jennions, M.D. (2002) 'How much variance can be explained by ecologists and evolutionary biologists?', *Oecologia*, 132(4), pp. 492–500. Available at: https://doi.org/10.1007/s00442-002-0952-2.

McElreath, Richard. 'Statistical rethinking: A Bayesian course with examples in R and Stan.' Vol. 122. Boca Raton, FL: CRC press, 2016.

Pinder, J.E., Wiener, J.G. and Smith, M.H. (1978) 'The Weibull Distribution: A New Method of Summarizing Survivorship Data', *Ecology*, 59(1), pp. 175–179. Available at: https://doi.org/10.2307/1936645.

Siler, W. (1979) 'A Competing-Risk Model for Animal Mortality', *Ecology*, 60(4), pp. 750–757. Available at: https://doi.org/10.2307/1936612.

Sivula, T. *et al.* (2020) 'Uncertainty in Bayesian Leave-One-Out Cross-Validation Based Model Comparison'. arXiv. Available at: https://doi.org/10.48550/ARXIV.2008.10296.

Winter, B. and Bürkner, P. (2021) 'Poisson regression for linguists: A tutorial introduction to modelling count data with brms', *Language and Linguistics Compass*, 15(11). Available at: https://doi.org/10.1111/lnc3.12439.

Winter, Bodo, and Paul-Christian Bürkner. 2021. 'Poisson Regression for Linguists: A Tutorial Introduction to Modelling Count Data with Brms'. *Language and Linguistics Compass* 15 (11). e12439. https://doi.org/10.1111/lnc3.12439.
